# The Neuronal Primary Cilium is a Key Regulator of Homeostatic Plasticity

**DOI:** 10.1101/2025.08.27.672341

**Authors:** Emma Dyke, Sofia Puvogel, Astrid Oudakker, Ronald Roepman, Brooke Latour, Nael Nadif Kasri

## Abstract

The capacity of neurons to maintain stable activity levels through homeostatic plasticity is essential for proper brain function. Primary cilia, which are non-motile, antenna-like organelles projecting from the surface of most vertebrate cells, serve as key hubs for signal transduction, playing crucial roles in tissue development and cellular homeostasis. In this study, we identify a previously unrecognised role for primary cilia in mediating neuronal homeostatic plasticity using human induced pluripotent stem cell-derived neurons. We show that neuronal cilia exhibit dynamic, bidirectional changes in volume in response to alterations in network activity: elongating during chronic activity suppression and shortening after increased activity. To assess the functional relevance of this ciliary plasticity, we modelled ciliary dysfunction in neurons carrying homozygous loss-of-function mutations in genes associated with neuronal ciliopathies, including *NPHP1* and *CEP290*. Mutations affecting ciliary function either increased ciliary length or led to ciliary loss, and these mutant neurons exhibited severe impairments in homeostatic regulation across multiple domains—morphological, functional, and transcriptional. Specifically, *NPHP1* and *CEP290* deficient neurons failed to adapt synaptic strength, intrinsic excitability, and ciliary morphology in response to prolonged activity suppression. They also displayed dysregulated baseline network activity, and exhibited blunted gene expression changes. Together, these findings establish the primary cilium as a critical regulator of homeostatic plasticity in human neurons and provide a new framework through which to examine neurodevelopmental and neuropsychiatric disorders linked to ciliary dysfunction.

**Key highlights:**

- Primary cilia adapt bidirectionally to neuronal activity by dynamically changing volume, expanding during suppressed activity and contracting with increased neuronal activity.
- Disruptions in ciliopathy-associated genes perturb homeostatic plasticity, impairing structural, functional, and transcriptional responses to activity changes.
- Primary cilia are essential for neuronal network development.

## Introduction

Neural circuits maintain their functional stability through homeostatic plasticity, a set of mechanisms that balance intrinsic excitability and synaptic strength to ensure stable and efficient information processing in the face of ongoing changes in network activity (Tien and Kerschensteiner, 2018, Turrigiano, 2012, Davis, 2013). This plasticity operates at multiple levels, encompassing structural and functional adaptations of axons, dendrites, and synapses that underlie both synaptic and intrinsic forms of homeostatic regulation (Turrigiano *et al*. 1998).

The primary cilium is a solitary, antenna-like structure that protrudes from the surface of most mammalian cell types, including those of the central nervous system, such as neurons, astrocytes, and oligodendrocyte precursor cells (Buchanan *et al*. 2022). Acting as a sensory and signalling hub, it detects and integrates extracellular signals to influence a broad range of cellular processes. While ciliary signalling is well recognised for its roles in neurodevelopment, such as neuronal migration, patterning, and differentiation (<u>Higginbotham *et al*. 2012</u>, <u>Higginbotham *et al*. 2013</u>, <u>Hasenpusch-Theil and Theil, 2021, Park *et al*. 2019, Karalis *et al*. 2022, Liu *et al*. 2021</u>), its role in post-mitotic neurons is less well understood, particularly in the context of neural plasticity.

Evidence from both human and rodent studies demonstrates that primary cilia persist in mature neurons and astrocytes (<u>Arellano *et al*. 2011</u>; <u>Ott *et al*. 2024</u>; Wu et al. 2024). Disruptions in ciliary function can alter neuronal connectivity and neuronal excitability (Guo *et al.,* 2017; Tereshko *et al*. 2021), and recent studies have shown that acute perturbation of cilia influences spontaneous activity and synaptic strength (Tereshko *et al*. 2021). Moreover, the primary cilium can form functional synapses *in vivo* and *in vitro* and mediate serotonergic signalling that impacts chromatin accessibility, presumably altering the transcriptome (Sheu *et al*. 2022). Finally, in primary human tissue single primary cilium can make numerous contacts with different cellular domains, including both excitatory and inhibitory dendrites and axons, and persist in proximity to tripartite synapses involving axon, dendrite and astrocytes (Wu et al. 2024). These observations suggest that primary cilia may play a role in maintaining neuronal homeostasis, though the underlying mechanisms remain largely undefined.

Homeostatic plasticity, the negative feedback counterpart to Hebbian plasticity, is essential for stabilising neural networks, enabling reliable information processing and supporting learning and memory. Its disruption is implicated in neurodevelopmental and neuropsychiatric conditions, including autism spectrum disorder (ASD) and intellectual disability (<u>Bülow *et al*. 2019, Ellingford *et al*. 2021, Tatavarty *et al*. 2020</u>; Benevento *et al*., 2016), and is considered a possible convergent mechanism for ASD risk gene function (<u>Genç *et al*. 2020</u>). Given the growing evidence that ciliary function is important to neuronal homeostasis, it is plausible that they also participate in homeostatic plasticity.

Here, we investigated the role of primary cilia in homeostatic plasticity using iPSC-derived neurons with mutations in Joubert syndrome-associated genes *NPHP1* and *CEP290*. We found that primary cilia exhibit activity-dependent morphological changes in control neurons, and that disruption of these ciliary proteins alters baseline network activity and impairs homeostatic adaptations. Using a well-established paradigm to induce homeostatic plasticity in human iPSC-derived neurons (Yuan *et al*. 2023), we found that ciliary mutants displayed blunted homeostatic responses, suggesting that the primary cilium plays a previously unrecognised role in regulating neuronal network stability.

## Results

### Neuronal ciliation and ciliary volume increase over development and ciliary morphology is altered by network activity perturbation

To characterise the dynamics of primary cilia in developing glutamatergic neurons, we differentiated control iPSCs using an *Ngn2* overexpression protocol (<u>Zhang *et al*. 2013</u>; <u>Mossink *et al*. 2021</u>). To assess ciliation *in vitro*, we tracked ciliary development over time by monitoring the expression of the ciliary-specific proteins ARL13B (ADP-Ribosylation Factor-Like GTPase 13B) and ADCY3 (Adenylate Cyclase 3) in MAP2 expressing neurons (Figure 1a, b). Previous studies have shown that ARL13B and ADCY3 exhibit temporal changes during neuronal maturation, with ARL13B levels decreasing as ADCY3 increases (<u>Kasahara</u> *<u>et al</u>*<u>. 2014; Sterpka and Chen 2018</u>). Consistent with these findings, we observed a progressive increase in the percentage of ciliated neurons over time (Figure 1c), accompanied by a gradual reduction in the proportion of ARL13B-positive cilia (Figure 1d). We also noted a significant increase in ciliary volume between day 7 and day 35 of differentiation (Figure 1e). Recent work has shown that the primary cilium can form contacts with axons both *in vivo* and *in vitro* in mouse hippocampal neurons (<u>Sheu *et al*. 2022</u>). Similarly, we observed that a subset of cilia established contacts with axons in our model (Supplementary Figure 1a). Together, these data support the use of iPSC-derived neurons to recapitulate key developmental features of neuronal cilia, including dynamic changes in structural maturation over time (<u>Kasahara *et al*. 2014</u>, <u>Ott *et al*. 2024</u>).

**Figure 1:**
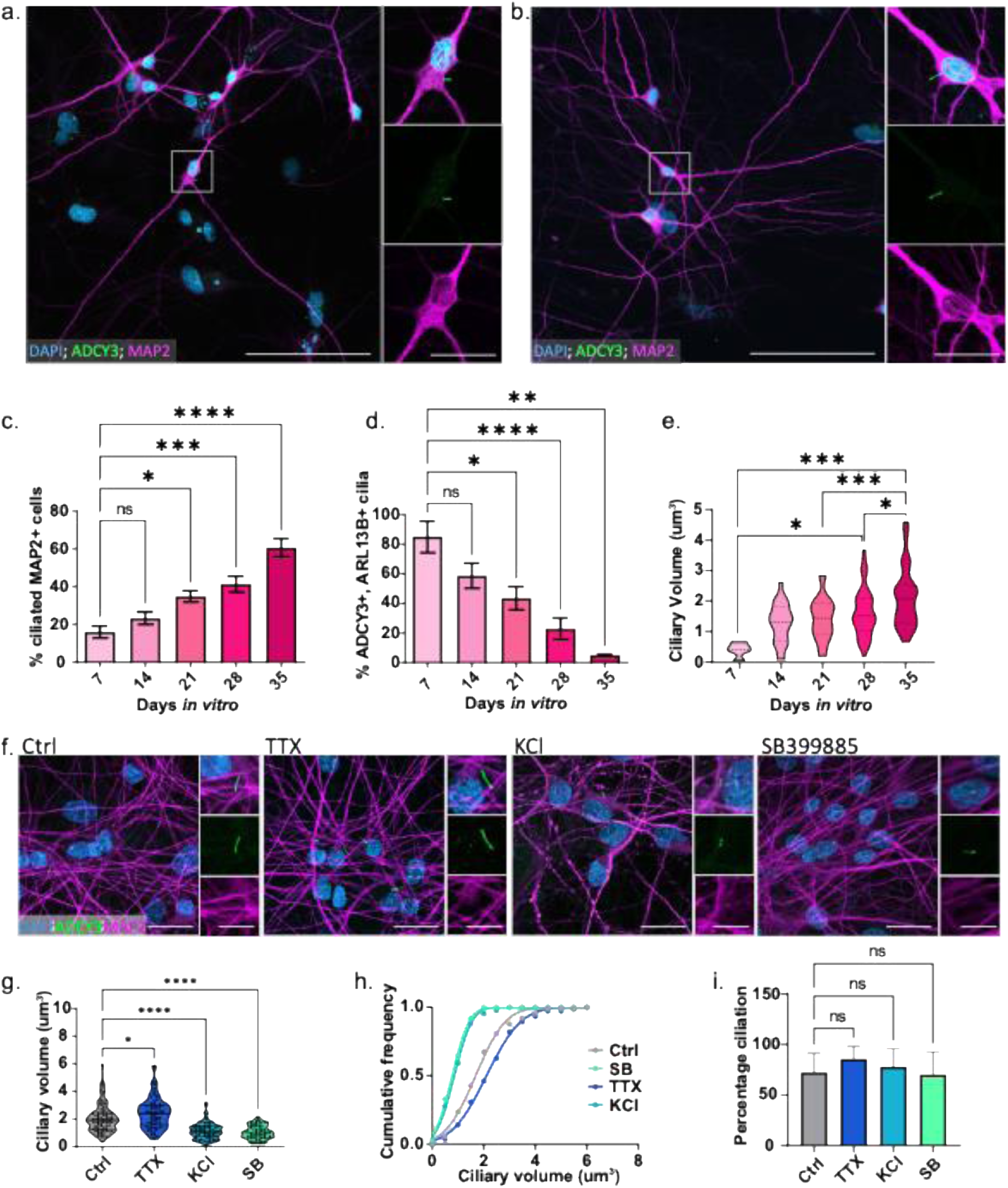
Cilia of hiPSC-derived neurons show temporal differences and bidirectional changes in response to changes in activity. Representative immunofluorescence of (a) DIV14 and (b) DIV28 neurons respectively, neurons are marked with MAP2 (magenta) and cilia with ADCY3 (green), the nucleus is shown with DAPI (blue), scale bar is 100 microns. Inset shows a zoom in on the primary cilium, scale bar is 20 microns. (c) Quantification of total percentage of ciliation (ARL13B and/or ADCY3-stained cilia as a percentage of MAP2 positive nuclei). N = independent experiments/coverslips assessed. N=4/6-20. (d) Quantification of percentage of cilia that are co-stained for both ARL13B and AC3 over time N=4/6-20. (e) Quantification of the volume of ADCY3-positive cilia over time. N= independent experiments/neurons assessed. N=4/14-62. Data represent means ± SEM. One-way ANOVA with Dunnett’s post-hoc correction (c-d) or Turkey’s correction (e) ns: not significant, *p < 0.05, **p < 0.01, ***p < 0.005, ****p < 0.0001. (f) Representative immunofluorescence staining of neurons treated at DIV35 with vehicle (ctrl) ddH_2_O for 24h, 1 μM TTX for 24h, 10 mM KCl for 3h, or 1 μM SB-399885 for 24h. MAP2 (magenta) and ADCY3 (green). Scale bars = 25 microns. Insets showing magnification of the cilium have scale bars of 10 microns. (g,h) shows the ciliary volume measured following treatment. N = independent experiments/neurons N=3/73-90 and (i) shows percentage ciliation per coverslip. (N=3/5). Analysis conducted with One-way ANOVA with Dunnett’s post-hoc correction, ns: not significant, *p < 0.05, ****p < 0.0001.

Neuronal compartments, such as dendritic spines, exhibit high structural plasticity in response to changes in activity, which is critical for maintaining homeostasis (<u>Matsuzaki *et al*. 2004</u>; <u>Zhou *et al*. 2004</u>). This dynamic remodelling also occurs in the cilium where ciliary composition and signalling are modified by various environmental cues (Corbit *et* al. 2005, Wang *et al*. 2009, Paillard *et* al. 2025). To determine whether neuronal activity directly modulates ciliary morphology, we manipulated network activity *in vitro* using potassium chloride (KCl, 10 mM) to increase activity, the sodium channel blocker tetrodotoxin (TTX) to suppress activity, and the serotonin receptor 6 (5-HT6) antagonist SB-399885 as a positive control, which has been reported to shorten cilia in striatal neurons (<u>Brodsky *et al*. 2017</u>). We found no differences in the volume of neuronal cilia between days *in vitro* (DIV) 35 and DIV 42 so we proceeded with the DIV 35 neurons. SB-399885 application for 24 h significantly reduced ciliary volume (Figure 1f–h). Similarly, KCl-induced hyperactivity for 3 h led to a reduction in ciliary size, whereas TTX treatment for 24 h, which silenced network activity (Supplementary Figure 1b– i), resulted in a significant increase in ciliary volume (Figure 1f–h). None of the treatments altered the proportion of ciliated neurons, which remained stable at 60–70% (Figure 1i). These results demonstrate that neuronal cilia exhibit bidirectional structural plasticity in response to network activity changes, with increased activity reducing and decreased activity enlarging ciliary volume.

### Increases in ciliary volume follow axon initial segment and synaptic remodelling during homeostatic plasticity

It is well established that prolonged blockage of network activity with TTX induces homeostatic plasticity (Turrigiano *et al*. 1998). To determine the temporal relationship between ciliary remodelling and other morphological adaptations during homeostatic plasticity, we monitored axon initial segment (AIS) length, synaptic GluA2 incorporation, and ciliary volume following chronic activity suppression with TTX, known to induce homeostatic plasticity. AIS length, measured using Ankyrin G, exhibited a rapid and robust elongation detectable within 30 minutes of TTX application and stabilised by 2 hours (Figure 2a, d). This aligns with previous reports of AIS plasticity during network silencing (<u>Grubb and Burrone, 2010</u>).

**Figure 2:**
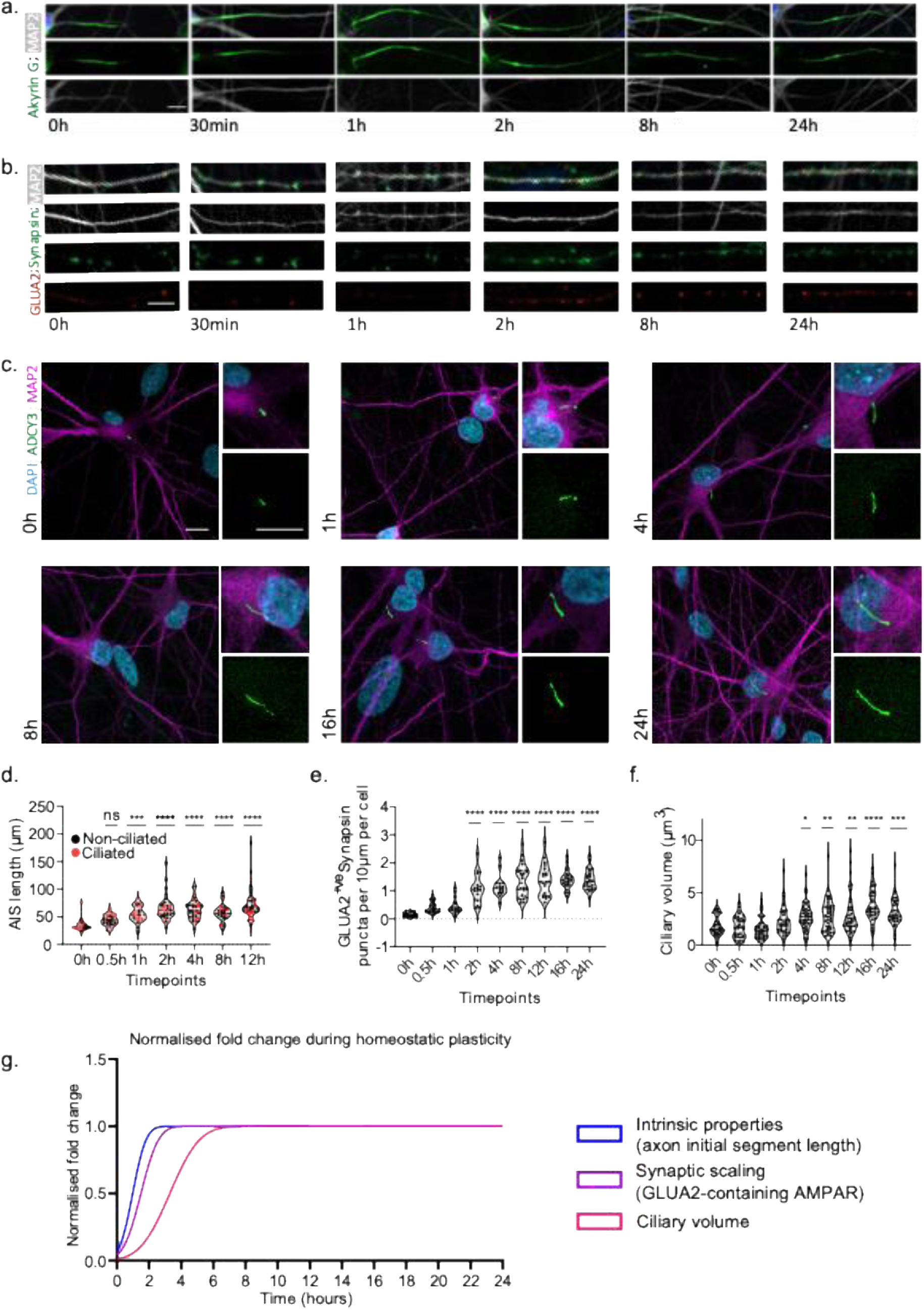
Synaptic scaling, intrinsic properties, and ciliary morphology change sequentially following sustained activity blockage. (a) Representative immunofluorescence staining of the AIS with AnkyrinG (green) and MAP2 (white) over time following TTX treatment of control neurons at DIV35 (b) shows MAP2 (white), Synapsin1 (green), and surface GluA2 (red), and (c) shows representative images of ciliary volume over time with MAP2 (purple) indicating neurons and ADCY3 (green) staining the primary cilium. Quantifications of these are shown in (d, e, and f) which was done across n=8 coverslips from N=3 independent experiments. 25-38 neurons per condition for (d), 16-26 for (e) and 32-42 for (f). (g) shows the normalised fold change in increase in number of GluA2-containing synapses, AIS or ciliary length normalised with respect to their largest value and scaled to 1. Scale bars = 10μm. One-way ANOVA with Dunnett’s multiple comparisons, ns = not significant,*p < 0.05, **p < 0.01, ***p<0.0005 ****p < 0.0001

Homeostatic synaptic scaling was evident as a progressive increase in the synaptic incorporation of the GluA2 AMPA receptor subunit. Co-localization analysis with Synapsin-1 revealed significant GluA2 enrichment at synapses after 2 hours of activity suppression, reaching a plateau by 12 hours (Figure 2b, e), consistent with previous reports (<u>Turrigiano *et*</u> *<u>al</u>*<u>. 1998</u>).

Ciliary volume, however, exhibited a delayed response. No significant changes were observed during the initial 2 hours of TTX treatment, but a marked increase emerged at 4 hours and persisted thereafter (Figure 2c, f). A normalised timeline of these morphological changes revealed that AIS and synaptic remodelling preceded ciliary enlargement (Figure 2g). These findings place ciliary remodelling downstream of early homeostatic adaptations, raising the possibility that increased ciliary volume contributes to the maintenance or consolidation of homeostatic plasticity rather than its initiation. Similar to dendritic spines, which undergo structural remodelling as part of plasticity processes, primary cilia may represent an active site of signalling adaptation during sustained activity perturbations.

### Loss of ciliary proteins alters ciliary morphology and neuronal network activity

To investigate whether ciliary integrity is required for homeostatic plasticity, we generated isogenic iPSC-derived neuronal lines lacking the Joubert syndrome–associated genes *CEP290* or *NPHP1* using CRISPR/Cas9 (*CEP290^−/-^* or *NPHP1^−/-^*; Supplementary Figure 2) (Dyke *et al*. 2023). Both proteins localise to the ciliary transition zone, where they play an important role in ciliary gating, preventing the passive diffusion of proteins in and out of the cilium. Consistent with previous observations (<u>Kobayashi *et al*. 2024</u>) (<u>Srivastava *et al*. 2017</u>), *NPHP1* deficiency was associated with elongated cilia, whereas *CEP290* deficiency resulted in an absence of detectable cilia. Neither *CEP290^−/-^* nor *NPHP1^−/-^* neurons exhibited deficits in differentiation, dendritic morphology, or synapse density compared to isogenic controls (Supplementary Figure 4). ADCY3 staining revealed a significant increase in ciliary length in *NPHP1^−/-^*neurons, accompanied by a reduced proportion of ciliated cells (Figure 3a-c). In contrast, *CEP290^−/-^* neurons lacked ADCY3 and ARL13B-positive cilia altogether (Supplementary Figure. 3).

**Figure 3:**
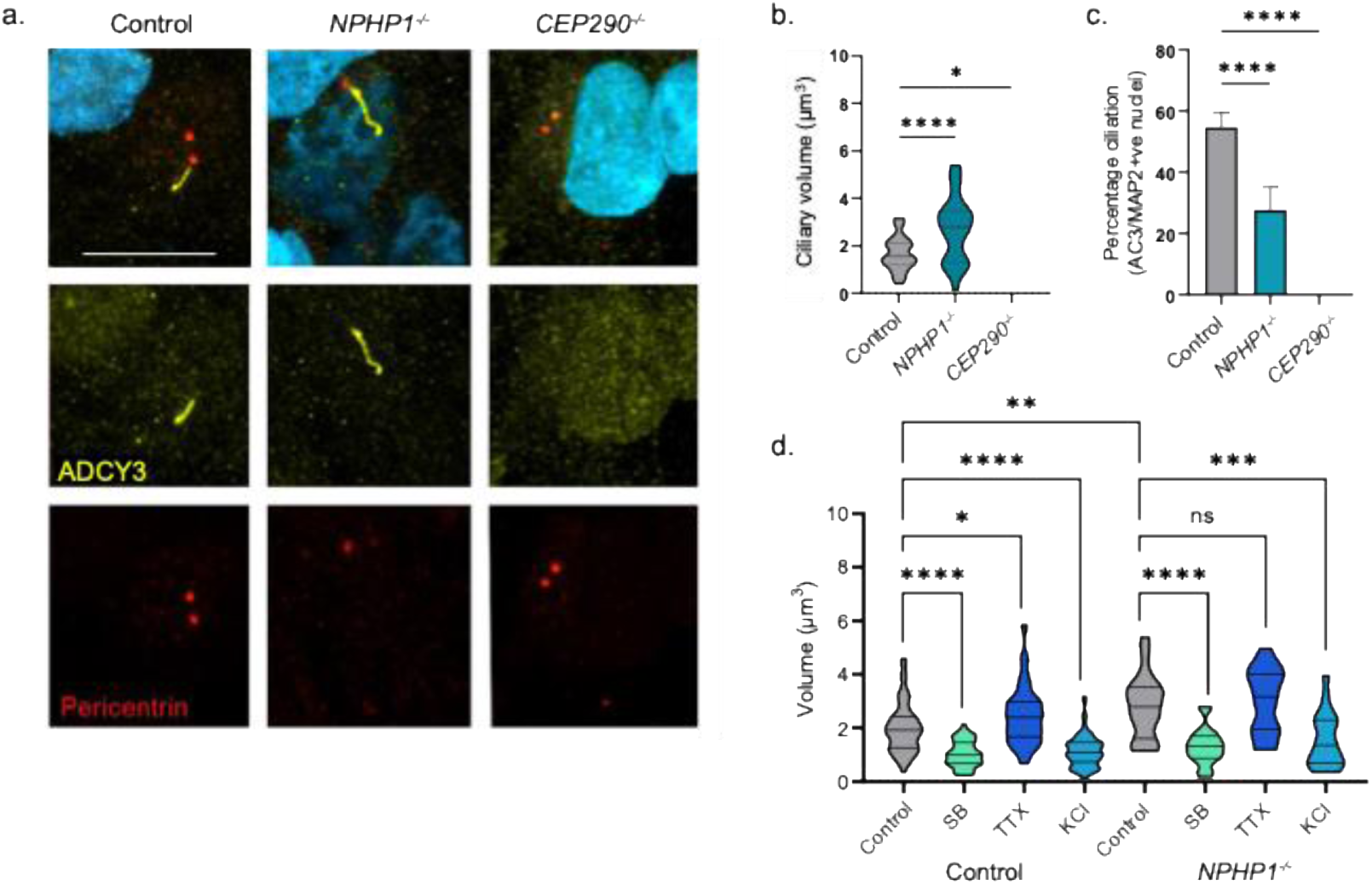
Characterisation of hiPSC-derived neurons generated using CRISPR-Cas9 with homozygous truncating mutations in *NPHP1* and *CEP290*. (a) Immunofluorescent staining of the cilia in green (ADCY3) of neurons generated from the CRISPR lines; pericentrin (red) is used to mark the centrosomes. Scale bar = 10μm. (b) shows the quantification of ciliary volume from the cell lines N = independent experiments/neurons, N=3/29-48, and (c) indicates the percentage ciliation. N = independent experiments/coverslips N=3/6. One-way ANOVA with Dunnett’s multiple comparisons, *p < 0.05, ****p < 0.0001. (d) shows the ciliary volume following application of vehicle (ddH_2_O), SB-399885, TTX, or KCl in the control and *NPHP1*^−/-^ neurons (DIV35). N = independent experiments/neurons, N=3/20-89. One-way ANOVA ANOVA with Turkey’s multiple comparisons, ns=not significant, *p < 0.05, **p<0.01, ***p<0.0005 ****p < 0.0001.

We next assessed whether these structural alterations impaired the bidirectional ciliary remodelling observed during activity perturbation. In *NPHP1^−/-^* neurons, KCl-induced hyperactivity reduced ciliary volume to control levels, but TTX-mediated activity suppression failed to elicit further ciliary enlargement (Fig. 3e), potentially because of an occlusion effect. Notably, the serotonin receptor antagonist SB-399885 reduced ciliary volume in *NPHP1^−/-^* neurons similarly to controls, indicating that this pathway remains intact. As expected, *CEP290^−/-^*neurons remained unciliated under all conditions (Supplementary Figure 5).

To evaluate the functional consequences of *NPHP1* and *CEP290* loss, we cultured iPSC-derived neurons on micro-electrode arrays, and monitored the network properties during development (DIV14-35). As previously described (Mossink *et al*. 2021), maturing neuronal networks exhibit synchronous bursting across electrodes, with bursts becoming shorter and more temporally confined as connectivity increases. Neuronal networks of all genotypes (control, *NPHP1^−/-^* and *CEP290^−/-^*) displayed this typical developmental trajectory, with progressive increases in network burst frequency (BF), spikes per burst, and the proportion of activity occurring within network bursts over time (Figure 4a-c). Interestingly, both *NPHP1^−/-^* and *CEP290^−/-^* networks deviated from controls by DIV28, exhibiting elevated network burst frequency but reduced spikes per burst compared to controls (Figure 4a-l). Detailed network analysis further revealed enhanced burst synchrony yet fewer spikes contributing to network bursts (Figure. 4d, e). Despite these changes in network dynamics, the mean firing rate (MFR) remained comparable across all lines, suggesting that ciliary dysfunction does not induce hyperactivity but rather reorganises the temporal structure of activity(Figure 4f, m).

**Figure 4:**
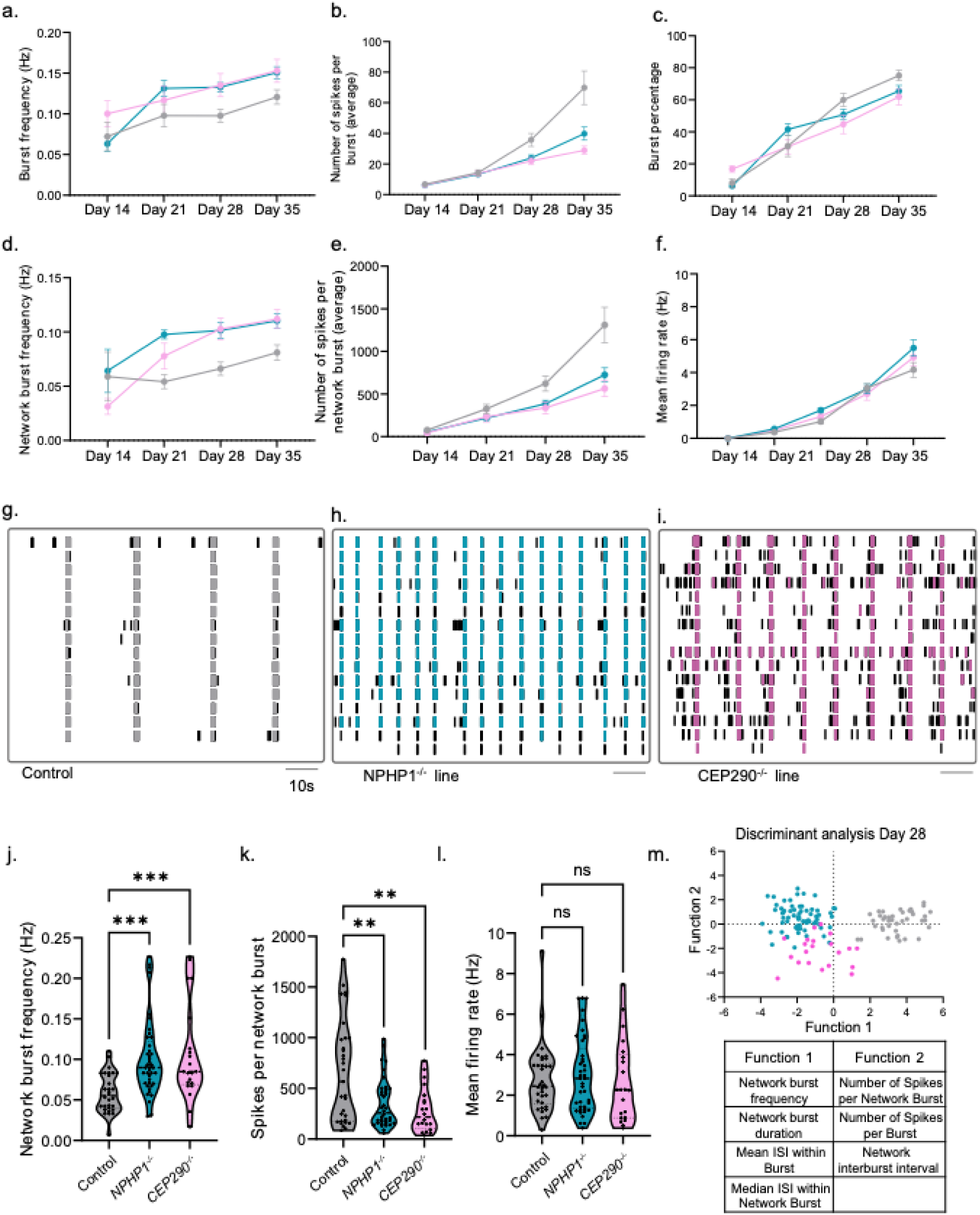
Loss of ciliary proteins alters development of neuronal networks and affects basal network activity on MEAs. (a-f) Developmental trajectory of control, *NPHP1^−/-^,* and *CEP290^−/-^* neurons on MEAs between DIV14-DIV35 across N=4 independent replicates, n= 30-42 wells. (g-i) Representative rasterplots from recordings of cell lines at DIV28. (j-l) results of an analysis of DIV28 recordings n=21-41 wells, one-way ANOVA with Dunnett’s multiple comparisons, ns=not significant, **p < 0.01, ***p < 0.0005 (m) shows the results from discriminant analysis of these cell lines with each point representing one well from DIV28 recordings.

Principal component analysis (PCA), encompassing 24 MEA parameters (Supplementary Table 1), further demonstrated that *NPHP1^−/-^* and *CEP290^−/-^*networks clustered together and apart from controls, with network burst frequency and network burst duration driving separation from controls, and spikes per network burst and network interburst interval differentiating between the two mutants (Figure 4m).Together, these data indicate that loss of two distinct Joubert syndrome–associated ciliary proteins, despite producing opposite effects on ciliary structure, converges on a shared phenotype of altered network organization independent of gross synaptic or dendritic abnormalities (Supplementary Figure 4).

### Loss of ciliary proteins alters neuronal transcriptional profiles

To better understand the molecular mechanisms underlying the neuronal network changes, we performed bulk RNA sequencing neurons. Principal component analysis (PCA) revealed that genotype accounted for the largest variation, with both *NPHP1*^−/-^ and *CEP290*^−/-^ neurons clustering separately from controls. Notably, the transcriptional profiles of the ciliary mutants showed minimal variation (Figure 5a), and this clustering pattern mirrored that observed in network activity PCA, where ciliary mutants clustered more closely together than the control.

**Figure 5:**
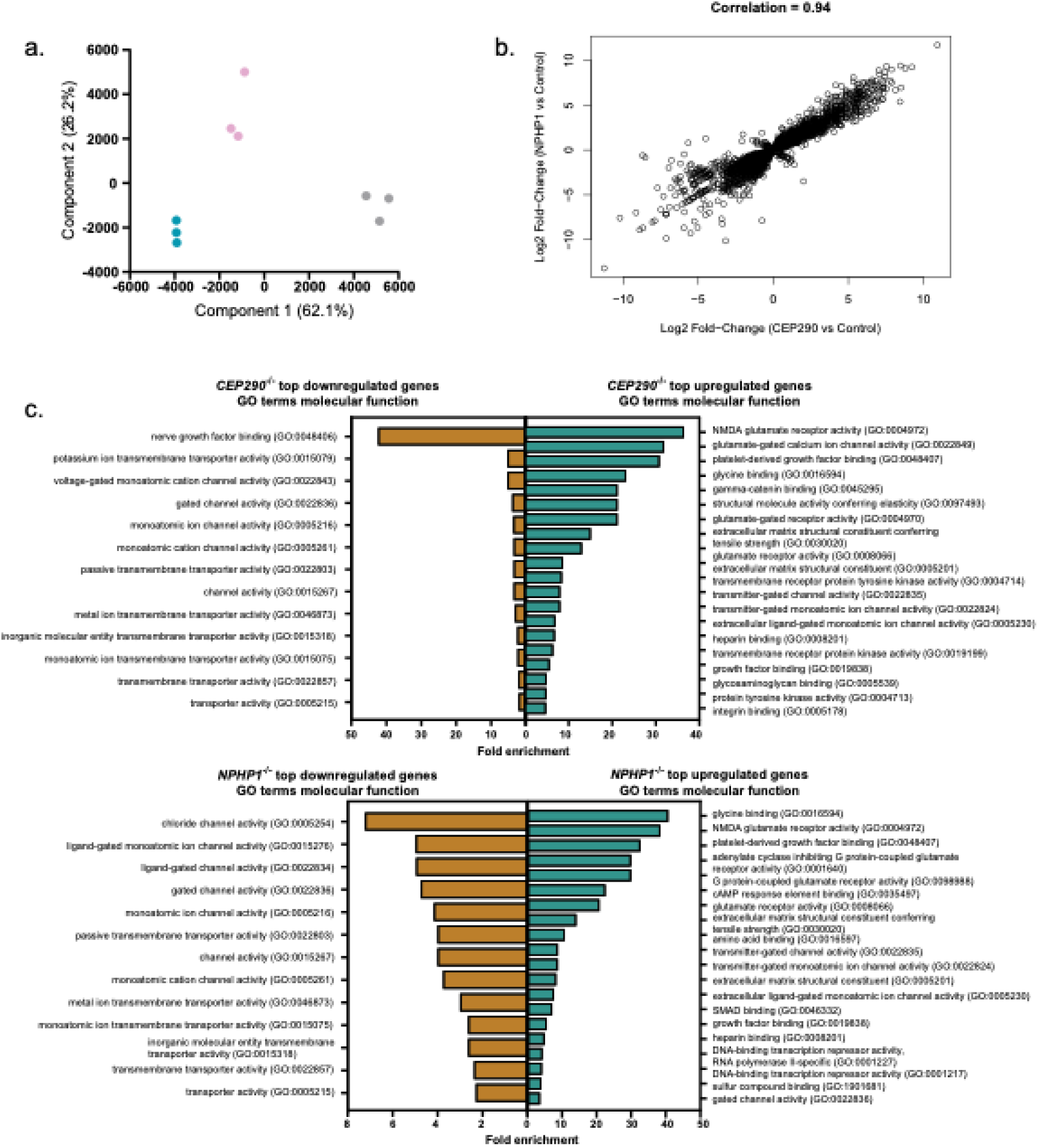
Transcriptome analysis of ciliary mutant neurons. (a) PCA of ciliary mutant lines at DIV36. (b) correlation of differentially expressed genes from *NPHP1^−/-^* and *CEP290^−/-^* with respect to the control line with Pearson correlation coefficient above. (c-d) gene ontology (GO) terms of the top 250 most upregulated and downregulated genes (by p value) for the *CEP290^−/-^*and *NPHP1^−/-^* neurons.

Differential expression analysis (using a |LogFC| >0.58 and a FDR p value of <0.05) revealed that 991 genes were significantly upregulated and 1029 genes were downregulated in the *CEP290^−/-^*neurons, while 1703 genes were significantly upregulated and 1648 genes were downregulated in the *NPHP1^−/-^* neurons. Of these, 731 genes were upregulated in both genotypes and 755 genes were downregulated in both genotypes (Supplementary figure 6a-d). The datasets are highly correlated, with a Pearson coefficient of 0.937 (Figure 5b), further indicating that despite the neurons having differential ciliary morphology, dysfunction of the primary cilium results in a convergent molecular phenotype. We focused on the most significantly differentially expressed genes and performed gene ontology (GO) analysis of the top 250 most significantly upregulated and downregulated genes (Figure 5b, Supplementary Table 4). Genes affecting receptor activity were upregulated, whereas downregulated genes were often associated with the activity of transmembrane transporters and ion channels (Figure 5c, d). There was an upregulation of glutamatergic subunit expression, including N-Methyl D-Aspartate (NMDA) receptor subunits *GRIN1, GRIN2B,* and *GRIN3A*, and α-amino-3-hydroxy-5-methyl-4-isoxazolepropionic acid (AMPA) receptor subunits *GRIA2* and *GRIA3* in both genotypes. This could, in part, explain the differences in network activity observed on the MEAs, as an increase in AMPA and NMDA receptors alter synaptic transmission, leading to an increase in excitability at a single cell level, and a positive feedback loop increasing spike rates and network synchronicity (<u>Turrigiano *et al*. 1998</u>).

In addition to genes related to glutamatergic transmission, we noted dysregulation of the expression of several ciliary genes, indicating loss of *NPHP1* or *CEP290* alters ciliary composition (Supplementary Table 5). Notably, although we do not observe ciliary structures in the *CEP290*^−/-^ lines, there is still considerable expression of ciliary genes. Overall, our transcriptional analysis reveals that despite distinct ciliary morphological defects, loss of either *NPHP1* or *CEP290* leads to a convergent molecular phenotype characterised by dysregulation of synaptic and ciliary gene networks, providing a potential mechanistic link to the observed alterations in neuronal network activity.

### Loss of ciliary proteins alters the homeostatic plasticity response

Given that activity-dependent ciliary remodelling was absent in both *NPHP1^−/-^* and *CEP290^−/-^* neurons (Figure 3d), we next examined whether this structural deficit impacts the expression of homeostatic plasticity. Neuronal networks were cultured on MEAs and subjected to TTX for 24 h, a time period at which AIS elongation, GluA2 upregulation, and ciliary enlargement are consistently observed in control neurons (Figure 2). Following TTX washout with conditioned medium, network activity was monitored for an additional 24 h (Figure 6a). In control networks, chronic activity suppression led to a robust homeostatic response, characterised by a sustained increase in network burst rate (NBR) following TTX removal (Figure 6b). Consistent with previous findings (Yuan *et al.,* 2023), this increase in bursting did not alter the mean firing rate (MFR) because the rise in NBR was offset by a reduction in spikes per burst (Figure 6c,d). In contrast, *NPHP1^−/-^*and *CEP290^−/-^* networks failed to exhibit comparable reorganization (Figue 6e–j). Their response to TTX washout remained indistinguishable from baseline (Figure 6k), indicating a blunted or absent homeostatic adaptation. Interestingly, the baseline activity patterns of both mutant networks, characterised by elevated NBR and reduced spikes per burst (Figure 4) resembled those of TTX-treated controls, suggesting a pre-existing, homeostatic-like state. These findings indicate that loss of *NPHP1* or *CEP290* impairs the network’s capacity to undergo homeostatic plasticity and may render the network functionally occluded to further adaptation.

**Figure 6:**
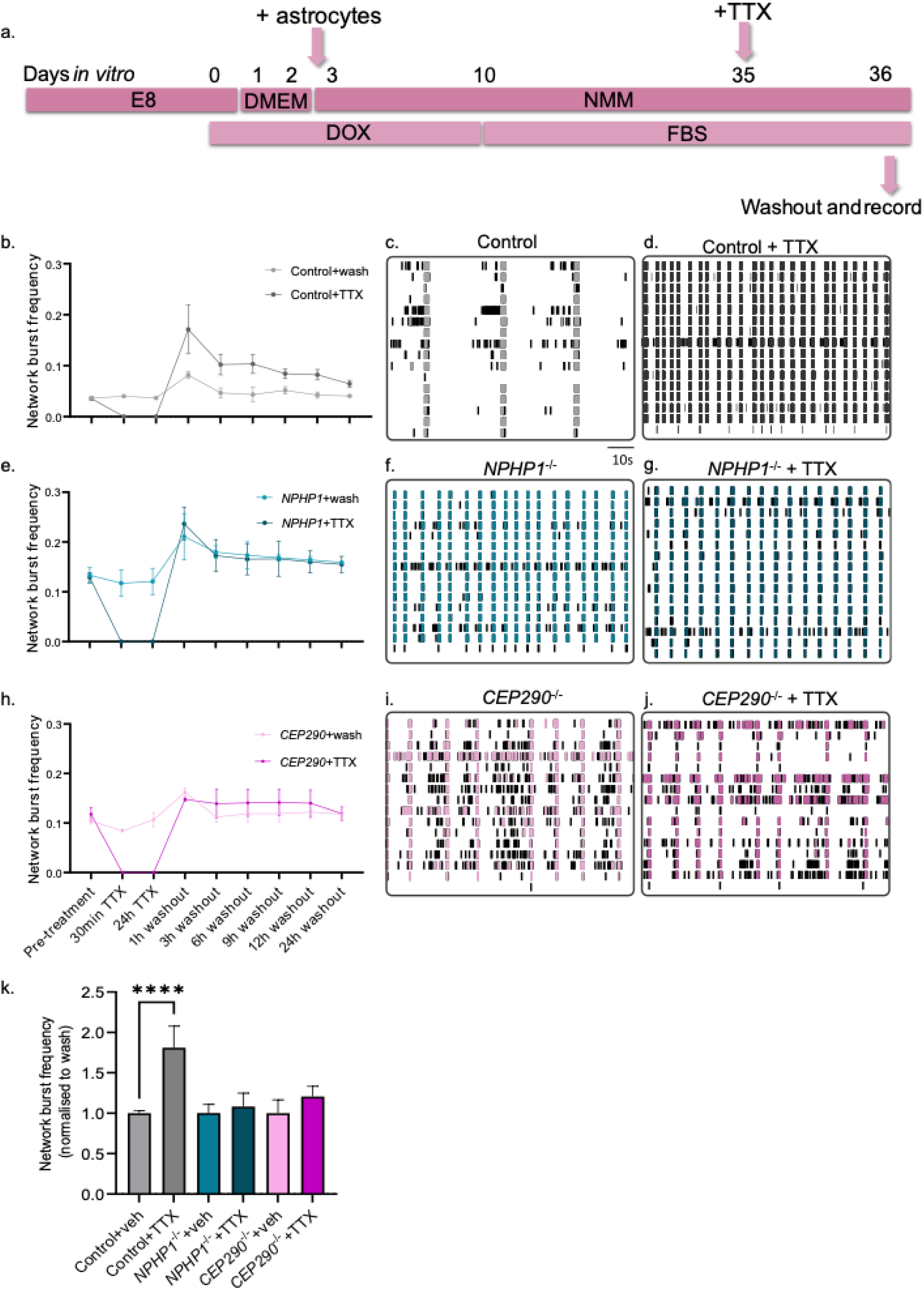
Functional cilia are necessary for proper homeostatic plasticity response at a functional level. (a) Protocol for inducing homeostatic plasticity on a MEA, neurons are treated at DIV35 with TTX for 24h (b, e, h) the network burst frequency of control, *NPHP1^−/-^* and *CEP290^−/-^* neurons following induction of homeostatic plasticity over time. Representative rasterplots at 3h can be seen in (c-d; f-h; i-j). Graphs represent network burst frequency from N=3 independent experiments across 9 wells. (k) shows the fold change in NBF at 1h following TTX washout normalised to the vehicle washout condition of each respective cell line. Data represents mean ± SEM, one-way ANOVA with Dunnett’s multiple comparisons, ****p < 0.0001.

Next, we asked if under the same circumstances AIS length increase and GluA2 incorporation were affected. Comparing to control neurons both *NPHP1^−/-^* and *CEP290^−/-^* neurons did not show significant changes in AIS length following 24h activity suppression (Figure 7a, c, d). Cumulative frequency analysis (Figure 7c) suggests that AIS length in ciliary mutants falls intermediate between control baseline and TTX-treated neurons, but differences were not statistically significant. We further investigated synaptic composition and found that the activity-dependent increase in synaptic GluA2 incorporation observed in control neurons was absent in both *NPHP1^−/-^* and *CEP290^−/-^* neurons (Figure 7b, e, f). Together, these data indicate that loss of ciliary proteins NPHP1 and CEP290 results in a blunted homeostatic plasticity response.

**Figure 7:**
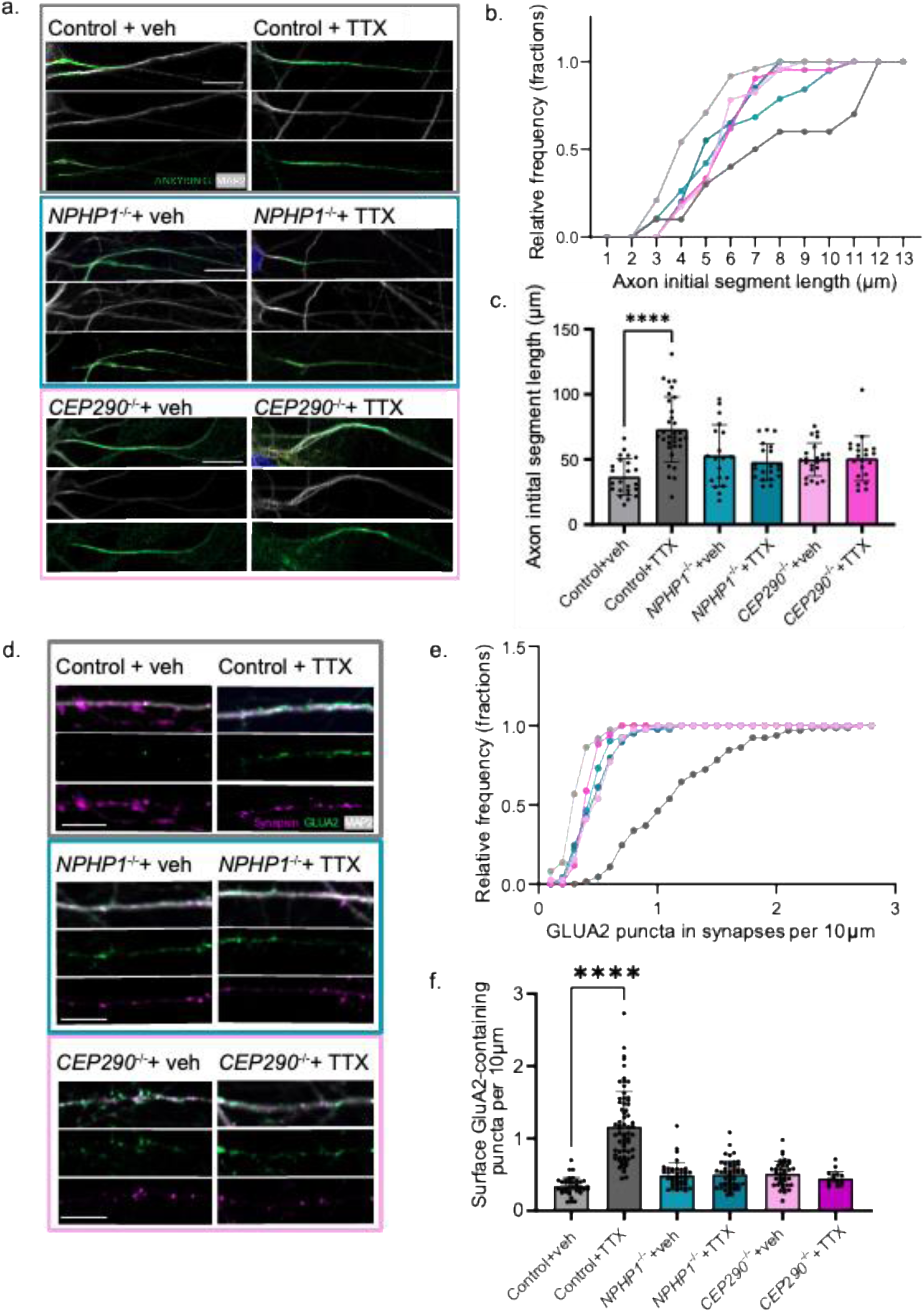
Loss of ciliary proteins leads to an attenuated homeostatic plasticity response at a morphological level. (a) Representative immunofluorescence staining of the AIS with AnkyrinG (green) and MAP2 (white) across the different cell lines, with vehicle-treated on the left and TTX-treated on the right. (b) cumulative frequency plot of AIS length in vehicle- and TTX-treated neurons between the different cell lines. (c) Measurement of AIS length in vehicle- and TTX-treated neurons between the different cell lines. N=3 independent experiments and 20-33 cells. Data represent mean ± SEM and asterixis denote significance following one-way ANOVA and Turkey’s multiple comparisons test. ****p < 0.0001. (d) Representative immunofluorescence staining of MAP2 (white), Synapsin1 (green), and surface GLUA2 (magenta) across the different cell lines, with vehicle treated on the left and TTX-treated on the right. (e) Cumulative frequency plot of incorporation of GLUA2 into synapses per 10μm of dendrite (f) Measurement of GLUA2-containing synapsin-positive puncta per 10 μm of dendrite. N=3 independent experiments and 22-65 cells. Data represent mean ± SEM and asterixis denote significance following one-way ANOVA and Turkey’s multiple comparisons test. ****p < 0.0001.

### Ciliary disruption shifts the transcriptome towards a perturbed network state

To further understand why the ciliary mutant neurons appear unable to respond to activity suppression, we compared transcriptome changes upon TTX-treatment in all genotypes. We hypothesised that the ciliary mutants would exhibit an attenuated homeostatic plasticity response because they are at a state of occlusion - whereby there is already an enrichment in the expression of genes that are necessary for induction of homeostatic plasticity.

Comparison of TTX-treated genotypes with the vehicle-treated ones (Figure 8a-c) revealed that several genes were differentially expressed between the two conditions across all the genotypes. Across the three genotypes, induction of homeostatic plasticity induced robust increase in expression of several genes including stress response gene *Thioredoxin-interacting protein* (*TXNIP*), and ubiquitination precursor *Arrestin Domain Containing 4* (*ARRDC4*).

**Figure 8:**
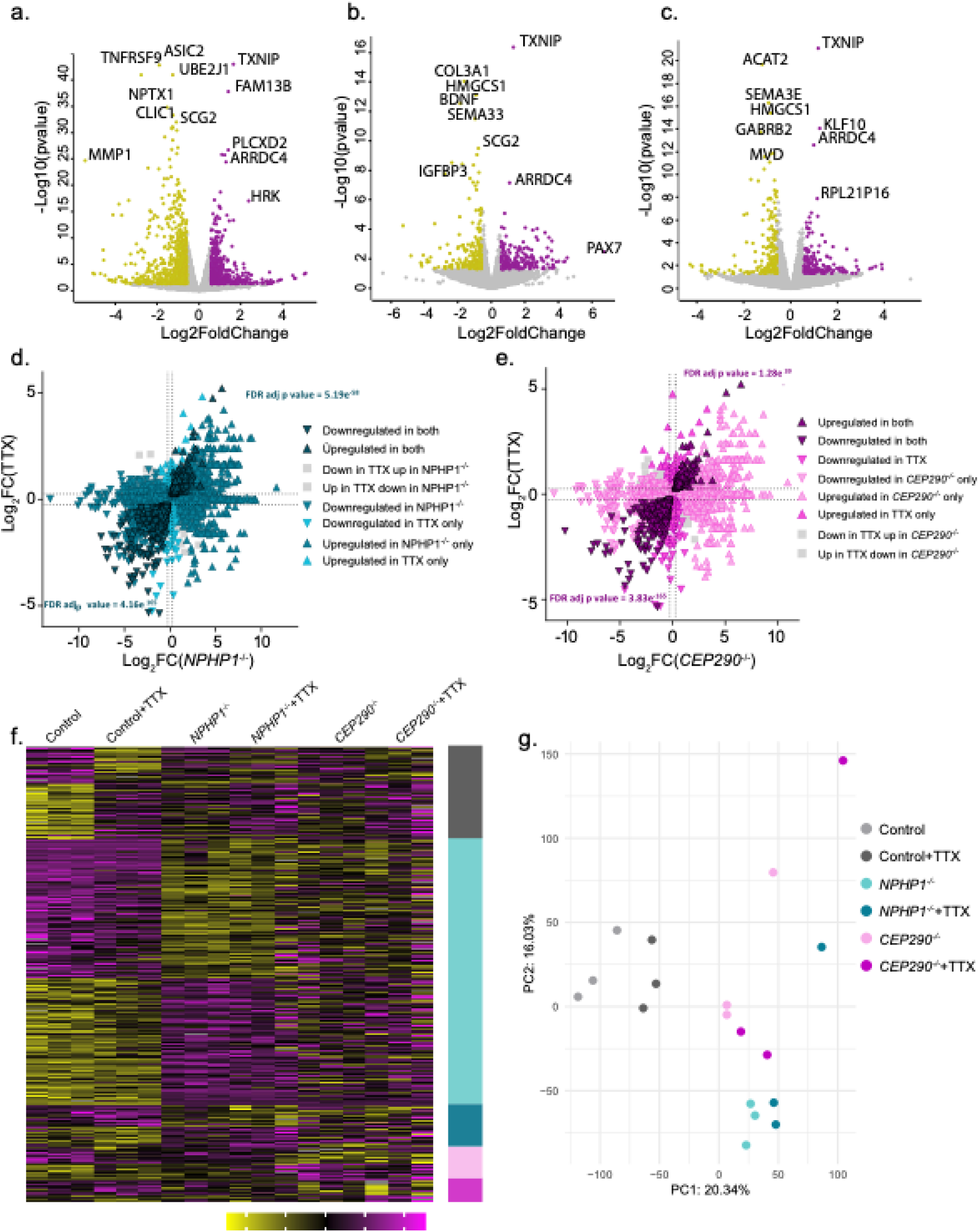
Transcriptome of ciliary mutants is less altered following homeostatic plasticity induction than the control line. (a) Volcano plot of *NPHP1^−/-^*+TTX with respect to *NPHP1^−/-^*+vehicle, highlighting significantly upregulated genes in purple and downregulated in yellow. (b) Volcano plot of Control +TTX with respect to Control +vehicle, (c) volcano plot of *CEP290^−/-^*+TTX with respect to *CEP290^−/-^*+vehicle. Genes with pvalue<0.05 and LogFDR<-0.58 shown in yellow and LogFDR>0.58 in purple. (d and e) show 4-way volcano plots of the gene expression of *NPHP1^−/-^* with and without TTX and *CEP290^−/-^* with and without TTX with respect to the control neuron gene expression. Significance of Fisher correlation is shown in the corners. (f) heatmap with clustering showing normalised differential gene expression between the 6 conditions. (g) principal component analysis (PCA) of RNA-sequencing data with three samples per condition (Control + vehicle, Control+ TTX, *NPHP1^−/-^*+vehicle, *NPHP1^−/-^*+TTX, *CEP290^−/-^*+vehicle and *CEP290^−/-^*+TTX).

To test our occlusion hypothesis, we compared the basal expression profiles of *CEP290^−/-^* and *NPHP1^−/-^* neurons with control neurons following homeostatic plasticity induction (Figure 8d, e). This analysis revealed a strong correlation between genes differentially expressed during homeostatic plasticity in controls and those already dysregulated at baseline in mutants, indicating pre-existing perturbations in homeostatic plasticity-associated gene networks.

Clustered heatmap visualisation (Figure 8f) showed clear segregation of genes by TTX treatment in controls, but less distinct separation between vehicle- and TTX-treated mutant neurons. Principal component analysis (PCA) further revealed that vehicle-treated *CEP290*^−/-^ and *NPHP1*^−/-^ neurons cluster separately from controls and closer to TTX-treated control neurons (Figure 8g). Collectively, these findings suggest that ciliary mutant neurons exist in a transcriptional state resembling chronically suppressed activity, thereby potentially occluding the normal homeostatic plasticity response.

Taken together, our transcriptomic analysis provides molecular evidence that ciliary dysfunction alters homeostatic plasticity machinery also at the gene expression level. There is a high correlation between the baseline gene expression in ciliary mutants and TTX-induced expression in control neurons, demonstrating that *CEP290*^−/-^ and *NPHP1^−/-^* neurons exist in a constitutive state of transcriptional activation resembling chronic homeostatic stress.

## Discussion

In this study, we used human iPSC-derived neurons to elucidate the role of the primary cilium in postmitotic neuronal function and plasticity. We find a critical role for the primary cilium in homeostatic plasticity. We demonstrate that changes in ciliary morphology occur bidirectionally alongside network activity alterations, proposing a timeline where intrinsic excitability and synaptic adaptations precede ciliary changes following chronic activity suppression. Using CRISPR-Cas9 targeting key ciliary genes involved in disease, we show that functional neuronal cilia are essential for establishing baseline network activity and for adaptive responses to prolonged activity deprivation, positioning the primary cilium as a critical cellular compartment for circuit plasticity with implications for learning and behaviour. Neuronal primary cilia remain understudied, especially in human neuronal models. We demonstrate that the size and composition of the cilium changes in accordance with what is known of cilia in the developing mammalian CNS (<u>Kasahara *et al*. 2014; Sterpka and Chen</u> <u>2018</u>).

Our investigation was inspired by the structural and functional parallels between the primary cilium and dendritic spines, both of which are specialised compartments with distinct receptor and ion channel populations regulated by diffusion barriers (the ciliary transition zone and spine neck, respectively) and responsive to extracellular stimuli (Nechipurenko *et al.,* 2013). Like dendritic spines, which show bidirectional morphological changes in response to altered activity (Hobbiss *et al*., 2018), cilia similarly exhibited bidirectional plasticity following hyper- and hypoactivity. To avoid confounding effects of KCl-induced depolarization on cell health and microtubule stability, we focused on chronic activity suppression to probe homeostatic plasticity. Our observations confirmed bidirectional changes in ciliary morphology following induction of network hyperactivity with KCl, or hypoactivity through use of TTX in the same direction as spine morphology changes.

Recent evidence suggests that the neuronal primary cilium may serve as a key locus for homeostatic plasticity by integrating compartmentalised signalling and intrinsic excitability control. Two recent studies (Macarelli *et al*., 2025; Kostyanovskaya *et al*., 2024) using proteomic approaches reported a broad array of receptors, channels, and synaptic proteins expressed in neuronal cilia, notably including SYNGAP1, STXBP1, PSD95, SCN2A, SLC6A1, and SHANK3 (Kostyanovskaya *et al*., 2024). This inventory suggests that cilia are not passive sensory organelles but active compartments capable of sensing network activity and influencing cellular excitability that integrates multiple dimensions of excitability and synaptic signalling. Supporting this view, disruption of ciliary signalling in pyramidal neurons was shown to rapidly strengthen excitatory synapses and increase firing rates, while pharmacological modulation of the cilium-localised GPCR SSTR3 bidirectionally adjusted excitatory synaptic strength, demonstrating that ciliary GPCR pathways can dynamically regulate excitatory transmission and circuit excitability (Tereshko *et al*., 2021). Together, these points support a model of the cilium as an active homeostatic sentinel, and a critical future step will be to manipulate specific receptors or channels within single cilia—using opto/chemogenetic tools, cilia-restricted degradation, or localised reporters—to directly test whether compartment-specific perturbations are sufficient to restore activity to baseline.

Examining the sequence of plasticity changes revealed that AIS elongation and increased synaptic GluA2 incorporation precede ciliary lengthening, reflecting differences in protein synthesis and trafficking constraints, as cilia lack local protein synthesis unlike dendritic spines (Bourne *et al*., 2007). While AIS and synaptic modifications have well-characterised functional consequences, the significance of ciliary elongation remains unclear; it may reflect accumulation of unknown membrane-bound proteins or enhanced sensitivity to extracellular stimuli.

Most models of ciliary dysfunction are developmental, where differences in circuit connectivity could be attributed to neurophysiological alterations, such as migration defects or reduction of synaptic integration. One study has addressed whether acute disruption of ciliary function is sufficient to alter activity. In line with our observations they found that treatment with shRNAs against a number of ciliary proteins led to a significant increase in GluA2 incorporation at excitatory synapses (<u>Tereshko *et al*. 2021</u>). One advantage of an inducible neuron system is that we can rapidly differentiate hiPSCs into glutamatergic neurons and thus bypass early developmental stages that ciliary function is already understood to play a critical role in. By studying hiPSC-derived neurons, we gain greater insight into the role of the cilium in post-mitotic neurons. Surprisingly, the network activity of the *NPHP1^−/-^* and *CEP290^−/-^*neurons showed no change in MFR despite the increase in NBR. We hypothesised that this activity resembles that of control lines following the induction of homeostatic plasticity, where there is no overall increase in activity, but rather a rearrangement of the network. Subsequent treatment of day 35 neurons with TTX revealed that there is an inability of the ciliary mutants to properly modulate synaptic and intrinsic properties in order to increase their activity in response to chronic activity suppression.

Finally, ciliopathy candidate genes increasingly overlap with neurodevelopmental disorder - associated proteins (Kostyanovskaya *et al.,* 2024), implicating ciliary dysfunction in diverse brain pathologies. Divergent ciliary morphologies yet convergent plasticity deficits highlight potential therapeutic targets for ciliopathy-related cognitive impairments. Future studies could investigate whether these changes are cell intrinsic or depend on astrocyte-derived cues. Culturing hiPSC-derived neurons without astrocytes could reveal whether ciliary changes are cell-intrinsic, and using *NPHP1*^⁻/⁻^ or *CEP290*^⁻/⁻^ astrocytes could uncover phenotypes masked by control astrocytes.

Together, our findings reveal a novel role for the primary cilium in establishing and maintaining neural network stability and enabling homeostatic plasticity. Dysfunction of this organelle disrupts key synaptic and intrinsic adaptations, likely contributing to the neurodevelopmental phenotypes observed in neuronal ciliopathies such as Joubert syndrome. The observed state of molecular occlusion, in which genes critical for homeostatic plasticity are already differentially expressed under basal conditions, illustrates how neurons lacking intact ciliary signalling fail to appropriately adapt to chronic activity suppression. More broadly, these findings position the primary cilium as an essential regulator of the homeostatic mechanisms that safeguard network function, highlighting its central role in linking cellular signalling to circuit-level stability.

## Material and Methods

### Human iPSC lines

All iPSC lines were cultured on Geltrex-(Gibco; A1413301) or Matrigel®-(Corning) coated plates in E8 Flex™ media (Gibco; A2858501) supplemented with 0.1 µg/ml Primocin® (Invivogen; ant-pm-1), 0.5 µg/ml puromycin (Sigma-Aldrich; P8833) and 50 µg/ml G418 (Sigma-Aldrich; G8168). hiPSCs were maintained at 37°C, 5% CO2, 21% O2. Cells were clump passaged (1:15) using ReleSR (Gibco) when ∼80% confluent. Cells were tested for mycoplasma every two weeks they were in culture with MycoAlertTM Mycoplasma Detection Kit (Lonza). Additionally Ngn2/rTTA hiPSCs were cultured in E8™ Flex medium (A2858501, Thermo Fisher Scientific) supplemented with 0.1 µg/ml Primocin® (ant-pm-1, Invivogen), 0.5 µg/ml puromycin (P8833, Sigma-Aldrich) and 50 µg/ml G418 (G8168, Sigma-Aldrich).

Control cell line (UCSFi001-A, Coriell Institute) was used to generate isogenic lines with frameshift variants in *CEP290* (UCSFi001-A-71) and *NPHP1* (UCSFi001-A-68) as described in Dyke *et al*. 2023. Briefly, a guide RNA targeting exon 20 of *CEP290* (5’-AGAAATATTGCAAGCAATTA-3’) was cloned into pSpCas9(BB)-2A-Puro (PX459) V2.0 (a gift from Feng Zhang; Addgene plasmid # 62988 ; http://n2t.net/addgene:62988; RRID:Addgene_62988). Following electroporation with P3 Primary Cell 4D-NucleofectorTM X kit (Lonza; V4XP-3024, program CA-137), resultant clones underwent puromycin selection (0.5μg/ml puromycin (Sigma Aldrich)) for 24h before surviving clones were manually picked and expanded. Sanger sequencing confirmed the mutations (Supplementary figure 2) indicating deletion events on both alleles. Gene expression was determined by RT-qPCR (Supplementary figure 2) revealing dramatically reduced transcript expression. Clones showed typical iPSC morphology and normal (46XY) karyotype in addition to STR analysis of 16 loci, all matched. Pluripotency and trilineage potential (using STEMdiff™ Trilineage Differentiation Kit (StemCell; 05230)) were assessed via immunofluorescence of pluripotency markers (OCT4 and NANOG) and via RT-qPCR (Supplementary table 2) (See Dyke *et al*. 2023 for *NPHP1^−/-^* and Supplementary figure 2 for *CEP290^−/-^*). Karyotyping was regularly checked with ddPCR (Stemgenomics).

### Generation of rtTA/Ngn2 stable iPSC lines and iNeuron culture

iNeurons were generated and maintained as previously described (<u>Frega *et al*. 2017</u>). In brief, for differentiation, hiPSCs were split into single cells using TrypLE Express (12604021, Gibco) and plated at a density of 20,000 cells per 1.9cm2 onto 12mm coverslips (631-0713, Thermo Fisher) coated with poly-L-ornithine hydrobromine (50 mg/mL, P3655, Sigma-Aldrich) and human recombinant laminin (5 ug/mL, LN521-05, Biolamina). On the day of plating, 0 days *in vitro* (DIV0), cells were plated in E8™ Flex medium supplemented with 0.1 µg/ml Primocin®, 4 μg/mL doxycycline, and 10 μg/mL RevitaCell (A2644501, Thermo Fisher).

### Micro-electrode array recording and analysis

Neurons were plated as above 24- and 48-well Cytoview MEA plates (Axion Biosystems, M384-tMEA-24W; M768-tMEA-48W) at a final concentration of 600 neurons/um^2^. Spontaneous activity was recorded weekly, 24h after a half medium change on a Maestro Pro MEA system equipped with AxIS Navigator software (Axion BioSystems, Atlanta, GA, USA) at 37°C and 5% CO2. Activity was recorded for 5 minutes following a 10 minute acclimation period in the system.

Activity parameters were extracted using AxIS Neural Metrics Tool (Axion BioSystems). Network bursts were defined with the built-in envelope algorithm, with a threshold factor of 1.5; minimum interburst interval 100ms; minimum burst inclusion set to 50%, and the minimum number of participating electrodes also set to 50%.

PCA and DIV28 analysis (total across 4 independent batches, n=40 for control, n=68 for *NPHP1*^−/-^, n=21 for *CEP290*^−/-^)

### Treatments with compounds

All compounds were added one day after half medium changes to minimise potential effects of the medium change on cellular activity. To increase neuronal activity, 10 μM potassium chloride (KCl, Sigma, P9333) was applied for 3 hours prior to fixation. To modify ciliary volume, 1 μM SB-399885 (Sigma-Aldrich, SML0604) was applied for 24 hours, as per the protocol used previously in striatal neurons (<u>Brodsky *et al*. 2017</u>). Tetrodotoxin citrate (TTX, Alomone Labs, T-550) was initially applied as previously published (Yuan *et al*. 2023), however the introduction of a different microelectrode array system meant we could reduce the treatments to 24 h of 1 μM TTX application at DIV35. In instances requiring washout, conditioned medium (a 1:1 mixture of medium from the previous medium changes and fresh medium) were used to wash the wells three times and finally to refresh the wells. Washes were performed on a warming plate at 37 ℃ (CooperSurgical, WP37). Vehicle treated wells underwent the same procedure, however using ddH2O as opposed to TTX.

### Immunofluorescence

Neurons were fixed with 4% paraformaldehyde (Sigma Aldrich; 441244)/4% sucrose (Sigma Aldrich; S7903) for 30 minutes before permeabilization in blocking buffer consisting of 5% normal goat serum (Invitrogen;), 5% normal donkey serum (Gibco), 5% normal horse serum (Jackson Immuno), 1% BSA (Sigma Aldrich), 1% glycine (Sigma Aldrich), 0.1% lysine (Sigma Aldrich), 0.4% Triton X-100 (Sigma Aldrich) in PBS (Sigma Aldrich; P5493) for 1 hour. Cells were incubated in primary antibodies diluted in blocking buffer overnight at 4°C in a humid chamber.

The following antibodies were used: Rabbit anti-ADCY3 (1:1000, Novus Biologicals; NBP1-92683), Mouse anti-ARL13B (1:1000, NeuroMab; 75-287), Guinea pig anti-MAP2 (1:1000, Synaptic Systems; 188 004), Mouse anti-ANKG (1:300, Antibodies Online; N106-36), Mouse anti-GluA2 (1:200, Invitrogen; 32-0300), Rabbit anti-Nanog (1:1000, Abcam, #ab21624), Rabbit anti-Oct4 (1:200, Abcam, ab19857), Rabbit anti-Synapsin (1:600, Sigma-Aldrich; AB1543P), Goat anti-Rabbit Alexa Fluor 488 (1:1000; Molecular probes; A11034), Goat anti-Mouse Alexa Fluor 568 (1:1000; Molecular probes; A11031), Goat anti-Guinea Pig Alexa Fluor 647 (1:1000; Molecular probes; A21450). Coverslips were incubated with secondary antibodies and DAPI (1:80,000 Thermo Fisher Scientific; D1306) at RT for 1h and mounted with Fluoromount-G® mounting medium (Southern Biotech; 0100-01). Images were acquired using Zeiss Axio Imager Z1, Zeiss LSM880 with Airyscan 2 module, Zeiss LSM900 with Airyscan 2 module, and Zeiss Axio-observer with Yokogawa CSU-X1.

### RNA-seq library preparation

For preparation of samples for RNA sequencing, 100,000 neurons were plated at a 1:1 ratio with rat astrocytes in 6 well plates (Corning). For quality control, the same cells were used to plate microelectrode arrays and coverslips for immunofluorescence in parallel, and all were grown using the same lots of reagents. Following 24h of TTX application (or vehicle), cells were harvested in 100 ul of DNA/RNA shield (Zymo Research Corporation, ZY-R1200) and RNA was extracted using the Zymo Quick-RNA™ Microprep Kit (Zymo Research Corporation, ZY-R1051). Samples were resuspended in 25 μl ddH2O. RNA integrity number was assessed using the Agilent 4150 TapeStation System, which resulted in RIN values between 9.4 and 9.8. cDNA libraries were prepared using The NEBNext Ultra II Directional RNA Library Prep Kit, and paired-end reads were sequenced on an Illumina NovaSeq 6000 platform at GenomeScan B.V. Leiden.

### RNA-seq data processing

Fastp, with a list of adapter sequences currently used by Illumina was used to eliminate PolyG artifacts and clip adapters. We utilised Seal from BBTools to segregate rat and human reads by aligning them to the respective reference genomes (Rnor_6.0 for rat and GRCh38 for human) while selecting the best mapping with the “ambig=first” setting. We mapped the human reads to the GRCh38 human reference genome using HISAT2, setting --rna-strandness = “RF”. SAM files were then sorted and indexed with SAMtools. UMI deduplication was performed with UMI-tools, and counting was done with Subread’s featureCounts.

### RNA-seq analysis

Raw count matrices were loaded into R version 4.2.1. For Ensembl IDs mapping to the same gene symbols, only the ID with the highest expression per sample was retained. Lowly expressed genes were filtered out by retaining only those with expression values greater than 0.1 counts per million in at least three samples. Counts were subsequently transformed into a DGEList using the DGEList function from edgeR package and setting the condition as the grouping parameter. Then, we applied voom normalisation using limma package, including normalization factors calculated with edgeR’s calcNormFactors. For principal component analysis (PCA), we used the prcomp function on scaled log2(voom normalised counts + 1). Gene expression was modelled using a generalised linear model with conditions as covariates without including an intercept. We used the model.matrix function from the stats package to generate the design matrix. Then, estimateDisp was applied to model data dispersion, and glmQLFit, both from the edgeR package, was used to fit a quasi-likelihood negative binomial generalised log-linear model and perform gene-wise statistical testing for the covariates included in the design matrix. Using the makeContrasts function from the limma package, we created a contrast matrix to specifically assess the effect of *NPHP1* and *CEP290* loss relative to control+Veh, and the effect of *NPHP1*+TTX and *CEP290*+TTX relative to control+TTX, to evaluate the impact of activity deprivation on each mutation compared to deprivation in the control condition. A quasi-likelihood F-test was performed for each contrast to statistically asses the comparisons, using glmQLFTest. Genes with FDR-adjusted p-value < 0.05 and absolute Log2 fold change > 0.58 were considered differentially expressed.

To test for the enrichment of homeostatic plasticity (TTX)-associated genes among the differentially expressed genes in the ciliary mutant lines, we performed a Fisher’s exact test using the fisher.test function with the parameter alternative=“greater”. We also performed gene ontology and pathway enrichment analyses separately for differentially expressed genes in each comparison, as well as for shared differentially expressed genes between ciliary mutant lines and control networks following activity deprivation. For this, we used the gost function from the gprofiler2 v0.2.1 R package. To manage redundancy among enriched gene ontology terms (including biological processes, cellular components, and molecular function), we performed clustering analysis and aggregated terms with high semantic similarity using the calculateSimMatrix and reduceSimMatrix functions from the rrvgo v1.2.0 R package, setting threshold=”0.7”.

### Quantifications and analysis

#### Cilia volume

Ciliary volume was assessed for the technical reason that the cilia of neurons do not lie flat but rather extend into the z plane, lying over the soma. Although it is well documented that ARL13B and ADCY3 are present in distinct domains of the cilium, utilising confocal microscopy, we were unable to detect non-contiguous staining along the cilia of neurons or astrocytes in our cultures. Z-stacks of 0.15µm steps were acquired using the Zeiss LSM880 with Airyscan 2 module, and Airyscan processing was conducted in ZEN 3.1 blue edition version 3.1.0 (Zeiss). Following acquisition, images were analysed in (Fiji Is Just) ImageJ 2.14 and underwent median filtering (radius 2.0 px) prior to volume measurement.

#### Synapse analysis

To calculate the incorporation of GLUA2 into the synapse, the number of synapsin and co-localised GLUA2 puncta per 10 μm were manually counted in (Fiji Is Just) ImageJ 2.14. Between 2 and 4 secondary neurites were assessed per neuron and the average was plotted.

#### Statistical analysis

Statistical analysis was performed with Prism 9.0 (GraphPad). Data were analysed using either Student’s *t*-test (two-group comparison) or one-way ANOVA with Tukey’s multiple-comparisons test when comparing multiple conditions to each other, and Dunnett’s when comparing to one control group. The threshold for significance was set as \**p* < 0.05, \*\**p* < 0.01 and \*\*\**p* < 0.001

## Supporting information

Dyke et al., Supplementary Files

## Data availability

Data reported in this manuscript are available from the corresponding authors upon request. The software used for data analysis is described in the corresponding Methods sections of the Supplementary Information.

## Acknowledgements

This work was financially supported by grants from the European Research Council (the European Union’s Horizon 2020 research and innovation program under the Marie Skłodowska-Curie grant agreement No. 861329 to R.R. and N.N.K.), the Dutch Research Council (NWO ENW-M2, OCENW.M20.216 to N.N.K. and B.L.L), the Simons Foundation Autism Research Initiative (SFARI, 890042 to N.N.K.), and the Koolen-de Vries Foundation (to B.L.L.).

## Author Information

E.D., B.L.L., and N.N.K designed the research. E.D. and A.O. performed the research. E.D. and S.P. analysed the data. R.R. and N.N.K acquired funding. R.R., B.L.L., and N.N.K supervised the project. E.D., B.L.L., and N.N.K. wrote the manuscript with input from all authors.

## Ethics declaration Competing interests

The authors declare no competing interests.

## Notes

### Competing Interest Statement

The authors have declared no competing interest.

