## Supplementary material for "The Neuronal Primary Cilium is a Key Regulator of Homeostatic Plasticity": Dyke et al., Supplementary Files

[1]Department of Human Genetics, Radboud University Medical Center, Nijmegen, the Netherlands
[2]Donders Institute for Brain, Cognition and Behaviour, Nijmegen, the Netherlands
[3]Radboud institute for Molecular Life Sciences, Nijmegen, the Netherlands
[4]Shared last authorship

Corresponding author:
[*]Brooke Latour:

**Supplementary data file**

- Supplementary materials and methods
- Supplementary table 1: MEA parameters used for PC
- Supplementary table 2: Primers used for Sanger sequencing of $CEP290^{CR}$ clones
- Supplementary table 3: Primers used for RT-qPCR following trilineage analysis
- Supplementary table 4: Gene ontology analysis for ciliary mutants
- Supplementary table 5: Ciliary genes differentially expressed in ciliary mutants
- Supplementary figures

**Supplementary materials and methods:**

PCR:

To confirm genomic alterations following CRISPR-Cas9 editing, clones of hiPSCs were characterised by PCR. PCR amplification was performed using AmpliTaq Gold™ 360 (Thermo Fisher Scientific, 4398881) in accordance with the manufacturer's instructions. Homozygous knockouts were confirmed by Sanger sequencing using primers around the predicted cut-site (Supplementary Table 1).

qPCR

To confirm loss of gene expression, we performed RT-qPCR. Cells were lysed and collected in DNA/RNA shield (Zymo Research corporation, ZY-R1200) prior to RNA extraction using Zymo Quick-RNA Microprep kit (Zymo Research corporation, ZY-R1051). cDNA was synthesised from the RNA using iScript™ cDNA Synthesis Kit (Bio-Rad, 1708890). qPCR was performed on QuantStudio™ 3 Real-Time PCR System (ThermoFisher, A28567), with GoTaq® qPCR Master Mix (Promega, A6001). GAPDH, 18S RNA, and GUSB were used as internal controls. Primers used can be found in Supplementary Table 2.

Karyotyping:

6 cells in metaphase were studied for at 300-500 bhps. Analysis performed by the Stem Cell Technology and Differentiation Centre (RadboudUMC, Netherlands).

Morphological analysis:

Structural analysis of hiPSC-derived neurons was conducted at DIV28. Briefly, cells were fixed with 4%PFA/4%sucrose and stained against Guinea pig anti-MAP2 (1:1000, Synaptic Systems; 188 004). 20x images of neurons were acquired on Zeiss Axio Imager with Apotome optical sectioning (5x), such that all the MAP2 signal from one neuron was visible in one field of view. Somatodendritic reconstructions were performed on NeuroLucida 360 software (v11, MBF-Bioscience, USA).

Heatmap generation:

To generate the heatmap (Supplementary Figure 5), RNAseq data was uploaded to usegalaxy.org (Afgan *et al.* 2018) and visualised with heatmap2 tool which utilises the heatmap.2 function from the R gplots package.

**Supplementary Table 1: Parameters used for principal component analysis**

| Parameters used for principal component analysis | |
|---|---|
| | Number of Spikes |
| | Mean Firing Rate (Hz) |
| | ISI Coefficient of Variation – Avg |
| | Weighted Mean Firing Rate (Hz) |
| | Number of Bursts |
| | Burst Duration - Avg (sec) |
| | Number of Spikes per Burst - Avg |
| | Mean ISI within Burst - Avg (sec) |
| | Median ISI within Burst - Avg (sec) |
| | Median/Mean ISI within Burst - Avg |
| | Inter-Burst Interval - Avg (sec) |
| | Burst Frequency - Avg (Hz) |

| | |
|---|---|
| | IBI Coefficient of Variation - Avg |
| | Burst Percentage - Avg |
| | Network Burst Frequency |
| | Network Burst Duration - Avg (sec) |
| | Number of Spikes per Network Burst – Avg |
| | Median/Mean ISI within Network Burst – Avg |
| | Number of Elecs Participating in Burst – Avg |
| | Number of Spikes per Network Burst per Channel – Avg |
| | Network Burst Percentage |
| | Network IBI Coefficient of Variation |
| | Network Normalized Duration IQR |
| | Area Under Normalized Cross-Correlation |

**Supplementary Table 2: Primers used for Sanger sequencing of CEP290$^{CR}$ clones**

| | Target | Forward/Reverse primer (5'-3') |
|---|---|---|
| gRNA oligonucleotide | CEP290 gRNA | AGAAATATTGCAAGCAATTA/ TAATTGCTTGCAATATTTCT |
| Targeted mutation analysis/sequencing | CEP290 (primer pair 1) | TGCACGTGATATAAATAGGTCTGTCA/ agcagctcaagaaaaactacactt |
| Targeted mutation analysis/sequencing | CEP290 (primer pair 2) | CTGATCATCCAATGAGAAAACTGTGT/ aaacgagctccaaagtcggtgt |
| Targeted mutation analysis expression | CEP290 (qPCR primer pair 1) | AGCCTTGAAAGACTAGTTAATGCTA/ GCTTTTGCCAACTGCTGTGA |
| Targeted mutation analysis expression | CEP290 (qPCR primer pair 2) | TAGCACCATCTAGTGCCAGT/ TCCAGGTCTCCTTTTCACTTAGG |
| Off target analysis | Off target 1 | ACAAGGGTGAGGAACGTTTG/ CCCATGGGTAGAAGGGACCTC |
| Off target analysis | Off target 2 | CGCATGCTAAGCACACGATC/ TCCCTCCCTCAACCCCTATGC |
| Off target analysis | Off target 3 | CATAGCAACAAGCAGGAACCA/ GCAAAGCCAAGTGTTTCAGG |
| Off target analysis | Off target 4 | CCCTACTTGTCTTTCTCTCCTTGT/ GTCAGGATTATGCTTGCGTTGT |

**Supplementary table 3: Primers used for RT-qPCR following trilineage analysis**

| | Target | Forward/Reverse primer (5'-3') |
|---|---|---|
| House-Keeping Genes (qPCR) | 18s RNA | GTAACCCGTTGAACCCCATT/CCATCCAATCGGTAGTAGCG |
| House-Keeping | GAPDH | GTGGACCTGACCTGCCGTCT/GGAGGAGTGGGTGTCGCTGT |

| | | |
|---|---|---|
| Genes (qPCR) | | |
| House-Keeping Genes (qPCR) | GUSB | AGAGTGGTGCTGAGGATTGG/CCCTCATGCTCTAGCGTGTC |
| Trilineage Genes (qPCR) | PAX6 | GCTGCAAAGAAATAGAACATCC/TTGGCTGCTAGTCTTTCTCG |
| Trilineage Genes (qPCR) | SOX17 | GAACGCTTTCATGGTGTGGG/CCTTCCACGACTTGCCCAG |
| Trilineage Genes (qPCR) | BRACHYURY (TBXT) | TGCTTCCCTGAGACCCAGTT/GATCACTTCTTTCCTTTGCATCAAG |
| Trilineage Genes (qPCR) | OTX2 | AGGAGGTGGCACTGAAAATC/TGACCTCCATTCTGCTGTTG |
| Trilineage Genes (qPCR) | HAND1 | GAACTCAAGAAGGCGGATGG/CGGTGCGTCCTTTAATCCTC |
| Trilineage Genes (qPCR) | CXCR4 | TGGAGGGGATCAGTATATACACT/ATGGTGGGCAGGAAGATTTT |

**Supplementary table 4: Gene ontology analysis for ciliary mutants**

**CEP290_Down:**

| | |
|---|---|
| Analysis Type: | PANTHER Overrepresentation Test (Released 20240807) |
| Annotation Version and Release Date: | GO Ontology database DOI: 10.5281/zenodo.12173881 Released 2024-06-17 |
| Analyzed List: | upload_1 (Homo sapiens) |
| Reference List: | Homo sapiens (all genes in database) |
| Test Type: | FISHER |
| Correction: | FDR |

| GO biological process complete | Homo sapiens - REFLIST (20580) | upload_1 (1480) | upload_1 (expected) | upload_1 (over/under) | upload_1 (fold Enrichment) | upload_1 (raw P-value) | upload_1 (FDR) |
|---|---|---|---|---|---|---|---|
| aminergic neurotransmitter loading into synaptic vesicle (GO:0015842) | 3 | 3 | 0.22 | + | 13.91 | 3.71E-04 | 1.79E-02 |
| L-leucine import across plasma membrane (GO:1903801) | 3 | 3 | 0.22 | + | 13.91 | 3.71E-04 | 1.78E-02 |
| protein localization to secretory granule (GO:0033366) | 4 | 3 | 0.29 | + | 10.43 | 1.40E-03 | 5.02E-02 |
| glutamate biosynthetic process (GO:0006537) | 4 | 3 | 0.29 | + | 10.43 | 1.40E-03 | 5.01E-02 |
| peripheral nervous system neuron axonogenesis (GO:0048936) | 4 | 3 | 0.29 | + | 10.43 | 1.40E-03 | 5.00E-02 |
| positive regulation of intracellular cholesterol transport (GO:0032385) | 4 | 3 | 0.29 | + | 10.43 | 1.40E-03 | 4.99E-02 |
| positive regulation of intracellular sterol transport (GO:0032382) | 4 | 3 | 0.29 | + | 10.43 | 1.40E-03 | 4.97E-02 |
| negative regulation of hyaluronan biosynthetic process (GO:1900126) | 4 | 3 | 0.29 | + | 10.43 | 1.40E-03 | 4.96E-02 |
| valine transport (GO:0015829) | 4 | 3 | 0.29 | + | 10.43 | 1.40E-03 | 4.95E-02 |
| peptide antigen assembly with MHC class I protein complex (GO:0002502) | 6 | 4 | 0.43 | + | 9.27 | 3.55E-04 | 1.73E-02 |
| MHC class I protein complex assembly (GO:0002397) | 6 | 4 | 0.43 | + | 9.27 | 3.55E-04 | 1.72E-02 |
| positive regulation of axon extension involved in axon guidance (GO:0048842) | 7 | 4 | 0.5 | + | 7.95 | 7.81E-04 | 3.15E-02 |

| | | | | | | | |
|---|---|---|---|---|---|---|---|
| neurotransmitter loading into synaptic vesicle (GO:0098700) | 7 | 4 | 0.5 | + | 7.95 | 7.81E-04 | 3.14E-02 |
| L-leucine transport (GO:0015820) | 10 | 5 | 0.72 | + | 6.95 | 3.54E-04 | 1.73E-02 |
| negative regulation of cellular response to hypoxia (GO:1900038) | 10 | 5 | 0.72 | + | 6.95 | 3.54E-04 | 1.73E-02 |
| NAD catabolic process (GO:0019677) | 21 | 9 | 1.51 | + | 5.96 | 6.67E-06 | 8.20E-04 |
| heparan sulfate proteoglycan biosynthetic process, enzymatic modification (GO:0015015) | 12 | 5 | 0.86 | + | 5.79 | 9.86E-04 | 3.75E-02 |
| dicarboxylic acid biosynthetic process (GO:0043650) | 12 | 5 | 0.86 | + | 5.79 | 9.86E-04 | 3.74E-02 |
| NADH metabolic process (GO:0006734) | 32 | 13 | 2.3 | + | 5.65 | 1.24E-07 | 2.76E-05 |
| neurotransmitter receptor localization to postsynaptic specialization membrane (GO:0099645) | 15 | 6 | 1.08 | + | 5.56 | 3.89E-04 | 1.86E-02 |
| protein localization to postsynaptic specialization membrane (GO:0099633) | 15 | 6 | 1.08 | + | 5.56 | 3.89E-04 | 1.86E-02 |
| detection of mechanical stimulus involved in sensory perception of pain (GO:0050966) | 15 | 6 | 1.08 | + | 5.56 | 3.89E-04 | 1.85E-02 |
| glycolytic process through glucose-6-phosphate (GO:0061620) | 20 | 8 | 1.44 | + | 5.56 | 4.04E-05 | 3.34E-03 |
| glycolytic process through fructose-6-phosphate (GO:0061615) | 20 | 8 | 1.44 | + | 5.56 | 4.04E-05 | 3.32E-03 |
| IMP metabolic process (GO:0046040) | 18 | 7 | 1.29 | + | 5.41 | 1.54E-04 | 9.42E-03 |
| NADH regeneration (GO:0006735) | 18 | 7 | 1.29 | + | 5.41 | 1.54E-04 | 9.39E-03 |
| regulation of protein kinase C signaling (GO:0090036) | 18 | 7 | 1.29 | + | 5.41 | 1.54E-04 | 9.35E-03 |
| glucose catabolic process to pyruvate (GO:0061718) | 18 | 7 | 1.29 | + | 5.41 | 1.54E-04 | 9.31E-03 |
| canonical glycolysis (GO:0061621) | 18 | 7 | 1.29 | + | 5.41 | 1.54E-04 | 9.27E-03 |
| heparan sulfate proteoglycan metabolic process (GO:0030201) | 16 | 6 | 1.15 | + | 5.21 | 5.85E-04 | 2.51E-02 |
| tail-anchored membrane protein insertion into ER membrane (GO:0071816) | 17 | 6 | 1.22 | + | 4.91 | 8.49E-04 | 3.33E-02 |

| | | | | | | | |
|---|---|---|---|---|---|---|---|
| synaptic transmission, dopaminergic (GO:0001963) | 17 | 6 | 1.22 | + | 4.91 | 8.49E-04 | 3.32E-02 |
| type B pancreatic cell proliferation (GO:0044342) | 17 | 6 | 1.22 | + | 4.91 | 8.49E-04 | 3.32E-02 |
| membrane depolarization during cardiac muscle cell action potential (GO:0086012) | 17 | 6 | 1.22 | + | 4.91 | 8.49E-04 | 3.31E-02 |
| peptide antigen assembly with MHC protein complex (GO:0002501) | 21 | 7 | 1.51 | + | 4.64 | 4.64E-04 | 2.12E-02 |
| MHC protein complex assembly (GO:0002396) | 21 | 7 | 1.51 | + | 4.64 | 4.64E-04 | 2.11E-02 |
| nucleoside monophosphate catabolic process (GO:0009125) | 18 | 6 | 1.29 | + | 4.64 | 1.20E-03 | 4.38E-02 |
| glucose catabolic process (GO:0006007) | 25 | 8 | 1.8 | + | 4.45 | 2.51E-04 | 1.32E-02 |
| membrane depolarization during action potential (GO:0086010) | 22 | 7 | 1.58 | + | 4.42 | 6.39E-04 | 2.68E-02 |
| detection of temperature stimulus (GO:0016048) | 26 | 8 | 1.87 | + | 4.28 | 3.39E-04 | 1.67E-02 |
| response to amphetamine (GO:0001975) | 36 | 11 | 2.59 | + | 4.25 | 2.87E-05 | 2.55E-03 |
| pyridine nucleotide catabolic process (GO:0019364) | 50 | 15 | 3.6 | + | 4.17 | 1.35E-06 | 2.19E-04 |
| cerebellar cortex formation (GO:0021697) | 27 | 8 | 1.94 | + | 4.12 | 4.52E-04 | 2.08E-02 |
| innervation (GO:0060384) | 27 | 8 | 1.94 | + | 4.12 | 4.52E-04 | 2.07E-02 |
| glutamine family amino acid catabolic process (GO:0009065) | 24 | 7 | 1.73 | + | 4.06 | 1.14E-03 | 4.23E-02 |
| detection of stimulus involved in sensory perception of pain (GO:0062149) | 28 | 8 | 2.01 | + | 3.97 | 5.94E-04 | 2.53E-02 |
| pyridine-containing compound catabolic process (GO:0072526) | 53 | 15 | 3.81 | + | 3.94 | 3.05E-06 | 4.44E-04 |
| glycolytic process (GO:0006096) | 46 | 13 | 3.31 | + | 3.93 | 1.41E-05 | 1.45E-03 |
| protein localization to postsynaptic membrane (GO:1903539) | 32 | 9 | 2.3 | + | 3.91 | 3.09E-04 | 1.54E-02 |
| heparan sulfate proteoglycan biosynthetic process (GO:0015012) | 29 | 8 | 2.09 | + | 3.84 | 7.68E-04 | 3.12E-02 |

| | | | | | | | |
|---|---|---|---|---|---|---|---|
| regulation of presynaptic membrane potential (GO:0099505) | 29 | 8 | 2.09 | + | 3.84 | 7.68E-04 | 3.11E-02 |
| protein localization to postsynapse (GO:0062237) | 33 | 9 | 2.37 | + | 3.79 | 3.98E-04 | 1.87E-02 |
| ADP catabolic process (GO:0046032) | 48 | 13 | 3.45 | + | 3.77 | 2.33E-05 | 2.19E-03 |
| hindbrain morphogenesis (GO:0021575) | 48 | 13 | 3.45 | + | 3.77 | 2.33E-05 | 2.18E-03 |
| cerebellum morphogenesis (GO:0021587) | 45 | 12 | 3.24 | + | 3.71 | 5.66E-05 | 4.34E-03 |
| neuronal action potential (GO:0019228) | 30 | 8 | 2.16 | + | 3.71 | 9.82E-04 | 3.74E-02 |
| response to electrical stimulus (GO:0051602) | 45 | 12 | 3.24 | + | 3.71 | 5.66E-05 | 4.32E-03 |
| gluconeogenesis (GO:0006094) | 49 | 13 | 3.52 | + | 3.69 | 2.97E-05 | 2.62E-03 |
| cerebellar cortex morphogenesis (GO:0021696) | 38 | 10 | 2.73 | + | 3.66 | 2.63E-04 | 1.38E-02 |
| 'de novo' protein folding (GO:0006458) | 42 | 11 | 3.02 | + | 3.64 | 1.37E-04 | 8.56E-03 |
| nucleoside diphosphate catabolic process (GO:0009134) | 58 | 15 | 4.17 | + | 3.6 | 1.03E-05 | 1.16E-03 |
| protein insertion into ER membrane (GO:0045048) | 31 | 8 | 2.23 | + | 3.59 | 1.24E-03 | 4.53E-02 |
| dopamine metabolic process (GO:0042417) | 31 | 8 | 2.23 | + | 3.59 | 1.24E-03 | 4.52E-02 |
| purine ribonucleoside diphosphate catabolic process (GO:0009181) | 51 | 13 | 3.67 | + | 3.54 | 4.70E-05 | 3.76E-03 |
| purine nucleoside diphosphate catabolic process (GO:0009137) | 51 | 13 | 3.67 | + | 3.54 | 4.70E-05 | 3.74E-03 |
| ADP metabolic process (GO:0046031) | 55 | 14 | 3.96 | + | 3.54 | 2.46E-05 | 2.26E-03 |
| potassium ion import across plasma membrane (GO:1990573) | 44 | 11 | 3.16 | + | 3.48 | 2.15E-04 | 1.18E-02 |
| hexose biosynthetic process (GO:0019319) | 52 | 13 | 3.74 | + | 3.48 | 5.86E-05 | 4.41E-03 |
| response to amine (GO:0014075) | 52 | 13 | 3.74 | + | 3.48 | 5.86E-05 | 4.38E-03 |

| | | | | | | | | |
|---|---|---|---|---|---|---|---|---|
| response to cocaine (GO:0042220) | 56 | 14 | 4.03 | + | | 3.48 | 3.07E-05 | 2.67E-03 |
| cerebellar cortex development (GO:0021695) | 53 | 13 | 3.81 | + | | 3.41 | 7.25E-05 | 4.98E-03 |
| positive regulation of axon extension (GO:0045773) | 37 | 9 | 2.66 | + | | 3.38 | 9.90E-04 | 3.74E-02 |
| 'de novo' post-translational protein folding (GO:0051084) | 37 | 9 | 2.66 | + | | 3.38 | 9.90E-04 | 3.73E-02 |
| pyruvate metabolic process (GO:0006090) | 70 | 17 | 5.03 | + | | 3.38 | 6.81E-06 | 8.23E-04 |
| NAD metabolic process (GO:0019674) | 62 | 15 | 4.46 | + | | 3.36 | 2.46E-05 | 2.27E-03 |
| ribonucleoside diphosphate catabolic process (GO:0009191) | 54 | 13 | 3.88 | + | | 3.35 | 8.93E-05 | 5.95E-03 |
| long-term synaptic potentiation (GO:0060291) | 50 | 12 | 3.6 | + | | 3.34 | 1.71E-04 | 9.93E-03 |
| regulation of catecholamine secretion (GO:0050433) | 42 | 10 | 3.02 | + | | 3.31 | 6.30E-04 | 2.66E-02 |
| positive regulation of synapse assembly (GO:0051965) | 64 | 15 | 4.6 | + | | 3.26 | 3.68E-05 | 3.14E-03 |
| neuron cellular homeostasis (GO:0070050) | 64 | 15 | 4.6 | + | | 3.26 | 3.68E-05 | 3.12E-03 |
| protein localization to synapse (GO:0035418) | 60 | 14 | 4.31 | + | | 3.24 | 6.99E-05 | 4.89E-03 |
| semaphorin-plexin signaling pathway (GO:0071526) | 43 | 10 | 3.09 | + | | 3.23 | 7.68E-04 | 3.13E-02 |
| monosaccharide biosynthetic process (GO:0046364) | 56 | 13 | 4.03 | + | | 3.23 | 1.33E-04 | 8.46E-03 |
| hexose catabolic process (GO:0019320) | 44 | 10 | 3.16 | + | | 3.16 | 9.31E-04 | 3.58E-02 |
| cerebellum development (GO:0021549) | 106 | 24 | 7.62 | + | | 3.15 | 3.80E-07 | 7.08E-05 |
| receptor localization to synapse (GO:0097120) | 45 | 10 | 3.24 | + | | 3.09 | 1.12E-03 | 4.16E-02 |
| purine-containing compound catabolic process (GO:0072523) | 105 | 23 | 7.55 | + | | 3.05 | 1.22E-06 | 2.01E-04 |
| synapse assembly (GO:0007416) | 119 | 26 | 8.56 | + | | 3.04 | 2.67E-07 | 5.23E-05 |

| | | | | | | | |
|---|---|---|---|---|---|---|---|
| positive regulation of insulin secretion (GO:0032024) | 87 | 19 | 6.26 | + | 3.04 | 1.06E-05 | 1.19E-03 |
| neural crest cell migration (GO:0001755) | 55 | 12 | 3.96 | + | 3.03 | 4.42E-04 | 2.04E-02 |
| regulation of potassium ion transport (GO:0043266) | 92 | 20 | 6.62 | + | 3.02 | 6.73E-06 | 8.21E-04 |
| transmission of nerve impulse (GO:0019226) | 60 | 13 | 4.31 | + | 3.01 | 2.78E-04 | 1.44E-02 |
| proteoglycan biosynthetic process (GO:0030166) | 60 | 13 | 4.31 | + | 3.01 | 2.78E-04 | 1.43E-02 |
| sensory perception of pain (GO:0019233) | 65 | 14 | 4.67 | + | 3 | 1.75E-04 | 1.02E-02 |
| mesenchymal cell migration (GO:0090497) | 57 | 12 | 4.1 | + | 2.93 | 6.23E-04 | 2.64E-02 |
| purine nucleotide catabolic process (GO:0006195) | 100 | 21 | 7.19 | + | 2.92 | 7.11E-06 | 8.46E-04 |
| purine ribonucleoside diphosphate metabolic process (GO:0009179) | 67 | 14 | 4.82 | + | 2.91 | 2.46E-04 | 1.31E-02 |
| purine nucleoside diphosphate metabolic process (GO:0009135) | 67 | 14 | 4.82 | + | 2.91 | 2.46E-04 | 1.30E-02 |
| metencephalon development (GO:0022037) | 116 | 24 | 8.34 | + | 2.88 | 2.14E-06 | 3.26E-04 |
| regulation of cytosolic calcium ion concentration (GO:0051480) | 58 | 12 | 4.17 | + | 2.88 | 7.35E-04 | 3.04E-02 |
| regulation of neuronal synaptic plasticity (GO:0048168) | 58 | 12 | 4.17 | + | 2.88 | 7.35E-04 | 3.03E-02 |
| positive regulation of peptide hormone secretion (GO:0090277) | 112 | 23 | 8.05 | + | 2.86 | 3.95E-06 | 5.53E-04 |
| multicellular organismal response to stress (GO:0033555) | 88 | 18 | 6.33 | + | 2.84 | 4.56E-05 | 3.69E-03 |
| negative regulation of leukocyte apoptotic process (GO:2000107) | 54 | 11 | 3.88 | + | 2.83 | 1.38E-03 | 4.95E-02 |
| regulation of receptor internalization (GO:0002090) | 64 | 13 | 4.6 | + | 2.82 | 5.40E-04 | 2.35E-02 |
| inorganic ion import across plasma membrane (GO:0099587) | 104 | 21 | 7.48 | + | 2.81 | 1.35E-05 | 1.42E-03 |
| inorganic cation import across plasma membrane (GO:0098659) | 104 | 21 | 7.48 | + | 2.81 | 1.35E-05 | 1.41E-03 |

| | | | | | | | |
|---|---|---|---|---|---|---|---|
| positive regulation of peptide secretion (GO:0002793) | 114 | 23 | 8.2 | + | 2.81 | 5.41E-06 | 6.93E-04 |
| purine ribonucleotide catabolic process (GO:0009154) | 91 | 18 | 6.54 | + | 2.75 | 7.25E-05 | 5.00E-03 |
| nucleoside phosphate catabolic process (GO:1901292) | 132 | 26 | 9.49 | + | 2.74 | 2.17E-06 | 3.29E-04 |
| regulation of synaptic plasticity (GO:0048167) | 204 | 40 | 14.67 | + | 2.73 | 5.12E-09 | 1.52E-06 |
| protein localization to cell junction (GO:1902414) | 87 | 17 | 6.26 | + | 2.72 | 1.33E-04 | 8.44E-03 |
| regulation of sodium ion transport (GO:0002028) | 82 | 16 | 5.9 | + | 2.71 | 2.12E-04 | 1.17E-02 |
| nucleotide catabolic process (GO:0009166) | 123 | 24 | 8.85 | + | 2.71 | 6.26E-06 | 7.89E-04 |
| pyridine nucleotide metabolic process (GO:0019362) | 118 | 23 | 8.49 | + | 2.71 | 9.88E-06 | 1.13E-03 |
| nicotinamide nucleotide metabolic process (GO:0046496) | 118 | 23 | 8.49 | + | 2.71 | 9.88E-06 | 1.12E-03 |
| regulation of potassium ion transmembrane transport (GO:1901379) | 77 | 15 | 5.54 | + | 2.71 | 3.36E-04 | 1.66E-02 |
| positive regulation of axonogenesis (GO:0050772) | 77 | 15 | 5.54 | + | 2.71 | 3.36E-04 | 1.66E-02 |
| regulation of amine transport (GO:0051952) | 83 | 16 | 5.97 | + | 2.68 | 2.45E-04 | 1.31E-02 |
| neural crest cell development (GO:0014032) | 78 | 15 | 5.61 | + | 2.67 | 3.89E-04 | 1.84E-02 |
| regulation of axon extension (GO:0030516) | 78 | 15 | 5.61 | + | 2.67 | 3.89E-04 | 1.84E-02 |
| carbohydrate catabolic process (GO:0016052) | 120 | 23 | 8.63 | + | 2.67 | 1.32E-05 | 1.40E-03 |
| hindbrain development (GO:0030902) | 157 | 30 | 11.29 | + | 2.66 | 7.29E-07 | 1.30E-04 |
| nucleoside diphosphate metabolic process (GO:0009132) | 84 | 16 | 6.04 | + | 2.65 | 2.83E-04 | 1.44E-02 |
| positive regulation of synaptic transmission (GO:0050806) | 143 | 27 | 10.28 | + | 2.63 | 3.26E-06 | 4.65E-04 |
| response to alkaloid (GO:0043279) | 101 | 19 | 7.26 | + | 2.62 | 9.42E-05 | 6.19E-03 |

| | | | | | | | |
|---|---|---|---|---|---|---|---|
| synapse organization (GO:0050808) | 337 | 63 | 24.24 | + | 2.6 | 1.93E-12 | 1.08E-09 |
| chaperone-mediated protein folding (GO:0061077) | 75 | 14 | 5.39 | + | 2.6 | 8.21E-04 | 3.26E-02 |
| antigen processing and presentation of peptide antigen (GO:0048002) | 70 | 13 | 5.03 | + | 2.58 | 1.31E-03 | 4.73E-02 |
| nerve development (GO:0021675) | 97 | 18 | 6.98 | + | 2.58 | 1.71E-04 | 1.00E-02 |
| ribonucleotide catabolic process (GO:0009261) | 97 | 18 | 6.98 | + | 2.58 | 1.71E-04 | 9.97E-03 |
| glucose metabolic process (GO:0006006) | 119 | 22 | 8.56 | + | 2.57 | 3.61E-05 | 3.10E-03 |
| organophosphate catabolic process (GO:0046434) | 184 | 34 | 13.23 | + | 2.57 | 3.13E-07 | 6.06E-05 |
| action potential (GO:0001508) | 125 | 23 | 8.99 | + | 2.56 | 2.62E-05 | 2.39E-03 |
| pyridine-containing compound metabolic process (GO:0072524) | 125 | 23 | 8.99 | + | 2.56 | 2.62E-05 | 2.37E-03 |
| vesicle-mediated transport in synapse (GO:0099003) | 154 | 28 | 11.07 | + | 2.53 | 4.65E-06 | 6.33E-04 |
| ribonucleoside diphosphate metabolic process (GO:0009185) | 77 | 14 | 5.54 | + | 2.53 | 1.07E-03 | 4.01E-02 |
| positive regulation of exocytosis (GO:0045921) | 77 | 14 | 5.54 | + | 2.53 | 1.07E-03 | 4.00E-02 |
| synaptic vesicle cycle (GO:0099504) | 144 | 26 | 10.36 | + | 2.51 | 1.14E-05 | 1.25E-03 |
| positive regulation of hormone secretion (GO:0046887) | 145 | 26 | 10.43 | + | 2.49 | 1.30E-05 | 1.39E-03 |
| regulation of extent of cell growth (GO:0061387) | 95 | 17 | 6.83 | + | 2.49 | 4.00E-04 | 1.87E-02 |
| stem cell development (GO:0048864) | 84 | 15 | 6.04 | + | 2.48 | 8.79E-04 | 3.41E-02 |
| regulation of G protein-coupled receptor signaling pathway (GO:0008277) | 157 | 28 | 11.29 | + | 2.48 | 6.82E-06 | 8.18E-04 |
| trans-synaptic signaling (GO:0099537) | 432 | 77 | 31.07 | + | 2.48 | 9.66E-14 | 7.31E-11 |
| regulation of neurotransmitter transport (GO:0051588) | 101 | 18 | 7.26 | + | 2.48 | 2.89E-04 | 1.46E-02 |

| | | | | | | | |
|---|---|---|---|---|---|---|---|
| regulation of axonogenesis (GO:0050770) | 141 | 25 | 10.14 | + | 2.47 | 2.31E-05 | 2.18E-03 |
| regulation of trans-synaptic signaling (GO:0099177) | 480 | 85 | 34.52 | + | 2.46 | 6.78E-15 | 1.28E-11 |
| hexose metabolic process (GO:0019318) | 159 | 28 | 11.43 | + | 2.45 | 8.75E-06 | 1.01E-03 |
| memory (GO:0007613) | 125 | 22 | 8.99 | + | 2.45 | 7.82E-05 | 5.33E-03 |
| chemical synaptic transmission (GO:0007268) | 415 | 73 | 29.84 | + | 2.45 | 8.09E-13 | 5.10E-10 |
| anterograde trans-synaptic signaling (GO:0098916) | 415 | 73 | 29.84 | + | 2.45 | 8.09E-13 | 4.89E-10 |
| modulation of chemical synaptic transmission (GO:0050804) | 479 | 84 | 34.45 | + | 2.44 | 2.45E-14 | 2.64E-11 |
| synaptic signaling (GO:0099536) | 462 | 81 | 33.22 | + | 2.44 | 7.77E-14 | 6.53E-11 |
| positive regulation of cell junction assembly (GO:1901890) | 109 | 19 | 7.84 | + | 2.42 | 2.67E-04 | 1.39E-02 |
| ATP metabolic process (GO:0046034) | 162 | 28 | 11.65 | + | 2.4 | 1.79E-05 | 1.78E-03 |
| regulation of synapse structure or activity (GO:0050803) | 255 | 44 | 18.34 | + | 2.4 | 5.50E-08 | 1.36E-05 |
| neurotransmitter transport (GO:0006836) | 140 | 24 | 10.07 | + | 2.38 | 8.65E-05 | 5.79E-03 |
| monosaccharide metabolic process (GO:0005996) | 181 | 31 | 13.02 | + | 2.38 | 6.64E-06 | 8.22E-04 |
| import across plasma membrane (GO:0098739) | 164 | 28 | 11.79 | + | 2.37 | 2.05E-05 | 1.98E-03 |
| positive regulation of protein secretion (GO:0050714) | 147 | 25 | 10.57 | + | 2.36 | 6.40E-05 | 4.65E-03 |
| regulation of synapse organization (GO:0050807) | 249 | 42 | 17.91 | + | 2.35 | 2.00E-07 | 4.03E-05 |
| positive regulation of secretion by cell (GO:1903532) | 291 | 49 | 20.93 | + | 2.34 | 2.75E-08 | 6.93E-06 |
| axonogenesis (GO:0007409) | 366 | 61 | 26.32 | + | 2.32 | 6.04E-10 | 2.28E-07 |
| neuromuscular process (GO:0050905) | 150 | 25 | 10.79 | + | 2.32 | 8.01E-05 | 5.43E-03 |

| | | | | | | | |
|---|---|---|---|---|---|---|---|
| regulation of synapse assembly (GO:0051963) | 121 | 20 | 8.7 | + | 2.3 | 5.02E-04 | 2.24E-02 |
| detection of abiotic stimulus (GO:0009582) | 135 | 22 | 9.71 | + | 2.27 | 2.89E-04 | 1.46E-02 |
| sulfur compound biosynthetic process (GO:0044272) | 160 | 26 | 11.51 | + | 2.26 | 8.10E-05 | 5.47E-03 |
| neuropeptide signaling pathway (GO:0007218) | 111 | 18 | 7.98 | + | 2.25 | 1.18E-03 | 4.33E-02 |
| regulation of exocytosis (GO:0017157) | 173 | 28 | 12.44 | + | 2.25 | 4.57E-05 | 3.68E-03 |
| establishment of protein localization to extracellular region (GO:0035592) | 136 | 22 | 9.78 | + | 2.25 | 3.15E-04 | 1.56E-02 |
| potassium ion transmembrane transport (GO:0071805) | 167 | 27 | 12.01 | + | 2.25 | 6.38E-05 | 4.66E-03 |
| purine ribonucleoside triphosphate metabolic process (GO:0009205) | 186 | 30 | 13.38 | + | 2.24 | 3.88E-05 | 3.26E-03 |
| purine nucleoside triphosphate metabolic process (GO:0009144) | 193 | 31 | 13.88 | + | 2.23 | 2.86E-05 | 2.56E-03 |
| cell morphogenesis involved in neuron differentiation (GO:0048667) | 443 | 71 | 31.86 | + | 2.23 | 1.60E-10 | 6.74E-08 |
| positive regulation of neuron projection development (GO:0010976) | 156 | 25 | 11.22 | + | 2.23 | 1.37E-04 | 8.60E-03 |
| endoplasmic reticulum to Golgi vesicle-mediated transport (GO:0006888) | 125 | 20 | 8.99 | + | 2.22 | 6.80E-04 | 2.84E-02 |
| positive regulation of secretion (GO:0051047) | 313 | 50 | 22.51 | + | 2.22 | 1.01E-07 | 2.35E-05 |
| detection of external stimulus (GO:0009581) | 132 | 21 | 9.49 | + | 2.21 | 5.25E-04 | 2.30E-02 |
| sensory perception of mechanical stimulus (GO:0050954) | 183 | 29 | 13.16 | + | 2.2 | 6.44E-05 | 4.66E-03 |
| protein secretion (GO:0009306) | 133 | 21 | 9.56 | + | 2.2 | 5.73E-04 | 2.48E-02 |
| potassium ion transport (GO:0006813) | 184 | 29 | 13.23 | + | 2.19 | 6.84E-05 | 4.85E-03 |
| axon development (GO:0061564) | 420 | 66 | 30.2 | + | 2.19 | 1.53E-09 | 5.02E-07 |
| regulation of monoatomic ion transmembrane transporter activity (GO:0032412) | 198 | 31 | 14.24 | + | 2.18 | 3.96E-05 | 3.31E-03 |

| | | | | | | | |
|---|---|---|---|---|---|---|---|
| ribonucleoside triphosphate metabolic process (GO:0009199) | 193 | 30 | 13.88 | + | 2.16 | 5.98E-05 | 4.43E-03 |
| protein localization to extracellular region (GO:0071692) | 142 | 22 | 10.21 | + | 2.15 | 8.03E-04 | 3.20E-02 |
| cellular response to metal ion (GO:0071248) | 188 | 29 | 13.52 | + | 2.14 | 9.02E-05 | 5.98E-03 |
| purine ribonucleotide biosynthetic process (GO:0009152) | 195 | 30 | 14.02 | + | 2.14 | 6.98E-05 | 4.91E-03 |
| regulation of neuron projection development (GO:0010975) | 437 | 67 | 31.43 | + | 2.13 | 3.31E-09 | 1.02E-06 |
| nucleoside triphosphate metabolic process (GO:0009141) | 209 | 32 | 15.03 | + | 2.13 | 6.32E-05 | 4.66E-03 |
| ribonucleotide biosynthetic process (GO:0009260) | 210 | 32 | 15.1 | + | 2.12 | 6.56E-05 | 4.68E-03 |
| locomotory behavior (GO:0007626) | 197 | 30 | 14.17 | + | 2.12 | 8.20E-05 | 5.51E-03 |
| ribose phosphate biosynthetic process (GO:0046390) | 217 | 33 | 15.61 | + | 2.11 | 4.87E-05 | 3.85E-03 |
| intracellular calcium ion homeostasis (GO:0006874) | 199 | 30 | 14.31 | + | 2.1 | 1.37E-04 | 8.57E-03 |
| regulation of transporter activity (GO:0032409) | 219 | 33 | 15.75 | + | 2.1 | 5.41E-05 | 4.24E-03 |
| regulation of transmembrane transporter activity (GO:0022898) | 206 | 31 | 14.81 | + | 2.09 | 1.01E-04 | 6.60E-03 |
| cell junction organization (GO:0034330) | 533 | 80 | 38.33 | + | 2.09 | 4.20E-10 | 1.63E-07 |
| carbohydrate derivative catabolic process (GO:1901136) | 222 | 33 | 15.97 | + | 2.07 | 6.53E-05 | 4.68E-03 |
| neuron projection morphogenesis (GO:0048812) | 485 | 72 | 34.88 | + | 2.06 | 3.82E-09 | 1.15E-06 |
| learning (GO:0007612) | 155 | 23 | 11.15 | + | 2.06 | 8.39E-04 | 3.30E-02 |
| cell projection morphogenesis (GO:0048858) | 495 | 73 | 35.6 | + | 2.05 | 5.20E-09 | 1.51E-06 |
| cell junction assembly (GO:0034329) | 285 | 42 | 20.5 | + | 2.05 | 8.22E-06 | 9.64E-04 |
| regulation of heart contraction (GO:0008016) | 197 | 29 | 14.17 | + | 2.05 | 2.33E-04 | 1.26E-02 |

| | | | | | | | |
|---|---|---|---|---|---|---|---|
| plasma membrane bounded cell projection morphogenesis (GO:0120039) | 490 | 72 | 35.24 | + | 2.04 | 7.55E-09 | 2.07E-06 |
| regulation of membrane potential (GO:0042391) | 456 | 67 | 32.79 | + | 2.04 | 2.10E-08 | 5.39E-06 |
| positive regulation of monoatomic ion transport (GO:0043270) | 205 | 30 | 14.74 | + | 2.03 | 1.88E-04 | 1.07E-02 |
| regulation of insulin secretion (GO:0050796) | 171 | 25 | 12.3 | + | 2.03 | 8.23E-04 | 3.26E-02 |
| purine-containing compound biosynthetic process (GO:0072522) | 240 | 35 | 17.26 | + | 2.03 | 6.94E-05 | 4.91E-03 |
| positive regulation of nervous system development (GO:0051962) | 291 | 42 | 20.93 | + | 2.01 | 1.63E-05 | 1.65E-03 |
| regulation of monoatomic ion transmembrane transport (GO:0034765) | 333 | 48 | 23.95 | + | 2 | 4.68E-06 | 6.32E-04 |
| protein folding (GO:0006457) | 223 | 32 | 16.04 | + | 2 | 1.97E-04 | 1.10E-02 |
| purine nucleotide biosynthetic process (GO:0006164) | 231 | 33 | 16.61 | + | 1.99 | 1.56E-04 | 9.34E-03 |
| axon guidance (GO:0007411) | 224 | 32 | 16.11 | + | 1.99 | 2.06E-04 | 1.15E-02 |
| neuron projection guidance (GO:0097485) | 225 | 32 | 16.18 | + | 1.98 | 2.16E-04 | 1.18E-02 |
| regulation of peptide transport (GO:0090087) | 204 | 29 | 14.67 | + | 1.98 | 5.14E-04 | 2.27E-02 |
| regulation of peptide secretion (GO:0002791) | 204 | 29 | 14.67 | + | 1.98 | 5.14E-04 | 2.26E-02 |
| regulation of cell junction assembly (GO:1901888) | 219 | 31 | 15.75 | + | 1.97 | 2.99E-04 | 1.50E-02 |
| purine nucleotide metabolic process (GO:0006163) | 468 | 66 | 33.66 | + | 1.96 | 1.54E-07 | 3.32E-05 |
| purine ribonucleotide metabolic process (GO:0009150) | 386 | 54 | 27.76 | + | 1.95 | 3.30E-06 | 4.66E-04 |
| regulation of peptide hormone secretion (GO:0090276) | 201 | 28 | 14.45 | + | 1.94 | 7.99E-04 | 3.19E-02 |
| export from cell (GO:0140352) | 490 | 68 | 35.24 | + | 1.93 | 1.83E-07 | 3.74E-05 |
| purine-containing compound metabolic process (GO:0072521) | 498 | 69 | 35.81 | + | 1.93 | 1.44E-07 | 3.16E-05 |

| | | | | | | | |
|---|---|---|---|---|---|---|---|
| cell morphogenesis (GO:0000902) | 686 | 95 | 49.33 | + | 1.93 | 6.24E-10 | 2.30E-07 |
| ribonucleotide metabolic process (GO:0009259) | 406 | 56 | 29.2 | + | 1.92 | 2.39E-06 | 3.57E-04 |
| ribose phosphate metabolic process (GO:0019693) | 414 | 57 | 29.77 | + | 1.91 | 2.79E-06 | 4.09E-04 |
| vascular process in circulatory system (GO:0003018) | 269 | 37 | 19.34 | + | 1.91 | 1.81E-04 | 1.04E-02 |
| behavior (GO:0007610) | 618 | 85 | 44.44 | + | 1.91 | 7.48E-09 | 2.09E-06 |
| regulation of transmembrane transport (GO:0034762) | 459 | 63 | 33.01 | + | 1.91 | 8.73E-07 | 1.50E-04 |
| regulation of secretion by cell (GO:1903530) | 547 | 75 | 39.34 | + | 1.91 | 8.10E-08 | 1.94E-05 |
| regulation of secretion (GO:0051046) | 600 | 82 | 43.15 | + | 1.9 | 1.94E-08 | 5.05E-06 |
| calcium ion homeostasis (GO:0055074) | 227 | 31 | 16.32 | + | 1.9 | 6.35E-04 | 2.67E-02 |
| secretion by cell (GO:0032940) | 425 | 58 | 30.56 | + | 1.9 | 2.41E-06 | 3.58E-04 |
| protein homooligomerization (GO:0051260) | 198 | 27 | 14.24 | + | 1.9 | 1.26E-03 | 4.59E-02 |
| cell-cell signaling (GO:0007267) | 831 | 113 | 59.76 | + | 1.89 | 3.96E-11 | 1.76E-08 |
| regulation of monoatomic ion transport (GO:0043269) | 449 | 61 | 32.29 | + | 1.89 | 1.74E-06 | 2.74E-04 |
| neuron projection development (GO:0031175) | 692 | 94 | 49.76 | + | 1.89 | 2.02E-09 | 6.36E-07 |
| neuron differentiation (GO:0030182) | 1082 | 146 | 77.81 | + | 1.88 | 7.43E-14 | 6.61E-11 |
| learning or memory (GO:0007611) | 276 | 37 | 19.85 | + | 1.86 | 2.33E-04 | 1.25E-02 |
| generation of neurons (GO:0048699) | 1160 | 154 | 83.42 | + | 1.85 | 5.28E-14 | 4.99E-11 |
| neuron development (GO:0048666) | 859 | 114 | 61.77 | + | 1.85 | 1.38E-10 | 5.94E-08 |
| regulation of blood circulation (GO:1903522) | 249 | 33 | 17.91 | + | 1.84 | 7.20E-04 | 2.99E-02 |

| | | | | | | | |
|---|---|---|---|---|---|---|---|
| regulation of metal ion transport (GO:0010959) | 378 | 50 | 27.18 | + | 1.84 | 2.99E-05 | 2.63E-03 |
| positive regulation of protein transport (GO:0051222) | 250 | 33 | 17.98 | + | 1.84 | 7.44E-04 | 3.06E-02 |
| exocytosis (GO:0006887) | 243 | 32 | 17.48 | + | 1.83 | 9.82E-04 | 3.75E-02 |
| generation of precursor metabolites and energy (GO:0006091) | 373 | 49 | 26.82 | + | 1.83 | 4.21E-05 | 3.44E-03 |
| muscle contraction (GO:0006936) | 236 | 31 | 16.97 | + | 1.83 | 1.30E-03 | 4.72E-02 |
| nucleotide biosynthetic process (GO:0009165) | 267 | 35 | 19.2 | + | 1.82 | 7.12E-04 | 2.97E-02 |
| neurogenesis (GO:0022008) | 1336 | 175 | 96.08 | + | 1.82 | 3.33E-15 | 7.19E-12 |
| protein localization to cell periphery (GO:1990778) | 237 | 31 | 17.04 | + | 1.82 | 1.34E-03 | 4.82E-02 |
| nucleoside phosphate metabolic process (GO:0006753) | 536 | 70 | 38.55 | + | 1.82 | 1.07E-06 | 1.80E-04 |
| positive regulation of transport (GO:0051050) | 835 | 109 | 60.05 | + | 1.82 | 1.11E-09 | 3.72E-07 |
| positive regulation of establishment of protein localization (GO:1904951) | 330 | 43 | 23.73 | + | 1.81 | 1.45E-04 | 8.93E-03 |
| nucleoside phosphate biosynthetic process (GO:1901293) | 269 | 35 | 19.34 | + | 1.81 | 7.44E-04 | 3.05E-02 |
| nucleotide metabolic process (GO:0009117) | 528 | 68 | 37.97 | + | 1.79 | 3.10E-06 | 4.47E-04 |
| cognition (GO:0050890) | 320 | 41 | 23.01 | + | 1.78 | 2.86E-04 | 1.46E-02 |
| regulation of monoatomic cation transmembrane transport (GO:1904062) | 297 | 38 | 21.36 | + | 1.78 | 5.88E-04 | 2.52E-02 |
| import into cell (GO:0098657) | 690 | 88 | 49.62 | + | 1.77 | 1.69E-07 | 3.51E-05 |
| central nervous system development (GO:0007417) | 1000 | 127 | 71.91 | + | 1.77 | 2.28E-10 | 9.06E-08 |
| regulation of nervous system development (GO:0051960) | 450 | 57 | 32.36 | + | 1.76 | 2.80E-05 | 2.52E-03 |
| phospholipid metabolic process (GO:0006644) | 364 | 46 | 26.18 | + | 1.76 | 1.95E-04 | 1.10E-02 |

| | | | | | | | |
|---|---|---|---|---|---|---|---|
| nucleobase-containing small molecule metabolic process (GO:0055086) | 605 | 76 | 43.51 | + | 1.75 | 1.92E-06 | 2.96E-04 |
| organophosphate metabolic process (GO:0019637) | 940 | 118 | 67.6 | + | 1.75 | 1.99E-09 | 6.40E-07 |
| regulation of vesicle-mediated transport (GO:0060627) | 510 | 64 | 36.68 | + | 1.74 | 1.21E-05 | 1.30E-03 |
| nervous system development (GO:0007399) | 2193 | 273 | 157.71 | + | 1.73 | 1.08E-20 | 1.63E-16 |
| positive regulation of cell projection organization (GO:0031346) | 354 | 44 | 25.46 | + | 1.73 | 3.70E-04 | 1.79E-02 |
| response to metal ion (GO:0010038) | 347 | 43 | 24.95 | + | 1.72 | 4.79E-04 | 2.15E-02 |
| organophosphate biosynthetic process (GO:0090407) | 549 | 68 | 39.48 | + | 1.72 | 1.10E-05 | 1.22E-03 |
| inorganic ion transmembrane transport (GO:0098660) | 733 | 90 | 52.71 | + | 1.71 | 5.82E-07 | 1.06E-04 |
| carbohydrate derivative biosynthetic process (GO:1901137) | 604 | 74 | 43.44 | + | 1.7 | 6.33E-06 | 7.91E-04 |
| circulatory system process (GO:0003013) | 501 | 61 | 36.03 | + | 1.69 | 4.97E-05 | 3.91E-03 |
| regulation of cell projection organization (GO:0031344) | 649 | 79 | 46.67 | + | 1.69 | 4.28E-06 | 5.88E-04 |
| regulation of plasma membrane bounded cell projection organization (GO:0120035) | 633 | 77 | 45.52 | + | 1.69 | 5.02E-06 | 6.60E-04 |
| carbohydrate metabolic process (GO:0005975) | 444 | 54 | 31.93 | + | 1.69 | 1.76E-04 | 1.02E-02 |
| regulation of transport (GO:0051049) | 1576 | 191 | 113.34 | + | 1.69 | 3.53E-13 | 2.32E-10 |
| secretion (GO:0046903) | 545 | 66 | 39.19 | + | 1.68 | 3.25E-05 | 2.81E-03 |
| regulation of establishment of protein localization (GO:0070201) | 529 | 64 | 38.04 | + | 1.68 | 3.82E-05 | 3.23E-03 |
| monoatomic cation transmembrane transport (GO:0098655) | 653 | 79 | 46.96 | + | 1.68 | 4.76E-06 | 6.37E-04 |
| inorganic cation transmembrane transport (GO:0098662) | 638 | 77 | 45.88 | + | 1.68 | 7.61E-06 | 8.99E-04 |
| protein localization to membrane (GO:0072657) | 459 | 55 | 33.01 | + | 1.67 | 2.26E-04 | 1.23E-02 |

| | | | | | | | |
|---|---|---|---|---|---|---|---|
| phosphate-containing compound metabolic process (GO:0006796) | 1579 | 188 | 113.55 | + | 1.66 | 2.49E-12 | 1.30E-09 |
| regulation of anatomical structure size (GO:0090066) | 496 | 59 | 35.67 | + | 1.65 | 1.37E-04 | 8.55E-03 |
| phosphorus metabolic process (GO:0006793) | 1610 | 191 | 115.78 | + | 1.65 | 1.98E-12 | 1.07E-09 |
| monoatomic ion transmembrane transport (GO:0034220) | 815 | 96 | 58.61 | + | 1.64 | 1.52E-06 | 2.42E-04 |
| regulation of hormone levels (GO:0010817) | 544 | 64 | 39.12 | + | 1.64 | 9.75E-05 | 6.38E-03 |
| monoatomic ion transport (GO:0006811) | 980 | 115 | 70.48 | + | 1.63 | 1.65E-07 | 3.46E-05 |
| blood circulation (GO:0008015) | 411 | 48 | 29.56 | + | 1.62 | 9.48E-04 | 3.63E-02 |
| regulation of biological quality (GO:0065008) | 2822 | 328 | 202.94 | + | 1.62 | 4.32E-20 | 3.27E-16 |
| regulation of localization (GO:0032879) | 1984 | 230 | 142.68 | + | 1.61 | 8.59E-14 | 6.83E-11 |
| protein maturation (GO:0051604) | 494 | 57 | 35.53 | + | 1.6 | 3.89E-04 | 1.83E-02 |
| brain development (GO:0007420) | 730 | 84 | 52.5 | + | 1.6 | 2.03E-05 | 1.98E-03 |
| forebrain development (GO:0030900) | 409 | 47 | 29.41 | + | 1.6 | 1.35E-03 | 4.85E-02 |
| regulation of protein transport (GO:0051223) | 427 | 49 | 30.71 | + | 1.6 | 1.19E-03 | 4.38E-02 |
| monoatomic cation transport (GO:0006812) | 787 | 90 | 56.6 | + | 1.59 | 1.13E-05 | 1.25E-03 |
| carbohydrate derivative metabolic process (GO:1901135) | 997 | 114 | 71.7 | + | 1.59 | 7.99E-07 | 1.40E-04 |
| head development (GO:0060322) | 780 | 89 | 56.09 | + | 1.59 | 1.43E-05 | 1.47E-03 |
| vesicle-mediated transport (GO:0016192) | 1298 | 148 | 93.34 | + | 1.59 | 1.66E-08 | 4.40E-06 |
| plasma membrane bounded cell projection organization (GO:0120036) | 1148 | 129 | 82.56 | + | 1.56 | 3.62E-07 | 6.83E-05 |
| cellular response to hormone stimulus (GO:0032870) | 508 | 57 | 36.53 | + | 1.56 | 8.80E-04 | 3.40E-02 |

| | | | | | | | |
|---|---|---|---|---|---|---|---|
| cellular homeostasis (GO:0019725) | 679 | 76 | 48.83 | + | 1.56 | 1.09E-04 | 7.10E-03 |
| metal ion transport (GO:0030001) | 646 | 72 | 46.46 | + | 1.55 | 1.92E-04 | 1.09E-02 |
| endocytosis (GO:0006897) | 503 | 56 | 36.17 | + | 1.55 | 1.14E-03 | 4.23E-02 |
| regulation of protein localization (GO:0032880) | 868 | 96 | 62.42 | + | 1.54 | 2.16E-05 | 2.06E-03 |
| cell projection organization (GO:0030030) | 1196 | 132 | 86.01 | + | 1.53 | 6.17E-07 | 1.11E-04 |
| membrane organization (GO:0061024) | 744 | 82 | 53.5 | + | 1.53 | 1.16E-04 | 7.49E-03 |
| cellular response to nitrogen compound (GO:1901699) | 573 | 63 | 41.21 | + | 1.53 | 7.31E-04 | 3.03E-02 |
| localization within membrane (GO:0051668) | 548 | 60 | 39.41 | + | 1.52 | 1.04E-03 | 3.92E-02 |
| positive regulation of cellular component organization (GO:0051130) | 1115 | 121 | 80.18 | + | 1.51 | 5.27E-06 | 6.86E-04 |
| regulation of MAPK cascade (GO:0043408) | 636 | 69 | 45.74 | + | 1.51 | 5.77E-04 | 2.49E-02 |
| regulation of cellular localization (GO:0060341) | 978 | 106 | 70.33 | + | 1.51 | 1.97E-05 | 1.93E-03 |
| cellular catabolic process (GO:0044248) | 788 | 85 | 56.67 | + | 1.5 | 1.80E-04 | 1.03E-02 |
| transmembrane transport (GO:0055085) | 1283 | 138 | 92.27 | + | 1.5 | 1.43E-06 | 2.30E-04 |
| regulation of anatomical structure morphogenesis (GO:0022603) | 828 | 89 | 59.55 | + | 1.49 | 1.47E-04 | 9.04E-03 |
| oxoacid metabolic process (GO:0043436) | 813 | 86 | 58.47 | + | 1.47 | 2.99E-04 | 1.51E-02 |
| response to hormone (GO:0009725) | 787 | 83 | 56.6 | + | 1.47 | 4.15E-04 | 1.93E-02 |
| response to organonitrogen compound (GO:0010243) | 902 | 95 | 64.87 | + | 1.46 | 1.63E-04 | 9.70E-03 |
| system development (GO:0048731) | 3527 | 371 | 253.64 | + | 1.46 | 1.03E-15 | 2.59E-12 |
| carboxylic acid metabolic process (GO:0019752) | 791 | 83 | 56.88 | + | 1.46 | 5.51E-04 | 2.39E-02 |

| | | | | | | | |
|---|---|---|---|---|---|---|---|
| regulation of cell motility (GO:2000145) | 1001 | 105 | 71.99 | + | 1.46 | 9.38E-05 | 6.19E-03 |
| organic acid metabolic process (GO:0006082) | 820 | 86 | 58.97 | + | 1.46 | 4.11E-04 | 1.92E-02 |
| response to abiotic stimulus (GO:0009628) | 1100 | 115 | 79.11 | + | 1.45 | 4.26E-05 | 3.46E-03 |
| anatomical structure morphogenesis (GO:0009653) | 2220 | 230 | 159.65 | + | 1.44 | 6.28E-09 | 1.79E-06 |
| positive regulation of signaling (GO:0023056) | 1743 | 180 | 125.35 | + | 1.44 | 5.40E-07 | 9.95E-05 |
| cell migration (GO:0016477) | 915 | 94 | 65.8 | + | 1.43 | 4.95E-04 | 2.21E-02 |
| regulation of locomotion (GO:0040012) | 1042 | 107 | 74.93 | + | 1.43 | 1.67E-04 | 9.80E-03 |
| chemical homeostasis (GO:0048878) | 887 | 91 | 63.79 | + | 1.43 | 5.36E-04 | 2.34E-02 |
| transport (GO:0006810) | 3665 | 376 | 263.57 | + | 1.43 | 3.39E-14 | 3.42E-11 |
| regulation of signaling (GO:0023051) | 3423 | 351 | 246.16 | + | 1.43 | 3.44E-13 | 2.37E-10 |
| organonitrogen compound catabolic process (GO:1901565) | 1116 | 114 | 80.26 | + | 1.42 | 1.30E-04 | 8.31E-03 |
| response to nitrogen compound (GO:1901698) | 1009 | 103 | 72.56 | + | 1.42 | 2.79E-04 | 1.43E-02 |
| regulation of cell migration (GO:0030334) | 941 | 96 | 67.67 | + | 1.42 | 4.70E-04 | 2.11E-02 |
| response to endogenous stimulus (GO:0009719) | 1375 | 140 | 98.88 | + | 1.42 | 2.35E-05 | 2.18E-03 |
| regulation of cell communication (GO:0010646) | 3428 | 349 | 246.52 | + | 1.42 | 1.32E-12 | 7.68E-10 |
| positive regulation of cell communication (GO:0010647) | 1741 | 177 | 125.2 | + | 1.41 | 1.85E-06 | 2.89E-04 |
| multicellular organism development (GO:0007275) | 3945 | 401 | 283.7 | + | 1.41 | 1.23E-14 | 1.56E-11 |
| cellular response to endogenous stimulus (GO:0071495) | 1112 | 113 | 79.97 | + | 1.41 | 1.64E-04 | 9.70E-03 |
| cell development (GO:0048468) | 2226 | 225 | 160.08 | + | 1.41 | 8.19E-08 | 1.93E-05 |

| | | | | | | | |
|---|---|---|---|---|---|---|---|
| positive regulation of molecular function (GO:0044093) | 1140 | 115 | 81.98 | + | 1.4 | 1.95E-04 | 1.09E-02 |
| establishment of localization (GO:0051234) | 3935 | 395 | 282.98 | + | 1.4 | 1.67E-13 | 1.20E-10 |
| negative regulation of cell communication (GO:0010648) | 1400 | 140 | 100.68 | + | 1.39 | 5.58E-05 | 4.33E-03 |
| negative regulation of signaling (GO:0023057) | 1401 | 140 | 100.75 | + | 1.39 | 5.64E-05 | 4.35E-03 |
| small molecule metabolic process (GO:0044281) | 1629 | 162 | 117.15 | + | 1.38 | 2.03E-05 | 1.97E-03 |
| localization (GO:0051179) | 4498 | 445 | 323.47 | + | 1.38 | 1.76E-14 | 2.05E-11 |
| cellular macromolecule localization (GO:0070727) | 1958 | 193 | 140.81 | + | 1.37 | 4.04E-06 | 5.61E-04 |
| negative regulation of signal transduction (GO:0009968) | 1309 | 129 | 94.14 | + | 1.37 | 2.07E-04 | 1.15E-02 |
| protein localization (GO:0008104) | 1950 | 192 | 140.23 | + | 1.37 | 4.81E-06 | 6.38E-04 |
| homeostatic process (GO:0042592) | 1429 | 139 | 102.77 | + | 1.35 | 2.40E-04 | 1.29E-02 |
| regulation of molecular function (GO:0065009) | 1903 | 185 | 136.85 | + | 1.35 | 1.74E-05 | 1.74E-03 |
| regulation of protein metabolic process (GO:0051246) | 2110 | 205 | 151.74 | + | 1.35 | 5.41E-06 | 6.98E-04 |
| cellular localization (GO:0051641) | 2659 | 258 | 191.22 | + | 1.35 | 2.44E-07 | 4.84E-05 |
| regulation of cellular component organization (GO:0051128) | 2413 | 234 | 173.53 | + | 1.35 | 1.08E-06 | 1.79E-04 |
| establishment of localization in cell (GO:0051649) | 1628 | 157 | 117.08 | + | 1.34 | 1.39E-04 | 8.64E-03 |
| catabolic process (GO:0009056) | 1944 | 185 | 139.8 | + | 1.32 | 5.79E-05 | 4.38E-03 |
| response to oxygen-containing compound (GO:1901700) | 1507 | 143 | 108.38 | + | 1.32 | 5.13E-04 | 2.27E-02 |
| regulation of apoptotic process (GO:0042981) | 1459 | 138 | 104.92 | + | 1.32 | 7.55E-04 | 3.09E-02 |
| macromolecule localization (GO:0033036) | 2391 | 226 | 171.95 | + | 1.31 | 1.14E-05 | 1.24E-03 |

| | | | | | | | |
|---|---|---|---|---|---|---|---|
| negative regulation of response to stimulus (GO:0048585) | 1656 | 156 | 119.09 | + | 1.31 | 4.18E-04 | 1.94E-02 |
| regulation of signal transduction (GO:0009966) | 3017 | 284 | 216.97 | + | 1.31 | 8.14E-07 | 1.41E-04 |
| regulation of programmed cell death (GO:0043067) | 1501 | 141 | 107.94 | + | 1.31 | 8.85E-04 | 3.41E-02 |
| cellular response to chemical stimulus (GO:0070887) | 1908 | 179 | 137.21 | + | 1.3 | 1.92E-04 | 1.09E-02 |
| anatomical structure development (GO:0048856) | 5208 | 488 | 374.53 | + | 1.3 | 7.54E-12 | 3.68E-09 |
| regulation of intracellular signal transduction (GO:1902531) | 1940 | 181 | 139.51 | + | 1.3 | 2.15E-04 | 1.18E-02 |
| animal organ development (GO:0048513) | 2843 | 257 | 204.45 | + | 1.26 | 6.50E-05 | 4.68E-03 |
| organonitrogen compound metabolic process (GO:1901564) | 4641 | 418 | 333.76 | + | 1.25 | 1.13E-07 | 2.60E-05 |
| developmental process (GO:0032502) | 5711 | 514 | 410.7 | + | 1.25 | 1.07E-09 | 3.69E-07 |
| cellular component assembly (GO:0022607) | 2459 | 220 | 176.84 | + | 1.24 | 4.69E-04 | 2.12E-02 |
| multicellular organismal process (GO:0032501) | 6239 | 556 | 448.67 | + | 1.24 | 6.53E-10 | 2.35E-07 |
| regulation of multicellular organismal process (GO:0051239) | 2931 | 260 | 210.78 | + | 1.23 | 2.08E-04 | 1.15E-02 |
| cell communication (GO:0007154) | 5158 | 457 | 370.93 | + | 1.23 | 1.62E-07 | 3.44E-05 |
| positive regulation of cellular process (GO:0048522) | 5601 | 493 | 402.79 | + | 1.22 | 7.77E-08 | 1.90E-05 |
| cell differentiation (GO:0030154) | 3638 | 320 | 261.62 | + | 1.22 | 5.44E-05 | 4.24E-03 |
| positive regulation of biological process (GO:0048518) | 6177 | 543 | 444.22 | + | 1.22 | 1.07E-08 | 2.89E-06 |
| cellular developmental process (GO:0048869) | 3641 | 320 | 261.84 | + | 1.22 | 6.32E-05 | 4.64E-03 |
| signaling (GO:0023052) | 5116 | 448 | 367.91 | + | 1.22 | 9.27E-07 | 1.57E-04 |
| regulation of response to stimulus (GO:0048583) | 4010 | 349 | 288.38 | + | 1.21 | 5.71E-05 | 4.34E-03 |

| | | | | | | | |
|---|---|---|---|---|---|---|---|
| negative regulation of biological process (GO:0048519) | 5171 | 441 | 371.87 | + | 1.19 | 2.30E-05 | 2.19E-03 |
| negative regulation of cellular process (GO:0048523) | 4749 | 405 | 341.52 | + | 1.19 | 7.04E-05 | 4.91E-03 |
| cellular metabolic process (GO:0044237) | 5659 | 480 | 406.96 | + | 1.18 | 1.34E-05 | 1.41E-03 |
| cellular component organization (GO:0016043) | 5623 | 471 | 404.38 | + | 1.16 | 7.20E-05 | 5.00E-03 |
| regulation of biological process (GO:0050789) | 11711 | 965 | 842.19 | + | 1.15 | 1.55E-11 | 7.12E-09 |
| regulation of cellular process (GO:0050794) | 11103 | 911 | 798.47 | + | 1.14 | 1.06E-09 | 3.71E-07 |
| cellular component organization or biogenesis (GO:0071840) | 5838 | 478 | 419.84 | + | 1.14 | 5.75E-04 | 2.48E-02 |
| biological regulation (GO:0065007) | 12149 | 989 | 873.69 | + | 1.13 | 1.81E-10 | 7.39E-08 |
| cellular process (GO:0009987) | 14769 | 1198 | 1062.1 | + | 1.13 | 3.59E-17 | 1.81E-13 |
| biological_process (GO:0008150) | 17791 | 1371 | 1279.43 | + | 1.07 | 1.08E-14 | 1.64E-11 |
| regulation of nucleobase-containing compound metabolic process (GO:0019219) | 3961 | 238 | 284.85 | - | 0.84 | 1.15E-03 | 4.23E-02 |
| regulation of DNA-templated transcription (GO:0006355) | 3383 | 199 | 243.29 | - | 0.82 | 1.05E-03 | 3.92E-02 |
| regulation of RNA metabolic process (GO:0051252) | 3657 | 215 | 262.99 | - | 0.82 | 6.11E-04 | 2.60E-02 |
| regulation of RNA biosynthetic process (GO:2001141) | 3402 | 199 | 244.65 | - | 0.81 | 8.23E-04 | 3.25E-02 |
| regulation of transcription by RNA polymerase II (GO:0006357) | 2584 | 133 | 185.83 | - | 0.72 | 8.75E-06 | 1.02E-03 |
| negative regulation of RNA metabolic process (GO:0051253) | 1375 | 68 | 98.88 | - | 0.69 | 6.43E-04 | 2.69E-02 |
| negative regulation of RNA biosynthetic process (GO:1902679) | 1274 | 63 | 91.62 | - | 0.69 | 9.37E-04 | 3.59E-02 |
| negative regulation of nucleobase-containing compound metabolic process (GO:0045934) | 1501 | 74 | 107.94 | - | 0.69 | 2.73E-04 | 1.42E-02 |
| RNA biosynthetic process (GO:0032774) | 1367 | 67 | 98.31 | - | 0.68 | 5.08E-04 | 2.26E-02 |

| | | | | | | | |
|---|---|---|---|---|---|---|---|
| nucleic acid biosynthetic process (GO:0141187) | 1455 | 71 | 104.64 | - | 0.68 | 2.71E-04 | 1.41E-02 |
| RNA metabolic process (GO:0016070) | 1519 | 74 | 109.24 | - | 0.68 | 1.60E-04 | 9.59E-03 |
| nucleic acid metabolic process (GO:0090304) | 2140 | 99 | 153.9 | - | 0.64 | 3.47E-07 | 6.64E-05 |
| immune response (GO:0006955) | 1662 | 76 | 119.52 | - | 0.64 | 6.22E-06 | 7.90E-04 |
| cell cycle process (GO:0022402) | 881 | 40 | 63.36 | - | 0.63 | 1.33E-03 | 4.79E-02 |
| sexual reproduction (GO:0019953) | 1049 | 47 | 75.44 | - | 0.62 | 2.82E-04 | 1.45E-02 |
| RNA processing (GO:0006396) | 865 | 37 | 62.21 | - | 0.59 | 4.46E-04 | 2.05E-02 |
| protein-DNA complex organization (GO:0071824) | 878 | 37 | 63.14 | - | 0.59 | 2.32E-04 | 1.26E-02 |
| chromatin organization (GO:0006325) | 786 | 32 | 56.52 | - | 0.57 | 3.09E-04 | 1.55E-02 |
| regulation of cell cycle process (GO:0010564) | 717 | 29 | 51.56 | - | 0.56 | 5.12E-04 | 2.27E-02 |
| adaptive immune response (GO:0002250) | 677 | 27 | 48.69 | - | 0.55 | 6.23E-04 | 2.65E-02 |
| Unclassified (UNCLASSIFIED) | 2789 | 109 | 200.57 | - | 0.54 | 1.08E-14 | 1.49E-11 |
| DNA metabolic process (GO:0006259) | 735 | 27 | 52.86 | - | 0.51 | 5.86E-05 | 4.37E-03 |
| detection of stimulus (GO:0051606) | 675 | 24 | 48.54 | - | 0.49 | 7.38E-05 | 5.05E-03 |
| DNA repair (GO:0006281) | 516 | 15 | 37.11 | - | 0.4 | 3.00E-05 | 2.62E-03 |
| regulation of cell cycle phase transition (GO:1901987) | 428 | 11 | 30.78 | - | 0.36 | 4.00E-05 | 3.32E-03 |
| chromosome organization (GO:0051276) | 467 | 12 | 33.58 | - | 0.36 | 1.68E-05 | 1.69E-03 |
| detection of stimulus involved in sensory perception (GO:0050906) | 558 | 12 | 40.13 | - | 0.3 | 1.18E-07 | 2.66E-05 |
| regulation of chromosome organization (GO:0033044) | 246 | 5 | 17.69 | - | 0.28 | 4.54E-04 | 2.07E-02 |

| | | | | | | | |
|---|---|---|---|---|---|---|---|
| rRNA metabolic process (GO:0016072) | 250 | 5 | 17.98 | - | 0.28 | 4.66E-04 | 2.11E-02 |
| rRNA processing (GO:0006364) | 217 | 4 | 15.61 | - | 0.26 | 7.96E-04 | 3.19E-02 |
| double-strand break repair (GO:0006302) | 217 | 4 | 15.61 | - | 0.26 | 7.96E-04 | 3.18E-02 |
| regulation of mitotic cell cycle phase transition (GO:1901990) | 332 | 6 | 23.88 | - | 0.25 | 1.40E-05 | 1.45E-03 |
| regulation of DNA repair (GO:0006282) | 224 | 3 | 16.11 | - | 0.19 | 1.19E-04 | 7.65E-03 |
| regulation of G1/S transition of mitotic cell cycle (GO:2000045) | 160 | 2 | 11.51 | - | 0.17 | 1.04E-03 | 3.92E-02 |
| DNA recombination (GO:0006310) | 249 | 3 | 17.91 | - | 0.17 | 1.94E-05 | 1.92E-03 |
| sensory perception of chemical stimulus (GO:0007606) | 542 | 5 | 38.98 | - | 0.13 | 4.72E-12 | 2.38E-09 |
| immunoglobulin production (GO:0002377) | 141 | 1 | 10.14 | - | 0.1 | 7.75E-04 | 3.13E-02 |
| production of molecular mediator of immune response (GO:0002440) | 147 | 1 | 10.57 | - | 0.09 | 3.35E-04 | 1.66E-02 |
| sensory perception of smell (GO:0007608) | 466 | 3 | 33.51 | - | 0.09 | 9.56E-12 | 4.52E-09 |
| detection of chemical stimulus (GO:0009593) | 520 | 1 | 37.4 | - | 0.03 | 8.73E-16 | 2.64E-12 |
| regulation of chromosome segregation (GO:0051983) | 131 | 0 | 9.42 | - | < 0.01 | 1.11E-04 | 7.20E-03 |
| regulation of sister chromatid segregation (GO:0033045) | 105 | 0 | 7.55 | - | < 0.01 | 8.58E-04 | 3.34E-02 |
| recombinational repair (GO:0000725) | 125 | 0 | 8.99 | - | < 0.01 | 1.64E-04 | 9.71E-03 |
| double-strand break repair via homologous recombination (GO:0000724) | 120 | 0 | 8.63 | - | < 0.01 | 2.48E-04 | 1.31E-02 |
| detection of chemical stimulus involved in sensory perception of smell (GO:0050911) | 440 | 0 | 31.64 | - | < 0.01 | 6.91E-15 | 1.16E-11 |
| detection of chemical stimulus involved in sensory perception (GO:0050907) | 488 | 0 | 35.09 | - | < 0.01 | 1.73E-16 | 6.54E-13 |

**CEP290_Up:**

| | |
|---|---|
| Analysis Type: | PANTHER Overrepresentation Test (Released 20240807) |
| Annotation Version and Release Date: | GO Ontology database DOI: 10.5281/zenodo.12173881 Released 2024-06-17 |
| Analyzed List: | upload_1 (Homo sapiens) |
| Reference List: | Homo sapiens (all genes in database) |
| Test Type: | FISHER |
| Correction: | FDR |

| GO molecular function complete | Homo sapiens - REFLIST (20580) | upload_1 (1288) | upload_1 (expected) | upload_1 (over/under) | upload_1 (fold Enrichment) | upload_1 (raw P-value) | upload_1 (FDR) |
|---|---|---|---|---|---|---|---|
| stearoyl-CoA 9-desaturase activity (GO:0004768) | 3 | 3 | 0.19 | + | 15.98 | 2.45E-04 | 1.53E-02 |
| acyl-CoA desaturase activity (GO:0016215) | 4 | 3 | 0.25 | + | 11.98 | 9.33E-04 | 4.50E-02 |
| gap junction channel activity involved in cardiac conduction electrical coupling (GO:0086075) | 4 | 3 | 0.25 | + | 11.98 | 9.33E-04 | 4.46E-02 |
| structural molecule activity conferring elasticity (GO:0097493) | 12 | 8 | 0.75 | + | 10.65 | 9.09E-08 | 1.15E-05 |
| platelet-derived growth factor binding (GO:0048407) | 11 | 7 | 0.69 | + | 10.17 | 9.77E-07 | 1.01E-04 |
| extracellular matrix constituent conferring elasticity (GO:0030023) | 10 | 6 | 0.63 | + | 9.59 | 1.00E-05 | 8.77E-04 |
| mitogen-activated protein kinase p38 binding (GO:0048273) | 7 | 4 | 0.44 | + | 9.13 | 4.59E-04 | 2.70E-02 |
| platelet-derived growth factor receptor binding (GO:0005161) | 14 | 7 | 0.88 | + | 7.99 | 8.60E-06 | 7.65E-04 |
| glycine binding (GO:0016594) | 11 | 5 | 0.69 | + | 7.26 | 3.20E-04 | 1.96E-02 |
| gamma-catenin binding (GO:0045295) | 12 | 5 | 0.75 | + | 6.66 | 5.21E-04 | 2.97E-02 |
| HMG box domain binding (GO:0071837) | 13 | 5 | 0.81 | + | 6.15 | 8.02E-04 | 4.03E-02 |

| | | | | | | | |
|---|---|---|---|---|---|---|---|
| glutamate receptor activity (GO:0008066) | 26 | 9 | 1.63 | + | 5.53 | 1.69E-05 | 1.43E-03 |
| ephrin receptor activity (GO:0005003) | 18 | 6 | 1.13 | + | 5.33 | 5.74E-04 | 3.06E-02 |
| extracellular matrix structural constituent conferring tensile strength (GO:0030020) | 45 | 15 | 2.82 | + | 5.33 | 4.73E-08 | 6.48E-06 |
| transmembrane receptor protein tyrosine kinase activity (GO:0004714) | 60 | 20 | 3.76 | + | 5.33 | 2.77E-10 | 5.61E-08 |
| RNA polymerase II core promoter sequence-specific DNA binding (GO:0000979) | 19 | 6 | 1.19 | + | 5.05 | 7.94E-04 | 4.03E-02 |
| collagen binding (GO:0005518) | 68 | 21 | 4.26 | + | 4.93 | 4.99E-10 | 8.72E-08 |
| SMAD binding (GO:0046332) | 78 | 24 | 4.88 | + | 4.92 | 3.17E-11 | 8.04E-09 |
| transmembrane receptor protein kinase activity (GO:0019199) | 79 | 24 | 4.94 | + | 4.85 | 4.28E-11 | 1.03E-08 |
| retinoic acid binding (GO:0001972) | 20 | 6 | 1.25 | + | 4.79 | 1.07E-03 | 4.96E-02 |
| transcription regulator inhibitor activity (GO:0140416) | 30 | 9 | 1.88 | + | 4.79 | 6.17E-05 | 4.60E-03 |
| extracellular matrix structural constituent (GO:0005201) | 166 | 49 | 10.39 | + | 4.72 | 1.32E-20 | 6.69E-17 |
| mitogen-activated protein kinase binding (GO:0051019) | 28 | 8 | 1.75 | + | 4.57 | 2.32E-04 | 1.49E-02 |
| miRNA binding (GO:0035198) | 32 | 9 | 2 | + | 4.49 | 1.08E-04 | 7.61E-03 |
| Wnt-protein binding (GO:0017147) | 32 | 9 | 2 | + | 4.49 | 1.08E-04 | 7.50E-03 |
| poly-purine tract binding (GO:0070717) | 31 | 8 | 1.94 | + | 4.12 | 4.97E-04 | 2.86E-02 |
| pre-mRNA binding (GO:0036002) | 39 | 10 | 2.44 | + | 4.1 | 1.06E-04 | 7.60E-03 |

| | | | | | | | |
|---|---|---|---|---|---|---|---|
| extracellular matrix binding (GO:0050840) | 64 | 15 | 4.01 | + | 3.74 | 7.10E-06 | 6.67E-04 |
| peptide hormone binding (GO:0017046) | 50 | 11 | 3.13 | + | 3.52 | 2.15E-04 | 1.40E-02 |
| core promoter sequence-specific DNA binding (GO:0001046) | 41 | 9 | 2.57 | + | 3.51 | 8.11E-04 | 4.03E-02 |
| protein tyrosine kinase activity (GO:0004713) | 138 | 30 | 8.64 | + | 3.47 | 1.55E-09 | 2.62E-07 |
| transmitter-gated channel activity (GO:0022835) | 63 | 13 | 3.94 | + | 3.3 | 1.18E-04 | 8.09E-03 |
| transmitter-gated monoatomic ion channel activity (GO:0022824) | 63 | 13 | 3.94 | + | 3.3 | 1.18E-04 | 7.98E-03 |
| growth factor binding (GO:0019838) | 135 | 27 | 8.45 | + | 3.2 | 6.47E-08 | 8.41E-06 |
| extracellular ligand-gated monoatomic ion channel activity (GO:0005230) | 73 | 13 | 4.57 | + | 2.85 | 5.44E-04 | 3.00E-02 |
| integrin binding (GO:0005178) | 158 | 28 | 9.89 | + | 2.83 | 5.24E-07 | 5.65E-05 |
| single-stranded RNA binding (GO:0003727) | 85 | 15 | 5.32 | + | 2.82 | 2.33E-04 | 1.47E-02 |
| heparin binding (GO:0008201) | 173 | 30 | 10.83 | + | 2.77 | 3.39E-07 | 3.90E-05 |
| beta-catenin binding (GO:0008013) | 106 | 18 | 6.63 | + | 2.71 | 9.66E-05 | 6.99E-03 |
| hormone binding (GO:0042562) | 92 | 15 | 5.76 | + | 2.61 | 5.62E-04 | 3.03E-02 |
| glycosaminoglycan binding (GO:0005539) | 241 | 38 | 15.08 | + | 2.52 | 1.31E-07 | 1.61E-05 |
| kinase activator activity (GO:0019209) | 142 | 22 | 8.89 | + | 2.48 | 7.23E-05 | 5.31E-03 |
| neurotransmitter receptor activity (GO:0030594) | 97 | 15 | 6.07 | + | 2.47 | 9.91E-04 | 4.61E-02 |

| | | | | | | | |
|---|---|---|---|---|---|---|---|
| transcription coregulator binding (GO:0001221) | 114 | 17 | 7.13 | + | 2.38 | 7.21E-04 | 3.69E-02 |
| transcription corepressor activity (GO:0003714) | 197 | 29 | 12.33 | + | 2.35 | 1.80E-05 | 1.47E-03 |
| organic acid binding (GO:0043177) | 195 | 27 | 12.2 | + | 2.21 | 1.32E-04 | 8.80E-03 |
| DNA-binding transcription activator activity (GO:0001216) | 482 | 66 | 30.17 | + | 2.19 | 1.72E-09 | 2.82E-07 |
| DNA-binding transcription activator activity, RNA polymerase II-specific (GO:0001228) | 476 | 65 | 29.79 | + | 2.18 | 3.94E-09 | 6.06E-07 |
| sulfur compound binding (GO:1901681) | 268 | 36 | 16.77 | + | 2.15 | 1.74E-05 | 1.45E-03 |
| DNA-binding transcription repressor activity (GO:0001217) | 273 | 36 | 17.09 | + | 2.11 | 2.25E-05 | 1.81E-03 |
| transcription factor binding (GO:0008134) | 600 | 79 | 37.55 | + | 2.1 | 3.19E-10 | 6.00E-08 |
| carboxylic acid binding (GO:0031406) | 184 | 24 | 11.52 | + | 2.08 | 6.01E-04 | 3.14E-02 |
| DNA-binding transcription factor binding (GO:0140297) | 478 | 62 | 29.92 | + | 2.07 | 4.85E-08 | 6.48E-06 |
| kinase regulator activity (GO:0019207) | 279 | 36 | 17.46 | + | 2.06 | 4.72E-05 | 3.68E-03 |
| DNA-binding transcription repressor activity, RNA polymerase II-specific (GO:0001227) | 264 | 34 | 16.52 | + | 2.06 | 5.77E-05 | 4.37E-03 |
| RNA polymerase II-specific DNA-binding transcription factor binding (GO:0061629) | 343 | 44 | 21.47 | + | 2.05 | 7.64E-06 | 7.05E-04 |
| mRNA binding (GO:0003729) | 343 | 44 | 21.47 | + | 2.05 | 7.64E-06 | 6.92E-04 |
| transcription coregulator activity (GO:0003712) | 504 | 63 | 31.54 | + | 2 | 1.84E-07 | 2.22E-05 |
| chromatin binding (GO:0003682) | 624 | 78 | 39.05 | + | 2 | 6.30E-09 | 9.40E-07 |

| | | | | | | | |
|---|---|---|---|---|---|---|---|
| guanyl-nucleotide exchange factor activity (GO:0005085) | 233 | 29 | 14.58 | + | 1.99 | 5.26E-04 | 2.93E-02 |
| protein kinase activity (GO:0004672) | 564 | 70 | 35.3 | + | 1.98 | 4.46E-08 | 6.28E-06 |
| calcium ion binding (GO:0005509) | 711 | 87 | 44.5 | + | 1.96 | 1.88E-09 | 2.98E-07 |
| transcription coactivator activity (GO:0003713) | 273 | 33 | 17.09 | + | 1.93 | 3.38E-04 | 2.04E-02 |
| protein kinase regulator activity (GO:0019887) | 241 | 29 | 15.08 | + | 1.92 | 6.95E-04 | 3.59E-02 |
| cell adhesion molecule binding (GO:0050839) | 552 | 66 | 34.55 | + | 1.91 | 4.18E-07 | 4.60E-05 |
| small GTPase binding (GO:0031267) | 282 | 33 | 17.65 | + | 1.87 | 4.75E-04 | 2.77E-02 |
| cis-regulatory region sequence-specific DNA binding (GO:0000987) | 1219 | 141 | 76.29 | + | 1.85 | 7.68E-13 | 3.54E-10 |
| GTPase binding (GO:0051020) | 314 | 36 | 19.65 | + | 1.83 | 5.55E-04 | 3.02E-02 |
| RNA polymerase II cis-regulatory region sequence-specific DNA binding (GO:0000978) | 1195 | 137 | 74.79 | + | 1.83 | 3.01E-12 | 1.27E-09 |
| phosphotransferase activity, alcohol group as acceptor (GO:0016773) | 673 | 77 | 42.12 | + | 1.83 | 2.57E-07 | 3.03E-05 |
| sequence-specific double-stranded DNA binding (GO:1990837) | 1573 | 178 | 98.45 | + | 1.81 | 3.00E-15 | 5.07E-12 |
| RNA polymerase II transcription regulatory region sequence-specific DNA binding (GO:0000977) | 1411 | 159 | 88.31 | + | 1.8 | 1.69E-13 | 8.55E-11 |
| transcription regulatory region nucleic acid binding (GO:0001067) | 1512 | 170 | 94.63 | + | 1.8 | 3.11E-14 | 1.97E-11 |
| transcription cis-regulatory region binding (GO:0000976) | 1510 | 169 | 94.5 | + | 1.79 | 4.85E-14 | 2.73E-11 |
| transcription regulator activity (GO:0140110) | 1972 | 219 | 123.42 | + | 1.77 | 7.79E-18 | 1.98E-14 |

| | | | | | | | |
|---|---|---|---|---|---|---|---|
| protein kinase binding (GO:0019901) | 717 | 79 | 44.87 | + | 1.76 | 8.91E-07 | 9.41E-05 |
| sequence-specific DNA binding (GO:0043565) | 1684 | 185 | 105.39 | + | 1.76 | 1.44E-14 | 1.21E-11 |
| double-stranded DNA binding (GO:0003690) | 1668 | 183 | 104.39 | + | 1.75 | 2.48E-14 | 1.80E-11 |
| DNA-binding transcription factor activity, RNA polymerase II-specific (GO:0000981) | 1382 | 151 | 86.49 | + | 1.75 | 1.05E-11 | 3.33E-09 |
| kinase activity (GO:0016301) | 727 | 79 | 45.5 | + | 1.74 | 1.58E-06 | 1.60E-04 |
| actin binding (GO:0003779) | 439 | 47 | 27.47 | + | 1.71 | 3.04E-04 | 1.88E-02 |
| DNA-binding transcription factor activity (GO:0003700) | 1473 | 157 | 92.19 | + | 1.7 | 2.92E-11 | 7.78E-09 |
| kinase binding (GO:0019900) | 796 | 84 | 49.82 | + | 1.69 | 2.21E-06 | 2.20E-04 |
| protein-containing complex binding (GO:0044877) | 1764 | 179 | 110.4 | + | 1.62 | 4.82E-11 | 1.11E-08 |
| GTPase regulator activity (GO:0030695) | 495 | 50 | 30.98 | + | 1.61 | 9.21E-04 | 4.53E-02 |
| nucleoside-triphosphatase regulator activity (GO:0060589) | 495 | 50 | 30.98 | + | 1.61 | 9.21E-04 | 4.49E-02 |
| DNA binding (GO:0003677) | 2518 | 251 | 157.59 | + | 1.59 | 1.39E-14 | 1.41E-11 |
| transferase activity, transferring phosphorus-containing groups (GO:0016772) | 900 | 88 | 56.33 | + | 1.56 | 2.97E-05 | 2.35E-03 |
| protein-macromolecule adaptor activity (GO:0030674) | 1017 | 99 | 63.65 | + | 1.56 | 1.05E-05 | 9.06E-04 |
| molecular adaptor activity (GO:0060090) | 1161 | 111 | 72.66 | + | 1.53 | 6.48E-06 | 6.20E-04 |
| protein domain specific binding (GO:0019904) | 652 | 62 | 40.81 | + | 1.52 | 9.69E-04 | 4.55E-02 |

| | | | | | | | |
|---|---|---|---|---|---|---|---|
| structural molecule activity (GO:0005198) | 786 | 74 | 49.19 | + | 1.5 | 4.02E-04 | 2.40E-02 |
| signaling receptor binding (GO:0005102) | 1517 | 140 | 94.94 | + | 1.47 | 2.60E-06 | 2.53E-04 |
| enzyme binding (GO:0019899) | 2095 | 187 | 131.12 | + | 1.43 | 4.12E-07 | 4.64E-05 |
| cytoskeletal protein binding (GO:0008092) | 994 | 88 | 62.21 | + | 1.41 | 9.67E-04 | 4.58E-02 |
| enzyme regulator activity (GO:0030234) | 1325 | 117 | 82.93 | + | 1.41 | 1.33E-04 | 8.75E-03 |
| nucleic acid binding (GO:0003676) | 4010 | 344 | 250.97 | + | 1.37 | 8.52E-11 | 1.88E-08 |
| cation binding (GO:0043169) | 4431 | 379 | 277.31 | + | 1.37 | 7.21E-12 | 2.44E-09 |
| metal ion binding (GO:0046872) | 4331 | 369 | 271.06 | + | 1.36 | 2.71E-11 | 7.64E-09 |
| molecular function regulator activity (GO:0098772) | 2112 | 170 | 132.18 | + | 1.29 | 5.25E-04 | 2.96E-02 |
| ion binding (GO:0043167) | 6101 | 491 | 381.83 | + | 1.29 | 1.70E-11 | 5.06E-09 |
| small molecule binding (GO:0036094) | 6296 | 497 | 394.04 | + | 1.26 | 3.12E-10 | 6.08E-08 |
| organic cyclic compound binding (GO:0097159) | 6040 | 469 | 378.01 | + | 1.24 | 1.74E-08 | 2.52E-06 |
| protein binding (GO:0005515) | 14060 | 980 | 879.95 | + | 1.11 | 2.56E-10 | 5.41E-08 |
| binding (GO:0005488) | 16418 | 1130 | 1027.52 | + | 1.1 | 1.30E-14 | 1.64E-11 |
| molecular_function (GO:0003674) | 18198 | 1211 | 1138.92 | + | 1.06 | 3.13E-12 | 1.22E-09 |
| Unclassified (UNCLASSIFIED) | 2382 | 77 | 149.08 | - | 0.52 | 3.13E-12 | 1.13E-09 |

| | | | | | | | |
|---|---|---|---|---|---|---|---|
| antigen binding (GO:0003823) | 200 | 1 | 12.52 | - | 0.08 | 5.34E-05 | 4.10E-03 |
| olfactory receptor activity (GO:0004984) | 438 | 2 | 27.41 | - | 0.07 | 3.70E-10 | 6.71E-08 |
| odorant binding (GO:0005549) | 125 | 0 | 7.82 | - | < 0.01 | 5.84E-04 | 3.08E-02 |

**NPHP1_Down:**

| | |
|---|---|
| Analysis Type: | PANTHER Overrepresentation Test (Released 20240807) |
| Annotation Version and Release Date: | GO Ontology database DOI: 10.5281/zenodo.12173881 Released 2024-06-17 |
| Analyzed List: | upload_1 (Homo sapiens) |
| Reference List: | Homo sapiens (all genes in database) |
| Test Type: | FISHER |
| Correction: | FDR |

| GO biological process complete | Homo sapiens - REFLIST (20580) | upload_1 (2477) | upload_1 (expected) | upload_1 (over/under) | upload_1 (fold Enrichment) | upload_1 (raw P-value) | upload_1 (FDR) |
|---|---|---|---|---|---|---|---|
| positive regulation of potassium ion import across plasma membrane (GO:1903288) | 3 | 3 | 0.36 | + | 8.31 | 1.74E-03 | 5.09E-02 |
| regulation of potassium ion import (GO:1903286) | 4 | 4 | 0.48 | + | 8.31 | 2.09E-04 | 9.48E-03 |
| calcium ion regulated lysosome exocytosis (GO:1990927) | 3 | 3 | 0.36 | + | 8.31 | 1.74E-03 | 5.08E-02 |
| regulation of endocannabinoid signaling pathway (GO:2000124) | 3 | 3 | 0.36 | + | 8.31 | 1.74E-03 | 5.07E-02 |
| negative regulation of synaptic plasticity (GO:0031914) | 3 | 3 | 0.36 | + | 8.31 | 1.74E-03 | 5.06E-02 |
| protein maturation by [2Fe-2S] cluster transfer (GO:0106034) | 3 | 3 | 0.36 | + | 8.31 | 1.74E-03 | 5.05E-02 |
| regulation of activated CD8-positive, alpha-beta T cell apoptotic process (GO:1905402) | 3 | 3 | 0.36 | + | 8.31 | 1.74E-03 | 5.04E-02 |
| protein targeting to lysosome involved in chaperone-mediated autophagy (GO:0061740) | 3 | 3 | 0.36 | + | 8.31 | 1.74E-03 | 5.04E-02 |
| positive regulation of mitochondrial electron transport, NADH to ubiquinone (GO:1902958) | 3 | 3 | 0.36 | + | 8.31 | 1.74E-03 | 5.03E-02 |
| cellular response to ammonium ion (GO:0071242) | 5 | 4 | 0.6 | + | 6.65 | 9.46E-04 | 3.14E-02 |
| negative regulation of IRE1-mediated unfolded protein response (GO:1903895) | 5 | 4 | 0.6 | + | 6.65 | 9.46E-04 | 3.14E-02 |

| | | | | | | | |
|---|---|---|---|---|---|---|---|
| short-chain fatty acid catabolic process (GO:0019626) | 7 | 5 | 0.84 | + | 5.93 | 4.28E-04 | 1.67E-02 |
| membrane hyperpolarization (GO:0060081) | 9 | 6 | 1.08 | + | 5.54 | 1.84E-04 | 8.46E-03 |
| negative regulation of heart rate (GO:0010459) | 11 | 7 | 1.32 | + | 5.29 | 7.70E-05 | 4.17E-03 |
| tetrahydrobiopterin metabolic process (GO:0046146) | 8 | 5 | 0.96 | + | 5.19 | 1.03E-03 | 3.36E-02 |
| protein maturation by [4Fe-4S] cluster transfer (GO:0106035) | 8 | 5 | 0.96 | + | 5.19 | 1.03E-03 | 3.35E-02 |
| PERK-mediated unfolded protein response (GO:0036499) | 8 | 5 | 0.96 | + | 5.19 | 1.03E-03 | 3.34E-02 |
| heparan sulfate proteoglycan biosynthetic process, enzymatic modification (GO:0015015) | 12 | 7 | 1.44 | + | 4.85 | 1.65E-04 | 7.80E-03 |
| heparan sulfate proteoglycan metabolic process (GO:0030201) | 16 | 9 | 1.93 | + | 4.67 | 2.70E-05 | 1.77E-03 |
| protein insertion into ER membrane by stop-transfer membrane-anchor sequence (GO:0045050) | 11 | 6 | 1.32 | + | 4.53 | 8.14E-04 | 2.78E-02 |
| protein folding in endoplasmic reticulum (GO:0034975) | 11 | 6 | 1.32 | + | 4.53 | 8.14E-04 | 2.78E-02 |
| synaptic transmission, dopaminergic (GO:0001963) | 17 | 9 | 2.05 | + | 4.4 | 5.12E-05 | 3.01E-03 |
| lactate metabolic process (GO:0006089) | 17 | 9 | 2.05 | + | 4.4 | 5.12E-05 | 3.00E-03 |
| magnesium ion transport (GO:0015693) | 19 | 10 | 2.29 | + | 4.37 | 2.06E-05 | 1.42E-03 |
| cellular response to purine-containing compound (GO:0071415) | 14 | 7 | 1.69 | + | 4.15 | 5.76E-04 | 2.12E-02 |
| enteric nervous system development (GO:0048484) | 12 | 6 | 1.44 | + | 4.15 | 1.46E-03 | 4.43E-02 |
| magnesium ion transmembrane transport (GO:1903830) | 18 | 9 | 2.17 | + | 4.15 | 9.15E-05 | 4.80E-03 |

| | | | | | | | |
|---|---|---|---|---|---|---|---|
| protein maturation by iron-sulfur cluster transfer (GO:0097428) | 16 | 8 | 1.93 | + | 4.15 | 2.29E-04 | 1.02E-02 |
| positive regulation of fibroblast migration (GO:0010763) | 14 | 7 | 1.69 | + | 4.15 | 5.76E-04 | 2.11E-02 |
| mitochondrial electron transport, cytochrome c to oxygen (GO:0006123) | 23 | 11 | 2.77 | + | 3.97 | 2.54E-05 | 1.70E-03 |
| mitochondrial electron transport, NADH to ubiquinone (GO:0006120) | 46 | 22 | 5.54 | + | 3.97 | 2.35E-09 | 3.52E-07 |
| membrane repolarization during action potential (GO:0086011) | 21 | 10 | 2.53 | + | 3.96 | 6.26E-05 | 3.52E-03 |
| long-term synaptic depression (GO:0060292) | 19 | 9 | 2.29 | + | 3.94 | 1.55E-04 | 7.36E-03 |
| neurotransmitter receptor localization to postsynaptic specialization membrane (GO:0099645) | 15 | 7 | 1.81 | + | 3.88 | 9.68E-04 | 3.19E-02 |
| protein localization to postsynaptic specialization membrane (GO:0099633) | 15 | 7 | 1.81 | + | 3.88 | 9.68E-04 | 3.19E-02 |
| positive regulation of sequestering of calcium ion (GO:0051284) | 15 | 7 | 1.81 | + | 3.88 | 9.68E-04 | 3.18E-02 |
| membrane repolarization (GO:0086009) | 26 | 12 | 3.13 | + | 3.83 | 1.70E-05 | 1.20E-03 |
| aerobic electron transport chain (GO:0019646) | 87 | 40 | 10.47 | + | 3.82 | 3.88E-15 | 1.68E-12 |
| mitochondrial ATP synthesis coupled electron transport (GO:0042775) | 92 | 42 | 11.07 | + | 3.79 | 1.10E-15 | 5.72E-13 |
| ATP synthesis coupled electron transport (GO:0042773) | 92 | 42 | 11.07 | + | 3.79 | 1.10E-15 | 5.53E-13 |
| cardiac muscle cell membrane repolarization (GO:0099622) | 20 | 9 | 2.41 | + | 3.74 | 2.52E-04 | 1.10E-02 |
| membrane repolarization during cardiac muscle cell action potential (GO:0086013) | 20 | 9 | 2.41 | + | 3.74 | 2.52E-04 | 1.10E-02 |
| regulation of protein kinase C signaling (GO:0090036) | 18 | 8 | 2.17 | + | 3.69 | 6.23E-04 | 2.26E-02 |

| | | | | | | | |
|---|---|---|---|---|---|---|---|
| regulation of catecholamine metabolic process (GO:0042069) | 18 | 8 | 2.17 | + | 3.69 | 6.23E-04 | 2.26E-02 |
| regulation of dopamine metabolic process (GO:0042053) | 18 | 8 | 2.17 | + | 3.69 | 6.23E-04 | 2.25E-02 |
| ATP biosynthetic process (GO:0006754) | 84 | 36 | 10.11 | + | 3.56 | 1.25E-12 | 3.27E-10 |
| negative regulation of phagocytosis (GO:0050765) | 21 | 9 | 2.53 | + | 3.56 | 3.94E-04 | 1.57E-02 |
| negative regulation of heart contraction (GO:0045822) | 21 | 9 | 2.53 | + | 3.56 | 3.94E-04 | 1.57E-02 |
| respiratory electron transport chain (GO:0022904) | 117 | 50 | 14.08 | + | 3.55 | 6.37E-17 | 4.19E-14 |
| proton motive force-driven ATP synthesis (GO:0015986) | 73 | 31 | 8.79 | + | 3.53 | 6.03E-11 | 1.22E-08 |
| proton motive force-driven mitochondrial ATP synthesis (GO:0042776) | 64 | 27 | 7.7 | + | 3.51 | 1.23E-09 | 1.90E-07 |
| positive regulation of autophagy of mitochondrion (GO:1903599) | 19 | 8 | 2.29 | + | 3.5 | 9.62E-04 | 3.18E-02 |
| electron transport chain (GO:0022900) | 127 | 53 | 15.29 | + | 3.47 | 2.64E-17 | 2.22E-14 |
| response to amphetamine (GO:0001975) | 36 | 15 | 4.33 | + | 3.46 | 7.19E-06 | 5.60E-04 |
| monoamine transport (GO:0015844) | 29 | 12 | 3.49 | + | 3.44 | 6.47E-05 | 3.61E-03 |
| NADH dehydrogenase complex assembly (GO:0010257) | 61 | 25 | 7.34 | + | 3.41 | 1.03E-08 | 1.41E-06 |
| mitochondrial respiratory chain complex I assembly (GO:0032981) | 61 | 25 | 7.34 | + | 3.41 | 1.03E-08 | 1.40E-06 |
| relaxation of muscle (GO:0090075) | 22 | 9 | 2.65 | + | 3.4 | 5.96E-04 | 2.18E-02 |
| negative regulation of blood circulation (GO:1903523) | 22 | 9 | 2.65 | + | 3.4 | 5.96E-04 | 2.18E-02 |

| | | | | | | | |
|---|---|---|---|---|---|---|---|
| NADH metabolic process (GO:0006734) | 32 | 13 | 3.85 | + | 3.38 | 4.05E-05 | 2.50E-03 |
| chaperone cofactor-dependent protein refolding (GO:0051085) | 32 | 13 | 3.85 | + | 3.38 | 4.05E-05 | 2.49E-03 |
| 'de novo' protein folding (GO:0006458) | 42 | 17 | 5.06 | + | 3.36 | 2.87E-06 | 2.45E-04 |
| oxidative phosphorylation (GO:0006119) | 119 | 48 | 14.32 | + | 3.35 | 4.12E-15 | 1.73E-12 |
| adenylate cyclase-activating adrenergic receptor signaling pathway (GO:0071880) | 20 | 8 | 2.41 | + | 3.32 | 1.44E-03 | 4.43E-02 |
| response to hydroperoxide (GO:0033194) | 20 | 8 | 2.41 | + | 3.32 | 1.44E-03 | 4.42E-02 |
| negative regulation of mitochondrial membrane permeability (GO:0035795) | 20 | 8 | 2.41 | + | 3.32 | 1.44E-03 | 4.41E-02 |
| dopamine metabolic process (GO:0042417) | 31 | 12 | 3.73 | + | 3.22 | 1.40E-04 | 6.76E-03 |
| chaperone-mediated protein folding (GO:0061077) | 75 | 29 | 9.03 | + | 3.21 | 3.48E-09 | 5.06E-07 |
| regulation of presynaptic membrane potential (GO:0099505) | 29 | 11 | 3.49 | + | 3.15 | 3.26E-04 | 1.35E-02 |
| purine ribonucleoside triphosphate biosynthetic process (GO:0009206) | 95 | 36 | 11.43 | + | 3.15 | 9.11E-11 | 1.70E-08 |
| 'de novo' post-translational protein folding (GO:0051084) | 37 | 14 | 4.45 | + | 3.14 | 5.28E-05 | 3.07E-03 |
| purine nucleoside triphosphate biosynthetic process (GO:0009145) | 96 | 36 | 11.55 | + | 3.12 | 1.29E-10 | 2.38E-08 |
| protein localization to postsynaptic membrane (GO:1903539) | 32 | 12 | 3.85 | + | 3.12 | 1.99E-04 | 9.09E-03 |
| negative regulation of amine transport (GO:0051953) | 27 | 10 | 3.25 | + | 3.08 | 7.58E-04 | 2.63E-02 |
| gluconeogenesis (GO:0006094) | 49 | 18 | 5.9 | + | 3.05 | 7.55E-06 | 5.85E-04 |

| | | | | | | | |
|---|---|---|---|---|---|---|---|
| neuronal action potential (GO:0019228) | 30 | 11 | 3.61 | + | 3.05 | 4.59E-04 | 1.77E-02 |
| response to amine (GO:0014075) | 52 | 19 | 6.26 | + | 3.04 | 4.65E-06 | 3.86E-04 |
| protein localization to postsynapse (GO:0062237) | 33 | 12 | 3.97 | + | 3.02 | 2.79E-04 | 1.19E-02 |
| adrenergic receptor signaling pathway (GO:0071875) | 25 | 9 | 3.01 | + | 2.99 | 1.75E-03 | 5.03E-02 |
| plasma membrane repair (GO:0001778) | 25 | 9 | 3.01 | + | 2.99 | 1.75E-03 | 5.02E-02 |
| maintenance of synapse structure (GO:0099558) | 25 | 9 | 3.01 | + | 2.99 | 1.75E-03 | 5.01E-02 |
| synaptic vesicle maturation (GO:0016188) | 25 | 9 | 3.01 | + | 2.99 | 1.75E-03 | 5.00E-02 |
| retina layer formation (GO:0010842) | 25 | 9 | 3.01 | + | 2.99 | 1.75E-03 | 4.99E-02 |
| protein refolding (GO:0042026) | 25 | 9 | 3.01 | + | 2.99 | 1.75E-03 | 4.98E-02 |
| cerebellar Purkinje cell layer development (GO:0021680) | 28 | 10 | 3.37 | + | 2.97 | 1.05E-03 | 3.41E-02 |
| regulation of catecholamine secretion (GO:0050433) | 42 | 15 | 5.06 | + | 2.97 | 6.30E-05 | 3.53E-03 |
| ribonucleoside triphosphate biosynthetic process (GO:0009201) | 101 | 36 | 12.16 | + | 2.96 | 6.71E-10 | 1.07E-07 |
| aerobic respiration (GO:0009060) | 163 | 58 | 19.62 | + | 2.96 | 4.90E-15 | 2.00E-12 |
| positive regulation of blood pressure (GO:0045777) | 31 | 11 | 3.73 | + | 2.95 | 6.35E-04 | 2.29E-02 |
| cellular respiration (GO:0045333) | 194 | 68 | 23.35 | + | 2.91 | 5.42E-17 | 3.72E-14 |
| hexose biosynthetic process (GO:0019319) | 52 | 18 | 6.26 | + | 2.88 | 1.95E-05 | 1.35E-03 |

| | | | | | | | |
|---|---|---|---|---|---|---|---|
| fatty-acyl-CoA biosynthetic process (GO:0046949) | 29 | 10 | 3.49 | + | 2.86 | 1.43E-03 | 4.44E-02 |
| pyruvate metabolic process (GO:0006090) | 70 | 24 | 8.43 | + | 2.85 | 1.03E-06 | 9.53E-05 |
| acyl-CoA biosynthetic process (GO:0071616) | 44 | 15 | 5.3 | + | 2.83 | 1.16E-04 | 5.91E-03 |
| thioester biosynthetic process (GO:0035384) | 44 | 15 | 5.3 | + | 2.83 | 1.16E-04 | 5.89E-03 |
| cerebellar cortex development (GO:0021695) | 53 | 18 | 6.38 | + | 2.82 | 2.63E-05 | 1.74E-03 |
| monosaccharide biosynthetic process (GO:0046364) | 56 | 19 | 6.74 | + | 2.82 | 1.61E-05 | 1.15E-03 |
| protein localization to synapse (GO:0035418) | 60 | 20 | 7.22 | + | 2.77 | 1.31E-05 | 9.50E-04 |
| insulin secretion (GO:0030073) | 48 | 16 | 5.78 | + | 2.77 | 9.42E-05 | 4.93E-03 |
| nucleoside triphosphate biosynthetic process (GO:0009142) | 109 | 36 | 13.12 | + | 2.74 | 7.13E-09 | 9.99E-07 |
| mitochondrial respiratory chain complex assembly (GO:0033108) | 103 | 34 | 12.4 | + | 2.74 | 1.88E-08 | 2.43E-06 |
| regulation of heart rate (GO:0002027) | 100 | 33 | 12.04 | + | 2.74 | 3.05E-08 | 3.82E-06 |
| regulation of heart rate by cardiac conduction (GO:0086091) | 40 | 13 | 4.81 | + | 2.7 | 5.56E-04 | 2.08E-02 |
| tricarboxylic acid cycle (GO:0006099) | 34 | 11 | 4.09 | + | 2.69 | 1.52E-03 | 4.59E-02 |
| purine ribonucleotide biosynthetic process (GO:0009152) | 195 | 63 | 23.47 | + | 2.68 | 5.93E-14 | 2.19E-11 |
| energy derivation by oxidation of organic compounds (GO:0015980) | 273 | 88 | 32.86 | + | 2.68 | 7.25E-19 | 8.44E-16 |
| positive regulation of insulin secretion (GO:0032024) | 87 | 28 | 10.47 | + | 2.67 | 5.94E-07 | 5.99E-05 |

| | | | | | | | | |
|---|---|---|---|---|---|---|---|---|
| ATP metabolic process (GO:0046034) | 162 | 52 | 19.5 | + | | 2.67 | 1.20E-11 | 2.84E-09 |
| inner mitochondrial membrane organization (GO:0007007) | 41 | 13 | 4.93 | + | | 2.63 | 7.25E-04 | 2.54E-02 |
| negative regulation of synaptic transmission (GO:0050805) | 41 | 13 | 4.93 | + | | 2.63 | 7.25E-04 | 2.54E-02 |
| cerebellar cortex morphogenesis (GO:0021696) | 38 | 12 | 4.57 | + | | 2.62 | 1.20E-03 | 3.82E-02 |
| purine nucleoside bisphosphate biosynthetic process (GO:0034033) | 54 | 17 | 6.5 | + | | 2.62 | 1.29E-04 | 6.36E-03 |
| ribonucleoside bisphosphate biosynthetic process (GO:0034030) | 54 | 17 | 6.5 | + | | 2.62 | 1.29E-04 | 6.33E-03 |
| nucleoside bisphosphate biosynthetic process (GO:0033866) | 54 | 17 | 6.5 | + | | 2.62 | 1.29E-04 | 6.31E-03 |
| ribonucleotide biosynthetic process (GO:0009260) | 210 | 66 | 25.28 | + | | 2.61 | 6.61E-14 | 2.32E-11 |
| hindbrain morphogenesis (GO:0021575) | 48 | 15 | 5.78 | + | | 2.6 | 3.46E-04 | 1.40E-02 |
| calcium-ion regulated exocytosis (GO:0017156) | 48 | 15 | 5.78 | + | | 2.6 | 3.46E-04 | 1.40E-02 |
| cerebellum morphogenesis (GO:0021587) | 45 | 14 | 5.42 | + | | 2.58 | 5.68E-04 | 2.10E-02 |
| response to electrical stimulus (GO:0051602) | 45 | 14 | 5.42 | + | | 2.58 | 5.68E-04 | 2.10E-02 |
| peptide hormone secretion (GO:0030072) | 71 | 22 | 8.55 | + | | 2.57 | 1.84E-05 | 1.29E-03 |
| neuron maturation (GO:0042551) | 42 | 13 | 5.06 | + | | 2.57 | 9.35E-04 | 3.12E-02 |
| ribose phosphate biosynthetic process (GO:0046390) | 217 | 67 | 26.12 | + | | 2.57 | 1.11E-13 | 3.63E-11 |
| purine-containing compound biosynthetic process (GO:0072522) | 240 | 74 | 28.89 | + | | 2.56 | 6.26E-15 | 2.49E-12 |

| | | | | | | | |
|---|---|---|---|---|---|---|---|
| protein transmembrane import into intracellular organelle (GO:0044743) | 39 | 12 | 4.69 | + | 2.56 | 1.54E-03 | 4.64E-02 |
| NAD metabolic process (GO:0019674) | 62 | 19 | 7.46 | + | 2.55 | 7.94E-05 | 4.27E-03 |
| regulation of sodium ion transmembrane transporter activity (GO:2000649) | 46 | 14 | 5.54 | + | 2.53 | 7.27E-04 | 2.54E-02 |
| regulation of sodium ion transmembrane transport (GO:1902305) | 56 | 17 | 6.74 | + | 2.52 | 2.11E-04 | 9.48E-03 |
| positive regulation of peptide hormone secretion (GO:0090277) | 112 | 34 | 13.48 | + | 2.52 | 1.89E-07 | 2.07E-05 |
| purine nucleotide biosynthetic process (GO:0006164) | 231 | 70 | 27.8 | + | 2.52 | 8.76E-14 | 3.01E-11 |
| positive regulation of receptor-mediated endocytosis (GO:0048260) | 53 | 16 | 6.38 | + | 2.51 | 3.45E-04 | 1.40E-02 |
| synaptic vesicle recycling (GO:0036465) | 63 | 19 | 7.58 | + | 2.51 | 1.01E-04 | 5.25E-03 |
| generation of precursor metabolites and energy (GO:0006091) | 373 | 112 | 44.89 | + | 2.49 | 7.80E-21 | 1.68E-17 |
| transmission of nerve impulse (GO:0019226) | 60 | 18 | 7.22 | + | 2.49 | 1.64E-04 | 7.76E-03 |
| long-term synaptic potentiation (GO:0060291) | 50 | 15 | 6.02 | + | 2.49 | 5.64E-04 | 2.09E-02 |
| positive regulation of peptide secretion (GO:0002793) | 114 | 34 | 13.72 | + | 2.48 | 3.02E-07 | 3.27E-05 |
| neuron cellular homeostasis (GO:0070050) | 64 | 19 | 7.7 | + | 2.47 | 1.28E-04 | 6.37E-03 |
| response to axon injury (GO:0048678) | 54 | 16 | 6.5 | + | 2.46 | 4.36E-04 | 1.69E-02 |
| purine ribonucleoside triphosphate metabolic process (GO:0009205) | 186 | 55 | 22.39 | + | 2.46 | 1.49E-10 | 2.69E-08 |
| regulation of autophagy of mitochondrion (GO:1903146) | 44 | 13 | 5.3 | + | 2.45 | 1.51E-03 | 4.56E-02 |

| | | | | | | | |
|---|---|---|---|---|---|---|---|
| cardiac muscle cell contraction (GO:0086003) | 51 | 15 | 6.14 | + | 2.44 | 7.11E-04 | 2.51E-02 |
| regulation of potassium ion transport (GO:0043266) | 92 | 27 | 11.07 | + | 2.44 | 6.78E-06 | 5.34E-04 |
| hormone secretion (GO:0046879) | 92 | 27 | 11.07 | + | 2.44 | 6.78E-06 | 5.31E-04 |
| peptide secretion (GO:0002790) | 75 | 22 | 9.03 | + | 2.44 | 4.73E-05 | 2.81E-03 |
| nucleoside diphosphate catabolic process (GO:0009134) | 58 | 17 | 6.98 | + | 2.44 | 3.37E-04 | 1.37E-02 |
| mitochondrial transmembrane transport (GO:1990542) | 99 | 29 | 11.92 | + | 2.43 | 3.27E-06 | 2.78E-04 |
| regulation of sodium ion transport (GO:0002028) | 82 | 24 | 9.87 | + | 2.43 | 2.27E-05 | 1.55E-03 |
| response to unfolded protein (GO:0006986) | 120 | 35 | 14.44 | + | 2.42 | 5.68E-07 | 5.76E-05 |
| hormone transport (GO:0009914) | 103 | 30 | 12.4 | + | 2.42 | 2.54E-06 | 2.21E-04 |
| synaptic vesicle endocytosis (GO:0048488) | 55 | 16 | 6.62 | + | 2.42 | 5.47E-04 | 2.06E-02 |
| ribonucleoside triphosphate metabolic process (GO:0009199) | 193 | 56 | 23.23 | + | 2.41 | 1.83E-10 | 3.22E-08 |
| purine nucleoside triphosphate metabolic process (GO:0009144) | 193 | 56 | 23.23 | + | 2.41 | 1.83E-10 | 3.19E-08 |
| regulation of amine transport (GO:0051952) | 83 | 24 | 9.99 | + | 2.4 | 2.83E-05 | 1.84E-03 |
| regulation of protein targeting (GO:1903533) | 80 | 23 | 9.63 | + | 2.39 | 4.56E-05 | 2.72E-03 |
| presynaptic endocytosis (GO:0140238) | 56 | 16 | 6.74 | + | 2.37 | 6.81E-04 | 2.43E-02 |
| cardiac muscle cell action potential (GO:0086001) | 56 | 16 | 6.74 | + | 2.37 | 6.81E-04 | 2.42E-02 |

| | | | | | | | |
|---|---|---|---|---|---|---|---|
| purine-containing compound catabolic process (GO:0072523) | 105 | 30 | 12.64 | + | 2.37 | 5.29E-06 | 4.35E-04 |
| nucleotide biosynthetic process (GO:0009165) | 267 | 76 | 32.14 | + | 2.36 | 3.26E-13 | 9.66E-11 |
| protein transmembrane transport (GO:0071806) | 74 | 21 | 8.91 | + | 2.36 | 1.92E-04 | 8.84E-03 |
| positive regulation of hormone secretion (GO:0046887) | 145 | 41 | 17.45 | + | 2.35 | 1.07E-07 | 1.22E-05 |
| nucleoside phosphate biosynthetic process (GO:1901293) | 269 | 76 | 32.38 | + | 2.35 | 4.65E-13 | 1.30E-10 |
| proton transmembrane transport (GO:1902600) | 146 | 41 | 17.57 | + | 2.33 | 1.29E-07 | 1.46E-05 |
| regulation of smooth muscle contraction (GO:0006940) | 57 | 16 | 6.86 | + | 2.33 | 8.43E-04 | 2.85E-02 |
| purine nucleotide catabolic process (GO:0006195) | 100 | 28 | 12.04 | + | 2.33 | 1.50E-05 | 1.08E-03 |
| action potential (GO:0001508) | 125 | 35 | 15.04 | + | 2.33 | 1.18E-06 | 1.07E-04 |
| fatty acid derivative biosynthetic process (GO:1901570) | 54 | 15 | 6.5 | + | 2.31 | 1.36E-03 | 4.25E-02 |
| ribonucleoside diphosphate catabolic process (GO:0009191) | 54 | 15 | 6.5 | + | 2.31 | 1.36E-03 | 4.24E-02 |
| nucleoside triphosphate metabolic process (GO:0009141) | 209 | 58 | 25.16 | + | 2.31 | 5.96E-10 | 9.69E-08 |
| response to topologically incorrect protein (GO:0035966) | 137 | 38 | 16.49 | + | 2.3 | 5.14E-07 | 5.28E-05 |
| peptide transport (GO:0015833) | 87 | 24 | 10.47 | + | 2.29 | 7.95E-05 | 4.26E-03 |
| purine ribonucleotide catabolic process (GO:0009154) | 91 | 25 | 10.95 | + | 2.28 | 5.86E-05 | 3.37E-03 |
| glycogen metabolic process (GO:0005977) | 62 | 17 | 7.46 | + | 2.28 | 1.14E-03 | 3.64E-02 |

| | | | | | | | |
|---|---|---|---|---|---|---|---|
| regulation of potassium ion transmembrane transport (GO:1901379) | 77 | 21 | 9.27 | + | 2.27 | 2.69E-04 | 1.15E-02 |
| synaptic vesicle cycle (GO:0099504) | 144 | 39 | 17.33 | + | 2.25 | 8.68E-07 | 8.26E-05 |
| axo-dendritic transport (GO:0008088) | 74 | 20 | 8.91 | + | 2.25 | 4.22E-04 | 1.65E-02 |
| glycosphingolipid metabolic process (GO:0006687) | 63 | 17 | 7.58 | + | 2.24 | 1.27E-03 | 4.01E-02 |
| positive regulation of ion transmembrane transporter activity (GO:0032414) | 78 | 21 | 9.39 | + | 2.24 | 3.10E-04 | 1.30E-02 |
| synapse assembly (GO:0007416) | 119 | 32 | 14.32 | + | 2.23 | 1.17E-05 | 8.62E-04 |
| cardiac conduction (GO:0061337) | 67 | 18 | 8.06 | + | 2.23 | 9.12E-04 | 3.05E-02 |
| ribonucleotide catabolic process (GO:0009261) | 97 | 26 | 11.67 | + | 2.23 | 6.05E-05 | 3.45E-03 |
| vesicle-mediated transport in synapse (GO:0099003) | 154 | 41 | 18.54 | + | 2.21 | 6.41E-07 | 6.37E-05 |
| glucan metabolic process (GO:0044042) | 64 | 17 | 7.7 | + | 2.21 | 1.43E-03 | 4.44E-02 |
| cerebellum development (GO:0021549) | 106 | 28 | 12.76 | + | 2.19 | 5.86E-05 | 3.36E-03 |
| regulation of heart contraction (GO:0008016) | 197 | 52 | 23.71 | + | 2.19 | 2.99E-08 | 3.76E-06 |
| protein secretion (GO:0009306) | 133 | 35 | 16.01 | + | 2.19 | 5.77E-06 | 4.64E-04 |
| regulation of monoatomic ion transmembrane transporter activity (GO:0032412) | 198 | 52 | 23.83 | + | 2.18 | 3.47E-08 | 4.26E-06 |
| sensory perception of pain (GO:0019233) | 65 | 17 | 7.82 | + | 2.17 | 1.64E-03 | 4.86E-02 |
| multicellular organismal response to stress (GO:0033555) | 88 | 23 | 10.59 | + | 2.17 | 2.36E-04 | 1.04E-02 |

| | | | | | | | |
|---|---|---|---|---|---|---|---|
| protein localization to extracellular region (GO:0071692) | 142 | 37 | 17.09 | + | 2.16 | 5.98E-06 | 4.76E-04 |
| regulation of cardiac muscle contraction (GO:0055117) | 73 | 19 | 8.79 | + | 2.16 | 8.89E-04 | 2.99E-02 |
| protein folding (GO:0006457) | 223 | 58 | 26.84 | + | 2.16 | 9.29E-09 | 1.29E-06 |
| regulation of cation channel activity (GO:2001257) | 100 | 26 | 12.04 | + | 2.16 | 1.37E-04 | 6.67E-03 |
| mitochondrial membrane organization (GO:0007006) | 112 | 29 | 13.48 | + | 2.15 | 5.28E-05 | 3.06E-03 |
| nerve development (GO:0021675) | 97 | 25 | 11.67 | + | 2.14 | 2.10E-04 | 9.47E-03 |
| regulation of neurotransmitter transport (GO:0051588) | 101 | 26 | 12.16 | + | 2.14 | 1.52E-04 | 7.25E-03 |
| establishment of protein localization to extracellular region (GO:0035592) | 136 | 35 | 16.37 | + | 2.14 | 1.35E-05 | 9.77E-04 |
| regulation of insulin secretion (GO:0050796) | 171 | 44 | 20.58 | + | 2.14 | 7.29E-07 | 7.07E-05 |
| neurotransmitter transport (GO:0006836) | 140 | 36 | 16.85 | + | 2.14 | 9.67E-06 | 7.35E-04 |
| antigen processing and presentation of peptide antigen (GO:0048002) | 70 | 18 | 8.43 | + | 2.14 | 1.39E-03 | 4.33E-02 |
| purine-containing compound metabolic process (GO:0072521) | 498 | 128 | 59.94 | + | 2.14 | 2.97E-17 | 2.36E-14 |
| purine ribonucleotide metabolic process (GO:0009150) | 386 | 99 | 46.46 | + | 2.13 | 1.61E-13 | 4.87E-11 |
| purine nucleotide metabolic process (GO:0006163) | 468 | 120 | 56.33 | + | 2.13 | 2.92E-16 | 1.77E-13 |
| positive regulation of transporter activity (GO:0032411) | 86 | 22 | 10.35 | + | 2.13 | 6.23E-04 | 2.25E-02 |
| mitochondrial transport (GO:0006839) | 180 | 46 | 21.66 | + | 2.12 | 6.92E-07 | 6.75E-05 |

| | | | | | | | |
|---|---|---|---|---|---|---|---|
| ribonucleotide metabolic process (GO:0009259) | 406 | 103 | 48.87 | + | 2.11 | 1.12E-13 | 3.60E-11 |
| negative regulation of cation transmembrane transport (GO:1904063) | 79 | 20 | 9.51 | + | 2.1 | 1.33E-03 | 4.19E-02 |
| protein localization to cell junction (GO:1902414) | 87 | 22 | 10.47 | + | 2.1 | 6.79E-04 | 2.43E-02 |
| heart contraction (GO:0060047) | 99 | 25 | 11.92 | + | 2.1 | 2.62E-04 | 1.13E-02 |
| regulation of transmembrane transporter activity (GO:0022898) | 206 | 52 | 24.79 | + | 2.1 | 1.39E-07 | 1.55E-05 |
| ribose phosphate metabolic process (GO:0019693) | 414 | 104 | 49.83 | + | 2.09 | 1.23E-13 | 3.80E-11 |
| metencephalon development (GO:0022037) | 116 | 29 | 13.96 | + | 2.08 | 1.34E-04 | 6.55E-03 |
| regulation of neurotransmitter secretion (GO:0046928) | 84 | 21 | 10.11 | + | 2.08 | 1.04E-03 | 3.36E-02 |
| regulation of G protein-coupled receptor signaling pathway (GO:0008277) | 157 | 39 | 18.9 | + | 2.06 | 1.00E-05 | 7.57E-04 |
| endoplasmic reticulum to Golgi vesicle-mediated transport (GO:0006888) | 125 | 31 | 15.04 | + | 2.06 | 7.66E-05 | 4.16E-03 |
| heart process (GO:0003015) | 113 | 28 | 13.6 | + | 2.06 | 2.01E-04 | 9.15E-03 |
| positive regulation of secretion by cell (GO:1903532) | 291 | 72 | 35.02 | + | 2.06 | 1.65E-09 | 2.50E-07 |
| adult locomotory behavior (GO:0008344) | 81 | 20 | 9.75 | + | 2.05 | 1.58E-03 | 4.74E-02 |
| regulation of transporter activity (GO:0032409) | 219 | 54 | 26.36 | + | 2.05 | 1.88E-07 | 2.08E-05 |
| positive regulation of secretion (GO:0051047) | 313 | 77 | 37.67 | + | 2.04 | 6.55E-10 | 1.05E-07 |
| regulation of peptide transport (GO:0090087) | 204 | 50 | 24.55 | + | 2.04 | 6.86E-07 | 6.78E-05 |

| | | | | | | | |
|---|---|---|---|---|---|---|---|
| regulation of peptide secretion (GO:0002791) | 204 | 50 | 24.55 | + | 2.04 | 6.86E-07 | 6.73E-05 |
| positive regulation of protein secretion (GO:0050714) | 147 | 36 | 17.69 | + | 2.03 | 3.07E-05 | 1.97E-03 |
| mitochondrion organization (GO:0007005) | 442 | 108 | 53.2 | + | 2.03 | 4.20E-13 | 1.20E-10 |
| organophosphate biosynthetic process (GO:0090407) | 549 | 134 | 66.08 | + | 2.03 | 3.96E-16 | 2.31E-13 |
| nucleotide catabolic process (GO:0009166) | 123 | 30 | 14.8 | + | 2.03 | 1.29E-04 | 6.31E-03 |
| regulation of peptide hormone secretion (GO:0090276) | 201 | 49 | 24.19 | + | 2.03 | 1.03E-06 | 9.59E-05 |
| negative regulation of monoatomic ion transport (GO:0043271) | 115 | 28 | 13.84 | + | 2.02 | 2.38E-04 | 1.05E-02 |
| signal release (GO:0023061) | 189 | 46 | 22.75 | + | 2.02 | 2.54E-06 | 2.20E-04 |
| neuropeptide signaling pathway (GO:0007218) | 111 | 27 | 13.36 | + | 2.02 | 3.27E-04 | 1.35E-02 |
| nucleoside phosphate metabolic process (GO:0006753) | 536 | 130 | 64.51 | + | 2.02 | 1.84E-15 | 8.99E-13 |
| nucleoside phosphate catabolic process (GO:1901292) | 132 | 32 | 15.89 | + | 2.01 | 1.22E-04 | 6.12E-03 |
| nucleotide metabolic process (GO:0009117) | 528 | 128 | 63.55 | + | 2.01 | 3.32E-15 | 1.48E-12 |
| ERAD pathway (GO:0036503) | 95 | 23 | 11.43 | + | 2.01 | 1.22E-03 | 3.88E-02 |
| regulation of mitochondrion organization (GO:0010821) | 149 | 36 | 17.93 | + | 2.01 | 3.66E-05 | 2.28E-03 |
| negative regulation of monoatomic ion transmembrane transport (GO:0034766) | 87 | 21 | 10.47 | + | 2.01 | 1.45E-03 | 4.42E-02 |
| chemical synaptic transmission (GO:0007268) | 415 | 100 | 49.95 | + | 2 | 7.59E-12 | 1.88E-09 |

| | | | | | | | |
|---|---|---|---|---|---|---|---|
| anterograde trans-synaptic signaling (GO:0098916) | 415 | 100 | 49.95 | + | 2 | 7.59E-12 | 1.85E-09 |
| negative regulation of establishment of protein localization (GO:1904950) | 125 | 30 | 15.04 | + | 1.99 | 2.50E-04 | 1.09E-02 |
| regulation of muscle contraction (GO:0006937) | 163 | 39 | 19.62 | + | 1.99 | 2.70E-05 | 1.78E-03 |
| organophosphate catabolic process (GO:0046434) | 184 | 44 | 22.15 | + | 1.99 | 6.31E-06 | 5.00E-04 |
| trans-synaptic signaling (GO:0099537) | 432 | 103 | 52 | + | 1.98 | 6.88E-12 | 1.73E-09 |
| phosphatidylinositol biosynthetic process (GO:0006661) | 126 | 30 | 15.17 | + | 1.98 | 2.63E-04 | 1.13E-02 |
| pyridine nucleotide metabolic process (GO:0019362) | 118 | 28 | 14.2 | + | 1.97 | 3.32E-04 | 1.37E-02 |
| nicotinamide nucleotide metabolic process (GO:0046496) | 118 | 28 | 14.2 | + | 1.97 | 3.32E-04 | 1.36E-02 |
| regulation of blood circulation (GO:1903522) | 249 | 59 | 29.97 | + | 1.97 | 3.49E-07 | 3.66E-05 |
| glycolipid metabolic process (GO:0006664) | 106 | 25 | 12.76 | + | 1.96 | 8.10E-04 | 2.78E-02 |
| regulation of synaptic plasticity (GO:0048167) | 204 | 48 | 24.55 | + | 1.95 | 3.94E-06 | 3.33E-04 |
| nucleobase-containing small molecule metabolic process (GO:0055086) | 605 | 142 | 72.82 | + | 1.95 | 1.94E-15 | 9.17E-13 |
| carbohydrate derivative catabolic process (GO:1901136) | 222 | 52 | 26.72 | + | 1.95 | 2.01E-06 | 1.75E-04 |
| liposaccharide metabolic process (GO:1903509) | 107 | 25 | 12.88 | + | 1.94 | 9.00E-04 | 3.02E-02 |
| positive regulation of synaptic transmission (GO:0050806) | 143 | 33 | 17.21 | + | 1.92 | 2.33E-04 | 1.03E-02 |
| antigen processing and presentation (GO:0019882) | 104 | 24 | 12.52 | + | 1.92 | 1.35E-03 | 4.23E-02 |

| | | | | | | | |
|---|---|---|---|---|---|---|---|
| negative regulation of protein localization (GO:1903828) | 209 | 48 | 25.16 | + | 1.91 | 8.55E-06 | 6.56E-04 |
| synaptic signaling (GO:0099536) | 462 | 106 | 55.61 | + | 1.91 | 3.63E-11 | 7.74E-09 |
| hindbrain development (GO:0030902) | 157 | 36 | 18.9 | + | 1.91 | 1.11E-04 | 5.68E-03 |
| regulation of sequestering of calcium ion (GO:0051282) | 131 | 30 | 15.77 | + | 1.9 | 4.06E-04 | 1.60E-02 |
| regulation of monoatomic ion transmembrane transport (GO:0034765) | 333 | 76 | 40.08 | + | 1.9 | 3.27E-08 | 4.06E-06 |
| mitochondrial translation (GO:0032543) | 110 | 25 | 13.24 | + | 1.89 | 1.67E-03 | 4.93E-02 |
| regulation of reactive oxygen species metabolic process (GO:2000377) | 141 | 32 | 16.97 | + | 1.89 | 3.64E-04 | 1.47E-02 |
| export from cell (GO:0140352) | 490 | 111 | 58.98 | + | 1.88 | 3.12E-11 | 6.83E-09 |
| regulation of monoatomic cation transmembrane transport (GO:1904062) | 297 | 67 | 35.75 | + | 1.87 | 3.21E-07 | 3.40E-05 |
| modulation of chemical synaptic transmission (GO:0050804) | 479 | 108 | 57.65 | + | 1.87 | 8.23E-11 | 1.60E-08 |
| regulation of trans-synaptic signaling (GO:0099177) | 480 | 108 | 57.77 | + | 1.87 | 8.67E-11 | 1.64E-08 |
| positive regulation of monoatomic ion transport (GO:0043270) | 205 | 46 | 24.67 | + | 1.86 | 3.09E-05 | 1.97E-03 |
| positive regulation of calcium ion transport (GO:0051928) | 116 | 26 | 13.96 | + | 1.86 | 1.44E-03 | 4.42E-02 |
| pyridine-containing compound metabolic process (GO:0072524) | 125 | 28 | 15.04 | + | 1.86 | 1.25E-03 | 3.98E-02 |
| secretion by cell (GO:0032940) | 425 | 95 | 51.15 | + | 1.86 | 1.58E-09 | 2.41E-07 |
| glycerophospholipid biosynthetic process (GO:0046474) | 202 | 45 | 24.31 | + | 1.85 | 4.41E-05 | 2.64E-03 |

| | | | | | | | |
|---|---|---|---|---|---|---|---|
| regulation of monoatomic ion transport (GO:0043269) | 449 | 100 | 54.04 | + | 1.85 | 6.87E-10 | 1.08E-07 |
| negative regulation of transmembrane transport (GO:0034763) | 117 | 26 | 14.08 | + | 1.85 | 1.58E-03 | 4.74E-02 |
| amide biosynthetic process (GO:0043604) | 145 | 32 | 17.45 | + | 1.83 | 7.14E-04 | 2.51E-02 |
| organophosphate metabolic process (GO:0019637) | 940 | 207 | 113.14 | + | 1.83 | 9.67E-19 | 9.75E-16 |
| hexose metabolic process (GO:0019318) | 159 | 35 | 19.14 | + | 1.83 | 3.34E-04 | 1.37E-02 |
| regulation of metal ion transport (GO:0010959) | 378 | 83 | 45.5 | + | 1.82 | 3.95E-08 | 4.78E-06 |
| protein localization to cell periphery (GO:1990778) | 237 | 52 | 28.53 | + | 1.82 | 1.88E-05 | 1.30E-03 |
| sphingolipid metabolic process (GO:0006665) | 160 | 35 | 19.26 | + | 1.82 | 5.28E-04 | 2.01E-02 |
| regulation of protein localization to membrane (GO:1905475) | 184 | 40 | 22.15 | + | 1.81 | 2.20E-04 | 9.82E-03 |
| regulation of hormone secretion (GO:0046883) | 258 | 56 | 31.05 | + | 1.8 | 1.12E-05 | 8.32E-04 |
| carbohydrate biosynthetic process (GO:0016051) | 129 | 28 | 15.53 | + | 1.8 | 1.63E-03 | 4.85E-02 |
| regulated exocytosis (GO:0045055) | 143 | 31 | 17.21 | + | 1.8 | 1.08E-03 | 3.47E-02 |
| intracellular calcium ion homeostasis (GO:0006874) | 199 | 43 | 23.95 | + | 1.8 | 1.11E-04 | 5.70E-03 |
| regulation of secretion by cell (GO:1903530) | 547 | 118 | 65.84 | + | 1.79 | 1.58E-10 | 2.81E-08 |
| monosaccharide metabolic process (GO:0005996) | 181 | 39 | 21.79 | + | 1.79 | 3.09E-04 | 1.30E-02 |
| carbohydrate derivative biosynthetic process (GO:1901137) | 604 | 130 | 72.7 | + | 1.79 | 2.33E-11 | 5.34E-09 |

| | | | | | | | |
|---|---|---|---|---|---|---|---|
| phosphatidylinositol metabolic process (GO:0046488) | 158 | 34 | 19.02 | + | 1.79 | 7.75E-04 | 2.68E-02 |
| phospholipid biosynthetic process (GO:0008654) | 242 | 52 | 29.13 | + | 1.79 | 2.55E-05 | 1.70E-03 |
| protein localization to plasma membrane (GO:0072659) | 196 | 42 | 23.59 | + | 1.78 | 2.25E-04 | 1.00E-02 |
| regulation of exocytosis (GO:0017157) | 173 | 37 | 20.82 | + | 1.78 | 5.61E-04 | 2.10E-02 |
| regulation of transmembrane transport (GO:0034762) | 459 | 98 | 55.25 | + | 1.77 | 1.06E-08 | 1.44E-06 |
| regulation of secretion (GO:0051046) | 600 | 128 | 72.22 | + | 1.77 | 7.42E-11 | 1.48E-08 |
| regulation of membrane potential (GO:0042391) | 456 | 97 | 54.88 | + | 1.77 | 1.51E-08 | 1.98E-06 |
| sulfur compound biosynthetic process (GO:0044272) | 160 | 34 | 19.26 | + | 1.77 | 8.65E-04 | 2.91E-02 |
| establishment of protein localization to membrane (GO:0090150) | 226 | 48 | 27.2 | + | 1.76 | 7.80E-05 | 4.21E-03 |
| regulation of macroautophagy (GO:0016241) | 184 | 39 | 22.15 | + | 1.76 | 3.68E-04 | 1.47E-02 |
| vesicle localization (GO:0051648) | 185 | 39 | 22.27 | + | 1.75 | 3.95E-04 | 1.57E-02 |
| inorganic cation transmembrane transport (GO:0098662) | 638 | 134 | 76.79 | + | 1.75 | 7.86E-11 | 1.54E-08 |
| glycerolipid biosynthetic process (GO:0045017) | 224 | 47 | 26.96 | + | 1.74 | 1.18E-04 | 5.93E-03 |
| monoatomic cation transmembrane transport (GO:0098655) | 653 | 137 | 78.59 | + | 1.74 | 5.63E-11 | 1.15E-08 |
| organic hydroxy compound transport (GO:0015850) | 167 | 35 | 20.1 | + | 1.74 | 1.09E-03 | 3.51E-02 |
| membrane lipid metabolic process (GO:0006643) | 206 | 43 | 24.79 | + | 1.73 | 3.24E-04 | 1.35E-02 |

| | | | | | | | |
|---|---|---|---|---|---|---|---|
| inorganic ion transmembrane transport (GO:0098660) | 733 | 153 | 88.22 | + | 1.73 | 5.26E-12 | 1.35E-09 |
| response to temperature stimulus (GO:0009266) | 163 | 34 | 19.62 | + | 1.73 | 1.46E-03 | 4.44E-02 |
| response to endoplasmic reticulum stress (GO:0034976) | 226 | 47 | 27.2 | + | 1.73 | 1.81E-04 | 8.38E-03 |
| synapse organization (GO:0050808) | 337 | 70 | 40.56 | + | 1.73 | 4.15E-06 | 3.47E-04 |
| regulation of protein secretion (GO:0050708) | 260 | 54 | 31.29 | + | 1.73 | 4.95E-05 | 2.92E-03 |
| sensory perception of mechanical stimulus (GO:0050954) | 183 | 38 | 22.03 | + | 1.73 | 8.08E-04 | 2.78E-02 |
| monoatomic ion transmembrane transport (GO:0034220) | 815 | 168 | 98.09 | + | 1.71 | 1.20E-12 | 3.19E-10 |
| positive regulation of establishment of protein localization (GO:1904951) | 330 | 68 | 39.72 | + | 1.71 | 7.58E-06 | 5.85E-04 |
| regulation of establishment of protein localization (GO:0070201) | 529 | 109 | 63.67 | + | 1.71 | 1.37E-08 | 1.82E-06 |
| establishment of vesicle localization (GO:0051650) | 170 | 35 | 20.46 | + | 1.71 | 1.28E-03 | 4.04E-02 |
| regulation of muscle system process (GO:0090257) | 229 | 47 | 27.56 | + | 1.71 | 2.11E-04 | 9.50E-03 |
| Golgi vesicle transport (GO:0048193) | 288 | 59 | 34.66 | + | 1.7 | 3.53E-05 | 2.20E-03 |
| carbohydrate derivative metabolic process (GO:1901135) | 997 | 204 | 120 | + | 1.7 | 8.26E-15 | 3.20E-12 |
| blood circulation (GO:0008015) | 411 | 84 | 49.47 | + | 1.7 | 1.09E-06 | 9.98E-05 |
| monoatomic cation transport (GO:0006812) | 787 | 160 | 94.72 | + | 1.69 | 1.31E-11 | 3.05E-09 |
| locomotory behavior (GO:0007626) | 197 | 40 | 23.71 | + | 1.69 | 8.58E-04 | 2.90E-02 |

| | | | | | | | |
|---|---|---|---|---|---|---|---|
| transmembrane transport (GO:0055085) | 1283 | 260 | 154.42 | + | 1.68 | 3.85E-18 | 3.43E-15 |
| protein localization to membrane (GO:0072657) | 459 | 93 | 55.25 | + | 1.68 | 4.28E-07 | 4.44E-05 |
| exocytosis (GO:0006887) | 243 | 49 | 29.25 | + | 1.68 | 3.09E-04 | 1.30E-02 |
| positive regulation of transport (GO:0051050) | 835 | 168 | 100.5 | + | 1.67 | 1.07E-11 | 2.56E-09 |
| protein maturation (GO:0051604) | 494 | 99 | 59.46 | + | 1.67 | 2.57E-07 | 2.80E-05 |
| regulation of synapse structure or activity (GO:0050803) | 255 | 51 | 30.69 | + | 1.66 | 2.96E-04 | 1.25E-02 |
| monoatomic ion transport (GO:0006811) | 980 | 195 | 117.95 | + | 1.65 | 5.68E-13 | 1.56E-10 |
| intracellular protein transport (GO:0006886) | 558 | 111 | 67.16 | + | 1.65 | 7.16E-08 | 8.40E-06 |
| calcium ion transport (GO:0006816) | 262 | 52 | 31.53 | + | 1.65 | 2.53E-04 | 1.10E-02 |
| calcium ion homeostasis (GO:0055074) | 227 | 45 | 27.32 | + | 1.65 | 6.58E-04 | 2.36E-02 |
| secretion (GO:0046903) | 545 | 108 | 65.6 | + | 1.65 | 1.52E-07 | 1.68E-05 |
| glycerophospholipid metabolic process (GO:0006650) | 293 | 58 | 35.27 | + | 1.64 | 1.28E-04 | 6.35E-03 |
| regulation of protein localization (GO:0032880) | 868 | 171 | 104.47 | + | 1.64 | 3.83E-11 | 8.04E-09 |
| positive regulation of protein localization (GO:1903829) | 484 | 95 | 58.25 | + | 1.63 | 1.26E-06 | 1.14E-04 |
| axon development (GO:0061564) | 420 | 82 | 50.55 | + | 1.62 | 9.36E-06 | 7.15E-04 |
| muscle contraction (GO:0006936) | 236 | 46 | 28.4 | + | 1.62 | 8.43E-04 | 2.86E-02 |

| | | | | | | | |
|---|---|---|---|---|---|---|---|
| axonogenesis (GO:0007409) | 366 | 71 | 44.05 | + | 1.61 | 4.43E-05 | 2.65E-03 |
| circulatory system process (GO:0003013) | 501 | 97 | 60.3 | + | 1.61 | 1.90E-06 | 1.68E-04 |
| response to wounding (GO:0009611) | 429 | 83 | 51.63 | + | 1.61 | 1.17E-05 | 8.64E-04 |
| regulation of calcium ion transport (GO:0051924) | 243 | 47 | 29.25 | + | 1.61 | 9.73E-04 | 3.19E-02 |
| cell junction assembly (GO:0034329) | 285 | 55 | 34.3 | + | 1.6 | 4.44E-04 | 1.72E-02 |
| regulation of synapse organization (GO:0050807) | 249 | 48 | 29.97 | + | 1.6 | 8.24E-04 | 2.81E-02 |
| positive regulation of nervous system development (GO:0051962) | 291 | 56 | 35.02 | + | 1.6 | 3.64E-04 | 1.46E-02 |
| phospholipid metabolic process (GO:0006644) | 364 | 70 | 43.81 | + | 1.6 | 6.20E-05 | 3.51E-03 |
| regulation of protein transport (GO:0051223) | 427 | 82 | 51.39 | + | 1.6 | 1.61E-05 | 1.15E-03 |
| positive regulation of protein transport (GO:0051222) | 250 | 48 | 30.09 | + | 1.6 | 1.14E-03 | 3.63E-02 |
| regulation of transport (GO:0051049) | 1576 | 300 | 189.69 | + | 1.58 | 6.47E-17 | 4.08E-14 |
| muscle system process (GO:0003012) | 289 | 55 | 34.78 | + | 1.58 | 5.07E-04 | 1.93E-02 |
| protein targeting (GO:0006605) | 247 | 47 | 29.73 | + | 1.58 | 1.51E-03 | 4.56E-02 |
| localization within membrane (GO:0051668) | 548 | 104 | 65.96 | + | 1.58 | 1.99E-06 | 1.75E-04 |
| negative regulation of transport (GO:0051051) | 406 | 77 | 48.87 | + | 1.58 | 5.45E-05 | 3.14E-03 |
| autophagy (GO:0006914) | 327 | 62 | 39.36 | + | 1.58 | 2.89E-04 | 1.23E-02 |

| | | | | | | | |
|---|---|---|---|---|---|---|---|
| process utilizing autophagic mechanism (GO:0061919) | 327 | 62 | 39.36 | + | 1.58 | 2.89E-04 | 1.22E-02 |
| regulation of localization (GO:0032879) | 1984 | 374 | 238.79 | + | 1.57 | 2.28E-20 | 4.31E-17 |
| regulation of cellular localization (GO:0060341) | 978 | 184 | 117.71 | + | 1.56 | 3.36E-10 | 5.71E-08 |
| phosphate-containing compound metabolic process (GO:0006796) | 1579 | 297 | 190.05 | + | 1.56 | 5.31E-16 | 2.97E-13 |
| response to oxidative stress (GO:0006979) | 362 | 68 | 43.57 | + | 1.56 | 1.66E-04 | 7.80E-03 |
| cell-cell signaling (GO:0007267) | 831 | 156 | 100.02 | + | 1.56 | 1.21E-08 | 1.62E-06 |
| regulation of hormone levels (GO:0010817) | 544 | 102 | 65.48 | + | 1.56 | 4.84E-06 | 4.00E-04 |
| cell morphogenesis involved in neuron differentiation (GO:0048667) | 443 | 83 | 53.32 | + | 1.56 | 4.35E-05 | 2.63E-03 |
| metal ion transport (GO:0030001) | 646 | 121 | 77.75 | + | 1.56 | 5.60E-07 | 5.72E-05 |

**NPHP1_Up:**

| | |
|---|---|
| Analysis Type: | PANTHER Overrepresentation Test (Released 20240807) |
| Annotation Version and Release Date: | GO Ontology database DOI: 10.5281/zenodo.12173881 Released 2024-06-17 |
| Analyzed List: | upload_1 (Homo sapiens) |
| Reference List: | Homo sapiens (all genes in database) |
| Test Type: | FISHER |
| Correction: | FDR |

| GO biological process complete | Homo sapiens - REFLIST (20580) | upload_1 (2372) | upload_1 (expected) | upload_1 (over/under) | upload_1 (fold Enrichment) | upload_1 (raw P-value) | upload_1 (FDR) |
|---|---|---|---|---|---|---|---|
| rhombomere 3 development (GO:0021569) | 3 | 3 | 0.35 | + | 8.68 | 1.53E-03 | 2.01E-02 |
| midbrain-hindbrain boundary morphogenesis (GO:0021555) | 3 | 3 | 0.35 | + | 8.68 | 1.53E-03 | 2.01E-02 |
| metanephric glomerular capillary formation (GO:0072277) | 3 | 3 | 0.35 | + | 8.68 | 1.53E-03 | 2.01E-02 |
| metanephric glomerulus vasculature morphogenesis (GO:0072276) | 3 | 3 | 0.35 | + | 8.68 | 1.53E-03 | 2.00E-02 |
| metanephric glomerulus morphogenesis (GO:0072275) | 3 | 3 | 0.35 | + | 8.68 | 1.53E-03 | 2.00E-02 |
| glomerular capillary formation (GO:0072104) | 6 | 6 | 0.69 | + | 8.68 | 2.33E-06 | 6.69E-05 |
| glomerulus vasculature morphogenesis (GO:0072103) | 6 | 6 | 0.69 | + | 8.68 | 2.33E-06 | 6.68E-05 |
| response to endothelin (GO:1990839) | 3 | 3 | 0.35 | + | 8.68 | 1.53E-03 | 2.00E-02 |
| cellular response to heparin (GO:0071504) | 4 | 4 | 0.46 | + | 8.68 | 1.76E-04 | 3.16E-03 |
| response to water-immersion restraint stress (GO:1990785) | 3 | 3 | 0.35 | + | 8.68 | 1.53E-03 | 2.00E-02 |
| cellular response to tumor cell (GO:0071228) | 4 | 4 | 0.46 | + | 8.68 | 1.76E-04 | 3.15E-03 |

| | | | | | | | |
|---|---|---|---|---|---|---|---|
| positive regulation of macrophage apoptotic process (GO:2000111) | 3 | 3 | 0.35 | + | 8.68 | 1.53E-03 | 2.00E-02 |
| N-terminal peptidyl-lysine acetylation (GO:0018076) | 3 | 3 | 0.35 | + | 8.68 | 1.53E-03 | 2.00E-02 |
| retrograde trans-synaptic signaling by trans-synaptic protein complex (GO:0098942) | 3 | 3 | 0.35 | + | 8.68 | 1.53E-03 | 1.99E-02 |
| regulation of cell proliferation in midbrain (GO:1904933) | 3 | 3 | 0.35 | + | 8.68 | 1.53E-03 | 1.99E-02 |
| melanocyte migration (GO:0097324) | 3 | 3 | 0.35 | + | 8.68 | 1.53E-03 | 1.99E-02 |
| lymphatic endothelial cell fate commitment (GO:0060838) | 3 | 3 | 0.35 | + | 8.68 | 1.53E-03 | 1.99E-02 |
| glomerulus morphogenesis (GO:0072102) | 9 | 8 | 1.04 | + | 7.71 | 2.49E-07 | 8.48E-06 |
| kidney vasculature morphogenesis (GO:0061439) | 8 | 7 | 0.92 | + | 7.59 | 1.93E-06 | 5.65E-05 |
| renal system vasculature morphogenesis (GO:0061438) | 8 | 7 | 0.92 | + | 7.59 | 1.93E-06 | 5.64E-05 |
| intramembranous ossification (GO:0001957) | 7 | 6 | 0.81 | + | 7.44 | 1.47E-05 | 3.61E-04 |
| direct ossification (GO:0036072) | 7 | 6 | 0.81 | + | 7.44 | 1.47E-05 | 3.60E-04 |
| renal vesicle formation (GO:0072033) | 6 | 5 | 0.69 | + | 7.23 | 1.10E-04 | 2.14E-03 |
| response to heparin (GO:0071503) | 6 | 5 | 0.69 | + | 7.23 | 1.10E-04 | 2.14E-03 |
| chemorepulsion of axon (GO:0061643) | 6 | 5 | 0.69 | + | 7.23 | 1.10E-04 | 2.13E-03 |
| regulation of basement membrane organization (GO:0110011) | 11 | 9 | 1.27 | + | 7.1 | 1.57E-07 | 5.60E-06 |
| glossopharyngeal nerve development (GO:0021563) | 5 | 4 | 0.58 | + | 6.94 | 7.99E-04 | 1.16E-02 |

| | | | | | | | |
|---|---|---|---|---|---|---|---|
| metanephric mesenchymal cell differentiation (GO:0072162) | 5 | 4 | 0.58 | + | 6.94 | 7.99E-04 | 1.16E-02 |
| tendon development (GO:0035989) | 5 | 4 | 0.58 | + | 6.94 | 7.99E-04 | 1.16E-02 |
| carbon catabolite activation of transcription (GO:0045991) | 8 | 6 | 0.92 | + | 6.51 | 5.31E-05 | 1.15E-03 |
| carbon catabolite regulation of transcription (GO:0045990) | 8 | 6 | 0.92 | + | 6.51 | 5.31E-05 | 1.15E-03 |
| positive regulation of transcription by glucose (GO:0046016) | 7 | 5 | 0.81 | + | 6.2 | 3.48E-04 | 5.66E-03 |
| anterior/posterior axon guidance (GO:0033564) | 7 | 5 | 0.81 | + | 6.2 | 3.48E-04 | 5.65E-03 |
| regulation of epithelial cell proliferation involved in lung morphogenesis (GO:2000794) | 7 | 5 | 0.81 | + | 6.2 | 3.48E-04 | 5.65E-03 |
| metanephric distal tubule development (GO:0072235) | 7 | 5 | 0.81 | + | 6.2 | 3.48E-04 | 5.64E-03 |
| cell proliferation involved in metanephros development (GO:0072203) | 7 | 5 | 0.81 | + | 6.2 | 3.48E-04 | 5.63E-03 |
| carbon catabolite activation of transcription from RNA polymerase II promoter (GO:0000436) | 7 | 5 | 0.81 | + | 6.2 | 3.48E-04 | 5.63E-03 |
| carbon catabolite regulation of transcription from RNA polymerase II promoter (GO:0000429) | 7 | 5 | 0.81 | + | 6.2 | 3.48E-04 | 5.62E-03 |
| hematopoietic stem cell migration (GO:0035701) | 7 | 5 | 0.81 | + | 6.2 | 3.48E-04 | 5.62E-03 |
| cell proliferation involved in kidney development (GO:0072111) | 10 | 7 | 1.15 | + | 6.07 | 2.34E-05 | 5.52E-04 |
| lung-associated mesenchyme development (GO:0060484) | 9 | 6 | 1.04 | + | 5.78 | 1.44E-04 | 2.67E-03 |
| galactolipid biosynthetic process (GO:0019375) | 6 | 4 | 0.69 | + | 5.78 | 2.18E-03 | 2.67E-02 |
| positive regulation of mammary gland epithelial cell proliferation (GO:0033601) | 6 | 4 | 0.69 | + | 5.78 | 2.18E-03 | 2.67E-02 |

| | | | | | | | |
|---|---|---|---|---|---|---|---|
| optic nerve morphogenesis (GO:0021631) | 6 | 4 | 0.69 | + | 5.78 | 2.18E-03 | 2.67E-02 |
| positive regulation of peptidyl-lysine acetylation (GO:2000758) | 6 | 4 | 0.69 | + | 5.78 | 2.18E-03 | 2.66E-02 |
| mesonephric duct development (GO:0072177) | 9 | 6 | 1.04 | + | 5.78 | 1.44E-04 | 2.67E-03 |
| galactosylceramide biosynthetic process (GO:0006682) | 6 | 4 | 0.69 | + | 5.78 | 2.18E-03 | 2.66E-02 |
| inhibition of neuroepithelial cell differentiation (GO:0002085) | 6 | 4 | 0.69 | + | 5.78 | 2.18E-03 | 2.66E-02 |
| luteolysis (GO:0001554) | 6 | 4 | 0.69 | + | 5.78 | 2.18E-03 | 2.66E-02 |
| retrograde trans-synaptic signaling (GO:0098917) | 6 | 4 | 0.69 | + | 5.78 | 2.18E-03 | 2.66E-02 |
| bone marrow development (GO:0048539) | 9 | 6 | 1.04 | + | 5.78 | 1.44E-04 | 2.66E-03 |
| dopaminergic neuron axon guidance (GO:0036514) | 6 | 4 | 0.69 | + | 5.78 | 2.18E-03 | 2.65E-02 |
| positive regulation of transcription from RNA polymerase II promoter by glucose (GO:0000432) | 6 | 4 | 0.69 | + | 5.78 | 2.18E-03 | 2.65E-02 |
| regulation of transcription from RNA polymerase II promoter by glucose (GO:0000430) | 6 | 4 | 0.69 | + | 5.78 | 2.18E-03 | 2.65E-02 |
| regulation of myofibroblast differentiation (GO:1904760) | 6 | 4 | 0.69 | + | 5.78 | 2.18E-03 | 2.65E-02 |
| positive regulation of hemoglobin biosynthetic process (GO:0046985) | 6 | 4 | 0.69 | + | 5.78 | 2.18E-03 | 2.65E-02 |
| positive regulation of cytoplasmic mRNA processing body assembly (GO:0010606) | 6 | 4 | 0.69 | + | 5.78 | 2.18E-03 | 2.64E-02 |
| planar cell polarity pathway involved in axis elongation (GO:0003402) | 6 | 4 | 0.69 | + | 5.78 | 2.18E-03 | 2.64E-02 |
| regulation of aspartic-type endopeptidase activity involved in amyloid precursor protein catabolic process (GO:1902959) | 6 | 4 | 0.69 | + | 5.78 | 2.18E-03 | 2.64E-02 |

| | | | | | | | |
|---|---|---|---|---|---|---|---|
| regulation of branching involved in lung morphogenesis (GO:0061046) | 6 | 4 | 0.69 | + | 5.78 | 2.18E-03 | 2.64E-02 |
| atrioventricular node development (GO:0003162) | 9 | 6 | 1.04 | + | 5.78 | 1.44E-04 | 2.66E-03 |
| negative regulation of neuron migration (GO:2001223) | 11 | 7 | 1.27 | + | 5.52 | 5.80E-05 | 1.23E-03 |
| metanephric glomerulus development (GO:0072224) | 11 | 7 | 1.27 | + | 5.52 | 5.80E-05 | 1.23E-03 |
| neuron projection extension involved in neuron projection guidance (GO:1902284) | 11 | 7 | 1.27 | + | 5.52 | 5.80E-05 | 1.23E-03 |
| axon extension involved in axon guidance (GO:0048846) | 11 | 7 | 1.27 | + | 5.52 | 5.80E-05 | 1.23E-03 |
| somatic stem cell division (GO:0048103) | 11 | 7 | 1.27 | + | 5.52 | 5.80E-05 | 1.23E-03 |
| rhombomere development (GO:0021546) | 8 | 5 | 0.92 | + | 5.42 | 8.40E-04 | 1.21E-02 |
| metanephric nephron tubule morphogenesis (GO:0072282) | 8 | 5 | 0.92 | + | 5.42 | 8.40E-04 | 1.21E-02 |
| positive regulation of vascular endothelial growth factor signaling pathway (GO:1900748) | 8 | 5 | 0.92 | + | 5.42 | 8.40E-04 | 1.21E-02 |
| otic vesicle formation (GO:0030916) | 8 | 5 | 0.92 | + | 5.42 | 8.40E-04 | 1.21E-02 |
| ventricular compact myocardium morphogenesis (GO:0003223) | 8 | 5 | 0.92 | + | 5.42 | 8.40E-04 | 1.20E-02 |
| membranous septum morphogenesis (GO:0003149) | 8 | 5 | 0.92 | + | 5.42 | 8.40E-04 | 1.20E-02 |
| commissural neuron axon guidance (GO:0071679) | 13 | 8 | 1.5 | + | 5.34 | 2.32E-05 | 5.48E-04 |
| renal vesicle development (GO:0072087) | 13 | 8 | 1.5 | + | 5.34 | 2.32E-05 | 5.48E-04 |
| nephric duct development (GO:0072176) | 15 | 9 | 1.73 | + | 5.21 | 9.23E-06 | 2.37E-04 |

| | | | | | | | |
|---|---|---|---|---|---|---|---|
| foregut morphogenesis (GO:0007440) | 10 | 6 | 1.15 | + | 5.21 | 3.24E-04 | 5.33E-03 |
| regulation of transcription by glucose (GO:0046015) | 10 | 6 | 1.15 | + | 5.21 | 3.24E-04 | 5.33E-03 |
| metanephric tubule morphogenesis (GO:0072173) | 10 | 6 | 1.15 | + | 5.21 | 3.24E-04 | 5.32E-03 |
| podocyte development (GO:0072015) | 10 | 6 | 1.15 | + | 5.21 | 3.24E-04 | 5.31E-03 |
| luteinization (GO:0001553) | 10 | 6 | 1.15 | + | 5.21 | 3.24E-04 | 5.31E-03 |
| elastic fiber assembly (GO:0048251) | 10 | 6 | 1.15 | + | 5.21 | 3.24E-04 | 5.30E-03 |
| glomerulus vasculature development (GO:0072012) | 22 | 13 | 2.54 | + | 5.13 | 1.11E-07 | 4.05E-06 |
| regulation of nephron tubule epithelial cell differentiation (GO:0072182) | 12 | 7 | 1.38 | + | 5.06 | 1.25E-04 | 2.37E-03 |
| positive regulation of integrin-mediated signaling pathway (GO:2001046) | 12 | 7 | 1.38 | + | 5.06 | 1.25E-04 | 2.37E-03 |
| renal vesicle morphogenesis (GO:0072077) | 12 | 7 | 1.38 | + | 5.06 | 1.25E-04 | 2.37E-03 |
| distal tubule development (GO:0072017) | 12 | 7 | 1.38 | + | 5.06 | 1.25E-04 | 2.36E-03 |
| positive regulation of myoblast proliferation (GO:2000288) | 12 | 7 | 1.38 | + | 5.06 | 1.25E-04 | 2.36E-03 |
| kidney vasculature development (GO:0061440) | 24 | 14 | 2.77 | + | 5.06 | 4.47E-08 | 1.78E-06 |
| renal system vasculature development (GO:0061437) | 24 | 14 | 2.77 | + | 5.06 | 4.47E-08 | 1.77E-06 |
| metanephric nephron morphogenesis (GO:0072273) | 19 | 11 | 2.19 | + | 5.02 | 1.45E-06 | 4.30E-05 |
| olfactory nerve development (GO:0021553) | 7 | 4 | 0.81 | + | 4.96 | 4.62E-03 | 4.96E-02 |

| | | | | | | | |
|---|---|---|---|---|---|---|---|
| metanephric glomerulus vasculature development (GO:0072239) | 7 | 4 | 0.81 | + | 4.96 | 4.62E-03 | 4.96E-02 |
| negative regulation of mesenchymal cell proliferation (GO:0072201) | 7 | 4 | 0.81 | + | 4.96 | 4.62E-03 | 4.95E-02 |
| mesenchymal cell differentiation involved in kidney development (GO:0072161) | 7 | 4 | 0.81 | + | 4.96 | 4.62E-03 | 4.95E-02 |
| maintenance of protein location in extracellular region (GO:0071694) | 7 | 4 | 0.81 | + | 4.96 | 4.62E-03 | 4.95E-02 |
| mesenchymal cell differentiation involved in renal system development (GO:2001012) | 7 | 4 | 0.81 | + | 4.96 | 4.62E-03 | 4.94E-02 |
| positive regulation of gastrulation (GO:2000543) | 7 | 4 | 0.81 | + | 4.96 | 4.62E-03 | 4.94E-02 |
| adiponectin-activated signaling pathway (GO:0033211) | 7 | 4 | 0.81 | + | 4.96 | 4.62E-03 | 4.93E-02 |
| dorsal root ganglion development (GO:1990791) | 7 | 4 | 0.81 | + | 4.96 | 4.62E-03 | 4.93E-02 |
| optic cup morphogenesis involved in camera-type eye development (GO:0002072) | 7 | 4 | 0.81 | + | 4.96 | 4.62E-03 | 4.93E-02 |
| positive regulation of lamellipodium morphogenesis (GO:2000394) | 7 | 4 | 0.81 | + | 4.96 | 4.62E-03 | 4.92E-02 |
| dendrite self-avoidance (GO:0070593) | 14 | 8 | 1.61 | + | 4.96 | 4.86E-05 | 1.06E-03 |
| detection of muscle stretch (GO:0035995) | 7 | 4 | 0.81 | + | 4.96 | 4.62E-03 | 4.92E-02 |
| midbrain-hindbrain boundary development (GO:0030917) | 7 | 4 | 0.81 | + | 4.96 | 4.62E-03 | 4.92E-02 |
| regulation of aspartic-type peptidase activity (GO:1905245) | 7 | 4 | 0.81 | + | 4.96 | 4.62E-03 | 4.91E-02 |
| post-embryonic eye morphogenesis (GO:0048050) | 7 | 4 | 0.81 | + | 4.96 | 4.62E-03 | 4.91E-02 |
| limb joint morphogenesis (GO:0036022) | 7 | 4 | 0.81 | + | 4.96 | 4.62E-03 | 4.91E-02 |

| | | | | | | | |
|---|---|---|---|---|---|---|---|
| negative regulation of hepatocyte apoptotic process (GO:1903944) | 7 | 4 | 0.81 | + | 4.96 | 4.62E-03 | 4.90E-02 |
| hippo signaling (GO:0035329) | 21 | 12 | 2.42 | + | 4.96 | 5.77E-07 | 1.86E-05 |
| glycosylceramide biosynthetic process (GO:0046476) | 7 | 4 | 0.81 | + | 4.96 | 4.62E-03 | 4.90E-02 |
| lamellipodium morphogenesis (GO:0072673) | 7 | 4 | 0.81 | + | 4.96 | 4.62E-03 | 4.90E-02 |
| mammary gland formation (GO:0060592) | 7 | 4 | 0.81 | + | 4.96 | 4.62E-03 | 4.89E-02 |
| metanephric nephron development (GO:0072210) | 32 | 18 | 3.69 | + | 4.88 | 1.15E-09 | 5.66E-08 |
| estrous cycle (GO:0044849) | 16 | 9 | 1.84 | + | 4.88 | 1.89E-05 | 4.53E-04 |
| facial nerve structural organization (GO:0021612) | 9 | 5 | 1.04 | + | 4.82 | 1.71E-03 | 2.20E-02 |
| cranial suture morphogenesis (GO:0060363) | 9 | 5 | 1.04 | + | 4.82 | 1.71E-03 | 2.20E-02 |
| bone trabecula formation (GO:0060346) | 9 | 5 | 1.04 | + | 4.82 | 1.71E-03 | 2.19E-02 |
| metanephric collecting duct development (GO:0072205) | 9 | 5 | 1.04 | + | 4.82 | 1.71E-03 | 2.19E-02 |
| otic vesicle morphogenesis (GO:0071600) | 9 | 5 | 1.04 | + | 4.82 | 1.71E-03 | 2.19E-02 |
| positive regulation of mitochondrial depolarization (GO:0051901) | 9 | 5 | 1.04 | + | 4.82 | 1.71E-03 | 2.19E-02 |
| DNA topological change (GO:0006265) | 9 | 5 | 1.04 | + | 4.82 | 1.71E-03 | 2.19E-02 |
| positive regulation of Wnt signaling pathway, planar cell polarity pathway (GO:2000096) | 9 | 5 | 1.04 | + | 4.82 | 1.71E-03 | 2.18E-02 |
| embryonic foregut morphogenesis (GO:0048617) | 9 | 5 | 1.04 | + | 4.82 | 1.71E-03 | 2.18E-02 |

| | | | | | | | |
|---|---|---|---|---|---|---|---|
| positive regulation of hormone biosynthetic process (GO:0046886) | 9 | 5 | 1.04 | + | 4.82 | 1.71E-03 | 2.18E-02 |
| positive regulation of cardiac epithelial to mesenchymal transition (GO:0062043) | 9 | 5 | 1.04 | + | 4.82 | 1.71E-03 | 2.18E-02 |
| RNA secondary structure unwinding (GO:0010501) | 9 | 5 | 1.04 | + | 4.82 | 1.71E-03 | 2.18E-02 |
| negative regulation of neuroblast proliferation (GO:0007406) | 11 | 6 | 1.27 | + | 4.73 | 6.43E-04 | 9.62E-03 |
| cranial nerve structural organization (GO:0021604) | 11 | 6 | 1.27 | + | 4.73 | 6.43E-04 | 9.61E-03 |
| glomerular epithelial cell development (GO:0072310) | 11 | 6 | 1.27 | + | 4.73 | 6.43E-04 | 9.60E-03 |
| nephric duct morphogenesis (GO:0072178) | 11 | 6 | 1.27 | + | 4.73 | 6.43E-04 | 9.59E-03 |
| Leydig cell differentiation (GO:0033327) | 11 | 6 | 1.27 | + | 4.73 | 6.43E-04 | 9.58E-03 |
| collagen-activated tyrosine kinase receptor signaling pathway (GO:0038063) | 11 | 6 | 1.27 | + | 4.73 | 6.43E-04 | 9.57E-03 |
| CRD-mediated mRNA stabilization (GO:0070934) | 11 | 6 | 1.27 | + | 4.73 | 6.43E-04 | 9.56E-03 |
| thymocyte apoptotic process (GO:0070242) | 11 | 6 | 1.27 | + | 4.73 | 6.43E-04 | 9.55E-03 |
| embryonic camera-type eye formation (GO:0060900) | 11 | 6 | 1.27 | + | 4.73 | 6.43E-04 | 9.54E-03 |
| olfactory bulb interneuron differentiation (GO:0021889) | 11 | 6 | 1.27 | + | 4.73 | 6.43E-04 | 9.54E-03 |
| positive regulation of mesenchymal cell proliferation (GO:0002053) | 26 | 14 | 3 | + | 4.67 | 1.76E-07 | 6.26E-06 |
| metanephros morphogenesis (GO:0003338) | 26 | 14 | 3 | + | 4.67 | 1.76E-07 | 6.24E-06 |
| positive regulation of non-canonical Wnt signaling pathway (GO:2000052) | 15 | 8 | 1.73 | + | 4.63 | 9.36E-05 | 1.87E-03 |

| | | | | | | | |
|---|---|---|---|---|---|---|---|
| preganglionic parasympathetic fiber development (GO:0021783) | 17 | 9 | 1.96 | + | 4.59 | 3.61E-05 | 8.10E-04 |
| regulation of gastrulation (GO:0010470) | 17 | 9 | 1.96 | + | 4.59 | 3.61E-05 | 8.09E-04 |
| definitive hemopoiesis (GO:0060216) | 19 | 10 | 2.19 | + | 4.57 | 1.40E-05 | 3.46E-04 |
| neuronal stem cell population maintenance (GO:0097150) | 25 | 13 | 2.88 | + | 4.51 | 8.32E-07 | 2.63E-05 |
| mRNA splice site recognition (GO:0006376) | 31 | 16 | 3.57 | + | 4.48 | 5.06E-08 | 1.98E-06 |
| regulation of mesenchymal cell proliferation (GO:0010464) | 33 | 17 | 3.8 | + | 4.47 | 2.00E-08 | 8.27E-07 |
| bronchus development (GO:0060433) | 10 | 5 | 1.15 | + | 4.34 | 3.10E-03 | 3.54E-02 |
| negative regulation of osteoblast proliferation (GO:0033689) | 12 | 6 | 1.38 | + | 4.34 | 1.16E-03 | 1.59E-02 |
| integrin activation (GO:0033622) | 10 | 5 | 1.15 | + | 4.34 | 3.10E-03 | 3.54E-02 |
| specification of animal organ identity (GO:0010092) | 18 | 9 | 2.07 | + | 4.34 | 6.48E-05 | 1.36E-03 |
| metanephric renal vesicle morphogenesis (GO:0072283) | 10 | 5 | 1.15 | + | 4.34 | 3.10E-03 | 3.54E-02 |
| metanephric nephron epithelium development (GO:0072243) | 20 | 10 | 2.31 | + | 4.34 | 2.51E-05 | 5.85E-04 |
| regulation of peptidyl-lysine acetylation (GO:2000756) | 10 | 5 | 1.15 | + | 4.34 | 3.10E-03 | 3.53E-02 |
| metanephric nephron tubule development (GO:0072234) | 18 | 9 | 2.07 | + | 4.34 | 6.48E-05 | 1.35E-03 |
| cell differentiation involved in metanephros development (GO:0072202) | 16 | 8 | 1.84 | + | 4.34 | 1.68E-04 | 3.05E-03 |
| regulation of epithelial cell differentiation involved in kidney development (GO:2000696) | 16 | 8 | 1.84 | + | 4.34 | 1.68E-04 | 3.05E-03 |

| | | | | | | | |
|---|---|---|---|---|---|---|---|
| metanephric tubule development (GO:0072170) | 20 | 10 | 2.31 | + | 4.34 | 2.51E-05 | 5.84E-04 |
| regulation of glomerular mesangial cell proliferation (GO:0072124) | 10 | 5 | 1.15 | + | 4.34 | 3.10E-03 | 3.53E-02 |
| collagen biosynthetic process (GO:0032964) | 10 | 5 | 1.15 | + | 4.34 | 3.10E-03 | 3.53E-02 |
| loop of Henle development (GO:0072070) | 12 | 6 | 1.38 | + | 4.34 | 1.16E-03 | 1.59E-02 |
| positive regulation of endoplasmic reticulum stress-induced intrinsic apoptotic signaling pathway (GO:1902237) | 10 | 5 | 1.15 | + | 4.34 | 3.10E-03 | 3.53E-02 |
| cellular response to X-ray (GO:0071481) | 10 | 5 | 1.15 | + | 4.34 | 3.10E-03 | 3.52E-02 |
| lens morphogenesis in camera-type eye (GO:0002089) | 22 | 11 | 2.54 | + | 4.34 | 9.75E-06 | 2.49E-04 |
| regulation of cell proliferation involved in kidney development (GO:1901722) | 14 | 7 | 1.61 | + | 4.34 | 4.40E-04 | 6.93E-03 |
| RISC complex assembly (GO:0070922) | 10 | 5 | 1.15 | + | 4.34 | 3.10E-03 | 3.52E-02 |
| regulation of myoblast proliferation (GO:2000291) | 16 | 8 | 1.84 | + | 4.34 | 1.68E-04 | 3.04E-03 |
| regulation of Wnt signaling pathway, planar cell polarity pathway (GO:2000095) | 16 | 8 | 1.84 | + | 4.34 | 1.68E-04 | 3.04E-03 |
| neuron projection arborization (GO:0140058) | 18 | 9 | 2.07 | + | 4.34 | 6.48E-05 | 1.35E-03 |
| negative regulation of oligodendrocyte differentiation (GO:0048715) | 14 | 7 | 1.61 | + | 4.34 | 4.40E-04 | 6.92E-03 |
| positive regulation of kidney development (GO:0090184) | 10 | 5 | 1.15 | + | 4.34 | 3.10E-03 | 3.52E-02 |
| neuron fate determination (GO:0048664) | 10 | 5 | 1.15 | + | 4.34 | 3.10E-03 | 3.52E-02 |
| regulation of extracellular matrix assembly (GO:1901201) | 20 | 10 | 2.31 | + | 4.34 | 2.51E-05 | 5.83E-04 |

| | | | | | | | |
|---|---|---|---|---|---|---|---|
| protein heterotetramerization (GO:0051290) | 16 | 8 | 1.84 | + | 4.34 | 1.68E-04 | 3.04E-03 |
| regulation of response to reactive oxygen species (GO:1901031) | 10 | 5 | 1.15 | + | 4.34 | 3.10E-03 | 3.51E-02 |
| sequestering of extracellular ligand from receptor (GO:0035581) | 10 | 5 | 1.15 | + | 4.34 | 3.10E-03 | 3.51E-02 |
| regulation of cardiac epithelial to mesenchymal transition (GO:0062042) | 10 | 5 | 1.15 | + | 4.34 | 3.10E-03 | 3.51E-02 |
| bone trabecula morphogenesis (GO:0061430) | 12 | 6 | 1.38 | + | 4.34 | 1.16E-03 | 1.59E-02 |
| retina vasculature morphogenesis in camera-type eye (GO:0061299) | 10 | 5 | 1.15 | + | 4.34 | 3.10E-03 | 3.51E-02 |
| skeletal myofibril assembly (GO:0014866) | 10 | 5 | 1.15 | + | 4.34 | 3.10E-03 | 3.50E-02 |
| heart field specification (GO:0003128) | 12 | 6 | 1.38 | + | 4.34 | 1.16E-03 | 1.59E-02 |
| glomerulus development (GO:0032835) | 59 | 29 | 6.8 | + | 4.26 | 9.44E-13 | 6.70E-11 |
| cell differentiation involved in kidney development (GO:0061005) | 47 | 23 | 5.42 | + | 4.25 | 2.39E-10 | 1.30E-08 |
| positive regulation of epithelial to mesenchymal transition (GO:0010718) | 56 | 27 | 6.45 | + | 4.18 | 1.03E-11 | 6.49E-10 |
| metanephric epithelium development (GO:0072207) | 23 | 11 | 2.65 | + | 4.15 | 1.67E-05 | 4.07E-04 |
| collagen fibril organization (GO:0030199) | 61 | 29 | 7.03 | + | 4.12 | 2.76E-12 | 1.83E-10 |
| cellular response to fluid shear stress (GO:0071498) | 19 | 9 | 2.19 | + | 4.11 | 1.11E-04 | 2.14E-03 |
| cartilage condensation (GO:0001502) | 19 | 9 | 2.19 | + | 4.11 | 1.11E-04 | 2.14E-03 |
| parasympathetic nervous system development (GO:0048486) | 19 | 9 | 2.19 | + | 4.11 | 1.11E-04 | 2.14E-03 |

| | | | | | | | |
|---|---|---|---|---|---|---|---|
| regulation of striated muscle tissue development (GO:0016202) | 19 | 9 | 2.19 | + | 4.11 | 1.11E-04 | 2.13E-03 |
| smooth muscle cell differentiation (GO:0051145) | 36 | 17 | 4.15 | + | 4.1 | 1.05E-07 | 3.84E-06 |
| ureter development (GO:0072189) | 17 | 8 | 1.96 | + | 4.08 | 2.86E-04 | 4.77E-03 |
| anatomical structure arrangement (GO:0048532) | 17 | 8 | 1.96 | + | 4.08 | 2.86E-04 | 4.76E-03 |
| mammary gland epithelial cell differentiation (GO:0060644) | 17 | 8 | 1.96 | + | 4.08 | 2.86E-04 | 4.76E-03 |
| regulation of mammary gland epithelial cell proliferation (GO:0033599) | 15 | 7 | 1.73 | + | 4.05 | 7.43E-04 | 1.09E-02 |
| collagen-activated signaling pathway (GO:0038065) | 15 | 7 | 1.73 | + | 4.05 | 7.43E-04 | 1.09E-02 |
| negative regulation of gene expression via chromosomal CpG island methylation (GO:0044027) | 15 | 7 | 1.73 | + | 4.05 | 7.43E-04 | 1.09E-02 |
| neuroblast division (GO:0055057) | 15 | 7 | 1.73 | + | 4.05 | 7.43E-04 | 1.09E-02 |
| negative regulation of epidermis development (GO:0045683) | 13 | 6 | 1.5 | + | 4 | 1.95E-03 | 2.43E-02 |
| negative regulation of epidermal cell differentiation (GO:0045605) | 13 | 6 | 1.5 | + | 4 | 1.95E-03 | 2.43E-02 |
| positive regulation of protein acetylation (GO:1901985) | 13 | 6 | 1.5 | + | 4 | 1.95E-03 | 2.43E-02 |
| chromatin looping (GO:0140588) | 13 | 6 | 1.5 | + | 4 | 1.95E-03 | 2.42E-02 |
| metanephric mesenchyme development (GO:0072075) | 13 | 6 | 1.5 | + | 4 | 1.95E-03 | 2.42E-02 |
| convergent extension (GO:0060026) | 13 | 6 | 1.5 | + | 4 | 1.95E-03 | 2.42E-02 |
| peptidyl-lysine acetylation (GO:0018394) | 13 | 6 | 1.5 | + | 4 | 1.95E-03 | 2.42E-02 |

| | | | | | | | |
|---|---|---|---|---|---|---|---|
| keratinocyte development (GO:0003334) | 13 | 6 | 1.5 | + | 4 | 1.95E-03 | 2.42E-02 |
| motor neuron axon guidance (GO:0008045) | 26 | 12 | 3 | + | 4 | 1.09E-05 | 2.76E-04 |
| eyelid development in camera-type eye (GO:0061029) | 13 | 6 | 1.5 | + | 4 | 1.95E-03 | 2.41E-02 |
| olfactory lobe development (GO:0021988) | 37 | 17 | 4.26 | + | 3.99 | 1.73E-07 | 6.16E-06 |
| smooth muscle tissue development (GO:0048745) | 24 | 11 | 2.77 | + | 3.98 | 2.77E-05 | 6.37E-04 |
| olfactory bulb development (GO:0021772) | 35 | 16 | 4.03 | + | 3.97 | 4.34E-07 | 1.43E-05 |
| regulation of heterochromatin formation (GO:0031445) | 22 | 10 | 2.54 | + | 3.94 | 7.06E-05 | 1.46E-03 |
| cranial nerve morphogenesis (GO:0021602) | 31 | 14 | 3.57 | + | 3.92 | 2.76E-06 | 7.79E-05 |
| dorsal spinal cord development (GO:0021516) | 20 | 9 | 2.31 | + | 3.9 | 1.80E-04 | 3.22E-03 |
| retina vasculature development in camera-type eye (GO:0061298) | 20 | 9 | 2.31 | + | 3.9 | 1.80E-04 | 3.22E-03 |
| ephrin receptor signaling pathway (GO:0048013) | 49 | 22 | 5.65 | + | 3.9 | 4.59E-09 | 2.03E-07 |
| embryonic skeletal system morphogenesis (GO:0048704) | 94 | 42 | 10.83 | + | 3.88 | 6.05E-16 | 5.47E-14 |
| negative regulation of anoikis (GO:2000811) | 18 | 8 | 2.07 | + | 3.86 | 4.63E-04 | 7.24E-03 |
| ossification involved in bone maturation (GO:0043931) | 18 | 8 | 2.07 | + | 3.86 | 4.63E-04 | 7.23E-03 |
| lymph vessel morphogenesis (GO:0036303) | 18 | 8 | 2.07 | + | 3.86 | 4.63E-04 | 7.22E-03 |
| lymph vessel development (GO:0001945) | 27 | 12 | 3.11 | + | 3.86 | 1.75E-05 | 4.23E-04 |

| | | | | | | | |
|---|---|---|---|---|---|---|---|
| regulation of non-canonical Wnt signaling pathway (GO:2000050) | 27 | 12 | 3.11 | + | 3.86 | 1.75E-05 | 4.22E-04 |
| negative regulation of miRNA transcription (GO:1902894) | 27 | 12 | 3.11 | + | 3.86 | 1.75E-05 | 4.22E-04 |
| cell fate determination (GO:0001709) | 41 | 18 | 4.73 | + | 3.81 | 1.77E-07 | 6.27E-06 |
| regulation of neuron migration (GO:2001222) | 48 | 21 | 5.53 | + | 3.8 | 1.81E-08 | 7.57E-07 |
| branching involved in salivary gland morphogenesis (GO:0060445) | 16 | 7 | 1.84 | + | 3.8 | 1.19E-03 | 1.62E-02 |
| kidney mesenchyme development (GO:0072074) | 16 | 7 | 1.84 | + | 3.8 | 1.19E-03 | 1.62E-02 |
| modification of synaptic structure (GO:0099563) | 16 | 7 | 1.84 | + | 3.8 | 1.19E-03 | 1.62E-02 |
| aorta morphogenesis (GO:0035909) | 32 | 14 | 3.69 | + | 3.8 | 4.38E-06 | 1.20E-04 |
| positive regulation of smooth muscle cell migration (GO:0014911) | 39 | 17 | 4.5 | + | 3.78 | 4.41E-07 | 1.44E-05 |
| positive regulation of cardiac muscle cell proliferation (GO:0060045) | 23 | 10 | 2.65 | + | 3.77 | 1.12E-04 | 2.16E-03 |
| stem cell division (GO:0017145) | 23 | 10 | 2.65 | + | 3.77 | 1.12E-04 | 2.16E-03 |
| ionotropic glutamate receptor signaling pathway (GO:0035235) | 23 | 10 | 2.65 | + | 3.77 | 1.12E-04 | 2.15E-03 |
| atrial septum development (GO:0003283) | 23 | 10 | 2.65 | + | 3.77 | 1.12E-04 | 2.15E-03 |
| epithelial cell differentiation involved in kidney development (GO:0035850) | 37 | 16 | 4.26 | + | 3.75 | 1.10E-06 | 3.34E-05 |
| positive regulation of BMP signaling pathway (GO:0030513) | 37 | 16 | 4.26 | + | 3.75 | 1.10E-06 | 3.33E-05 |
| lymphangiogenesis (GO:0001946) | 14 | 6 | 1.61 | + | 3.72 | 3.07E-03 | 3.55E-02 |

| | | | | | | | |
|---|---|---|---|---|---|---|---|
| negative regulation of miRNA metabolic process (GO:2000629) | 28 | 12 | 3.23 | + | 3.72 | 2.75E-05 | 6.33E-04 |
| collecting duct development (GO:0072044) | 14 | 6 | 1.61 | + | 3.72 | 3.07E-03 | 3.55E-02 |
| otic vesicle development (GO:0071599) | 14 | 6 | 1.61 | + | 3.72 | 3.07E-03 | 3.54E-02 |
| left/right axis specification (GO:0070986) | 14 | 6 | 1.61 | + | 3.72 | 3.07E-03 | 3.54E-02 |
| dorsal/ventral axis specification (GO:0009950) | 14 | 6 | 1.61 | + | 3.72 | 3.07E-03 | 3.54E-02 |
| positive regulation of extracellular matrix assembly (GO:1901203) | 14 | 6 | 1.61 | + | 3.72 | 3.07E-03 | 3.53E-02 |
| regulation of thymocyte apoptotic process (GO:0070243) | 14 | 6 | 1.61 | + | 3.72 | 3.07E-03 | 3.53E-02 |
| positive regulation of vascular endothelial growth factor receptor signaling pathway (GO:0030949) | 14 | 6 | 1.61 | + | 3.72 | 3.07E-03 | 3.53E-02 |
| positive regulation of execution phase of apoptosis (GO:1900119) | 14 | 6 | 1.61 | + | 3.72 | 3.07E-03 | 3.53E-02 |
| cell communication by electrical coupling involved in cardiac conduction (GO:0086064) | 14 | 6 | 1.61 | + | 3.72 | 3.07E-03 | 3.52E-02 |
| bone maturation (GO:0070977) | 21 | 9 | 2.42 | + | 3.72 | 2.84E-04 | 4.74E-03 |
| cell aggregation (GO:0098743) | 21 | 9 | 2.42 | + | 3.72 | 2.84E-04 | 4.74E-03 |
| ovulation cycle process (GO:0022602) | 47 | 20 | 5.42 | + | 3.69 | 6.99E-08 | 2.67E-06 |
| regulation of chromatin organization (GO:1902275) | 33 | 14 | 3.8 | + | 3.68 | 6.80E-06 | 1.80E-04 |
| aorta development (GO:0035904) | 59 | 25 | 6.8 | + | 3.68 | 1.83E-09 | 8.55E-08 |
| regulation of vascular associated smooth muscle cell migration (GO:1904752) | 26 | 11 | 3 | + | 3.67 | 6.88E-05 | 1.43E-03 |

| | | | | | | | |
|---|---|---|---|---|---|---|---|
| secondary palate development (GO:0062009) | 26 | 11 | 3 | + | 3.67 | 6.88E-05 | 1.43E-03 |
| DNA damage response, signal transduction by p53 class mediator resulting in cell cycle arrest (GO:0006977) | 19 | 8 | 2.19 | + | 3.65 | 7.19E-04 | 1.06E-02 |
| paraxial mesoderm development (GO:0048339) | 19 | 8 | 2.19 | + | 3.65 | 7.19E-04 | 1.06E-02 |
| outflow tract septum morphogenesis (GO:0003148) | 24 | 10 | 2.77 | + | 3.62 | 1.72E-04 | 3.10E-03 |
| ovulation cycle (GO:0042698) | 70 | 29 | 8.07 | + | 3.59 | 1.80E-10 | 9.93E-09 |
| regulation of extracellular matrix organization (GO:1903053) | 63 | 26 | 7.26 | + | 3.58 | 1.73E-09 | 8.16E-08 |
| regulation of sister chromatid cohesion (GO:0007063) | 17 | 7 | 1.96 | + | 3.57 | 1.82E-03 | 2.30E-02 |
| embryonic brain development (GO:1990403) | 17 | 7 | 1.96 | + | 3.57 | 1.82E-03 | 2.30E-02 |
| white fat cell differentiation (GO:0050872) | 17 | 7 | 1.96 | + | 3.57 | 1.82E-03 | 2.29E-02 |
| cell migration involved in heart development (GO:0060973) | 17 | 7 | 1.96 | + | 3.57 | 1.82E-03 | 2.29E-02 |
| podocyte differentiation (GO:0072112) | 22 | 9 | 2.54 | + | 3.55 | 4.31E-04 | 6.85E-03 |
| renal filtration cell differentiation (GO:0061318) | 22 | 9 | 2.54 | + | 3.55 | 4.31E-04 | 6.84E-03 |
| cardiac septum morphogenesis (GO:0060411) | 71 | 29 | 8.18 | + | 3.54 | 2.71E-10 | 1.46E-08 |
| epithelial tube branching involved in lung morphogenesis (GO:0060441) | 27 | 11 | 3.11 | + | 3.53 | 1.04E-04 | 2.06E-03 |
| chondrocyte development (GO:0002063) | 27 | 11 | 3.11 | + | 3.53 | 1.04E-04 | 2.05E-03 |
| axis elongation (GO:0003401) | 27 | 11 | 3.11 | + | 3.53 | 1.04E-04 | 2.05E-03 |

| | | | | | | | |
|---|---|---|---|---|---|---|---|
| mesenchymal cell proliferation (GO:0010463) | 27 | 11 | 3.11 | + | 3.53 | 1.04E-04 | 2.05E-03 |
| atrioventricular valve morphogenesis (GO:0003181) | 27 | 11 | 3.11 | + | 3.53 | 1.04E-04 | 2.05E-03 |
| positive regulation of synapse assembly (GO:0051965) | 64 | 26 | 7.38 | + | 3.52 | 2.59E-09 | 1.20E-07 |
| embryonic skeletal joint development (GO:0072498) | 15 | 6 | 1.73 | + | 3.47 | 4.63E-03 | 4.90E-02 |
| atrial septum morphogenesis (GO:0060413) | 15 | 6 | 1.73 | + | 3.47 | 4.63E-03 | 4.89E-02 |
| regulation of cellular response to vascular endothelial growth factor stimulus (GO:1902547) | 25 | 10 | 2.88 | + | 3.47 | 2.58E-04 | 4.42E-03 |
| epithelial cell fate commitment (GO:0072148) | 15 | 6 | 1.73 | + | 3.47 | 4.63E-03 | 4.89E-02 |
| regulation of integrin-mediated signaling pathway (GO:2001044) | 20 | 8 | 2.31 | + | 3.47 | 1.08E-03 | 1.49E-02 |
| branching involved in ureteric bud morphogenesis (GO:0001658) | 45 | 18 | 5.19 | + | 3.47 | 9.54E-07 | 2.93E-05 |
| negative regulation of neural precursor cell proliferation (GO:2000178) | 25 | 10 | 2.88 | + | 3.47 | 2.58E-04 | 4.41E-03 |
| animal organ maturation (GO:0048799) | 25 | 10 | 2.88 | + | 3.47 | 2.58E-04 | 4.41E-03 |
| embryonic cranial skeleton morphogenesis (GO:0048701) | 45 | 18 | 5.19 | + | 3.47 | 9.54E-07 | 2.93E-05 |
| positive regulation of chondrocyte differentiation (GO:0032332) | 20 | 8 | 2.31 | + | 3.47 | 1.08E-03 | 1.49E-02 |
| negative regulation of axon regeneration (GO:0048681) | 15 | 6 | 1.73 | + | 3.47 | 4.63E-03 | 4.89E-02 |
| regulation of steroid hormone biosynthetic process (GO:0090030) | 15 | 6 | 1.73 | + | 3.47 | 4.63E-03 | 4.88E-02 |
| response to muscle stretch (GO:0035994) | 25 | 10 | 2.88 | + | 3.47 | 2.58E-04 | 4.40E-03 |

| | | | | | | | |
|---|---|---|---|---|---|---|---|
| chondrocyte proliferation (GO:0035988) | 15 | 6 | 1.73 | + | 3.47 | 4.63E-03 | 4.88E-02 |
| negative regulation of DNA damage response, signal transduction by p53 class mediator (GO:0043518) | 15 | 6 | 1.73 | + | 3.47 | 4.63E-03 | 4.88E-02 |
| positive regulation of vascular associated smooth muscle cell migration (GO:1904754) | 15 | 6 | 1.73 | + | 3.47 | 4.63E-03 | 4.87E-02 |
| embryonic eye morphogenesis (GO:0048048) | 35 | 14 | 4.03 | + | 3.47 | 1.54E-05 | 3.76E-04 |
| regulation of extracellular matrix disassembly (GO:0010715) | 15 | 6 | 1.73 | + | 3.47 | 4.63E-03 | 4.87E-02 |
| extracellular matrix assembly (GO:0085029) | 30 | 12 | 3.46 | + | 3.47 | 6.26E-05 | 1.32E-03 |
| forebrain neuron development (GO:0021884) | 25 | 10 | 2.88 | + | 3.47 | 2.58E-04 | 4.40E-03 |
| pyramidal neuron differentiation (GO:0021859) | 15 | 6 | 1.73 | + | 3.47 | 4.63E-03 | 4.87E-02 |
| pulmonary valve morphogenesis (GO:0003184) | 20 | 8 | 2.31 | + | 3.47 | 1.08E-03 | 1.48E-02 |
| atrioventricular valve development (GO:0003171) | 30 | 12 | 3.46 | + | 3.47 | 6.26E-05 | 1.31E-03 |
| embryonic skeletal system development (GO:0048706) | 128 | 51 | 14.75 | + | 3.46 | 1.74E-16 | 1.64E-14 |
| axon guidance (GO:0007411) | 224 | 89 | 25.82 | + | 3.45 | 1.22E-27 | 1.92E-25 |
| neuron projection guidance (GO:0097485) | 225 | 89 | 25.93 | + | 3.43 | 1.80E-27 | 2.74E-25 |
| spinal cord development (GO:0021510) | 99 | 39 | 11.41 | + | 3.42 | 9.34E-13 | 6.66E-11 |
| gastrulation with mouth forming second (GO:0001702) | 28 | 11 | 3.23 | + | 3.41 | 1.54E-04 | 2.82E-03 |
| nephron development (GO:0072006) | 140 | 55 | 16.14 | + | 3.41 | 2.43E-17 | 2.45E-15 |

| | | | | | | | |
|---|---|---|---|---|---|---|---|
| glomerular epithelial cell differentiation (GO:0072311) | 23 | 9 | 2.65 | + | 3.4 | 6.36E-04 | 9.57E-03 |
| negative chemotaxis (GO:0050919) | 46 | 18 | 5.3 | + | 3.4 | 1.40E-06 | 4.14E-05 |
| regulation of vascular endothelial growth factor signaling pathway (GO:1900746) | 23 | 9 | 2.65 | + | 3.4 | 6.36E-04 | 9.56E-03 |
| regulation of nuclear-transcribed mRNA catabolic process, deadenylation-dependent decay (GO:1900151) | 23 | 9 | 2.65 | + | 3.4 | 6.36E-04 | 9.55E-03 |
| negative regulation of smooth muscle cell migration (GO:0014912) | 23 | 9 | 2.65 | + | 3.4 | 6.36E-04 | 9.54E-03 |
| cranial nerve development (GO:0021545) | 64 | 25 | 7.38 | + | 3.39 | 1.36E-08 | 5.76E-07 |
| negative regulation of cell-substrate adhesion (GO:0010812) | 59 | 23 | 6.8 | + | 3.38 | 5.36E-08 | 2.08E-06 |
| negative regulation of cell-matrix adhesion (GO:0001953) | 36 | 14 | 4.15 | + | 3.37 | 2.25E-05 | 5.36E-04 |
| spinal cord patterning (GO:0021511) | 18 | 7 | 2.07 | + | 3.37 | 2.69E-03 | 3.17E-02 |
| angiogenesis involved in wound healing (GO:0060055) | 18 | 7 | 2.07 | + | 3.37 | 2.69E-03 | 3.16E-02 |
| genomic imprinting (GO:0071514) | 18 | 7 | 2.07 | + | 3.37 | 2.69E-03 | 3.16E-02 |
| regulation of smooth muscle cell differentiation (GO:0051150) | 36 | 14 | 4.15 | + | 3.37 | 2.25E-05 | 5.35E-04 |
| vascular associated smooth muscle cell differentiation (GO:0035886) | 18 | 7 | 2.07 | + | 3.37 | 2.69E-03 | 3.16E-02 |
| endochondral ossification (GO:0001958) | 31 | 12 | 3.57 | + | 3.36 | 9.15E-05 | 1.84E-03 |
| replacement ossification (GO:0036075) | 31 | 12 | 3.57 | + | 3.36 | 9.15E-05 | 1.84E-03 |
| mammary gland duct morphogenesis (GO:0060603) | 31 | 12 | 3.57 | + | 3.36 | 9.15E-05 | 1.83E-03 |

| | | | | | | | |
|---|---|---|---|---|---|---|---|
| heart valve morphogenesis (GO:0003179) | 62 | 24 | 7.15 | + | 3.36 | 3.25E-08 | 1.31E-06 |
| cellular response to transforming growth factor beta stimulus (GO:0071560) | 153 | 59 | 17.63 | + | 3.35 | 4.84E-18 | 5.09E-16 |
| endoderm development (GO:0007492) | 83 | 32 | 9.57 | + | 3.35 | 1.96E-10 | 1.07E-08 |
| negative regulation of stem cell differentiation (GO:2000737) | 26 | 10 | 3 | + | 3.34 | 3.76E-04 | 6.03E-03 |
| glomerular epithelium development (GO:0072010) | 26 | 10 | 3 | + | 3.34 | 3.76E-04 | 6.03E-03 |
| positive regulation of focal adhesion assembly (GO:0051894) | 26 | 10 | 3 | + | 3.34 | 3.76E-04 | 6.02E-03 |
| synaptic membrane adhesion (GO:0099560) | 26 | 10 | 3 | + | 3.34 | 3.76E-04 | 6.01E-03 |
| embryonic camera-type eye morphogenesis (GO:0048596) | 26 | 10 | 3 | + | 3.34 | 3.76E-04 | 6.01E-03 |
| coronary vasculature development (GO:0060976) | 47 | 18 | 5.42 | + | 3.32 | 2.02E-06 | 5.85E-05 |
| cardiac muscle tissue growth (GO:0055017) | 34 | 13 | 3.92 | + | 3.32 | 5.45E-05 | 1.18E-03 |
| regulation of alternative mRNA splicing, via spliceosome (GO:0000381) | 55 | 21 | 6.34 | + | 3.31 | 3.07E-07 | 1.04E-05 |
| response to transforming growth factor beta (GO:0071559) | 160 | 61 | 18.44 | + | 3.31 | 2.56E-18 | 2.76E-16 |
| ventricular septum morphogenesis (GO:0060412) | 42 | 16 | 4.84 | + | 3.31 | 8.08E-06 | 2.09E-04 |
| Sertoli cell differentiation (GO:0060008) | 21 | 8 | 2.42 | + | 3.31 | 1.57E-03 | 2.03E-02 |
| vascular endothelial growth factor signaling pathway (GO:0038084) | 21 | 8 | 2.42 | + | 3.31 | 1.57E-03 | 2.03E-02 |
| cell migration involved in sprouting angiogenesis (GO:0002042) | 21 | 8 | 2.42 | + | 3.31 | 1.57E-03 | 2.03E-02 |

| | | | | | | | | |
|---|---|---|---|---|---|---|---|---|
| cellular response to platelet-derived growth factor stimulus (GO:0036120) | 21 | 8 | 2.42 | + | | 3.31 | 1.57E-03 | 2.03E-02 |
| retinal ganglion cell axon guidance (GO:0031290) | 21 | 8 | 2.42 | + | | 3.31 | 1.57E-03 | 2.03E-02 |
| salivary gland morphogenesis (GO:0007435) | 29 | 11 | 3.34 | + | | 3.29 | 2.22E-04 | 3.85E-03 |
| mammary gland morphogenesis (GO:0060443) | 45 | 17 | 5.19 | + | | 3.28 | 4.83E-06 | 1.31E-04 |
| renal tubule development (GO:0061326) | 90 | 34 | 10.37 | + | | 3.28 | 1.03E-10 | 5.82E-09 |
| nephron tubule development (GO:0072080) | 85 | 32 | 9.8 | + | | 3.27 | 4.02E-10 | 2.12E-08 |
| metanephros development (GO:0001656) | 85 | 32 | 9.8 | + | | 3.27 | 4.02E-10 | 2.11E-08 |
| face morphogenesis (GO:0060325) | 32 | 12 | 3.69 | + | | 3.25 | 1.31E-04 | 2.45E-03 |
| nuclear migration (GO:0007097) | 24 | 9 | 2.77 | + | | 3.25 | 9.14E-04 | 1.30E-02 |
| nephron morphogenesis (GO:0072028) | 72 | 27 | 8.3 | + | | 3.25 | 1.02E-08 | 4.39E-07 |
| protein localization to chromosome, centromeric region (GO:0071459) | 24 | 9 | 2.77 | + | | 3.25 | 9.14E-04 | 1.30E-02 |
| hematopoietic stem cell proliferation (GO:0071425) | 24 | 9 | 2.77 | + | | 3.25 | 9.14E-04 | 1.30E-02 |
| regulation of brown fat cell differentiation (GO:0090335) | 24 | 9 | 2.77 | + | | 3.25 | 9.14E-04 | 1.30E-02 |
| negative regulation of chondrocyte differentiation (GO:0032331) | 24 | 9 | 2.77 | + | | 3.25 | 9.14E-04 | 1.30E-02 |
| regulation of muscle organ development (GO:0048634) | 24 | 9 | 2.77 | + | | 3.25 | 9.14E-04 | 1.29E-02 |
| alternative mRNA splicing, via spliceosome (GO:0000380) | 24 | 9 | 2.77 | + | | 3.25 | 9.14E-04 | 1.29E-02 |

| | | | | | | | |
|---|---|---|---|---|---|---|---|
| renal tubule morphogenesis (GO:0061333) | 72 | 27 | 8.3 | + | 3.25 | 1.02E-08 | 4.38E-07 |
| positive regulation of cartilage development (GO:0061036) | 32 | 12 | 3.69 | + | 3.25 | 1.31E-04 | 2.45E-03 |
| forebrain neuron differentiation (GO:0021879) | 40 | 15 | 4.61 | + | 3.25 | 1.93E-05 | 4.61E-04 |
| cardiac septum development (GO:0003279) | 107 | 40 | 12.33 | + | 3.24 | 3.49E-12 | 2.28E-10 |
| roof of mouth development (GO:0060021) | 91 | 34 | 10.49 | + | 3.24 | 1.46E-10 | 8.15E-09 |
| nephron epithelium development (GO:0072009) | 110 | 41 | 12.68 | + | 3.23 | 2.11E-12 | 1.43E-10 |
| cell-substrate junction organization (GO:0150115) | 43 | 16 | 4.96 | + | 3.23 | 1.15E-05 | 2.89E-04 |
| regulation of kidney development (GO:0090183) | 35 | 13 | 4.03 | + | 3.22 | 7.76E-05 | 1.58E-03 |
| platelet-derived growth factor receptor signaling pathway (GO:0048008) | 35 | 13 | 4.03 | + | 3.22 | 7.76E-05 | 1.57E-03 |
| kidney morphogenesis (GO:0060993) | 89 | 33 | 10.26 | + | 3.22 | 3.43E-10 | 1.81E-08 |
| telencephalon glial cell migration (GO:0022030) | 27 | 10 | 3.11 | + | 3.21 | 5.35E-04 | 8.23E-03 |
| positive regulation of cardiac muscle tissue growth (GO:0055023) | 27 | 10 | 3.11 | + | 3.21 | 5.35E-04 | 8.22E-03 |
| cerebral cortex radial glia-guided migration (GO:0021801) | 27 | 10 | 3.11 | + | 3.21 | 5.35E-04 | 8.22E-03 |
| ventricular septum development (GO:0003281) | 73 | 27 | 8.41 | + | 3.21 | 1.44E-08 | 6.08E-07 |
| glutamate receptor signaling pathway (GO:0007215) | 46 | 17 | 5.3 | + | 3.21 | 6.85E-06 | 1.81E-04 |
| regulation of muscle tissue development (GO:1901861) | 46 | 17 | 5.3 | + | 3.21 | 6.85E-06 | 1.80E-04 |

| | | | | | | | |
|---|---|---|---|---|---|---|---|
| endodermal cell differentiation (GO:0035987) | 46 | 17 | 5.3 | + | 3.21 | 6.85E-06 | 1.80E-04 |
| neuron recognition (GO:0008038) | 46 | 17 | 5.3 | + | 3.21 | 6.85E-06 | 1.80E-04 |
| regulation of smooth muscle cell migration (GO:0014910) | 65 | 24 | 7.49 | + | 3.2 | 9.42E-08 | 3.51E-06 |
| trachea development (GO:0060438) | 19 | 7 | 2.19 | + | 3.2 | 3.84E-03 | 4.24E-02 |
| head morphogenesis (GO:0060323) | 38 | 14 | 4.38 | + | 3.2 | 4.59E-05 | 1.00E-03 |
| mesonephric tubule morphogenesis (GO:0072171) | 57 | 21 | 6.57 | + | 3.2 | 6.18E-07 | 1.98E-05 |
| Wnt signaling pathway, planar cell polarity pathway (GO:0060071) | 38 | 14 | 4.38 | + | 3.2 | 4.59E-05 | 1.00E-03 |
| notochord development (GO:0030903) | 19 | 7 | 2.19 | + | 3.2 | 3.84E-03 | 4.24E-02 |
| fibroblast migration (GO:0010761) | 19 | 7 | 2.19 | + | 3.2 | 3.84E-03 | 4.23E-02 |
| cell communication by electrical coupling (GO:0010644) | 19 | 7 | 2.19 | + | 3.2 | 3.84E-03 | 4.23E-02 |
| cardiac right ventricle morphogenesis (GO:0003215) | 19 | 7 | 2.19 | + | 3.2 | 3.84E-03 | 4.23E-02 |
| negative regulation of transcription elongation by RNA polymerase II (GO:0034244) | 19 | 7 | 2.19 | + | 3.2 | 3.84E-03 | 4.22E-02 |
| negative regulation of osteoblast differentiation (GO:0045668) | 49 | 18 | 5.65 | + | 3.19 | 4.08E-06 | 1.13E-04 |
| positive regulation of extracellular matrix organization (GO:1903055) | 30 | 11 | 3.46 | + | 3.18 | 3.14E-04 | 5.20E-03 |
| negative regulation of cell junction assembly (GO:1901889) | 30 | 11 | 3.46 | + | 3.18 | 3.14E-04 | 5.19E-03 |
| T cell apoptotic process (GO:0070231) | 30 | 11 | 3.46 | + | 3.18 | 3.14E-04 | 5.19E-03 |

| | | | | | | | |
|---|---|---|---|---|---|---|---|
| neuron projection regeneration (GO:0031102) | 30 | 11 | 3.46 | + | 3.18 | 3.14E-04 | 5.18E-03 |
| negative regulation of cartilage development (GO:0061037) | 30 | 11 | 3.46 | + | 3.18 | 3.14E-04 | 5.18E-03 |
| cell-substrate junction assembly (GO:0007044) | 41 | 15 | 4.73 | + | 3.17 | 2.72E-05 | 6.28E-04 |
| forebrain generation of neurons (GO:0021872) | 52 | 19 | 5.99 | + | 3.17 | 2.43E-06 | 6.95E-05 |
| regulation of focal adhesion assembly (GO:0051893) | 63 | 23 | 7.26 | + | 3.17 | 2.21E-07 | 7.67E-06 |
| regulation of cell-substrate junction assembly (GO:0090109) | 63 | 23 | 7.26 | + | 3.17 | 2.21E-07 | 7.65E-06 |
| regulation of attachment of spindle microtubules to kinetochore (GO:0051988) | 22 | 8 | 2.54 | + | 3.15 | 2.21E-03 | 2.67E-02 |
| negative regulation of mRNA processing (GO:0050686) | 22 | 8 | 2.54 | + | 3.15 | 2.21E-03 | 2.67E-02 |
| peptidyl-tyrosine dephosphorylation (GO:0035335) | 33 | 12 | 3.8 | + | 3.15 | 1.85E-04 | 3.28E-03 |
| embryonic digit morphogenesis (GO:0042733) | 58 | 21 | 6.68 | + | 3.14 | 8.63E-07 | 2.70E-05 |
| kidney epithelium development (GO:0072073) | 141 | 51 | 16.25 | + | 3.14 | 1.82E-14 | 1.46E-12 |
| heart growth (GO:0060419) | 36 | 13 | 4.15 | + | 3.13 | 1.09E-04 | 2.12E-03 |
| endocrine pancreas development (GO:0031018) | 36 | 13 | 4.15 | + | 3.13 | 1.09E-04 | 2.12E-03 |
| sprouting angiogenesis (GO:0002040) | 61 | 22 | 7.03 | + | 3.13 | 5.15E-07 | 1.68E-05 |
| negative regulation of myoblast differentiation (GO:0045662) | 25 | 9 | 2.88 | + | 3.12 | 1.28E-03 | 1.73E-02 |
| SMAD protein signal transduction (GO:0060395) | 25 | 9 | 2.88 | + | 3.12 | 1.28E-03 | 1.73E-02 |

| | | | | | | | |
|---|---|---|---|---|---|---|---|
| positive regulation of collagen biosynthetic process (GO:0032967) | 25 | 9 | 2.88 | + | 3.12 | 1.28E-03 | 1.72E-02 |
| cardiac muscle cell proliferation (GO:0060038) | 25 | 9 | 2.88 | + | 3.12 | 1.28E-03 | 1.72E-02 |
| cochlea development (GO:0090102) | 50 | 18 | 5.76 | + | 3.12 | 5.69E-06 | 1.52E-04 |
| regulation of miRNA transcription (GO:1902893) | 75 | 27 | 8.64 | + | 3.12 | 2.82E-08 | 1.15E-06 |
| mesonephric tubule development (GO:0072164) | 89 | 32 | 10.26 | + | 3.12 | 1.56E-09 | 7.42E-08 |
| mesonephric epithelium development (GO:0072163) | 89 | 32 | 10.26 | + | 3.12 | 1.56E-09 | 7.39E-08 |
| regulation of epithelial to mesenchymal transition (GO:0010717) | 103 | 37 | 11.87 | + | 3.12 | 8.68E-11 | 4.95E-09 |
| aortic valve morphogenesis (GO:0003180) | 39 | 14 | 4.5 | + | 3.11 | 6.41E-05 | 1.34E-03 |
| face development (GO:0060324) | 53 | 19 | 6.11 | + | 3.11 | 3.38E-06 | 9.45E-05 |
| non-canonical Wnt signaling pathway (GO:0035567) | 53 | 19 | 6.11 | + | 3.11 | 3.38E-06 | 9.43E-05 |
| endoderm formation (GO:0001706) | 56 | 20 | 6.45 | + | 3.1 | 2.01E-06 | 5.84E-05 |
| somatic stem cell population maintenance (GO:0035019) | 56 | 20 | 6.45 | + | 3.1 | 2.01E-06 | 5.83E-05 |
| ureteric bud morphogenesis (GO:0060675) | 56 | 20 | 6.45 | + | 3.1 | 2.01E-06 | 5.82E-05 |
| nephron epithelium morphogenesis (GO:0072088) | 70 | 25 | 8.07 | + | 3.1 | 1.09E-07 | 4.00E-06 |
| chondrocyte differentiation (GO:0002062) | 84 | 30 | 9.68 | + | 3.1 | 6.02E-09 | 2.63E-07 |
| skeletal system morphogenesis (GO:0048705) | 227 | 81 | 26.16 | + | 3.1 | 1.02E-21 | 1.23E-19 |

| | | | | | | | |
|---|---|---|---|---|---|---|---|
| regulation of osteoblast differentiation (GO:0045667) | 129 | 46 | 14.87 | + | 3.09 | 6.25E-13 | 4.63E-11 |
| endochondral bone morphogenesis (GO:0060350) | 59 | 21 | 6.8 | + | 3.09 | 1.20E-06 | 3.60E-05 |
| neural tube closure (GO:0001843) | 90 | 32 | 10.37 | + | 3.08 | 2.15E-09 | 9.99E-08 |
| positive regulation of heart growth (GO:0060421) | 31 | 11 | 3.57 | + | 3.08 | 4.36E-04 | 6.92E-03 |
| neural tube patterning (GO:0021532) | 31 | 11 | 3.57 | + | 3.08 | 4.36E-04 | 6.91E-03 |
| nucleus localization (GO:0051647) | 31 | 11 | 3.57 | + | 3.08 | 4.36E-04 | 6.90E-03 |
| positive regulation of filopodium assembly (GO:0051491) | 31 | 11 | 3.57 | + | 3.08 | 4.36E-04 | 6.89E-03 |
| regulation of neuron projection regeneration (GO:0070570) | 31 | 11 | 3.57 | + | 3.08 | 4.36E-04 | 6.89E-03 |
| mesonephros development (GO:0001823) | 93 | 33 | 10.72 | + | 3.08 | 1.29E-09 | 6.26E-08 |
| nephron tubule morphogenesis (GO:0072078) | 68 | 24 | 7.84 | + | 3.06 | 2.52E-07 | 8.57E-06 |
| artery morphogenesis (GO:0048844) | 68 | 24 | 7.84 | + | 3.06 | 2.52E-07 | 8.55E-06 |
| regulation of BMP signaling pathway (GO:0030510) | 105 | 37 | 12.1 | + | 3.06 | 1.65E-10 | 9.16E-09 |
| ureteric bud development (GO:0001657) | 88 | 31 | 10.14 | + | 3.06 | 4.95E-09 | 2.18E-07 |
| morphogenesis of an epithelial sheet (GO:0002011) | 54 | 19 | 6.22 | + | 3.05 | 4.65E-06 | 1.28E-04 |
| protein sumoylation (GO:0016925) | 54 | 19 | 6.22 | + | 3.05 | 4.65E-06 | 1.27E-04 |
| tube closure (GO:0060606) | 91 | 32 | 10.49 | + | 3.05 | 2.96E-09 | 1.35E-07 |

| | | | | | | | |
|---|---|---|---|---|---|---|---|
| regulation of collagen metabolic process (GO:0010712) | 37 | 13 | 4.26 | + | 3.05 | 1.50E-04 | 2.77E-03 |
| cardiac atrium development (GO:0003230) | 37 | 13 | 4.26 | + | 3.05 | 1.50E-04 | 2.77E-03 |
| mesenchymal cell migration (GO:0090497) | 57 | 20 | 6.57 | + | 3.04 | 2.76E-06 | 7.79E-05 |
| artery development (GO:0060840) | 100 | 35 | 11.53 | + | 3.04 | 6.31E-10 | 3.22E-08 |
| glandular epithelial cell differentiation (GO:0002067) | 63 | 22 | 7.26 | + | 3.03 | 9.75E-07 | 2.98E-05 |
| body morphogenesis (GO:0010171) | 46 | 16 | 5.3 | + | 3.02 | 3.07E-05 | 6.99E-04 |
| positive regulation of osteoblast differentiation (GO:0045669) | 69 | 24 | 7.95 | + | 3.02 | 3.45E-07 | 1.15E-05 |
| axon extension (GO:0048675) | 46 | 16 | 5.3 | + | 3.02 | 3.07E-05 | 6.98E-04 |
| autonomic nervous system development (GO:0048483) | 46 | 16 | 5.3 | + | 3.02 | 3.07E-05 | 6.97E-04 |
| negative regulation of vascular associated smooth muscle cell proliferation (GO:1904706) | 23 | 8 | 2.65 | + | 3.02 | 3.06E-03 | 3.54E-02 |
| response to platelet-derived growth factor (GO:0036119) | 23 | 8 | 2.65 | + | 3.02 | 3.06E-03 | 3.53E-02 |
| hematopoietic stem cell homeostasis (GO:0061484) | 23 | 8 | 2.65 | + | 3.02 | 3.06E-03 | 3.53E-02 |
| vasculogenesis (GO:0001570) | 72 | 25 | 8.3 | + | 3.01 | 2.05E-07 | 7.18E-06 |
| negative regulation of glial cell differentiation (GO:0045686) | 26 | 9 | 3 | + | 3 | 1.76E-03 | 2.25E-02 |
| regulation of chondrocyte differentiation (GO:0032330) | 52 | 18 | 5.99 | + | 3 | 1.07E-05 | 2.73E-04 |
| positive regulation of collagen metabolic process (GO:0010714) | 26 | 9 | 3 | + | 3 | 1.76E-03 | 2.24E-02 |

| | | | | | | | |
|---|---|---|---|---|---|---|---|
| adherens junction organization (GO:0034332) | 52 | 18 | 5.99 | + | 3 | 1.07E-05 | 2.72E-04 |
| intestinal epithelial cell differentiation (GO:0060575) | 26 | 9 | 3 | + | 3 | 1.76E-03 | 2.24E-02 |
| branching morphogenesis of an epithelial tube (GO:0048754) | 136 | 47 | 15.68 | + | 3 | 1.31E-12 | 9.15E-11 |
| positive regulation of blood vessel endothelial cell migration (GO:0043536) | 55 | 19 | 6.34 | + | 3 | 6.35E-06 | 1.68E-04 |
| basement membrane organization (GO:0071711) | 29 | 10 | 3.34 | + | 2.99 | 1.02E-03 | 1.43E-02 |
| glial cell proliferation (GO:0014009) | 29 | 10 | 3.34 | + | 2.99 | 1.02E-03 | 1.43E-02 |
| regulation of axon regeneration (GO:0048679) | 29 | 10 | 3.34 | + | 2.99 | 1.02E-03 | 1.42E-02 |
| regulation of post-transcriptional gene silencing by regulatory ncRNA (GO:1900368) | 29 | 10 | 3.34 | + | 2.99 | 1.02E-03 | 1.42E-02 |
| focal adhesion assembly (GO:0048041) | 29 | 10 | 3.34 | + | 2.99 | 1.02E-03 | 1.42E-02 |
| central nervous system projection neuron axonogenesis (GO:0021952) | 29 | 10 | 3.34 | + | 2.99 | 1.02E-03 | 1.42E-02 |
| salivary gland development (GO:0007431) | 32 | 11 | 3.69 | + | 2.98 | 5.96E-04 | 9.11E-03 |
| transforming growth factor beta receptor signaling pathway (GO:0007179) | 96 | 33 | 11.06 | + | 2.98 | 3.27E-09 | 1.48E-07 |
| regulation of collagen biosynthetic process (GO:0032965) | 32 | 11 | 3.69 | + | 2.98 | 5.96E-04 | 9.10E-03 |
| primary neural tube formation (GO:0014020) | 96 | 33 | 11.06 | + | 2.98 | 3.27E-09 | 1.48E-07 |
| dendrite morphogenesis (GO:0048813) | 64 | 22 | 7.38 | + | 2.98 | 1.32E-06 | 3.94E-05 |
| epigenetic programming of gene expression (GO:0043045) | 32 | 11 | 3.69 | + | 2.98 | 5.96E-04 | 9.09E-03 |

| | | | | | | | |
|---|---|---|---|---|---|---|---|
| positive regulation of transmembrane receptor protein serine/threonine kinase signaling pathway (GO:0090100) | 102 | 35 | 11.76 | + | 2.98 | 1.16E-09 | 5.69E-08 |
| pancreas development (GO:0031016) | 70 | 24 | 8.07 | + | 2.97 | 4.68E-07 | 1.53E-05 |
| regulation of stem cell proliferation (GO:0072091) | 82 | 28 | 9.45 | + | 2.96 | 5.86E-08 | 2.27E-06 |
| DNA damage response, signal transduction by p53 class mediator (GO:0030330) | 41 | 14 | 4.73 | + | 2.96 | 1.20E-04 | 2.29E-03 |
| regulation of miRNA metabolic process (GO:2000628) | 85 | 29 | 9.8 | + | 2.96 | 3.49E-08 | 1.40E-06 |
| exocrine system development (GO:0035272) | 44 | 15 | 5.07 | + | 2.96 | 7.05E-05 | 1.46E-03 |
| mesenchyme development (GO:0060485) | 244 | 83 | 28.12 | + | 2.95 | 1.09E-20 | 1.27E-18 |
| cell differentiation in spinal cord (GO:0021515) | 50 | 17 | 5.76 | + | 2.95 | 2.45E-05 | 5.75E-04 |
| regulation of neuroblast proliferation (GO:1902692) | 53 | 18 | 6.11 | + | 2.95 | 1.45E-05 | 3.57E-04 |
| negative regulation of fat cell differentiation (GO:0045599) | 53 | 18 | 6.11 | + | 2.95 | 1.45E-05 | 3.57E-04 |
| positive regulation of cell junction assembly (GO:1901890) | 109 | 37 | 12.56 | + | 2.95 | 5.58E-10 | 2.89E-08 |
| regulation of vascular associated smooth muscle cell proliferation (GO:1904705) | 59 | 20 | 6.8 | + | 2.94 | 5.08E-06 | 1.37E-04 |
| mammary gland epithelium development (GO:0061180) | 59 | 20 | 6.8 | + | 2.94 | 5.08E-06 | 1.37E-04 |
| positive regulation of neuron apoptotic process (GO:0043525) | 65 | 22 | 7.49 | + | 2.94 | 1.79E-06 | 5.25E-05 |
| regulation of cell-substrate junction organization (GO:0150116) | 68 | 23 | 7.84 | + | 2.93 | 1.06E-06 | 3.23E-05 |
| heart valve development (GO:0003170) | 74 | 25 | 8.53 | + | 2.93 | 3.74E-07 | 1.24E-05 |

| | | | | | | | |
|---|---|---|---|---|---|---|---|
| outflow tract morphogenesis (GO:0003151) | 77 | 26 | 8.87 | + | 2.93 | 2.23E-07 | 7.70E-06 |
| epithelial tube formation (GO:0072175) | 131 | 44 | 15.1 | + | 2.91 | 2.02E-11 | 1.24E-09 |
| neural tube development (GO:0021915) | 158 | 53 | 18.21 | + | 2.91 | 1.94E-13 | 1.48E-11 |
| cerebral cortex radially oriented cell migration (GO:0021799) | 36 | 12 | 4.15 | + | 2.89 | 4.68E-04 | 7.29E-03 |
| negative regulation of neuron differentiation (GO:0045665) | 72 | 24 | 8.3 | + | 2.89 | 8.42E-07 | 2.65E-05 |
| positive regulation of erythrocyte differentiation (GO:0045648) | 30 | 10 | 3.46 | + | 2.89 | 1.38E-03 | 1.83E-02 |
| regulation of post-transcriptional gene silencing (GO:0060147) | 30 | 10 | 3.46 | + | 2.89 | 1.38E-03 | 1.83E-02 |
| positive regulation of stem cell population maintenance (GO:1902459) | 48 | 16 | 5.53 | + | 2.89 | 5.57E-05 | 1.20E-03 |
| regulation of cardiac muscle cell proliferation (GO:0060043) | 36 | 12 | 4.15 | + | 2.89 | 4.68E-04 | 7.28E-03 |
| mitotic G1/S transition checkpoint signaling (GO:0044819) | 30 | 10 | 3.46 | + | 2.89 | 1.38E-03 | 1.83E-02 |
| regulation of protein sumoylation (GO:0033233) | 24 | 8 | 2.77 | + | 2.89 | 4.13E-03 | 4.48E-02 |
| regulation of anoikis (GO:2000209) | 24 | 8 | 2.77 | + | 2.89 | 4.13E-03 | 4.48E-02 |
| regulation of stem cell population maintenance (GO:2000036) | 72 | 24 | 8.3 | + | 2.89 | 8.42E-07 | 2.65E-05 |
| neuron fate commitment (GO:0048663) | 72 | 24 | 8.3 | + | 2.89 | 8.42E-07 | 2.64E-05 |
| cochlea morphogenesis (GO:0090103) | 24 | 8 | 2.77 | + | 2.89 | 4.13E-03 | 4.48E-02 |
| protein heterooligomerization (GO:0051291) | 30 | 10 | 3.46 | + | 2.89 | 1.38E-03 | 1.83E-02 |

| | | | | | | | |
|---|---|---|---|---|---|---|---|
| negative regulation of axonogenesis (GO:0050771) | 51 | 17 | 5.88 | + | 2.89 | 3.29E-05 | 7.40E-04 |
| mitotic G1 DNA damage checkpoint signaling (GO:0031571) | 30 | 10 | 3.46 | + | 2.89 | 1.38E-03 | 1.83E-02 |
| semi-lunar valve development (GO:1905314) | 48 | 16 | 5.53 | + | 2.89 | 5.57E-05 | 1.20E-03 |
| negative regulation of epithelial cell differentiation (GO:0030857) | 48 | 16 | 5.53 | + | 2.89 | 5.57E-05 | 1.20E-03 |
| regulation of hormone biosynthetic process (GO:0046885) | 24 | 8 | 2.77 | + | 2.89 | 4.13E-03 | 4.47E-02 |
| gland morphogenesis (GO:0022612) | 105 | 35 | 12.1 | + | 2.89 | 2.81E-09 | 1.29E-07 |
| striated muscle cell proliferation (GO:0014855) | 36 | 12 | 4.15 | + | 2.89 | 4.68E-04 | 7.28E-03 |
| response to fluid shear stress (GO:0034405) | 33 | 11 | 3.8 | + | 2.89 | 8.02E-04 | 1.16E-02 |
| endocardial cushion development (GO:0003197) | 51 | 17 | 5.88 | + | 2.89 | 3.29E-05 | 7.39E-04 |
| pulmonary valve development (GO:0003177) | 24 | 8 | 2.77 | + | 2.89 | 4.13E-03 | 4.47E-02 |
| neuroepithelial cell differentiation (GO:0060563) | 33 | 11 | 3.8 | + | 2.89 | 8.02E-04 | 1.16E-02 |
| response to X-ray (GO:0010165) | 27 | 9 | 3.11 | + | 2.89 | 2.38E-03 | 2.84E-02 |
| negative regulation of gliogenesis (GO:0014014) | 42 | 14 | 4.84 | + | 2.89 | 1.61E-04 | 2.93E-03 |
| animal organ formation (GO:0048645) | 39 | 13 | 4.5 | + | 2.89 | 2.74E-04 | 4.64E-03 |
| mesodermal cell differentiation (GO:0048333) | 27 | 9 | 3.11 | + | 2.89 | 2.38E-03 | 2.84E-02 |
| negative regulation of telomere maintenance via telomere lengthening (GO:1904357) | 27 | 9 | 3.11 | + | 2.89 | 2.38E-03 | 2.83E-02 |

| | | | | | | | |
|---|---|---|---|---|---|---|---|
| embryonic organ morphogenesis (GO:0048562) | 295 | 98 | 34 | + | 2.88 | 2.55E-23 | 3.16E-21 |
| renal system development (GO:0072001) | 314 | 104 | 36.19 | + | 2.87 | 1.48E-24 | 1.98E-22 |
| tube formation (GO:0035148) | 145 | 48 | 16.71 | + | 2.87 | 4.54E-12 | 2.94E-10 |
| canonical Wnt signaling pathway (GO:0060070) | 106 | 35 | 12.22 | + | 2.86 | 3.73E-09 | 1.68E-07 |
| neural tube formation (GO:0001841) | 103 | 34 | 11.87 | + | 2.86 | 6.26E-09 | 2.71E-07 |
| cartilage development (GO:0051216) | 173 | 57 | 19.94 | + | 2.86 | 5.79E-14 | 4.51E-12 |
| transforming growth factor beta receptor superfamily signaling pathway (GO:0141091) | 170 | 56 | 19.59 | + | 2.86 | 9.71E-14 | 7.45E-12 |
| negative regulation of nervous system development (GO:0051961) | 137 | 45 | 15.79 | + | 2.85 | 2.83E-11 | 1.71E-09 |
| negative regulation of neurogenesis (GO:0050768) | 131 | 43 | 15.1 | + | 2.85 | 7.94E-11 | 4.55E-09 |
| dendrite development (GO:0016358) | 113 | 37 | 13.02 | + | 2.84 | 1.76E-09 | 8.28E-08 |
| mesenchyme morphogenesis (GO:0072132) | 55 | 18 | 6.34 | + | 2.84 | 2.58E-05 | 5.98E-04 |
| neural crest cell migration (GO:0001755) | 55 | 18 | 6.34 | + | 2.84 | 2.58E-05 | 5.98E-04 |
| epithelial tube morphogenesis (GO:0060562) | 312 | 102 | 35.96 | + | 2.84 | 1.29E-23 | 1.64E-21 |
| digestive tract morphogenesis (GO:0048546) | 49 | 16 | 5.65 | + | 2.83 | 7.39E-05 | 1.52E-03 |
| visual learning (GO:0008542) | 49 | 16 | 5.65 | + | 2.83 | 7.39E-05 | 1.51E-03 |
| cell-matrix adhesion (GO:0007160) | 138 | 45 | 15.91 | + | 2.83 | 3.73E-11 | 2.23E-09 |

| | | | | | | | |
|---|---|---|---|---|---|---|---|
| calcium-dependent cell-cell adhesion via plasma membrane cell adhesion molecules (GO:0016339) | 46 | 15 | 5.3 | + | 2.83 | 1.25E-04 | 2.36E-03 |
| glial cell migration (GO:0008347) | 46 | 15 | 5.3 | + | 2.83 | 1.25E-04 | 2.36E-03 |
| regulation of embryonic development (GO:0045995) | 92 | 30 | 10.6 | + | 2.83 | 6.56E-08 | 2.52E-06 |
| cellular response to BMP stimulus (GO:0071773) | 89 | 29 | 10.26 | + | 2.83 | 1.10E-07 | 4.02E-06 |
| response to BMP (GO:0071772) | 89 | 29 | 10.26 | + | 2.83 | 1.10E-07 | 4.01E-06 |
| digestive tract development (GO:0048565) | 132 | 43 | 15.21 | + | 2.83 | 1.05E-10 | 5.91E-09 |
| kidney development (GO:0001822) | 304 | 99 | 35.04 | + | 2.83 | 8.16E-23 | 1.00E-20 |
| semaphorin-plexin signaling pathway (GO:0071526) | 43 | 14 | 4.96 | + | 2.82 | 2.13E-04 | 3.75E-03 |
| cellular response to epidermal growth factor stimulus (GO:0071364) | 43 | 14 | 4.96 | + | 2.82 | 2.13E-04 | 3.75E-03 |
| positive regulation of substrate adhesion-dependent cell spreading (GO:1900026) | 43 | 14 | 4.96 | + | 2.82 | 2.13E-04 | 3.74E-03 |
| positive regulation of organ growth (GO:0046622) | 43 | 14 | 4.96 | + | 2.82 | 2.13E-04 | 3.74E-03 |
| maintenance of cell number (GO:0098727) | 120 | 39 | 13.83 | + | 2.82 | 8.25E-10 | 4.16E-08 |
| regulation of stem cell differentiation (GO:2000736) | 80 | 26 | 9.22 | + | 2.82 | 5.22E-07 | 1.70E-05 |
| neuron projection extension (GO:1990138) | 80 | 26 | 9.22 | + | 2.82 | 5.22E-07 | 1.69E-05 |
| secondary alcohol biosynthetic process (GO:1902653) | 37 | 12 | 4.26 | + | 2.81 | 6.21E-04 | 9.40E-03 |
| cholesterol biosynthetic process (GO:0006695) | 37 | 12 | 4.26 | + | 2.81 | 6.21E-04 | 9.39E-03 |

| | | | | | | | |
|---|---|---|---|---|---|---|---|
| protein localization to chromatin (GO:0071168) | 37 | 12 | 4.26 | + | 2.81 | 6.21E-04 | 9.38E-03 |
| regulation of DNA damage response, signal transduction by p53 class mediator (GO:0043516) | 37 | 12 | 4.26 | + | 2.81 | 6.21E-04 | 9.37E-03 |
| regulation of hippo signaling (GO:0035330) | 37 | 12 | 4.26 | + | 2.81 | 6.21E-04 | 9.36E-03 |
| cell surface receptor protein serine/threonine kinase signaling pathway (GO:0007178) | 182 | 59 | 20.98 | + | 2.81 | 4.73E-14 | 3.75E-12 |
| regulation of cartilage development (GO:0061035) | 71 | 23 | 8.18 | + | 2.81 | 2.48E-06 | 7.08E-05 |
| embryonic heart tube morphogenesis (GO:0003143) | 71 | 23 | 8.18 | + | 2.81 | 2.48E-06 | 7.07E-05 |
| cardiac cell development (GO:0055006) | 68 | 22 | 7.84 | + | 2.81 | 4.18E-06 | 1.15E-04 |
| negative regulation of smoothened signaling pathway (GO:0045879) | 34 | 11 | 3.92 | + | 2.81 | 1.06E-03 | 1.47E-02 |
| bone morphogenesis (GO:0060349) | 99 | 32 | 11.41 | + | 2.8 | 3.06E-08 | 1.24E-06 |
| peptidyl-tyrosine phosphorylation (GO:0018108) | 65 | 21 | 7.49 | + | 2.8 | 7.05E-06 | 1.85E-04 |
| eye morphogenesis (GO:0048592) | 161 | 52 | 18.56 | + | 2.8 | 1.75E-12 | 1.20E-10 |
| regulation of transforming growth factor beta production (GO:0071634) | 31 | 10 | 3.57 | + | 2.8 | 1.83E-03 | 2.30E-02 |
| negative regulation of developmental growth (GO:0048640) | 93 | 30 | 10.72 | + | 2.8 | 8.62E-08 | 3.23E-06 |
| negative regulation of axon extension (GO:0030517) | 31 | 10 | 3.57 | + | 2.8 | 1.83E-03 | 2.29E-02 |
| regulation of substrate adhesion-dependent cell spreading (GO:1900024) | 62 | 20 | 7.15 | + | 2.8 | 1.19E-05 | 2.97E-04 |
| positive regulation of cell-substrate junction organization (GO:0150117) | 31 | 10 | 3.57 | + | 2.8 | 1.83E-03 | 2.29E-02 |

| | | | | | | | |
|---|---|---|---|---|---|---|---|
| regulation of gene silencing by regulatory ncRNA (GO:0060966) | 31 | 10 | 3.57 | + | 2.8 | 1.83E-03 | 2.29E-02 |
| regulation of synapse assembly (GO:0051963) | 121 | 39 | 13.95 | + | 2.8 | 1.08E-09 | 5.37E-08 |
| associative learning (GO:0008306) | 87 | 28 | 10.03 | + | 2.79 | 2.43E-07 | 8.30E-06 |
| digestive system development (GO:0055123) | 143 | 46 | 16.48 | + | 2.79 | 3.85E-11 | 2.29E-09 |
| regulation of osteoblast proliferation (GO:0033688) | 28 | 9 | 3.23 | + | 2.79 | 3.15E-03 | 3.56E-02 |
| subpallium development (GO:0021544) | 28 | 9 | 3.23 | + | 2.79 | 3.15E-03 | 3.56E-02 |
| miRNA-mediated post-transcriptional gene silencing (GO:0035195) | 28 | 9 | 3.23 | + | 2.79 | 3.15E-03 | 3.55E-02 |
| dendritic spine development (GO:0060996) | 28 | 9 | 3.23 | + | 2.79 | 3.15E-03 | 3.55E-02 |
| embryonic hindlimb morphogenesis (GO:0035116) | 28 | 9 | 3.23 | + | 2.79 | 3.15E-03 | 3.55E-02 |
| regulation of miRNA-mediated gene silencing (GO:0060964) | 28 | 9 | 3.23 | + | 2.79 | 3.15E-03 | 3.55E-02 |
| endochondral bone growth (GO:0003416) | 28 | 9 | 3.23 | + | 2.79 | 3.15E-03 | 3.54E-02 |
| cardiac chamber development (GO:0003205) | 168 | 54 | 19.36 | + | 2.79 | 8.17E-13 | 5.88E-11 |
| morphogenesis of a branching epithelium (GO:0061138) | 165 | 53 | 19.02 | + | 2.79 | 1.37E-12 | 9.53E-11 |
| camera-type eye morphogenesis (GO:0048593) | 134 | 43 | 15.44 | + | 2.78 | 1.80E-10 | 9.92E-09 |
| heterophilic cell-cell adhesion via plasma membrane cell adhesion molecules (GO:0007157) | 53 | 17 | 6.11 | + | 2.78 | 5.74E-05 | 1.23E-03 |
| regulation of morphogenesis of a branching structure (GO:0060688) | 53 | 17 | 6.11 | + | 2.78 | 5.74E-05 | 1.22E-03 |

| | | | | | | | |
|---|---|---|---|---|---|---|---|
| mesenchymal cell differentiation (GO:0048762) | 172 | 55 | 19.82 | + | 2.77 | 6.38E-13 | 4.68E-11 |
| axonogenesis (GO:0007409) | 366 | 117 | 42.18 | + | 2.77 | 5.93E-26 | 8.38E-24 |
| development of primary female sexual characteristics (GO:0046545) | 97 | 31 | 11.18 | + | 2.77 | 6.72E-08 | 2.57E-06 |
| stem cell proliferation (GO:0072089) | 72 | 23 | 8.3 | + | 2.77 | 3.25E-06 | 9.11E-05 |
| positive regulation of stem cell proliferation (GO:2000648) | 47 | 15 | 5.42 | + | 2.77 | 1.65E-04 | 3.00E-03 |
| response to epidermal growth factor (GO:0070849) | 47 | 15 | 5.42 | + | 2.77 | 1.65E-04 | 3.00E-03 |
| stem cell population maintenance (GO:0019827) | 116 | 37 | 13.37 | + | 2.77 | 3.97E-09 | 1.78E-07 |
| developmental growth involved in morphogenesis (GO:0060560) | 138 | 44 | 15.91 | + | 2.77 | 1.41E-10 | 7.88E-09 |
| plasma membrane bounded cell projection morphogenesis (GO:0120039) | 490 | 156 | 56.48 | + | 2.76 | 4.62E-34 | 1.07E-31 |
| columnar/cuboidal epithelial cell development (GO:0002066) | 44 | 14 | 5.07 | + | 2.76 | 2.80E-04 | 4.70E-03 |
| sterol biosynthetic process (GO:0016126) | 44 | 14 | 5.07 | + | 2.76 | 2.80E-04 | 4.69E-03 |
| aortic valve development (GO:0003176) | 44 | 14 | 5.07 | + | 2.76 | 2.80E-04 | 4.69E-03 |
| columnar/cuboidal epithelial cell differentiation (GO:0002065) | 107 | 34 | 12.33 | + | 2.76 | 1.86E-08 | 7.76E-07 |
| embryonic heart tube development (GO:0035050) | 85 | 27 | 9.8 | + | 2.76 | 5.32E-07 | 1.72E-05 |
| cardiac chamber morphogenesis (GO:0003206) | 126 | 40 | 14.52 | + | 2.75 | 1.10E-09 | 5.45E-08 |
| morphogenesis of embryonic epithelium (GO:0016331) | 145 | 46 | 16.71 | + | 2.75 | 6.55E-11 | 3.82E-09 |

| | | | | | | | |
|---|---|---|---|---|---|---|---|
| embryonic epithelial tube formation (GO:0001838) | 123 | 39 | 14.18 | + | 2.75 | 1.84E-09 | 8.61E-08 |
| regulation of keratinocyte differentiation (GO:0045616) | 38 | 12 | 4.38 | + | 2.74 | 8.13E-04 | 1.17E-02 |
| neuron projection morphogenesis (GO:0048812) | 485 | 153 | 55.9 | + | 2.74 | 6.53E-33 | 1.33E-30 |
| cell projection morphogenesis (GO:0048858) | 495 | 156 | 57.05 | + | 2.73 | 1.73E-33 | 3.69E-31 |
| cranial skeletal system development (GO:1904888) | 73 | 23 | 8.41 | + | 2.73 | 4.24E-06 | 1.16E-04 |
| cardiac ventricle development (GO:0003231) | 127 | 40 | 14.64 | + | 2.73 | 1.43E-09 | 6.90E-08 |
| lung alveolus development (GO:0048286) | 54 | 17 | 6.22 | + | 2.73 | 7.48E-05 | 1.53E-03 |
| axon development (GO:0061564) | 420 | 132 | 48.41 | + | 2.73 | 2.46E-28 | 4.01E-26 |
| cardiac ventricle morphogenesis (GO:0003208) | 70 | 22 | 8.07 | + | 2.73 | 7.12E-06 | 1.85E-04 |
| peptidyl-tyrosine modification (GO:0018212) | 67 | 21 | 7.72 | + | 2.72 | 1.20E-05 | 2.99E-04 |
| limb morphogenesis (GO:0035108) | 150 | 47 | 17.29 | + | 2.72 | 6.59E-11 | 3.83E-09 |
| appendage morphogenesis (GO:0035107) | 150 | 47 | 17.29 | + | 2.72 | 6.59E-11 | 3.82E-09 |
| morphogenesis of a branching structure (GO:0001763) | 176 | 55 | 20.29 | + | 2.71 | 1.81E-12 | 1.24E-10 |
| cerebral cortex cell migration (GO:0021795) | 48 | 15 | 5.53 | + | 2.71 | 2.14E-04 | 3.74E-03 |
| cartilage development involved in endochondral bone morphogenesis (GO:0060351) | 32 | 10 | 3.69 | + | 2.71 | 2.39E-03 | 2.84E-02 |
| ligand-gated ion channel signaling pathway (GO:1990806) | 32 | 10 | 3.69 | + | 2.71 | 2.39E-03 | 2.84E-02 |

| | | | | | | | |
|---|---|---|---|---|---|---|---|
| modulation of excitatory postsynaptic potential (GO:0098815) | 48 | 15 | 5.53 | + | 2.71 | 2.14E-04 | 3.74E-03 |
| female sex differentiation (GO:0046660) | 112 | 35 | 12.91 | + | 2.71 | 1.87E-08 | 7.78E-07 |
| cardiac muscle hypertrophy (GO:0003300) | 32 | 10 | 3.69 | + | 2.71 | 2.39E-03 | 2.84E-02 |
| female gonad development (GO:0008585) | 93 | 29 | 10.72 | + | 2.71 | 3.18E-07 | 1.07E-05 |
| connective tissue development (GO:0061448) | 231 | 72 | 26.62 | + | 2.7 | 8.14E-16 | 7.20E-14 |
| regulation of cell-substrate adhesion (GO:0010810) | 215 | 67 | 24.78 | + | 2.7 | 8.24E-15 | 6.70E-13 |
| ameboidal-type cell migration (GO:0001667) | 199 | 62 | 22.94 | + | 2.7 | 8.32E-14 | 6.45E-12 |
| cell morphogenesis involved in neuron differentiation (GO:0048667) | 443 | 138 | 51.06 | + | 2.7 | 3.85E-29 | 6.69E-27 |
| embryonic limb morphogenesis (GO:0030326) | 122 | 38 | 14.06 | + | 2.7 | 5.18E-09 | 2.27E-07 |
| embryonic appendage morphogenesis (GO:0035113) | 122 | 38 | 14.06 | + | 2.7 | 5.18E-09 | 2.26E-07 |
| synaptic transmission, glutamatergic (GO:0035249) | 45 | 14 | 5.19 | + | 2.7 | 3.64E-04 | 5.86E-03 |
| regulation of epidermal cell differentiation (GO:0045604) | 58 | 18 | 6.68 | + | 2.69 | 5.73E-05 | 1.23E-03 |
| spinal cord motor neuron differentiation (GO:0021522) | 29 | 9 | 3.34 | + | 2.69 | 4.11E-03 | 4.47E-02 |
| cardiac atrium morphogenesis (GO:0003209) | 29 | 9 | 3.34 | + | 2.69 | 4.11E-03 | 4.47E-02 |
| mammary gland development (GO:0030879) | 126 | 39 | 14.52 | + | 2.69 | 3.99E-09 | 1.78E-07 |
| myoblast differentiation (GO:0045445) | 42 | 13 | 4.84 | + | 2.69 | 6.18E-04 | 9.37E-03 |

| | | | | | | | | |
|---|---|---|---|---|---|---|---|---|
| nerve development (GO:0021675) | 97 | 30 | 11.18 | + | | 2.68 | 2.45E-07 | 8.36E-06 |
| negative regulation of myeloid cell differentiation (GO:0045638) | 81 | 25 | 9.34 | + | | 2.68 | 2.51E-06 | 7.13E-05 |
| cell-substrate adhesion (GO:0031589) | 185 | 57 | 21.32 | + | | 2.67 | 1.39E-12 | 9.64E-11 |
| blood vessel morphogenesis (GO:0048514) | 432 | 133 | 49.79 | + | | 2.67 | 1.46E-27 | 2.27E-25 |
| positive regulation of SMAD protein signal transduction (GO:0060391) | 39 | 12 | 4.5 | + | | 2.67 | 1.05E-03 | 1.46E-02 |
| neural crest cell differentiation (GO:0014033) | 91 | 28 | 10.49 | + | | 2.67 | 6.86E-07 | 2.19E-05 |
| neural crest cell development (GO:0014032) | 78 | 24 | 8.99 | + | | 2.67 | 4.20E-06 | 1.16E-04 |
| cellular response to alkaloid (GO:0071312) | 39 | 12 | 4.5 | + | | 2.67 | 1.05E-03 | 1.46E-02 |
| spliceosomal complex assembly (GO:0000245) | 75 | 23 | 8.64 | + | | 2.66 | 7.05E-06 | 1.84E-04 |
| anterior/posterior pattern specification (GO:0009952) | 212 | 65 | 24.43 | + | | 2.66 | 4.88E-14 | 3.85E-12 |
| heart morphogenesis (GO:0003007) | 258 | 79 | 29.74 | + | | 2.66 | 1.01E-16 | 9.82E-15 |
| lung morphogenesis (GO:0060425) | 49 | 15 | 5.65 | + | | 2.66 | 2.76E-04 | 4.65E-03 |
| thymus development (GO:0048538) | 49 | 15 | 5.65 | + | | 2.66 | 2.76E-04 | 4.65E-03 |
| epithelial to mesenchymal transition (GO:0001837) | 85 | 26 | 9.8 | + | | 2.65 | 1.92E-06 | 5.64E-05 |
| regulation of cell-matrix adhesion (GO:0001952) | 121 | 37 | 13.95 | + | | 2.65 | 1.43E-08 | 6.02E-07 |
| positive regulation of vascular associated smooth muscle cell proliferation (GO:1904707) | 36 | 11 | 4.15 | + | | 2.65 | 1.80E-03 | 2.27E-02 |

| | | | | | | | |
|---|---|---|---|---|---|---|---|
| regulatory ncRNA-mediated post-transcriptional gene silencing (GO:0035194) | 36 | 11 | 4.15 | + | 2.65 | 1.80E-03 | 2.27E-02 |
| hippocampus development (GO:0021766) | 95 | 29 | 10.95 | + | 2.65 | 5.26E-07 | 1.70E-05 |
| inner ear morphogenesis (GO:0042472) | 105 | 32 | 12.1 | + | 2.64 | 1.44E-07 | 5.18E-06 |
| ventral spinal cord development (GO:0021517) | 46 | 14 | 5.3 | + | 2.64 | 4.68E-04 | 7.30E-03 |
| trabecula morphogenesis (GO:0061383) | 46 | 14 | 5.3 | + | 2.64 | 4.68E-04 | 7.30E-03 |
| regulation of neural precursor cell proliferation (GO:2000177) | 102 | 31 | 11.76 | + | 2.64 | 2.41E-07 | 8.26E-06 |
| positive regulation of miRNA transcription (GO:1902895) | 56 | 17 | 6.45 | + | 2.63 | 1.24E-04 | 2.36E-03 |
| regulation of neuron differentiation (GO:0045664) | 188 | 57 | 21.67 | + | 2.63 | 2.91E-12 | 1.91E-10 |
| regulation of transmembrane receptor protein serine/threonine kinase signaling pathway (GO:0090092) | 264 | 80 | 30.43 | + | 2.63 | 1.26E-16 | 1.20E-14 |
| striated muscle hypertrophy (GO:0014897) | 33 | 10 | 3.8 | + | 2.63 | 3.08E-03 | 3.52E-02 |
| cellular response to growth factor stimulus (GO:0071363) | 472 | 143 | 54.4 | + | 2.63 | 9.32E-29 | 1.58E-26 |
| neuron migration (GO:0001764) | 142 | 43 | 16.37 | + | 2.63 | 1.39E-09 | 6.72E-08 |
| BMP signaling pathway (GO:0030509) | 76 | 23 | 8.76 | + | 2.63 | 9.01E-06 | 2.32E-04 |
| synapse assembly (GO:0007416) | 119 | 36 | 13.72 | + | 2.62 | 3.04E-08 | 1.23E-06 |
| regulation of erythrocyte differentiation (GO:0045646) | 43 | 13 | 4.96 | + | 2.62 | 7.94E-04 | 1.16E-02 |
| cellular response to vascular endothelial growth factor stimulus (GO:0035924) | 43 | 13 | 4.96 | + | 2.62 | 7.94E-04 | 1.15E-02 |

| | | | | | | | |
|---|---|---|---|---|---|---|---|
| positive regulation of double-strand break repair via homologous recombination (GO:1905168) | 43 | 13 | 4.96 | + | 2.62 | 7.94E-04 | 1.15E-02 |
| tube morphogenesis (GO:0035239) | 682 | 206 | 78.61 | + | 2.62 | 5.61E-41 | 2.50E-38 |
| blood vessel development (GO:0001568) | 527 | 159 | 60.74 | + | 2.62 | 1.18E-31 | 2.21E-29 |
| urogenital system development (GO:0001655) | 63 | 19 | 7.26 | + | 2.62 | 5.60E-05 | 1.20E-03 |
| regulation of fat cell differentiation (GO:0045598) | 133 | 40 | 15.33 | + | 2.61 | 6.39E-09 | 2.76E-07 |
| response to growth factor (GO:0070848) | 503 | 151 | 57.97 | + | 2.6 | 7.79E-30 | 1.39E-27 |
| telencephalon cell migration (GO:0022029) | 60 | 18 | 6.92 | + | 2.6 | 9.40E-05 | 1.88E-03 |
| developmental cell growth (GO:0048588) | 110 | 33 | 12.68 | + | 2.6 | 1.40E-07 | 5.05E-06 |
| cardiac muscle tissue morphogenesis (GO:0055008) | 60 | 18 | 6.92 | + | 2.6 | 9.40E-05 | 1.87E-03 |
| cellular response to nutrient (GO:0031670) | 40 | 12 | 4.61 | + | 2.6 | 1.35E-03 | 1.80E-02 |
| cAMP-mediated signaling (GO:0019933) | 40 | 12 | 4.61 | + | 2.6 | 1.35E-03 | 1.80E-02 |
| endocardial cushion morphogenesis (GO:0003203) | 40 | 12 | 4.61 | + | 2.6 | 1.35E-03 | 1.80E-02 |
| lactation (GO:0007595) | 40 | 12 | 4.61 | + | 2.6 | 1.35E-03 | 1.80E-02 |
| embryonic morphogenesis (GO:0048598) | 588 | 176 | 67.77 | + | 2.6 | 1.74E-34 | 4.17E-32 |
| positive regulation of muscle cell differentiation (GO:0051149) | 67 | 20 | 7.72 | + | 2.59 | 4.24E-05 | 9.36E-04 |
| gastrulation (GO:0007369) | 171 | 51 | 19.71 | + | 2.59 | 7.82E-11 | 4.50E-09 |

| | | | | | | | |
|---|---|---|---|---|---|---|---|
| neuroblast proliferation (GO:0007405) | 57 | 17 | 6.57 | + | 2.59 | 1.58E-04 | 2.89E-03 |
| leukocyte apoptotic process (GO:0071887) | 47 | 14 | 5.42 | + | 2.58 | 5.97E-04 | 9.09E-03 |
| ventricular cardiac muscle tissue morphogenesis (GO:0055010) | 47 | 14 | 5.42 | + | 2.58 | 5.97E-04 | 9.08E-03 |
| stem cell development (GO:0048864) | 84 | 25 | 9.68 | + | 2.58 | 5.21E-06 | 1.40E-04 |
| cellular response to fibroblast growth factor stimulus (GO:0044344) | 84 | 25 | 9.68 | + | 2.58 | 5.21E-06 | 1.40E-04 |
| formation of primary germ layer (GO:0001704) | 121 | 36 | 13.95 | + | 2.58 | 4.89E-08 | 1.92E-06 |
| positive regulation of neuroblast proliferation (GO:0002052) | 37 | 11 | 4.26 | + | 2.58 | 2.30E-03 | 2.77E-02 |
| astrocyte development (GO:0014002) | 37 | 11 | 4.26 | + | 2.58 | 2.30E-03 | 2.77E-02 |
| lymphocyte apoptotic process (GO:0070227) | 37 | 11 | 4.26 | + | 2.58 | 2.30E-03 | 2.76E-02 |
| positive regulation of morphogenesis of an epithelium (GO:1905332) | 37 | 11 | 4.26 | + | 2.58 | 2.30E-03 | 2.76E-02 |
| embryonic camera-type eye development (GO:0031076) | 37 | 11 | 4.26 | + | 2.58 | 2.30E-03 | 2.76E-02 |
| positive regulation of smoothened signaling pathway (GO:0045880) | 37 | 11 | 4.26 | + | 2.58 | 2.30E-03 | 2.76E-02 |
| cardiac conduction system development (GO:0003161) | 37 | 11 | 4.26 | + | 2.58 | 2.30E-03 | 2.75E-02 |
| regulation of cellular response to growth factor stimulus (GO:0090287) | 313 | 93 | 36.08 | + | 2.58 | 1.77E-18 | 1.94E-16 |
| regulation of cell junction assembly (GO:1901888) | 219 | 65 | 25.24 | + | 2.58 | 2.63E-13 | 1.97E-11 |
| stem cell differentiation (GO:0048863) | 182 | 54 | 20.98 | + | 2.57 | 2.73E-11 | 1.66E-09 |

| | | | | | | | |
|---|---|---|---|---|---|---|---|
| embryonic organ development (GO:0048568) | 452 | 134 | 52.1 | + | 2.57 | 5.36E-26 | 7.64E-24 |
| positive regulation of ossification (GO:0045778) | 54 | 16 | 6.22 | + | 2.57 | 2.66E-04 | 4.52E-03 |
| muscle tissue morphogenesis (GO:0060415) | 71 | 21 | 8.18 | + | 2.57 | 3.21E-05 | 7.27E-04 |
| cellular response to ionizing radiation (GO:0071479) | 71 | 21 | 8.18 | + | 2.57 | 3.21E-05 | 7.26E-04 |
| vasculature development (GO:0001944) | 548 | 162 | 63.16 | + | 2.56 | 4.31E-31 | 7.94E-29 |
| positive regulation of bone mineralization (GO:0030501) | 44 | 13 | 5.07 | + | 2.56 | 1.01E-03 | 1.41E-02 |
| actin filament bundle assembly (GO:0051017) | 61 | 18 | 7.03 | + | 2.56 | 1.19E-04 | 2.28E-03 |
| cell growth (GO:0016049) | 112 | 33 | 12.91 | + | 2.56 | 2.24E-07 | 7.75E-06 |
| proximal/distal pattern formation (GO:0009954) | 34 | 10 | 3.92 | + | 2.55 | 3.91E-03 | 4.29E-02 |
| neuron fate specification (GO:0048665) | 34 | 10 | 3.92 | + | 2.55 | 3.91E-03 | 4.29E-02 |
| vascular endothelial growth factor receptor signaling pathway (GO:0048010) | 34 | 10 | 3.92 | + | 2.55 | 3.91E-03 | 4.28E-02 |
| miRNA processing (GO:0035196) | 34 | 10 | 3.92 | + | 2.55 | 3.91E-03 | 4.28E-02 |
| cardiocyte differentiation (GO:0035051) | 119 | 35 | 13.72 | + | 2.55 | 1.03E-07 | 3.77E-06 |
| morphogenesis of an epithelium (GO:0002009) | 456 | 134 | 52.56 | + | 2.55 | 1.36E-25 | 1.90E-23 |
| ear morphogenesis (GO:0042471) | 126 | 37 | 14.52 | + | 2.55 | 4.69E-08 | 1.84E-06 |
| sensory organ morphogenesis (GO:0090596) | 276 | 81 | 31.81 | + | 2.55 | 6.30E-16 | 5.64E-14 |

| | | | | | | | |
|---|---|---|---|---|---|---|---|
| positive regulation of small GTPase mediated signal transduction (GO:0051057) | 75 | 22 | 8.64 | + | 2.55 | 2.42E-05 | 5.68E-04 |
| regulation of cell fate commitment (GO:0010453) | 41 | 12 | 4.73 | + | 2.54 | 1.71E-03 | 2.20E-02 |
| heart looping (GO:0001947) | 65 | 19 | 7.49 | + | 2.54 | 8.97E-05 | 1.80E-03 |
| negative regulation of canonical Wnt signaling pathway (GO:0090090) | 137 | 40 | 15.79 | + | 2.53 | 1.62E-08 | 6.80E-07 |
| response to amyloid-beta (GO:1904645) | 48 | 14 | 5.53 | + | 2.53 | 7.54E-04 | 1.10E-02 |
| extracellular matrix organization (GO:0030198) | 275 | 80 | 31.7 | + | 2.52 | 1.65E-15 | 1.42E-13 |
| bone development (GO:0060348) | 196 | 57 | 22.59 | + | 2.52 | 1.89E-11 | 1.16E-09 |
| regulation of SMAD protein signal transduction (GO:0060390) | 62 | 18 | 7.15 | + | 2.52 | 1.50E-04 | 2.76E-03 |
| positive regulation of miRNA metabolic process (GO:2000630) | 62 | 18 | 7.15 | + | 2.52 | 1.50E-04 | 2.76E-03 |
| positive regulation of cell-substrate adhesion (GO:0010811) | 124 | 36 | 14.29 | + | 2.52 | 9.75E-08 | 3.60E-06 |
| negative regulation of Wnt signaling pathway (GO:0030178) | 169 | 49 | 19.48 | + | 2.52 | 5.39E-10 | 2.80E-08 |
| Rho protein signal transduction (GO:0007266) | 69 | 20 | 7.95 | + | 2.51 | 6.74E-05 | 1.40E-03 |
| telencephalon development (GO:0021537) | 276 | 80 | 31.81 | + | 2.51 | 2.06E-15 | 1.76E-13 |
| extracellular structure organization (GO:0043062) | 276 | 80 | 31.81 | + | 2.51 | 2.06E-15 | 1.75E-13 |
| cellular response to amyloid-beta (GO:1904646) | 38 | 11 | 4.38 | + | 2.51 | 2.90E-03 | 3.37E-02 |
| lung development (GO:0030324) | 190 | 55 | 21.9 | + | 2.51 | 5.16E-11 | 3.02E-09 |

| | | | | | | | |
|---|---|---|---|---|---|---|---|
| external encapsulating structure organization (GO:0045229) | 277 | 80 | 31.93 | + | 2.51 | 4.03E-15 | 3.37E-13 |
| positive regulation of neuron projection development (GO:0010976) | 156 | 45 | 17.98 | + | 2.5 | 3.22E-09 | 1.46E-07 |
| regulation of mRNA splicing, via spliceosome (GO:0048024) | 111 | 32 | 12.79 | + | 2.5 | 5.86E-07 | 1.88E-05 |
| central nervous system neuron differentiation (GO:0021953) | 177 | 51 | 20.4 | + | 2.5 | 3.08E-10 | 1.63E-08 |
| synapse organization (GO:0050808) | 337 | 97 | 38.84 | + | 2.5 | 4.40E-18 | 4.68E-16 |
| homophilic cell adhesion via plasma membrane adhesion molecules (GO:0007156) | 167 | 48 | 19.25 | + | 2.49 | 1.11E-09 | 5.47E-08 |
| regulation of organ growth (GO:0046620) | 87 | 25 | 10.03 | + | 2.49 | 1.03E-05 | 2.64E-04 |
| integrin-mediated signaling pathway (GO:0007229) | 94 | 27 | 10.83 | + | 2.49 | 4.69E-06 | 1.28E-04 |
| limbic system development (GO:0021761) | 126 | 36 | 14.52 | + | 2.48 | 1.52E-07 | 5.44E-06 |
| cell adhesion mediated by integrin (GO:0033627) | 42 | 12 | 4.84 | + | 2.48 | 2.15E-03 | 2.64E-02 |
| positive regulation of canonical Wnt signaling pathway (GO:0090263) | 105 | 30 | 12.1 | + | 2.48 | 1.60E-06 | 4.73E-05 |
| determination of heart left/right asymmetry (GO:0061371) | 70 | 20 | 8.07 | + | 2.48 | 8.42E-05 | 1.70E-03 |
| development of primary male sexual characteristics (GO:0046546) | 147 | 42 | 16.94 | + | 2.48 | 1.45E-08 | 6.08E-07 |
| ventricular cardiac muscle tissue development (GO:0003229) | 63 | 18 | 7.26 | + | 2.48 | 1.88E-04 | 3.33E-03 |
| somite development (GO:0061053) | 77 | 22 | 8.87 | + | 2.48 | 3.80E-05 | 8.48E-04 |
| forebrain cell migration (GO:0021885) | 63 | 18 | 7.26 | + | 2.48 | 1.88E-04 | 3.33E-03 |

| | | | | | | | |
|---|---|---|---|---|---|---|---|
| regulation of synapse organization (GO:0050807) | 249 | 71 | 28.7 | + | 2.47 | 2.37E-13 | 1.79E-11 |
| male sex differentiation (GO:0046661) | 169 | 48 | 19.48 | + | 2.46 | 2.66E-09 | 1.22E-07 |
| tissue morphogenesis (GO:0048729) | 567 | 161 | 65.35 | + | 2.46 | 1.23E-28 | 2.07E-26 |
| angiogenesis (GO:0001525) | 342 | 97 | 39.42 | + | 2.46 | 1.11E-17 | 1.15E-15 |
| respiratory tube development (GO:0030323) | 194 | 55 | 22.36 | + | 2.46 | 1.67E-10 | 9.23E-09 |
| negative regulation of neuron projection development (GO:0010977) | 127 | 36 | 14.64 | + | 2.46 | 1.89E-07 | 6.63E-06 |
| skeletal system development (GO:0001501) | 501 | 142 | 57.74 | + | 2.46 | 3.14E-25 | 4.32E-23 |
| response to fibroblast growth factor (GO:0071774) | 92 | 26 | 10.6 | + | 2.45 | 9.66E-06 | 2.47E-04 |
| post-transcriptional gene silencing (GO:0016441) | 46 | 13 | 5.3 | + | 2.45 | 1.59E-03 | 2.05E-02 |
| cell fate commitment (GO:0045165) | 255 | 72 | 29.39 | + | 2.45 | 2.43E-13 | 1.83E-11 |
| ossification (GO:0001503) | 287 | 81 | 33.08 | + | 2.45 | 7.60E-15 | 6.21E-13 |
| blood vessel remodeling (GO:0001974) | 39 | 11 | 4.5 | + | 2.45 | 3.62E-03 | 4.01E-02 |
| neuron projection development (GO:0031175) | 692 | 195 | 79.76 | + | 2.44 | 5.16E-34 | 1.18E-31 |
| regulation of epidermis development (GO:0045682) | 64 | 18 | 7.38 | + | 2.44 | 2.33E-04 | 4.04E-03 |
| positive regulation of neural precursor cell proliferation (GO:2000179) | 64 | 18 | 7.38 | + | 2.44 | 2.33E-04 | 4.03E-03 |
| actin filament bundle organization (GO:0061572) | 64 | 18 | 7.38 | + | 2.44 | 2.33E-04 | 4.03E-03 |

| | | | | | | | |
|---|---|---|---|---|---|---|---|
| forebrain development (GO:0030900) | 409 | 115 | 47.14 | + | 2.44 | 2.59E-20 | 2.97E-18 |
| male gonad development (GO:0008584) | 146 | 41 | 16.83 | + | 2.44 | 5.27E-08 | 2.05E-06 |
| visual behavior (GO:0007632) | 57 | 16 | 6.57 | + | 2.44 | 5.22E-04 | 8.05E-03 |
| regulation of filopodium assembly (GO:0051489) | 50 | 14 | 5.76 | + | 2.43 | 1.18E-03 | 1.61E-02 |
| cellular response to nerve growth factor stimulus (GO:1990090) | 50 | 14 | 5.76 | + | 2.43 | 1.18E-03 | 1.60E-02 |
| enzyme-linked receptor protein signaling pathway (GO:0007167) | 611 | 171 | 70.42 | + | 2.43 | 1.58E-29 | 2.78E-27 |
| limb development (GO:0060173) | 186 | 52 | 21.44 | + | 2.43 | 7.96E-10 | 4.04E-08 |
| appendage development (GO:0048736) | 186 | 52 | 21.44 | + | 2.43 | 7.96E-10 | 4.03E-08 |
| epithelial cell migration (GO:0010631) | 104 | 29 | 11.99 | + | 2.42 | 4.06E-06 | 1.13E-04 |
| regulation of synapse structure or activity (GO:0050803) | 255 | 71 | 29.39 | + | 2.42 | 7.04E-13 | 5.12E-11 |
| regulation of nervous system development (GO:0051960) | 450 | 125 | 51.87 | + | 2.41 | 1.37E-21 | 1.65E-19 |
| positive regulation of fibroblast proliferation (GO:0048146) | 54 | 15 | 6.22 | + | 2.41 | 8.70E-04 | 1.25E-02 |
| protein autophosphorylation (GO:0046777) | 155 | 43 | 17.86 | + | 2.41 | 3.08E-08 | 1.24E-06 |
| lens development in camera-type eye (GO:0002088) | 83 | 23 | 9.57 | + | 2.4 | 4.35E-05 | 9.59E-04 |
| development of primary sexual characteristics (GO:0045137) | 231 | 64 | 26.62 | + | 2.4 | 1.21E-11 | 7.64E-10 |
| regulation of morphogenesis of an epithelium (GO:1905330) | 65 | 18 | 7.49 | + | 2.4 | 2.88E-04 | 4.79E-03 |

| | | | | | | | |
|---|---|---|---|---|---|---|---|
| regulation of RNA splicing (GO:0043484) | 185 | 51 | 21.32 | + | 2.39 | 1.82E-09 | 8.54E-08 |
| cell morphogenesis (GO:0000902) | 686 | 189 | 79.07 | + | 2.39 | 1.11E-31 | 2.13E-29 |
| negative regulation of G1/S transition of mitotic cell cycle (GO:2000134) | 69 | 19 | 7.95 | + | 2.39 | 2.14E-04 | 3.75E-03 |
| regulation of actin filament-based movement (GO:1903115) | 40 | 11 | 4.61 | + | 2.39 | 4.49E-03 | 4.83E-02 |
| mesoderm development (GO:0007498) | 131 | 36 | 15.1 | + | 2.38 | 5.38E-07 | 1.74E-05 |
| cardiac muscle tissue development (GO:0048738) | 193 | 53 | 22.24 | + | 2.38 | 9.97E-10 | 4.97E-08 |
| tube development (GO:0035295) | 900 | 247 | 103.73 | + | 2.38 | 6.23E-41 | 2.69E-38 |
| head development (GO:0060322) | 780 | 214 | 89.9 | + | 2.38 | 1.62E-35 | 4.00E-33 |
| tissue migration (GO:0090130) | 113 | 31 | 13.02 | + | 2.38 | 3.74E-06 | 1.04E-04 |
| cardiac muscle cell development (GO:0055013) | 62 | 17 | 7.15 | + | 2.38 | 4.78E-04 | 7.39E-03 |
| endothelial cell migration (GO:0043542) | 73 | 20 | 8.41 | + | 2.38 | 1.59E-04 | 2.91E-03 |
| regulation of extent of cell growth (GO:0061387) | 95 | 26 | 10.95 | + | 2.37 | 2.67E-05 | 6.17E-04 |
| ear development (GO:0043583) | 223 | 61 | 25.7 | + | 2.37 | 9.86E-11 | 5.60E-09 |
| muscle cell proliferation (GO:0033002) | 44 | 12 | 5.07 | + | 2.37 | 3.29E-03 | 3.68E-02 |
| regulation of cellular response to transforming growth factor beta stimulus (GO:1903844) | 143 | 39 | 16.48 | + | 2.37 | 2.06E-07 | 7.19E-06 |
| multicellular organism growth (GO:0035264) | 88 | 24 | 10.14 | + | 2.37 | 6.02E-05 | 1.27E-03 |

| | | | | | | | |
|---|---|---|---|---|---|---|---|
| positive regulation of double-strand break repair (GO:2000781) | 92 | 25 | 10.6 | + | 2.36 | 4.21E-05 | 9.33E-04 |
| regulation of blood vessel endothelial cell migration (GO:0043535) | 92 | 25 | 10.6 | + | 2.36 | 4.21E-05 | 9.31E-04 |
| diencephalon development (GO:0021536) | 81 | 22 | 9.34 | + | 2.36 | 1.36E-04 | 2.54E-03 |
| mesoderm formation (GO:0001707) | 70 | 19 | 8.07 | + | 2.35 | 2.63E-04 | 4.47E-03 |
| osteoblast differentiation (GO:0001649) | 140 | 38 | 16.14 | + | 2.35 | 3.32E-07 | 1.11E-05 |
| pattern specification process (GO:0007389) | 457 | 124 | 52.67 | + | 2.35 | 2.69E-20 | 3.06E-18 |
| somitogenesis (GO:0001756) | 59 | 16 | 6.8 | + | 2.35 | 7.91E-04 | 1.15E-02 |
| epithelium migration (GO:0090132) | 107 | 29 | 12.33 | + | 2.35 | 9.47E-06 | 2.43E-04 |
| regulation of mRNA processing (GO:0050684) | 133 | 36 | 15.33 | + | 2.35 | 7.31E-07 | 2.33E-05 |
| inner ear development (GO:0048839) | 196 | 53 | 22.59 | + | 2.35 | 2.62E-09 | 1.21E-07 |
| gonad development (GO:0008406) | 226 | 61 | 26.05 | + | 2.34 | 1.39E-10 | 7.81E-09 |
| cellular senescence (GO:0090398) | 63 | 17 | 7.26 | + | 2.34 | 5.85E-04 | 8.95E-03 |
| regulation of axonogenesis (GO:0050770) | 141 | 38 | 16.25 | + | 2.34 | 3.93E-07 | 1.30E-05 |
| Wnt signaling pathway (GO:0016055) | 282 | 76 | 32.5 | + | 2.34 | 6.86E-13 | 5.01E-11 |
| positive regulation of Wnt signaling pathway (GO:0030177) | 141 | 38 | 16.25 | + | 2.34 | 3.93E-07 | 1.30E-05 |
| muscle organ morphogenesis (GO:0048644) | 78 | 21 | 8.99 | + | 2.34 | 2.14E-04 | 3.74E-03 |

| | | | | | | | |
|---|---|---|---|---|---|---|---|
| response to nerve growth factor (GO:1990089) | 52 | 14 | 5.99 | + | 2.34 | 1.78E-03 | 2.26E-02 |
| hair cell differentiation (GO:0035315) | 52 | 14 | 5.99 | + | 2.34 | 1.78E-03 | 2.25E-02 |
| positive regulation of fat cell differentiation (GO:0045600) | 67 | 18 | 7.72 | + | 2.33 | 7.02E-04 | 1.04E-02 |
| positive regulation of epithelial cell proliferation (GO:0050679) | 201 | 54 | 23.17 | + | 2.33 | 2.06E-09 | 9.61E-08 |
| heart development (GO:0007507) | 564 | 151 | 65.01 | + | 2.32 | 4.80E-24 | 6.37E-22 |
| dorsal/ventral pattern formation (GO:0009953) | 86 | 23 | 9.91 | + | 2.32 | 1.06E-04 | 2.07E-03 |
| central nervous system neuron development (GO:0021954) | 86 | 23 | 9.91 | + | 2.32 | 1.06E-04 | 2.07E-03 |
| cellular response to alcohol (GO:0097306) | 101 | 27 | 11.64 | + | 2.32 | 2.41E-05 | 5.68E-04 |
| brain development (GO:0007420) | 730 | 195 | 84.14 | + | 2.32 | 1.26E-30 | 2.29E-28 |
| sex differentiation (GO:0007548) | 281 | 75 | 32.39 | + | 2.32 | 2.22E-12 | 1.50E-10 |
| circulatory system development (GO:0072359) | 926 | 247 | 106.73 | + | 2.31 | 7.69E-39 | 2.58E-36 |
| regulation of canonical Wnt signaling pathway (GO:0060828) | 255 | 68 | 29.39 | + | 2.31 | 1.87E-11 | 1.15E-09 |
| positive regulation of neuron differentiation (GO:0045666) | 90 | 24 | 10.37 | + | 2.31 | 7.51E-05 | 1.53E-03 |
| signal transduction by p53 class mediator (GO:0072331) | 90 | 24 | 10.37 | + | 2.31 | 7.51E-05 | 1.53E-03 |
| embryonic pattern specification (GO:0009880) | 75 | 20 | 8.64 | + | 2.31 | 3.38E-04 | 5.51E-03 |
| cardiac muscle cell differentiation (GO:0055007) | 90 | 24 | 10.37 | + | 2.31 | 7.51E-05 | 1.53E-03 |

| | | | | | | | |
|---|---|---|---|---|---|---|---|
| lamellipodium organization (GO:0097581) | 45 | 12 | 5.19 | + | 2.31 | 4.03E-03 | 4.39E-02 |
| positive regulation of endothelial cell migration (GO:0010595) | 105 | 28 | 12.1 | + | 2.31 | 1.74E-05 | 4.21E-04 |
| negative regulation of cell cycle G1/S phase transition (GO:1902807) | 75 | 20 | 8.64 | + | 2.31 | 3.38E-04 | 5.51E-03 |
| neuron development (GO:0048666) | 859 | 229 | 99.01 | + | 2.31 | 6.91E-36 | 1.74E-33 |
| generation of neurons (GO:0048699) | 1160 | 309 | 133.7 | + | 2.31 | 9.91E-49 | 7.49E-46 |
| striated muscle tissue development (GO:0014706) | 199 | 53 | 22.94 | + | 2.31 | 3.72E-09 | 1.68E-07 |
| response to ionizing radiation (GO:0010212) | 139 | 37 | 16.02 | + | 2.31 | 7.56E-07 | 2.40E-05 |
| pallium development (GO:0021543) | 192 | 51 | 22.13 | + | 2.3 | 8.16E-09 | 3.51E-07 |
| regulation of Ras protein signal transduction (GO:0046578) | 49 | 13 | 5.65 | + | 2.3 | 2.95E-03 | 3.42E-02 |
| neuron differentiation (GO:0030182) | 1082 | 287 | 124.71 | + | 2.3 | 9.52E-45 | 4.96E-42 |
| cellular response to amino acid stimulus (GO:0071230) | 83 | 22 | 9.57 | + | 2.3 | 1.68E-04 | 3.05E-03 |
| organ growth (GO:0035265) | 102 | 27 | 11.76 | + | 2.3 | 2.79E-05 | 6.41E-04 |
| positive regulation of nervous system development (GO:0051962) | 291 | 77 | 33.54 | + | 2.3 | 1.48E-12 | 1.02E-10 |
| cell junction organization (GO:0034330) | 533 | 141 | 61.43 | + | 2.3 | 7.90E-22 | 9.63E-20 |
| regulation of smoothened signaling pathway (GO:0008589) | 87 | 23 | 10.03 | + | 2.29 | 1.20E-04 | 2.28E-03 |
| regulation of muscle cell differentiation (GO:0051147) | 140 | 37 | 16.14 | + | 2.29 | 9.02E-07 | 2.81E-05 |

| | | | | | | | |
|---|---|---|---|---|---|---|---|
| regulation of transforming growth factor beta receptor signaling pathway (GO:0017015) | 140 | 37 | 16.14 | + | 2.29 | 9.02E-07 | 2.80E-05 |
| ovarian follicle development (GO:0001541) | 53 | 14 | 6.11 | + | 2.29 | 2.16E-03 | 2.66E-02 |
| regulation of neurogenesis (GO:0050767) | 371 | 98 | 42.76 | + | 2.29 | 1.49E-15 | 1.29E-13 |
| mesoderm morphogenesis (GO:0048332) | 72 | 19 | 8.3 | + | 2.29 | 5.34E-04 | 8.23E-03 |
| cellular response to acid chemical (GO:0071229) | 91 | 24 | 10.49 | + | 2.29 | 8.58E-05 | 1.73E-03 |
| reproductive structure development (GO:0048608) | 296 | 78 | 34.12 | + | 2.29 | 1.26E-12 | 8.93E-11 |
| neurogenesis (GO:0022008) | 1336 | 352 | 153.98 | + | 2.29 | 1.36E-54 | 2.29E-51 |
| reproductive system development (GO:0061458) | 300 | 79 | 34.58 | + | 2.28 | 9.29E-13 | 6.65E-11 |
| chordate embryonic development (GO:0043009) | 650 | 171 | 74.92 | + | 2.28 | 3.56E-26 | 5.12E-24 |
| negative regulation of cell migration (GO:0030336) | 289 | 76 | 33.31 | + | 2.28 | 2.76E-12 | 1.84E-10 |
| respiratory system development (GO:0060541) | 217 | 57 | 25.01 | + | 2.28 | 1.47E-09 | 7.07E-08 |
| hematopoietic or lymphoid organ development (GO:0048534) | 99 | 26 | 11.41 | + | 2.28 | 4.48E-05 | 9.84E-04 |
| axis specification (GO:0009798) | 99 | 26 | 11.41 | + | 2.28 | 4.48E-05 | 9.82E-04 |
| regulation of dendrite development (GO:0050773) | 99 | 26 | 11.41 | + | 2.28 | 4.48E-05 | 9.81E-04 |
| cell-cell adhesion via plasma-membrane adhesion molecules (GO:0098742) | 263 | 69 | 30.31 | + | 2.28 | 3.38E-11 | 2.03E-09 |
| odontogenesis (GO:0042476) | 122 | 32 | 14.06 | + | 2.28 | 5.84E-06 | 1.55E-04 |

| | | | | | | | |
|---|---|---|---|---|---|---|---|
| regulation of actin filament bundle assembly (GO:0032231) | 103 | 27 | 11.87 | + | 2.27 | 3.25E-05 | 7.34E-04 |
| cell surface receptor protein tyrosine kinase signaling pathway (GO:0007169) | 420 | 110 | 48.41 | + | 2.27 | 5.35E-17 | 5.33E-15 |
| regionalization (GO:0003002) | 409 | 107 | 47.14 | + | 2.27 | 1.64E-16 | 1.55E-14 |
| cell fate specification (GO:0001708) | 88 | 23 | 10.14 | + | 2.27 | 1.37E-04 | 2.56E-03 |
| embryo development ending in birth or egg hatching (GO:0009792) | 671 | 175 | 77.34 | + | 2.26 | 2.83E-26 | 4.15E-24 |
| segmentation (GO:0035282) | 96 | 25 | 11.06 | + | 2.26 | 7.18E-05 | 1.47E-03 |
| regulation of actomyosin structure organization (GO:0110020) | 100 | 26 | 11.53 | + | 2.26 | 5.22E-05 | 1.13E-03 |
| positive regulation of epithelial cell migration (GO:0010634) | 150 | 39 | 17.29 | + | 2.26 | 9.03E-07 | 2.80E-05 |
| endocrine system development (GO:0035270) | 127 | 33 | 14.64 | + | 2.25 | 7.90E-06 | 2.05E-04 |
| positive regulation of axonogenesis (GO:0050772) | 77 | 20 | 8.87 | + | 2.25 | 4.26E-04 | 6.77E-03 |
| neural precursor cell proliferation (GO:0061351) | 104 | 27 | 11.99 | + | 2.25 | 3.82E-05 | 8.51E-04 |
| central nervous system development (GO:0007417) | 1000 | 259 | 115.26 | + | 2.25 | 2.50E-38 | 7.56E-36 |
| animal organ morphogenesis (GO:0009887) | 977 | 253 | 112.61 | + | 2.25 | 2.19E-37 | 5.93E-35 |
| regulation of heart growth (GO:0060420) | 58 | 15 | 6.68 | + | 2.24 | 2.73E-03 | 3.21E-02 |
| negative regulation of BMP signaling pathway (GO:0030514) | 58 | 15 | 6.68 | + | 2.24 | 2.73E-03 | 3.20E-02 |
| eye development (GO:0001654) | 383 | 99 | 44.14 | + | 2.24 | 4.83E-15 | 3.99E-13 |

| | | | | | | | |
|---|---|---|---|---|---|---|---|
| rhythmic process (GO:0048511) | 271 | 70 | 31.23 | + | 2.24 | 6.94E-11 | 4.01E-09 |
| growth (GO:0040007) | 426 | 110 | 49.1 | + | 2.24 | 1.92E-16 | 1.79E-14 |
| developmental growth (GO:0048589) | 426 | 110 | 49.1 | + | 2.24 | 1.92E-16 | 1.78E-14 |
| gliogenesis (GO:0042063) | 275 | 71 | 31.7 | + | 2.24 | 4.90E-11 | 2.89E-09 |
| positive regulation of protein localization to nucleus (GO:1900182) | 93 | 24 | 10.72 | + | 2.24 | 1.15E-04 | 2.21E-03 |
| negative regulation of transforming growth factor beta receptor signaling pathway (GO:0030512) | 93 | 24 | 10.72 | + | 2.24 | 1.15E-04 | 2.20E-03 |
| sensory system development (GO:0048880) | 393 | 101 | 45.3 | + | 2.23 | 3.44E-15 | 2.89E-13 |
| negative regulation of cell motility (GO:2000146) | 304 | 78 | 35.04 | + | 2.23 | 5.58E-12 | 3.59E-10 |
| cell junction assembly (GO:0034329) | 285 | 73 | 32.85 | + | 2.22 | 3.13E-11 | 1.88E-09 |
| camera-type eye development (GO:0043010) | 336 | 86 | 38.73 | + | 2.22 | 6.34E-13 | 4.68E-11 |
| visual system development (GO:0150063) | 387 | 99 | 44.6 | + | 2.22 | 1.35E-14 | 1.09E-12 |
| regulation of stress fiber assembly (GO:0051492) | 90 | 23 | 10.37 | + | 2.22 | 1.84E-04 | 3.28E-03 |
| protein localization to chromosome (GO:0034502) | 90 | 23 | 10.37 | + | 2.22 | 1.84E-04 | 3.28E-03 |
| negative regulation of transcription by RNA polymerase II (GO:0000122) | 920 | 235 | 106.04 | + | 2.22 | 1.15E-33 | 2.53E-31 |
| negative regulation of cell differentiation (GO:0045596) | 647 | 165 | 74.57 | + | 2.21 | 1.01E-23 | 1.31E-21 |
| negative regulation of locomotion (GO:0040013) | 322 | 82 | 37.11 | + | 2.21 | 2.98E-12 | 1.95E-10 |

| | | | | | | | |
|---|---|---|---|---|---|---|---|
| regulation of epithelial cell migration (GO:0010632) | 228 | 58 | 26.28 | + | 2.21 | 4.01E-09 | 1.78E-07 |
| regulation of protein import into nucleus (GO:0042306) | 59 | 15 | 6.8 | + | 2.21 | 3.03E-03 | 3.51E-02 |
| regulation of neuron projection development (GO:0010975) | 437 | 111 | 50.37 | + | 2.2 | 5.09E-16 | 4.64E-14 |
| collagen metabolic process (GO:0032963) | 63 | 16 | 7.26 | + | 2.2 | 2.13E-03 | 2.62E-02 |
| cellular response to retinoic acid (GO:0071300) | 67 | 17 | 7.72 | + | 2.2 | 1.51E-03 | 1.99E-02 |
| regulation of insulin receptor signaling pathway (GO:0046626) | 67 | 17 | 7.72 | + | 2.2 | 1.51E-03 | 1.99E-02 |
| regulation of Wnt signaling pathway (GO:0030111) | 328 | 83 | 37.8 | + | 2.2 | 2.71E-12 | 1.81E-10 |
| hair cycle process (GO:0022405) | 87 | 22 | 10.03 | + | 2.19 | 2.95E-04 | 4.90E-03 |
| molting cycle process (GO:0022404) | 87 | 22 | 10.03 | + | 2.19 | 2.95E-04 | 4.89E-03 |
| regulation of myeloid cell differentiation (GO:0045637) | 198 | 50 | 22.82 | + | 2.19 | 6.50E-08 | 2.50E-06 |
| cell adhesion (GO:0007155) | 959 | 242 | 110.53 | + | 2.19 | 9.61E-34 | 2.17E-31 |
| embryo development (GO:0009790) | 1043 | 263 | 120.21 | + | 2.19 | 1.02E-36 | 2.65E-34 |
| cellular response to hypoxia (GO:0071456) | 127 | 32 | 14.64 | + | 2.19 | 1.66E-05 | 4.03E-04 |
| negative regulation of cell development (GO:0010721) | 262 | 66 | 30.2 | + | 2.19 | 5.64E-10 | 2.91E-08 |
| regulation of endothelial cell migration (GO:0010594) | 167 | 42 | 19.25 | + | 2.18 | 1.02E-06 | 3.11E-05 |
| positive regulation of cell projection organization (GO:0031346) | 354 | 89 | 40.8 | + | 2.18 | 7.54E-13 | 5.46E-11 |

| | | | | | | | |
|---|---|---|---|---|---|---|---|
| epithelial cell development (GO:0002064) | 191 | 48 | 22.01 | + | 2.18 | 1.40E-07 | 5.06E-06 |
| positive regulation of endothelial cell proliferation (GO:0001938) | 92 | 23 | 10.6 | + | 2.17 | 4.02E-04 | 6.41E-03 |
| regulation of bone mineralization (GO:0030500) | 80 | 20 | 9.22 | + | 2.17 | 6.48E-04 | 9.60E-03 |
| retina development in camera-type eye (GO:0060041) | 156 | 39 | 17.98 | + | 2.17 | 3.10E-06 | 8.71E-05 |
| RNA export from nucleus (GO:0006405) | 84 | 21 | 9.68 | + | 2.17 | 4.71E-04 | 7.31E-03 |
| positive regulation of smooth muscle cell proliferation (GO:0048661) | 84 | 21 | 9.68 | + | 2.17 | 4.71E-04 | 7.30E-03 |
| regulation of cell migration (GO:0030334) | 941 | 234 | 108.46 | + | 2.16 | 1.18E-31 | 2.22E-29 |
| regulation of epithelial cell differentiation (GO:0030856) | 153 | 38 | 17.63 | + | 2.15 | 4.72E-06 | 1.28E-04 |
| regulation of smooth muscle cell proliferation (GO:0048660) | 137 | 34 | 15.79 | + | 2.15 | 1.15E-05 | 2.89E-04 |
| memory (GO:0007613) | 125 | 31 | 14.41 | + | 2.15 | 2.95E-05 | 6.75E-04 |
| regulation of Notch signaling pathway (GO:0008593) | 97 | 24 | 11.18 | + | 2.15 | 3.02E-04 | 5.01E-03 |
| cellular response to carbohydrate stimulus (GO:0071322) | 93 | 23 | 10.72 | + | 2.15 | 4.29E-04 | 6.82E-03 |
| post-embryonic development (GO:0009791) | 89 | 22 | 10.26 | + | 2.14 | 6.12E-04 | 9.29E-03 |
| regulation of fibroblast proliferation (GO:0048145) | 89 | 22 | 10.26 | + | 2.14 | 6.12E-04 | 9.28E-03 |
| regulation of viral genome replication (GO:0045069) | 85 | 21 | 9.8 | + | 2.14 | 8.77E-04 | 1.25E-02 |
| adult locomotory behavior (GO:0008344) | 81 | 20 | 9.34 | + | 2.14 | 7.52E-04 | 1.10E-02 |

| | | | | | | | |
|---|---|---|---|---|---|---|---|
| regulation of myoblast differentiation (GO:0045661) | 73 | 18 | 8.41 | + | 2.14 | 1.42E-03 | 1.88E-02 |
| regulation of steroid biosynthetic process (GO:0050810) | 73 | 18 | 8.41 | + | 2.14 | 1.42E-03 | 1.88E-02 |
| regulation of double-strand break repair (GO:2000779) | 142 | 35 | 16.37 | + | 2.14 | 1.43E-05 | 3.54E-04 |
| cellular response to decreased oxygen levels (GO:0036294) | 134 | 33 | 15.44 | + | 2.14 | 1.81E-05 | 4.34E-04 |
| positive regulation of cell differentiation (GO:0045597) | 865 | 213 | 99.7 | + | 2.14 | 3.66E-28 | 5.89E-26 |
| peptidyl-lysine modification (GO:0018205) | 130 | 32 | 14.98 | + | 2.14 | 2.47E-05 | 5.78E-04 |
| cellular response to hydrogen peroxide (GO:0070301) | 65 | 16 | 7.49 | + | 2.14 | 2.74E-03 | 3.21E-02 |
| negative regulation of cell projection organization (GO:0031345) | 179 | 44 | 20.63 | + | 2.13 | 8.42E-07 | 2.64E-05 |
| learning (GO:0007612) | 155 | 38 | 17.86 | + | 2.13 | 5.69E-06 | 1.51E-04 |
| positive regulation of myeloid cell differentiation (GO:0045639) | 102 | 25 | 11.76 | + | 2.13 | 2.36E-04 | 4.07E-03 |
| negative regulation of transmembrane receptor protein serine/threonine kinase signaling pathway (GO:0090101) | 147 | 36 | 16.94 | + | 2.12 | 1.10E-05 | 2.79E-04 |
| regulation of animal organ morphogenesis (GO:2000027) | 90 | 22 | 10.37 | + | 2.12 | 6.57E-04 | 9.73E-03 |
| negative regulation of developmental process (GO:0051093) | 896 | 219 | 103.27 | + | 2.12 | 1.60E-28 | 2.63E-26 |
| regulation of small GTPase mediated signal transduction (GO:0051056) | 303 | 74 | 34.92 | + | 2.12 | 2.64E-10 | 1.43E-08 |
| cerebral cortex development (GO:0021987) | 127 | 31 | 14.64 | + | 2.12 | 3.89E-05 | 8.64E-04 |
| cellular response to glucose stimulus (GO:0071333) | 78 | 19 | 8.99 | + | 2.11 | 1.20E-03 | 1.63E-02 |

| | | | | | | | |
|---|---|---|---|---|---|---|---|
| regulation of axon extension (GO:0030516) | 78 | 19 | 8.99 | + | 2.11 | 1.20E-03 | 1.62E-02 |
| glial cell differentiation (GO:0010001) | 218 | 53 | 25.13 | + | 2.11 | 9.65E-08 | 3.57E-06 |
| positive regulation of developmental process (GO:0051094) | 1323 | 321 | 152.49 | + | 2.11 | 2.35E-41 | 1.08E-38 |
| astrocyte differentiation (GO:0048708) | 66 | 16 | 7.61 | + | 2.1 | 3.14E-03 | 3.55E-02 |
| hindbrain development (GO:0030902) | 157 | 38 | 18.1 | + | 2.1 | 7.17E-06 | 1.86E-04 |
| extrinsic apoptotic signaling pathway (GO:0097191) | 112 | 27 | 12.91 | + | 2.09 | 1.56E-04 | 2.86E-03 |
| endothelial cell differentiation (GO:0045446) | 83 | 20 | 9.57 | + | 2.09 | 1.43E-03 | 1.88E-02 |
| regulation of epithelial cell proliferation (GO:0050678) | 357 | 86 | 41.15 | + | 2.09 | 2.38E-11 | 1.45E-09 |
| negative regulation of DNA-templated transcription (GO:0045892) | 1259 | 303 | 145.11 | + | 2.09 | 3.09E-38 | 8.99E-36 |
| gland development (GO:0048732) | 424 | 102 | 48.87 | + | 2.09 | 2.65E-13 | 1.97E-11 |
| muscle structure development (GO:0061061) | 516 | 124 | 59.47 | + | 2.08 | 7.00E-16 | 6.22E-14 |
| nervous system development (GO:0007399) | 2193 | 527 | 252.76 | + | 2.08 | 5.03E-69 | 1.52E-65 |
| negative regulation of RNA biosynthetic process (GO:1902679) | 1274 | 306 | 146.84 | + | 2.08 | 2.18E-38 | 6.73E-36 |
| positive regulation of transcription by RNA polymerase II (GO:0045944) | 1245 | 299 | 143.5 | + | 2.08 | 1.55E-37 | 4.26E-35 |
| negative regulation of growth (GO:0045926) | 225 | 54 | 25.93 | + | 2.08 | 1.38E-07 | 4.99E-06 |
| sensory organ development (GO:0007423) | 584 | 140 | 67.31 | + | 2.08 | 1.25E-17 | 1.29E-15 |

| | | | | | | | |
|---|---|---|---|---|---|---|---|
| positive regulation of cell migration (GO:0030335) | 547 | 131 | 63.05 | + | 2.08 | 1.43E-16 | 1.36E-14 |
| response to decreased oxygen levels (GO:0036293) | 301 | 72 | 34.69 | + | 2.08 | 1.43E-09 | 6.91E-08 |
| mRNA export from nucleus (GO:0006406) | 67 | 16 | 7.72 | + | 2.07 | 3.60E-03 | 3.99E-02 |
| regulation of developmental growth (GO:0048638) | 302 | 72 | 34.81 | + | 2.07 | 1.55E-09 | 7.39E-08 |
| negative regulation of blood vessel morphogenesis (GO:2000181) | 105 | 25 | 12.1 | + | 2.07 | 3.34E-04 | 5.46E-03 |
| regulation of cell differentiation (GO:0045595) | 1534 | 365 | 176.81 | + | 2.06 | 3.88E-45 | 2.09E-42 |
| regulation of protein localization to nucleus (GO:1900180) | 143 | 34 | 16.48 | + | 2.06 | 3.23E-05 | 7.30E-04 |
| regulation of cell motility (GO:2000145) | 1001 | 238 | 115.37 | + | 2.06 | 5.82E-29 | 9.99E-27 |
| regulation of ossification (GO:0030278) | 122 | 29 | 14.06 | + | 2.06 | 1.50E-04 | 2.77E-03 |
| response to alkaloid (GO:0043279) | 101 | 24 | 11.64 | + | 2.06 | 4.60E-04 | 7.22E-03 |
| response to glucose (GO:0009749) | 135 | 32 | 15.56 | + | 2.06 | 6.12E-05 | 1.29E-03 |
| response to hypoxia (GO:0001666) | 287 | 68 | 33.08 | + | 2.06 | 6.26E-09 | 2.72E-07 |
| anatomical structure formation involved in morphogenesis (GO:0048646) | 969 | 229 | 111.68 | + | 2.05 | 1.77E-27 | 2.74E-25 |
| regulation of cellular response to insulin stimulus (GO:1900076) | 72 | 17 | 8.3 | + | 2.05 | 4.39E-03 | 4.73E-02 |
| smoothened signaling pathway (GO:0007224) | 89 | 21 | 10.26 | + | 2.05 | 1.22E-03 | 1.65E-02 |
| regulation of signal transduction by p53 class mediator (GO:1901796) | 106 | 25 | 12.22 | + | 2.05 | 3.80E-04 | 6.07E-03 |

| | | | | | | | |
|---|---|---|---|---|---|---|---|
| negative regulation of vasculature development (GO:1901343) | 106 | 25 | 12.22 | + | 2.05 | 3.80E-04 | 6.07E-03 |
| regulation of multicellular organismal development (GO:2000026) | 1384 | 326 | 159.52 | + | 2.04 | 4.93E-39 | 1.73E-36 |
| negative regulation of RNA metabolic process (GO:0051253) | 1375 | 323 | 158.48 | + | 2.04 | 1.75E-38 | 5.52E-36 |
| cellular response to oxygen levels (GO:0071453) | 149 | 35 | 17.17 | + | 2.04 | 4.40E-05 | 9.68E-04 |
| establishment of cell polarity (GO:0030010) | 132 | 31 | 15.21 | + | 2.04 | 9.39E-05 | 1.88E-03 |
| cell-cell junction organization (GO:0045216) | 179 | 42 | 20.63 | + | 2.04 | 5.49E-06 | 1.48E-04 |
| cellular response to hexose stimulus (GO:0071331) | 81 | 19 | 9.34 | + | 2.04 | 2.38E-03 | 2.84E-02 |
| negative regulation of translation (GO:0017148) | 128 | 30 | 14.75 | + | 2.03 | 1.29E-04 | 2.42E-03 |
| regulation of locomotion (GO:0040012) | 1042 | 244 | 120.1 | + | 2.03 | 1.31E-28 | 2.18E-26 |
| anatomical structure morphogenesis (GO:0009653) | 2220 | 519 | 255.87 | + | 2.03 | 1.36E-63 | 3.44E-60 |
| positive regulation of proteasomal ubiquitin-dependent protein catabolic process (GO:0032436) | 77 | 18 | 8.87 | + | 2.03 | 3.34E-03 | 3.73E-02 |
| positive regulation of DNA recombination (GO:0045911) | 77 | 18 | 8.87 | + | 2.03 | 3.34E-03 | 3.72E-02 |
| erythrocyte differentiation (GO:0030218) | 103 | 24 | 11.87 | + | 2.02 | 8.81E-04 | 1.26E-02 |
| cell migration (GO:0016477) | 915 | 213 | 105.46 | + | 2.02 | 9.42E-25 | 1.27E-22 |
| positive regulation of developmental growth (GO:0048639) | 159 | 37 | 18.33 | + | 2.02 | 2.66E-05 | 6.15E-04 |
| embryonic placenta development (GO:0001892) | 86 | 20 | 9.91 | + | 2.02 | 1.88E-03 | 2.36E-02 |

| | | | | | | | |
|---|---|---|---|---|---|---|---|
| response to oxygen levels (GO:0070482) | 327 | 76 | 37.69 | + | 2.02 | 1.55E-09 | 7.40E-08 |
| positive regulation of neurogenesis (GO:0050769) | 241 | 56 | 27.78 | + | 2.02 | 2.34E-07 | 8.03E-06 |
| regulation of plasma membrane bounded cell projection organization (GO:0120035) | 633 | 147 | 72.96 | + | 2.01 | 3.12E-17 | 3.12E-15 |
| positive regulation of cell motility (GO:2000147) | 573 | 133 | 66.04 | + | 2.01 | 1.32E-15 | 1.16E-13 |
| learning or memory (GO:0007611) | 276 | 64 | 31.81 | + | 2.01 | 3.53E-08 | 1.41E-06 |
| regulation of double-strand break repair via homologous recombination (GO:0010569) | 82 | 19 | 9.45 | + | 2.01 | 2.61E-03 | 3.08E-02 |
| positive regulation of locomotion (GO:0040017) | 587 | 136 | 67.66 | + | 2.01 | 6.20E-16 | 5.58E-14 |
| response to hydrogen peroxide (GO:0042542) | 95 | 22 | 10.95 | + | 2.01 | 1.11E-03 | 1.52E-02 |
| response to amino acid (GO:0043200) | 121 | 28 | 13.95 | + | 2.01 | 2.72E-04 | 4.62E-03 |
| regulation of G1/S transition of mitotic cell cycle (GO:2000045) | 160 | 37 | 18.44 | + | 2.01 | 2.93E-05 | 6.72E-04 |
| glucose homeostasis (GO:0042593) | 199 | 46 | 22.94 | + | 2.01 | 3.21E-06 | 9.01E-05 |
| negative regulation of cellular response to growth factor stimulus (GO:0090288) | 104 | 24 | 11.99 | + | 2 | 9.35E-04 | 1.32E-02 |
| negative regulation of angiogenesis (GO:0016525) | 104 | 24 | 11.99 | + | 2 | 9.35E-04 | 1.32E-02 |
| muscle tissue development (GO:0060537) | 347 | 80 | 39.99 | + | 2 | 8.90E-10 | 4.47E-08 |
| carbohydrate homeostasis (GO:0033500) | 200 | 46 | 23.05 | + | 2 | 3.58E-06 | 9.98E-05 |
| positive regulation of DNA repair (GO:0045739) | 135 | 31 | 15.56 | + | 1.99 | 1.90E-04 | 3.37E-03 |

| | | | | | | | |
|---|---|---|---|---|---|---|---|
| response to acid chemical (GO:0001101) | 135 | 31 | 15.56 | + | 1.99 | 1.90E-04 | 3.36E-03 |
| positive regulation of RNA biosynthetic process (GO:1902680) | 1681 | 386 | 193.75 | + | 1.99 | 5.00E-44 | 2.52E-41 |
| cell-cell adhesion (GO:0098609) | 549 | 126 | 63.28 | + | 1.99 | 1.77E-14 | 1.43E-12 |
| positive regulation of DNA-templated transcription (GO:0045893) | 1678 | 385 | 193.4 | + | 1.99 | 1.17E-43 | 5.69E-41 |
| negative regulation of cell growth (GO:0030308) | 170 | 39 | 19.59 | + | 1.99 | 2.81E-05 | 6.45E-04 |
| chromatin organization (GO:0006325) | 786 | 180 | 90.59 | + | 1.99 | 4.33E-20 | 4.89E-18 |
| hair follicle development (GO:0001942) | 83 | 19 | 9.57 | + | 1.99 | 2.90E-03 | 3.38E-02 |
| cellular response to monosaccharide stimulus (GO:0071326) | 83 | 19 | 9.57 | + | 1.99 | 2.90E-03 | 3.38E-02 |
| regulation of gliogenesis (GO:0014013) | 105 | 24 | 12.1 | + | 1.98 | 1.00E-03 | 1.41E-02 |
| regulation of telomere maintenance (GO:0032204) | 105 | 24 | 12.1 | + | 1.98 | 1.00E-03 | 1.41E-02 |
| regulation of postsynapse organization (GO:0099175) | 105 | 24 | 12.1 | + | 1.98 | 1.00E-03 | 1.40E-02 |
| response to hexose (GO:0009746) | 140 | 32 | 16.14 | + | 1.98 | 1.46E-04 | 2.70E-03 |
| regulation of cell development (GO:0060284) | 815 | 186 | 93.93 | + | 1.98 | 1.35E-20 | 1.56E-18 |
| placenta development (GO:0001890) | 149 | 34 | 17.17 | + | 1.98 | 8.36E-05 | 1.69E-03 |
| positive regulation of RNA metabolic process (GO:0051254) | 1806 | 412 | 208.16 | + | 1.98 | 2.12E-46 | 1.34E-43 |
| regulation of biomineral tissue development (GO:0070167) | 101 | 23 | 11.64 | + | 1.98 | 1.40E-03 | 1.85E-02 |

| | | | | | | | |
|---|---|---|---|---|---|---|---|
| endothelium development (GO:0003158) | 101 | 23 | 11.64 | + | 1.98 | 1.40E-03 | 1.85E-02 |
| response to mechanical stimulus (GO:0009612) | 220 | 50 | 25.36 | + | 1.97 | 2.05E-06 | 5.93E-05 |
| regulation of Rho protein signal transduction (GO:0035023) | 88 | 20 | 10.14 | + | 1.97 | 2.35E-03 | 2.81E-02 |
| regulation of cell adhesion (GO:0030155) | 785 | 178 | 90.48 | + | 1.97 | 2.18E-19 | 2.45E-17 |
| regulation of cell projection organization (GO:0031344) | 649 | 147 | 74.8 | + | 1.97 | 3.74E-16 | 3.45E-14 |
| response to retinoic acid (GO:0032526) | 106 | 24 | 12.22 | + | 1.96 | 1.09E-03 | 1.50E-02 |
| myeloid cell differentiation (GO:0030099) | 296 | 67 | 34.12 | + | 1.96 | 5.95E-08 | 2.29E-06 |
| peripheral nervous system development (GO:0007422) | 84 | 19 | 9.68 | + | 1.96 | 3.23E-03 | 3.62E-02 |
| regulation of developmental process (GO:0050793) | 2409 | 543 | 277.66 | + | 1.96 | 5.30E-61 | 1.15E-57 |
| regulation of cold-induced thermogenesis (GO:0120161) | 151 | 34 | 17.4 | + | 1.95 | 1.03E-04 | 2.03E-03 |
| negative regulation of nucleobase-containing compound metabolic process (GO:0045934) | 1501 | 337 | 173 | + | 1.95 | 5.04E-36 | 1.29E-33 |
| regulation of cell morphogenesis (GO:0022604) | 245 | 55 | 28.24 | + | 1.95 | 9.49E-07 | 2.92E-05 |
| Notch signaling pathway (GO:0007219) | 116 | 26 | 13.37 | + | 1.94 | 7.03E-04 | 1.04E-02 |
| regulation of cell cycle G1/S phase transition (GO:1902806) | 183 | 41 | 21.09 | + | 1.94 | 3.36E-05 | 7.55E-04 |
| erythrocyte homeostasis (GO:0034101) | 112 | 25 | 12.91 | + | 1.94 | 9.60E-04 | 1.35E-02 |
| regulation of angiogenesis (GO:0045765) | 288 | 64 | 33.19 | + | 1.93 | 2.02E-07 | 7.10E-06 |

| | | | | | | | |
|---|---|---|---|---|---|---|---|
| response to carbohydrate (GO:0009743) | 171 | 38 | 19.71 | + | 1.93 | 5.76E-05 | 1.23E-03 |
| cellular response to UV (GO:0034644) | 90 | 20 | 10.37 | + | 1.93 | 4.07E-03 | 4.44E-02 |
| plasma membrane bounded cell projection organization (GO:0120036) | 1148 | 255 | 132.32 | + | 1.93 | 2.85E-26 | 4.14E-24 |
| regulation of vasculature development (GO:1901342) | 294 | 65 | 33.89 | + | 1.92 | 1.79E-07 | 6.32E-06 |
| negative regulation of extrinsic apoptotic signaling pathway (GO:2001237) | 95 | 21 | 10.95 | + | 1.92 | 3.14E-03 | 3.55E-02 |
| regulation of anatomical structure morphogenesis (GO:0022603) | 828 | 183 | 95.43 | + | 1.92 | 1.18E-18 | 1.31E-16 |
| positive regulation of nucleobase-containing compound metabolic process (GO:0045935) | 2009 | 444 | 231.55 | + | 1.92 | 2.89E-46 | 1.75E-43 |
| positive regulation of phosphatidylinositol 3-kinase/protein kinase B signal transduction (GO:0051897) | 181 | 40 | 20.86 | + | 1.92 | 5.17E-05 | 1.12E-03 |
| positive regulation of angiogenesis (GO:0045766) | 163 | 36 | 18.79 | + | 1.92 | 1.07E-04 | 2.08E-03 |
| positive regulation of cellular component biogenesis (GO:0044089) | 512 | 113 | 59.01 | + | 1.91 | 7.73E-12 | 4.91E-10 |
| signal transduction in response to DNA damage (GO:0042770) | 145 | 32 | 16.71 | + | 1.91 | 3.30E-04 | 5.40E-03 |
| response to estradiol (GO:0032355) | 118 | 26 | 13.6 | + | 1.91 | 1.18E-03 | 1.61E-02 |
| import into nucleus (GO:0051170) | 118 | 26 | 13.6 | + | 1.91 | 1.18E-03 | 1.61E-02 |
| tissue remodeling (GO:0048771) | 109 | 24 | 12.56 | + | 1.91 | 2.18E-03 | 2.64E-02 |
| postsynapse organization (GO:0099173) | 109 | 24 | 12.56 | + | 1.91 | 2.18E-03 | 2.63E-02 |
| hair cycle (GO:0042633) | 100 | 22 | 11.53 | + | 1.91 | 2.47E-03 | 2.93E-02 |

| | | | | | | | |
|---|---|---|---|---|---|---|---|
| molting cycle (GO:0042303) | 100 | 22 | 11.53 | + | 1.91 | 2.47E-03 | 2.92E-02 |
| system development (GO:0048731) | 3527 | 773 | 406.51 | + | 1.9 | 3.20E-86 | 2.42E-82 |
| cellular response to nutrient levels (GO:0031669) | 233 | 51 | 26.86 | + | 1.9 | 6.39E-06 | 1.69E-04 |
| regulation of DNA repair (GO:0006282) | 224 | 49 | 25.82 | + | 1.9 | 1.15E-05 | 2.89E-04 |
| epithelium development (GO:0060429) | 1093 | 239 | 125.98 | + | 1.9 | 1.05E-23 | 1.35E-21 |
| cellular response to endogenous stimulus (GO:0071495) | 1112 | 243 | 128.17 | + | 1.9 | 5.04E-24 | 6.62E-22 |
| muscle cell differentiation (GO:0042692) | 284 | 62 | 32.73 | + | 1.89 | 7.52E-07 | 2.39E-05 |
| muscle organ development (GO:0007517) | 312 | 68 | 35.96 | + | 1.89 | 2.26E-07 | 7.78E-06 |
| intracellular glucose homeostasis (GO:0001678) | 101 | 22 | 11.64 | + | 1.89 | 2.71E-03 | 3.18E-02 |
| positive regulation of cold-induced thermogenesis (GO:0120162) | 101 | 22 | 11.64 | + | 1.89 | 2.71E-03 | 3.18E-02 |
| regulation of myeloid leukocyte differentiation (GO:0002761) | 124 | 27 | 14.29 | + | 1.89 | 9.94E-04 | 1.39E-02 |
| response to monosaccharide (GO:0034284) | 147 | 32 | 16.94 | + | 1.89 | 3.72E-04 | 5.99E-03 |
| cell projection organization (GO:0030030) | 1196 | 260 | 137.85 | + | 1.89 | 2.83E-25 | 3.92E-23 |
| positive regulation of vasculature development (GO:1904018) | 166 | 36 | 19.13 | + | 1.88 | 1.95E-04 | 3.45E-03 |
| double-strand break repair via homologous recombination (GO:0000724) | 120 | 26 | 13.83 | + | 1.88 | 1.36E-03 | 1.81E-02 |
| recombinational repair (GO:0000725) | 125 | 27 | 14.41 | + | 1.87 | 1.08E-03 | 1.49E-02 |

| | | | | | | | |
|---|---|---|---|---|---|---|---|
| positive regulation of cell adhesion (GO:0045785) | 482 | 104 | 55.55 | + | 1.87 | 1.85E-10 | 1.01E-08 |
| cognition (GO:0050890) | 320 | 69 | 36.88 | + | 1.87 | 2.17E-07 | 7.57E-06 |
| regulation of chromosome organization (GO:0033044) | 246 | 53 | 28.35 | + | 1.87 | 7.09E-06 | 1.85E-04 |
| protein-DNA complex organization (GO:0071824) | 878 | 189 | 101.2 | + | 1.87 | 4.68E-18 | 4.95E-16 |
| modulation of chemical synaptic transmission (GO:0050804) | 479 | 103 | 55.21 | + | 1.87 | 2.69E-10 | 1.45E-08 |
| regulation of postsynaptic membrane potential (GO:0060078) | 107 | 23 | 12.33 | + | 1.86 | 3.32E-03 | 3.71E-02 |
| neuron apoptotic process (GO:0051402) | 107 | 23 | 12.33 | + | 1.86 | 3.32E-03 | 3.71E-02 |
| mRNA transport (GO:0051028) | 135 | 29 | 15.56 | + | 1.86 | 9.62E-04 | 1.35E-02 |
| regulation of trans-synaptic signaling (GO:0099177) | 480 | 103 | 55.32 | + | 1.86 | 2.92E-10 | 1.56E-08 |
| DNA damage checkpoint signaling (GO:0000077) | 112 | 24 | 12.91 | + | 1.86 | 2.56E-03 | 3.02E-02 |
| tissue development (GO:0009888) | 1748 | 374 | 201.47 | + | 1.86 | 2.44E-35 | 5.96E-33 |
| multicellular organism development (GO:0007275) | 3945 | 843 | 454.69 | + | 1.85 | 8.72E-90 | 1.32E-85 |
| regulation of neuron apoptotic process (GO:0043523) | 239 | 51 | 27.55 | + | 1.85 | 1.39E-05 | 3.44E-04 |
| in utero embryonic development (GO:0001701) | 394 | 84 | 45.41 | + | 1.85 | 2.43E-08 | 9.94E-07 |
| protein import into nucleus (GO:0006606) | 113 | 24 | 13.02 | + | 1.84 | 2.75E-03 | 3.22E-02 |
| mitotic cell cycle phase transition (GO:0044772) | 146 | 31 | 16.83 | + | 1.84 | 6.41E-04 | 9.61E-03 |

| | | | | | | | |
|---|---|---|---|---|---|---|---|
| chromatin remodeling (GO:0006338) | 637 | 135 | 73.42 | + | 1.84 | 1.30E-12 | 9.13E-11 |
| cell motility (GO:0048870) | 1098 | 232 | 126.55 | + | 1.83 | 7.73E-21 | 9.06E-19 |
| regeneration (GO:0031099) | 161 | 34 | 18.56 | + | 1.83 | 4.36E-04 | 6.88E-03 |
| positive regulation of cell development (GO:0010720) | 451 | 95 | 51.98 | + | 1.83 | 3.98E-09 | 1.78E-07 |
| regulation of endothelial cell proliferation (GO:0001936) | 133 | 28 | 15.33 | + | 1.83 | 1.49E-03 | 1.96E-02 |
| negative regulation of mitotic cell cycle (GO:0045930) | 214 | 45 | 24.67 | + | 1.82 | 5.61E-05 | 1.20E-03 |
| skin epidermis development (GO:0098773) | 119 | 25 | 13.72 | + | 1.82 | 3.44E-03 | 3.82E-02 |
| regulation of nervous system process (GO:0031644) | 119 | 25 | 13.72 | + | 1.82 | 3.44E-03 | 3.82E-02 |
| nucleus organization (GO:0006997) | 143 | 30 | 16.48 | + | 1.82 | 1.36E-03 | 1.82E-02 |
| cellular response to radiation (GO:0071478) | 172 | 36 | 19.82 | + | 1.82 | 4.16E-04 | 6.62E-03 |
| animal organ development (GO:0048513) | 2843 | 595 | 327.68 | + | 1.82 | 1.89E-55 | 3.58E-52 |
| myeloid leukocyte differentiation (GO:0002573) | 153 | 32 | 17.63 | + | 1.81 | 8.04E-04 | 1.16E-02 |
| positive regulation of intracellular transport (GO:0032388) | 139 | 29 | 16.02 | + | 1.81 | 1.27E-03 | 1.72E-02 |
| regulation of cell shape (GO:0008360) | 139 | 29 | 16.02 | + | 1.81 | 1.27E-03 | 1.72E-02 |
| locomotory behavior (GO:0007626) | 197 | 41 | 22.71 | + | 1.81 | 1.75E-04 | 3.15E-03 |
| regulation of cellular response to stress (GO:0080135) | 516 | 107 | 59.47 | + | 1.8 | 1.19E-09 | 5.80E-08 |

| | | | | | | | |
|---|---|---|---|---|---|---|---|
| negative regulation of cell population proliferation (GO:0008285) | 700 | 145 | 80.68 | + | 1.8 | 1.48E-12 | 1.02E-10 |
| myeloid cell homeostasis (GO:0002262) | 140 | 29 | 16.14 | + | 1.8 | 1.93E-03 | 2.42E-02 |
| regulation of transcription by RNA polymerase II (GO:0006357) | 2584 | 535 | 297.83 | + | 1.8 | 1.62E-47 | 1.17E-44 |
| regulation of cell size (GO:0008361) | 174 | 36 | 20.05 | + | 1.8 | 4.62E-04 | 7.24E-03 |
| cell-cell junction assembly (GO:0007043) | 126 | 26 | 14.52 | + | 1.79 | 2.93E-03 | 3.40E-02 |
| small GTPase-mediated signal transduction (GO:0007264) | 262 | 54 | 30.2 | + | 1.79 | 2.21E-05 | 5.26E-04 |
| regulation of chromosome segregation (GO:0051983) | 131 | 27 | 15.1 | + | 1.79 | 2.32E-03 | 2.78E-02 |
| cellular response to reactive oxygen species (GO:0034614) | 131 | 27 | 15.1 | + | 1.79 | 2.32E-03 | 2.78E-02 |
| negative regulation of mitotic cell cycle phase transition (GO:1901991) | 165 | 34 | 19.02 | + | 1.79 | 8.02E-04 | 1.16E-02 |
| regulation of synaptic plasticity (GO:0048167) | 204 | 42 | 23.51 | + | 1.79 | 1.58E-04 | 2.89E-03 |
| cell development (GO:0048468) | 2226 | 458 | 256.56 | + | 1.79 | 2.85E-39 | 1.05E-36 |
| protein localization to nucleus (GO:0034504) | 190 | 39 | 21.9 | + | 1.78 | 3.37E-04 | 5.50E-03 |
| response to endogenous stimulus (GO:0009719) | 1375 | 282 | 158.48 | + | 1.78 | 2.32E-23 | 2.89E-21 |
| RNA transport (GO:0050658) | 161 | 33 | 18.56 | + | 1.78 | 1.09E-03 | 1.50E-02 |
| nucleic acid transport (GO:0050657) | 161 | 33 | 18.56 | + | 1.78 | 1.09E-03 | 1.49E-02 |
| positive regulation of kinase activity (GO:0033674) | 290 | 59 | 33.42 | + | 1.77 | 1.14E-05 | 2.87E-04 |

| | | | | | | | |
|---|---|---|---|---|---|---|---|
| regulation of cellular component biogenesis (GO:0044087) | 979 | 199 | 112.84 | + | 1.76 | 3.79E-16 | 3.47E-14 |
| positive regulation of protein serine/threonine kinase activity (GO:0071902) | 123 | 25 | 14.18 | + | 1.76 | 4.23E-03 | 4.57E-02 |
| regulation of mitotic cell cycle (GO:0007346) | 497 | 101 | 57.28 | + | 1.76 | 1.31E-08 | 5.57E-07 |
| striated muscle cell differentiation (GO:0051146) | 222 | 45 | 25.59 | + | 1.76 | 1.80E-04 | 3.22E-03 |
| regulation of hemopoiesis (GO:1903706) | 405 | 82 | 46.68 | + | 1.76 | 3.57E-07 | 1.19E-05 |
| establishment of RNA localization (GO:0051236) | 163 | 33 | 18.79 | + | 1.76 | 1.20E-03 | 1.62E-02 |
| negative regulation of macromolecule biosynthetic process (GO:0010558) | 2055 | 416 | 236.85 | + | 1.76 | 1.15E-33 | 2.56E-31 |
| regulation of cell population proliferation (GO:0042127) | 1665 | 337 | 191.9 | + | 1.76 | 6.63E-27 | 9.92E-25 |
| actomyosin structure organization (GO:0031032) | 124 | 25 | 14.29 | + | 1.75 | 4.54E-03 | 4.88E-02 |
| regulation of actin filament-based process (GO:0032970) | 382 | 77 | 44.03 | + | 1.75 | 9.18E-07 | 2.83E-05 |
| negative regulation of epithelial cell proliferation (GO:0050680) | 134 | 27 | 15.44 | + | 1.75 | 3.88E-03 | 4.26E-02 |
| negative regulation of cytoskeleton organization (GO:0051494) | 154 | 31 | 17.75 | + | 1.75 | 2.07E-03 | 2.55E-02 |
| hemopoiesis (GO:0030097) | 686 | 138 | 79.07 | + | 1.75 | 4.65E-11 | 2.76E-09 |
| mRNA splicing, via spliceosome (GO:0000398) | 244 | 49 | 28.12 | + | 1.74 | 1.06E-04 | 2.07E-03 |
| RNA splicing, via transesterification reactions with bulged adenosine as nucleophile (GO:0000377) | 244 | 49 | 28.12 | + | 1.74 | 1.06E-04 | 2.07E-03 |
| negative regulation of cellular biosynthetic process (GO:0031327) | 2097 | 421 | 241.7 | + | 1.74 | 2.97E-33 | 6.16E-31 |

| | | | | | | | |
|---|---|---|---|---|---|---|---|
| skin development (GO:0043588) | 284 | 57 | 32.73 | + | 1.74 | 3.10E-05 | 7.03E-04 |
| regulation of mRNA metabolic process (GO:1903311) | 304 | 61 | 35.04 | + | 1.74 | 1.68E-05 | 4.08E-04 |
| protein phosphorylation (GO:0006468) | 489 | 98 | 56.36 | + | 1.74 | 3.67E-08 | 1.46E-06 |
| negative regulation of biosynthetic process (GO:0009890) | 2121 | 425 | 244.46 | + | 1.74 | 2.44E-33 | 5.13E-31 |
| determination of left/right symmetry (GO:0007368) | 135 | 27 | 15.56 | + | 1.74 | 4.07E-03 | 4.44E-02 |
| positive regulation of apoptotic signaling pathway (GO:2001235) | 135 | 27 | 15.56 | + | 1.74 | 4.07E-03 | 4.43E-02 |
| regulation of phosphatidylinositol 3-kinase/protein kinase B signal transduction (GO:0051896) | 255 | 51 | 29.39 | + | 1.74 | 9.87E-05 | 1.96E-03 |
| protein-RNA complex assembly (GO:0022618) | 195 | 39 | 22.48 | + | 1.74 | 6.22E-04 | 9.37E-03 |
| positive regulation of cell population proliferation (GO:0008284) | 942 | 188 | 108.57 | + | 1.73 | 1.90E-14 | 1.51E-12 |
| regulation of mitotic cell cycle phase transition (GO:1901990) | 332 | 66 | 38.27 | + | 1.72 | 8.25E-06 | 2.13E-04 |
| left/right pattern formation (GO:0060972) | 141 | 28 | 16.25 | + | 1.72 | 3.43E-03 | 3.82E-02 |
| cell recognition (GO:0008037) | 136 | 27 | 15.68 | + | 1.72 | 4.31E-03 | 4.65E-02 |
| positive regulation of apoptotic process (GO:0043065) | 510 | 101 | 58.78 | + | 1.72 | 4.63E-08 | 1.82E-06 |
| RNA splicing, via transesterification reactions (GO:0000375) | 248 | 49 | 28.58 | + | 1.71 | 1.81E-04 | 3.23E-03 |
| peptidyl-serine phosphorylation (GO:0018105) | 157 | 31 | 18.1 | + | 1.71 | 2.42E-03 | 2.87E-02 |
| adult behavior (GO:0030534) | 147 | 29 | 16.94 | + | 1.71 | 3.90E-03 | 4.28E-02 |

| | | | | | | | |
|---|---|---|---|---|---|---|---|
| positive regulation of multicellular organismal process (GO:0051240) | 1635 | 322 | 188.45 | + | 1.71 | 1.55E-23 | 1.96E-21 |
| negative regulation of cellular component organization (GO:0051129) | 683 | 134 | 78.72 | + | 1.7 | 5.03E-10 | 2.62E-08 |
| cellular response to organic cyclic compound (GO:0071407) | 459 | 90 | 52.9 | + | 1.7 | 3.85E-07 | 1.27E-05 |
| response to radiation (GO:0009314) | 424 | 83 | 48.87 | + | 1.7 | 1.07E-06 | 3.25E-05 |
| positive regulation of programmed cell death (GO:0043068) | 527 | 103 | 60.74 | + | 1.7 | 7.66E-08 | 2.90E-06 |
| epithelial cell proliferation (GO:0050673) | 174 | 34 | 20.05 | + | 1.7 | 1.81E-03 | 2.28E-02 |
| positive regulation of supramolecular fiber organization (GO:1902905) | 174 | 34 | 20.05 | + | 1.7 | 1.81E-03 | 2.28E-02 |
| protein-RNA complex organization (GO:0071826) | 205 | 40 | 23.63 | + | 1.69 | 8.60E-04 | 1.23E-02 |
| cell cycle phase transition (GO:0044770) | 159 | 31 | 18.33 | + | 1.69 | 3.72E-03 | 4.11E-02 |
| cell population proliferation (GO:0008283) | 719 | 140 | 82.87 | + | 1.69 | 2.98E-10 | 1.58E-08 |
| trans-synaptic signaling (GO:0099537) | 432 | 84 | 49.79 | + | 1.69 | 1.30E-06 | 3.89E-05 |
| epigenetic regulation of gene expression (GO:0040029) | 247 | 48 | 28.47 | + | 1.69 | 2.70E-04 | 4.59E-03 |
| positive regulation of DNA metabolic process (GO:0051054) | 294 | 57 | 33.89 | + | 1.68 | 9.51E-05 | 1.89E-03 |
| response to alcohol (GO:0097305) | 258 | 50 | 29.74 | + | 1.68 | 2.45E-04 | 4.21E-03 |
| regulation of microtubule cytoskeleton organization (GO:0070507) | 160 | 31 | 18.44 | + | 1.68 | 3.85E-03 | 4.24E-02 |
| positive regulation of cellular component organization (GO:0051130) | 1115 | 216 | 128.51 | + | 1.68 | 4.62E-15 | 3.84E-13 |

| | | | | | | | |
|---|---|---|---|---|---|---|---|
| positive regulation of leukocyte differentiation (GO:1902107) | 191 | 37 | 22.01 | + | 1.68 | 1.35E-03 | 1.80E-02 |
| positive regulation of hemopoiesis (GO:1903708) | 191 | 37 | 22.01 | + | 1.68 | 1.35E-03 | 1.80E-02 |
| regulation of actin cytoskeleton organization (GO:0032956) | 341 | 66 | 39.3 | + | 1.68 | 2.33E-05 | 5.50E-04 |
| regulation of growth (GO:0040008) | 589 | 114 | 67.89 | + | 1.68 | 2.00E-08 | 8.25E-07 |
| cell surface receptor signaling pathway (GO:0007166) | 2057 | 398 | 237.08 | + | 1.68 | 9.15E-28 | 1.46E-25 |
| mRNA processing (GO:0006397) | 456 | 88 | 52.56 | + | 1.67 | 1.14E-06 | 3.44E-05 |
| homeostasis of number of cells (GO:0048872) | 280 | 54 | 32.27 | + | 1.67 | 1.42E-04 | 2.65E-03 |
| negative regulation of cell cycle process (GO:0010948) | 270 | 52 | 31.12 | + | 1.67 | 2.31E-04 | 4.01E-03 |
| cellular developmental process (GO:0048869) | 3641 | 701 | 419.65 | + | 1.67 | 5.96E-52 | 6.93E-49 |
| cell differentiation (GO:0030154) | 3638 | 700 | 419.31 | + | 1.67 | 1.27E-51 | 1.37E-48 |
| regulation of RNA biosynthetic process (GO:2001141) | 3402 | 654 | 392.11 | + | 1.67 | 1.70E-47 | 1.17E-44 |
| regulation of DNA-templated transcription (GO:0006355) | 3383 | 650 | 389.92 | + | 1.67 | 4.13E-47 | 2.72E-44 |
| establishment or maintenance of cell polarity (GO:0007163) | 203 | 39 | 23.4 | + | 1.67 | 1.25E-03 | 1.68E-02 |
| regulation of RNA metabolic process (GO:0051252) | 3657 | 701 | 421.5 | + | 1.66 | 3.47E-51 | 3.50E-48 |
| phosphorylation (GO:0016310) | 548 | 105 | 63.16 | + | 1.66 | 1.38E-07 | 5.00E-06 |
| RNA splicing (GO:0008380) | 376 | 72 | 43.34 | + | 1.66 | 1.33E-05 | 3.31E-04 |

| | | | | | | | |
|---|---|---|---|---|---|---|---|
| epithelial cell differentiation (GO:0030855) | 632 | 121 | 72.84 | + | 1.66 | 1.38E-08 | 5.86E-07 |
| positive regulation of cytoskeleton organization (GO:0051495) | 183 | 35 | 21.09 | + | 1.66 | 2.35E-03 | 2.81E-02 |
| behavior (GO:0007610) | 618 | 118 | 71.23 | + | 1.66 | 2.86E-08 | 1.16E-06 |
| synaptic signaling (GO:0099536) | 462 | 88 | 53.25 | + | 1.65 | 1.96E-06 | 5.72E-05 |
| positive regulation of macromolecule biosynthetic process (GO:0010557) | 2568 | 489 | 295.98 | + | 1.65 | 6.92E-33 | 1.39E-30 |
| regulation of cytoskeleton organization (GO:0051493) | 520 | 99 | 59.93 | + | 1.65 | 4.39E-07 | 1.44E-05 |
| nucleocytoplasmic transport (GO:0006913) | 247 | 47 | 28.47 | + | 1.65 | 5.79E-04 | 8.88E-03 |
| negative regulation of cell cycle (GO:0045786) | 358 | 68 | 41.26 | + | 1.65 | 3.65E-05 | 8.16E-04 |
| cellular response to starvation (GO:0009267) | 174 | 33 | 20.05 | + | 1.65 | 3.89E-03 | 4.27E-02 |
| supramolecular fiber organization (GO:0097435) | 596 | 113 | 68.69 | + | 1.64 | 7.50E-08 | 2.84E-06 |
| nuclear transport (GO:0051169) | 248 | 47 | 28.58 | + | 1.64 | 6.00E-04 | 9.12E-03 |
| negative regulation of cellular metabolic process (GO:0031324) | 2425 | 459 | 279.5 | + | 1.64 | 5.04E-30 | 9.07E-28 |
| actin cytoskeleton organization (GO:0030036) | 550 | 104 | 63.39 | + | 1.64 | 3.09E-07 | 1.04E-05 |
| negative regulation of cell adhesion (GO:0007162) | 291 | 55 | 33.54 | + | 1.64 | 1.98E-04 | 3.49E-03 |
| anatomical structure development (GO:0048856) | 5208 | 984 | 600.26 | + | 1.64 | 1.47E-75 | 7.41E-72 |
| multicellular organismal-level homeostasis (GO:0048871) | 614 | 116 | 70.77 | + | 1.64 | 7.83E-08 | 2.95E-06 |

| | | | | | | | |
|---|---|---|---|---|---|---|---|
| regulation of actin filament organization (GO:0110053) | 270 | 51 | 31.12 | + | 1.64 | 3.51E-04 | 5.66E-03 |
| regulation of MAPK cascade (GO:0043408) | 636 | 120 | 73.3 | + | 1.64 | 4.48E-08 | 1.77E-06 |
| regulation of supramolecular fiber organization (GO:1902903) | 387 | 73 | 44.6 | + | 1.64 | 2.49E-05 | 5.83E-04 |
| positive regulation of transferase activity (GO:0051347) | 361 | 68 | 41.61 | + | 1.63 | 4.06E-05 | 9.00E-04 |
| positive regulation of cellular biosynthetic process (GO:0031328) | 2625 | 494 | 302.55 | + | 1.63 | 6.24E-32 | 1.21E-29 |
| regulation of DNA metabolic process (GO:0051052) | 505 | 95 | 58.21 | + | 1.63 | 1.32E-06 | 3.94E-05 |
| chemical synaptic transmission (GO:0007268) | 415 | 78 | 47.83 | + | 1.63 | 1.19E-05 | 2.97E-04 |
| anterograde trans-synaptic signaling (GO:0098916) | 415 | 78 | 47.83 | + | 1.63 | 1.19E-05 | 2.96E-04 |
| positive regulation of biosynthetic process (GO:0009891) | 2668 | 501 | 307.51 | + | 1.63 | 3.47E-32 | 6.82E-30 |
| actin filament-based process (GO:0030029) | 619 | 116 | 71.34 | + | 1.63 | 9.32E-08 | 3.48E-06 |
| regulation of nucleobase-containing compound metabolic process (GO:0019219) | 3961 | 742 | 456.54 | + | 1.63 | 1.13E-50 | 1.00E-47 |
| cellular response to organonitrogen compound (GO:0071417) | 497 | 93 | 57.28 | + | 1.62 | 2.23E-06 | 6.42E-05 |
| regulation of cell cycle phase transition (GO:1901987) | 428 | 80 | 49.33 | + | 1.62 | 1.13E-05 | 2.86E-04 |
| positive regulation of MAPK cascade (GO:0043410) | 455 | 85 | 52.44 | + | 1.62 | 7.08E-06 | 1.85E-04 |
| positive regulation of signal transduction (GO:0009967) | 1558 | 291 | 179.57 | + | 1.62 | 7.36E-18 | 7.68E-16 |
| positive regulation of growth (GO:0045927) | 247 | 46 | 28.47 | + | 1.62 | 8.95E-04 | 1.27E-02 |

| | | | | | | | |
|---|---|---|---|---|---|---|---|
| negative regulation of cell cycle phase transition (GO:1901988) | 226 | 42 | 26.05 | + | 1.61 | 1.59E-03 | 2.05E-02 |
| regulation of kinase activity (GO:0043549) | 506 | 94 | 58.32 | + | 1.61 | 2.75E-06 | 7.79E-05 |
| anatomical structure homeostasis (GO:0060249) | 221 | 41 | 25.47 | + | 1.61 | 1.98E-03 | 2.46E-02 |
| tissue homeostasis (GO:0001894) | 221 | 41 | 25.47 | + | 1.61 | 1.98E-03 | 2.46E-02 |
| cellular response to oxidative stress (GO:0034599) | 221 | 41 | 25.47 | + | 1.61 | 1.98E-03 | 2.45E-02 |
| peptidyl-amino acid modification (GO:0018193) | 523 | 97 | 60.28 | + | 1.61 | 2.01E-06 | 5.85E-05 |
| regulation of multicellular organismal process (GO:0051239) | 2931 | 543 | 337.82 | + | 1.61 | 1.49E-33 | 3.21E-31 |
| regulation of cellular component organization (GO:0051128) | 2413 | 447 | 278.12 | + | 1.61 | 4.42E-27 | 6.68E-25 |
| cellular response to abiotic stimulus (GO:0071214) | 319 | 59 | 36.77 | + | 1.6 | 2.59E-04 | 4.42E-03 |
| cellular response to environmental stimulus (GO:0104004) | 319 | 59 | 36.77 | + | 1.6 | 2.59E-04 | 4.42E-03 |
| positive regulation of macromolecule metabolic process (GO:0010604) | 3364 | 622 | 387.73 | + | 1.6 | 4.49E-39 | 1.62E-36 |
| regulation of cell cycle process (GO:0010564) | 717 | 132 | 82.64 | + | 1.6 | 3.58E-08 | 1.43E-06 |
| intracellular signaling cassette (GO:0141124) | 826 | 152 | 95.2 | + | 1.6 | 4.01E-09 | 1.78E-07 |
| wound healing (GO:0042060) | 321 | 59 | 37 | + | 1.59 | 2.76E-04 | 4.66E-03 |
| apoptotic signaling pathway (GO:0097190) | 316 | 58 | 36.42 | + | 1.59 | 3.46E-04 | 5.63E-03 |
| regulation of cell communication (GO:0010646) | 3428 | 629 | 395.1 | + | 1.59 | 1.72E-38 | 5.53E-36 |

| | | | | | | | |
|---|---|---|---|---|---|---|---|
| positive regulation of cellular metabolic process (GO:0031325) | 3124 | 573 | 360.06 | + | 1.59 | 1.86E-34 | 4.38E-32 |
| regulation of signaling (GO:0023051) | 3423 | 627 | 394.53 | + | 1.59 | 4.76E-38 | 1.36E-35 |
| regulation of signal transduction (GO:0009966) | 3017 | 551 | 347.73 | + | 1.58 | 2.02E-32 | 4.02E-30 |
| response to light stimulus (GO:0009416) | 313 | 57 | 36.08 | + | 1.58 | 4.61E-04 | 7.23E-03 |
| response to wounding (GO:0009611) | 429 | 78 | 49.45 | + | 1.58 | 4.53E-05 | 9.91E-04 |
| regulation of binding (GO:0051098) | 226 | 41 | 26.05 | + | 1.57 | 3.16E-03 | 3.55E-02 |
| response to starvation (GO:0042594) | 215 | 39 | 24.78 | + | 1.57 | 3.64E-03 | 4.03E-02 |
| negative regulation of multicellular organismal process (GO:0051241) | 1092 | 198 | 125.86 | + | 1.57 | 4.98E-11 | 2.93E-09 |
| mRNA metabolic process (GO:0016071) | 607 | 110 | 69.96 | + | 1.57 | 1.15E-06 | 3.47E-05 |
| regulation of protein serine/threonine kinase activity (GO:0071900) | 254 | 46 | 29.28 | + | 1.57 | 2.01E-03 | 2.48E-02 |
| developmental process (GO:0032502) | 5711 | 1034 | 658.24 | + | 1.57 | 1.54E-69 | 5.80E-66 |
| positive regulation of protein kinase activity (GO:0045860) | 244 | 44 | 28.12 | + | 1.56 | 2.38E-03 | 2.84E-02 |
| positive regulation of metabolic process (GO:0009893) | 3677 | 663 | 423.8 | + | 1.56 | 2.95E-38 | 8.75E-36 |
| regulation of cell growth (GO:0001558) | 394 | 71 | 45.41 | + | 1.56 | 1.21E-04 | 2.30E-03 |
| cellular response to chemical stress (GO:0062197) | 278 | 50 | 32.04 | + | 1.56 | 1.25E-03 | 1.69E-02 |
| positive regulation of cell communication (GO:0010647) | 1741 | 313 | 200.66 | + | 1.56 | 1.10E-16 | 1.06E-14 |

| | | | | | | | |
|---|---|---|---|---|---|---|---|
| cellular response to nitrogen compound (GO:1901699) | 573 | 103 | 66.04 | + | 1.56 | 4.10E-06 | 1.13E-04 |
| positive regulation of signaling (GO:0023056) | 1743 | 313 | 200.89 | + | 1.56 | 1.18E-16 | 1.14E-14 |
| negative regulation of macromolecule metabolic process (GO:0010605) | 2602 | 467 | 299.9 | + | 1.56 | 3.87E-25 | 5.28E-23 |
| response to hormone (GO:0009725) | 787 | 141 | 90.71 | + | 1.55 | 7.35E-08 | 2.79E-06 |
| response to peptide hormone (GO:0043434) | 374 | 67 | 43.11 | + | 1.55 | 2.21E-04 | 3.84E-03 |
| negative regulation of organelle organization (GO:0010639) | 335 | 60 | 38.61 | + | 1.55 | 5.16E-04 | 7.97E-03 |
| regulation of leukocyte differentiation (GO:1902105) | 324 | 58 | 37.34 | + | 1.55 | 5.95E-04 | 9.10E-03 |
| anatomical structure maturation (GO:0071695) | 236 | 42 | 27.2 | + | 1.54 | 3.93E-03 | 4.30E-02 |
| cellular response to hormone stimulus (GO:0032870) | 508 | 90 | 58.55 | + | 1.54 | 3.00E-05 | 6.86E-04 |
| positive regulation of phosphorus metabolic process (GO:0010562) | 683 | 121 | 78.72 | + | 1.54 | 1.28E-06 | 3.83E-05 |
| positive regulation of phosphate metabolic process (GO:0045937) | 683 | 121 | 78.72 | + | 1.54 | 1.28E-06 | 3.83E-05 |
| response to nutrient levels (GO:0031667) | 487 | 86 | 56.13 | + | 1.53 | 5.25E-05 | 1.14E-03 |
| regulation of cell cycle (GO:0051726) | 1082 | 191 | 124.71 | + | 1.53 | 1.05E-09 | 5.24E-08 |
| negative regulation of signal transduction (GO:0009968) | 1309 | 231 | 150.87 | + | 1.53 | 1.48E-11 | 9.22E-10 |
| response to abiotic stimulus (GO:0009628) | 1100 | 194 | 126.78 | + | 1.53 | 7.78E-10 | 3.96E-08 |
| cellular response to oxygen-containing compound (GO:1901701) | 1010 | 178 | 116.41 | + | 1.53 | 5.03E-09 | 2.21E-07 |

| | | | | | | | |
|---|---|---|---|---|---|---|---|
| positive regulation of cell cycle process (GO:0090068) | 256 | 45 | 29.51 | + | 1.53 | 4.05E-03 | 4.42E-02 |
| negative regulation of cellular process (GO:0048523) | 4749 | 834 | 547.36 | + | 1.52 | 5.52E-46 | 3.09E-43 |
| response to steroid hormone (GO:0048545) | 285 | 50 | 32.85 | + | 1.52 | 2.63E-03 | 3.10E-02 |
| positive regulation of phosphorylation (GO:0042327) | 605 | 106 | 69.73 | + | 1.52 | 9.94E-06 | 2.54E-04 |
| negative regulation of cell communication (GO:0010648) | 1400 | 245 | 161.36 | + | 1.52 | 7.66E-12 | 4.89E-10 |
| positive regulation of cell-cell adhesion (GO:0022409) | 326 | 57 | 37.57 | + | 1.52 | 1.54E-03 | 2.00E-02 |
| negative regulation of metabolic process (GO:0009892) | 2815 | 491 | 324.45 | + | 1.51 | 1.03E-23 | 1.33E-21 |
| negative regulation of signaling (GO:0023057) | 1401 | 244 | 161.48 | + | 1.51 | 1.43E-11 | 8.99E-10 |
| regulation of apoptotic process (GO:0042981) | 1459 | 254 | 168.16 | + | 1.51 | 5.70E-12 | 3.65E-10 |
| regulation of ERK1 and ERK2 cascade (GO:0070372) | 270 | 47 | 31.12 | + | 1.51 | 3.83E-03 | 4.23E-02 |
| regulation of GTPase activity (GO:0043087) | 276 | 48 | 31.81 | + | 1.51 | 4.17E-03 | 4.51E-02 |
| regulation of organelle organization (GO:0033043) | 1186 | 206 | 136.7 | + | 1.51 | 9.18E-10 | 4.60E-08 |
| intracellular signal transduction (GO:0035556) | 1694 | 294 | 195.25 | + | 1.51 | 1.12E-13 | 8.53E-12 |
| positive regulation of cell cycle (GO:0045787) | 323 | 56 | 37.23 | + | 1.5 | 1.98E-03 | 2.45E-02 |
| regulation of intracellular signal transduction (GO:1902531) | 1940 | 336 | 223.6 | + | 1.5 | 1.71E-15 | 1.46E-13 |
| positive regulation of cellular process (GO:0048522) | 5601 | 969 | 645.56 | + | 1.5 | 6.63E-53 | 8.35E-50 |

| | | | | | | | |
|---|---|---|---|---|---|---|---|
| cellular response to lipid (GO:0071396) | 515 | 89 | 59.36 | + | 1.5 | 8.50E-05 | 1.72E-03 |
| actin filament organization (GO:0007015) | 278 | 48 | 32.04 | + | 1.5 | 4.33E-03 | 4.67E-02 |
| negative regulation of gene expression (GO:0010629) | 985 | 170 | 113.53 | + | 1.5 | 5.07E-08 | 1.98E-06 |
| regulation of cellular component size (GO:0032535) | 354 | 61 | 40.8 | + | 1.5 | 1.34E-03 | 1.80E-02 |
| leukocyte differentiation (GO:0002521) | 424 | 73 | 48.87 | + | 1.49 | 5.06E-04 | 7.82E-03 |
| negative regulation of apoptotic process (GO:0043066) | 884 | 152 | 101.89 | + | 1.49 | 3.63E-07 | 1.21E-05 |
| regulation of programmed cell death (GO:0043067) | 1501 | 258 | 173 | + | 1.49 | 1.46E-11 | 9.13E-10 |
| positive regulation of intracellular signal transduction (GO:1902533) | 1102 | 189 | 127.01 | + | 1.49 | 1.21E-08 | 5.17E-07 |
| epidermis development (GO:0008544) | 339 | 58 | 39.07 | + | 1.48 | 1.95E-03 | 2.42E-02 |
| cell-cell signaling (GO:0007267) | 831 | 142 | 95.78 | + | 1.48 | 1.25E-06 | 3.75E-05 |
| negative regulation of programmed cell death (GO:0043069) | 913 | 156 | 105.23 | + | 1.48 | 3.23E-07 | 1.08E-05 |
| regulation of DNA-binding transcription factor activity (GO:0051090) | 346 | 59 | 39.88 | + | 1.48 | 2.13E-03 | 2.61E-02 |
| regulation of gene expression (GO:0010468) | 4865 | 829 | 560.73 | + | 1.48 | 4.68E-40 | 1.91E-37 |
| negative regulation of biological process (GO:0048519) | 5171 | 880 | 596 | + | 1.48 | 4.15E-43 | 1.96E-40 |
| response to organonitrogen compound (GO:0010243) | 902 | 153 | 103.96 | + | 1.47 | 8.24E-07 | 2.61E-05 |
| regulation of macromolecule biosynthetic process (GO:0010556) | 4989 | 846 | 575.02 | + | 1.47 | 2.80E-40 | 1.18E-37 |

| | | | | | | | |
|---|---|---|---|---|---|---|---|
| regulation of primary metabolic process (GO:0080090) | 5544 | 939 | 638.99 | + | 1.47 | 4.31E-46 | 2.50E-43 |
| regulation of phosphorylation (GO:0042325) | 963 | 163 | 110.99 | + | 1.47 | 3.60E-07 | 1.20E-05 |
| cell division (GO:0051301) | 520 | 88 | 59.93 | + | 1.47 | 2.13E-04 | 3.75E-03 |
| regulation of transferase activity (GO:0051338) | 609 | 103 | 70.19 | + | 1.47 | 6.05E-05 | 1.28E-03 |
| regulation of protein kinase activity (GO:0045859) | 444 | 75 | 51.17 | + | 1.47 | 6.83E-04 | 1.01E-02 |
| regulation of cellular biosynthetic process (GO:0031326) | 5078 | 856 | 585.28 | + | 1.46 | 8.28E-40 | 3.29E-37 |
| negative regulation of response to stimulus (GO:0048585) | 1656 | 279 | 190.87 | + | 1.46 | 1.70E-11 | 1.06E-09 |
| response to oxygen-containing compound (GO:1901700) | 1507 | 253 | 173.69 | + | 1.46 | 2.73E-10 | 1.47E-08 |
| regulation of phosphate metabolic process (GO:0019220) | 1121 | 188 | 129.2 | + | 1.46 | 8.20E-08 | 3.08E-06 |
| regulation of phosphorus metabolic process (GO:0051174) | 1122 | 188 | 129.32 | + | 1.45 | 8.36E-08 | 3.14E-06 |
| positive regulation of biological process (GO:0048518) | 6177 | 1034 | 711.95 | + | 1.45 | 3.33E-50 | 2.79E-47 |
| regulation of biosynthetic process (GO:0009889) | 5150 | 862 | 593.58 | + | 1.45 | 5.59E-39 | 1.92E-36 |
| positive regulation of protein modification process (GO:0031401) | 742 | 124 | 85.52 | + | 1.45 | 1.79E-05 | 4.29E-04 |
| positive regulation of cell activation (GO:0050867) | 377 | 63 | 43.45 | + | 1.45 | 2.49E-03 | 2.94E-02 |
| response to organic cyclic compound (GO:0014070) | 839 | 140 | 96.7 | + | 1.45 | 5.51E-06 | 1.48E-04 |
| cytoskeleton organization (GO:0007010) | 1256 | 209 | 144.76 | + | 1.44 | 2.37E-08 | 9.73E-07 |

| | | | | | | | |
|---|---|---|---|---|---|---|---|
| multicellular organismal process (GO:0032501) | 6239 | 1038 | 719.09 | + | 1.44 | 5.38E-49 | 4.28E-46 |
| positive regulation of protein phosphorylation (GO:0001934) | 560 | 93 | 64.54 | + | 1.44 | 2.76E-04 | 4.66E-03 |
| regulation of response to stimulus (GO:0048583) | 4010 | 665 | 462.18 | + | 1.44 | 7.98E-27 | 1.18E-24 |
| mononuclear cell differentiation (GO:1903131) | 362 | 60 | 41.72 | + | 1.44 | 3.57E-03 | 3.96E-02 |
| apoptotic process (GO:0006915) | 1033 | 171 | 119.06 | + | 1.44 | 8.66E-07 | 2.70E-05 |
| regulation of apoptotic signaling pathway (GO:2001233) | 375 | 62 | 43.22 | + | 1.43 | 3.23E-03 | 3.63E-02 |
| regulation of anatomical structure size (GO:0090066) | 496 | 82 | 57.17 | + | 1.43 | 7.82E-04 | 1.14E-02 |
| positive regulation of response to stimulus (GO:0048584) | 2241 | 370 | 258.29 | + | 1.43 | 8.56E-14 | 6.61E-12 |
| positive regulation of organelle organization (GO:0010638) | 509 | 84 | 58.67 | + | 1.43 | 7.06E-04 | 1.04E-02 |
| mitotic cell cycle (GO:0000278) | 595 | 98 | 68.58 | + | 1.43 | 2.52E-04 | 4.33E-03 |
| regulation of protein-containing complex assembly (GO:0043254) | 419 | 69 | 48.29 | + | 1.43 | 2.46E-03 | 2.92E-02 |
| positive regulation of hydrolase activity (GO:0051345) | 426 | 70 | 49.1 | + | 1.43 | 2.11E-03 | 2.59E-02 |
| cellular component organization (GO:0016043) | 5623 | 922 | 648.09 | + | 1.42 | 1.07E-38 | 3.53E-36 |
| protein-containing complex organization (GO:0043933) | 2044 | 335 | 235.59 | + | 1.42 | 4.35E-12 | 2.82E-10 |
| mitotic cell cycle process (GO:1903047) | 513 | 84 | 59.13 | + | 1.42 | 9.52E-04 | 1.34E-02 |
| response to nitrogen compound (GO:1901698) | 1009 | 165 | 116.29 | + | 1.42 | 3.03E-06 | 8.52E-05 |

| | | | | | | | |
|---|---|---|---|---|---|---|---|
| regulation of cellular metabolic process (GO:0031323) | 5837 | 953 | 672.76 | + | 1.42 | 1.17E-39 | 4.40E-37 |
| regulation of cell-cell adhesion (GO:0022407) | 490 | 80 | 56.48 | + | 1.42 | 1.24E-03 | 1.67E-02 |
| cell death (GO:0008219) | 1072 | 175 | 123.56 | + | 1.42 | 1.37E-06 | 4.07E-05 |
| negative regulation of intracellular signal transduction (GO:1902532) | 650 | 106 | 74.92 | + | 1.41 | 2.19E-04 | 3.82E-03 |
| regulation of macromolecule metabolic process (GO:0060255) | 5998 | 974 | 691.31 | + | 1.41 | 9.57E-40 | 3.71E-37 |
| cell cycle process (GO:0022402) | 881 | 143 | 101.54 | + | 1.41 | 1.93E-05 | 4.60E-04 |
| programmed cell death (GO:0012501) | 1067 | 173 | 122.98 | + | 1.41 | 2.68E-06 | 7.61E-05 |
| cellular component organization or biogenesis (GO:0071840) | 5838 | 942 | 672.87 | + | 1.4 | 9.66E-37 | 2.56E-34 |
| response to lipid (GO:0033993) | 841 | 135 | 96.93 | + | 1.39 | 6.79E-05 | 1.41E-03 |
| cellular response to chemical stimulus (GO:0070887) | 1908 | 306 | 219.91 | + | 1.39 | 5.87E-10 | 3.02E-08 |
| regulation of system process (GO:0044057) | 544 | 87 | 62.7 | + | 1.39 | 1.68E-03 | 2.17E-02 |
| regulation of protein phosphorylation (GO:0001932) | 886 | 141 | 102.12 | + | 1.38 | 6.54E-05 | 1.36E-03 |
| positive regulation of molecular function (GO:0044093) | 1140 | 181 | 131.39 | + | 1.38 | 5.51E-06 | 1.48E-04 |
| regulation of metabolic process (GO:0019222) | 6519 | 1032 | 751.36 | + | 1.37 | 7.96E-38 | 2.23E-35 |
| microtubule cytoskeleton organization (GO:0000226) | 557 | 88 | 64.2 | + | 1.37 | 2.39E-03 | 2.84E-02 |
| cellular response to stress (GO:0033554) | 1589 | 251 | 183.14 | + | 1.37 | 9.83E-08 | 3.62E-06 |

| | | | | | | | |
|---|---|---|---|---|---|---|---|
| positive regulation of catalytic activity (GO:0043085) | 820 | 129 | 94.51 | + | 1.36 | 2.22E-04 | 3.85E-03 |
| positive regulation of protein metabolic process (GO:0051247) | 1221 | 192 | 140.73 | + | 1.36 | 5.60E-06 | 1.50E-04 |
| positive regulation of catabolic process (GO:0009896) | 522 | 82 | 60.16 | + | 1.36 | 3.48E-03 | 3.86E-02 |
| DNA damage response (GO:0006974) | 780 | 122 | 89.9 | + | 1.36 | 4.70E-04 | 7.29E-03 |
| DNA-templated transcription (GO:0006351) | 546 | 85 | 62.93 | + | 1.35 | 4.22E-03 | 4.56E-02 |
| regulation of protein modification process (GO:0031399) | 1203 | 187 | 138.65 | + | 1.35 | 1.45E-05 | 3.58E-04 |
| homeostatic process (GO:0042592) | 1429 | 222 | 164.7 | + | 1.35 | 2.20E-06 | 6.32E-05 |
| cell cycle (GO:0007049) | 973 | 150 | 112.15 | + | 1.34 | 1.69E-04 | 3.05E-03 |
| negative regulation of protein metabolic process (GO:0051248) | 797 | 122 | 91.86 | + | 1.33 | 1.01E-03 | 1.41E-02 |
| developmental process involved in reproduction (GO:0003006) | 995 | 152 | 114.68 | + | 1.33 | 2.42E-04 | 4.18E-03 |
| regulation of protein metabolic process (GO:0051246) | 2110 | 320 | 243.19 | + | 1.32 | 9.44E-08 | 3.51E-06 |
| regulation of biological quality (GO:0065008) | 2822 | 427 | 325.26 | + | 1.31 | 4.52E-10 | 2.37E-08 |
| regulation of hydrolase activity (GO:0051336) | 676 | 102 | 77.91 | + | 1.31 | 4.74E-03 | 4.97E-02 |
| regulation of molecular function (GO:0065009) | 1903 | 287 | 219.34 | + | 1.31 | 9.13E-07 | 2.82E-05 |
| cell communication (GO:0007154) | 5158 | 774 | 594.5 | + | 1.3 | 1.20E-18 | 1.32E-16 |
| positive regulation of gene expression (GO:0010628) | 1180 | 177 | 136 | + | 1.3 | 2.02E-04 | 3.56E-03 |

| | | | | | | | |
|---|---|---|---|---|---|---|---|
| signaling (GO:0023052) | 5116 | 767 | 589.66 | + | 1.3 | 2.41E-18 | 2.62E-16 |
| regulation of catabolic process (GO:0009894) | 1017 | 152 | 117.22 | + | 1.3 | 7.23E-04 | 1.06E-02 |
| regulation of catalytic activity (GO:0050790) | 1323 | 197 | 152.49 | + | 1.29 | 1.26E-04 | 2.37E-03 |
| signal transduction (GO:0007165) | 4794 | 710 | 552.54 | + | 1.28 | 2.36E-15 | 1.99E-13 |
| RNA biosynthetic process (GO:0032774) | 1367 | 202 | 157.56 | + | 1.28 | 1.60E-04 | 2.91E-03 |
| regulation of cellular process (GO:0050794) | 11103 | 1625 | 1279.7 | + | 1.27 | 5.81E-53 | 7.99E-50 |
| RNA metabolic process (GO:0016070) | 1519 | 220 | 175.08 | + | 1.26 | 2.75E-04 | 4.66E-03 |
| regulation of biological process (GO:0050789) | 11711 | 1695 | 1349.78 | + | 1.26 | 4.07E-54 | 6.15E-51 |
| cellular response to stimulus (GO:0051716) | 6306 | 910 | 726.81 | + | 1.25 | 1.61E-17 | 1.64E-15 |
| regulation of cellular localization (GO:0060341) | 978 | 141 | 112.72 | + | 1.25 | 4.73E-03 | 4.96E-02 |
| regulation of response to stress (GO:0080134) | 1376 | 198 | 158.59 | + | 1.25 | 8.80E-04 | 1.26E-02 |
| nucleic acid metabolic process (GO:0090304) | 2140 | 306 | 246.65 | + | 1.24 | 3.82E-05 | 8.51E-04 |
| nucleic acid biosynthetic process (GO:0141187) | 1455 | 208 | 167.7 | + | 1.24 | 8.80E-04 | 1.26E-02 |
| protein modification process (GO:0036211) | 2055 | 293 | 236.85 | + | 1.24 | 7.08E-05 | 1.46E-03 |
| reproductive process (GO:0022414) | 1473 | 210 | 169.77 | + | 1.24 | 9.43E-04 | 1.33E-02 |
| biological regulation (GO:0065007) | 12149 | 1732 | 1400.26 | + | 1.24 | 4.33E-51 | 4.10E-48 |

| | | | | | | | |
|---|---|---|---|---|---|---|---|
| cellular component assembly (GO:0022607) | 2459 | 344 | 283.42 | + | 1.21 | 7.03E-05 | 1.45E-03 |
| macromolecule modification (GO:0043412) | 2239 | 307 | 258.06 | + | 1.19 | 7.60E-04 | 1.11E-02 |
| protein localization (GO:0008104) | 1950 | 267 | 224.75 | + | 1.19 | 1.97E-03 | 2.44E-02 |
| cellular macromolecule localization (GO:0070727) | 1958 | 268 | 225.67 | + | 1.19 | 2.01E-03 | 2.48E-02 |
| cellular component biogenesis (GO:0044085) | 2705 | 367 | 311.77 | + | 1.18 | 4.80E-04 | 7.42E-03 |
| response to stimulus (GO:0050896) | 8080 | 1089 | 931.28 | + | 1.17 | 2.89E-12 | 1.91E-10 |
| response to chemical (GO:0042221) | 3643 | 489 | 419.88 | + | 1.16 | 9.98E-05 | 1.98E-03 |
| cellular localization (GO:0051641) | 2659 | 351 | 306.47 | + | 1.15 | 4.18E-03 | 4.53E-02 |
| response to stress (GO:0006950) | 3411 | 450 | 393.14 | + | 1.14 | 1.01E-03 | 1.41E-02 |
| macromolecule metabolic process (GO:0043170) | 5602 | 727 | 645.67 | + | 1.13 | 7.84E-05 | 1.59E-03 |
| cellular process (GO:0009987) | 14769 | 1877 | 1702.24 | + | 1.1 | 4.01E-18 | 4.30E-16 |
| biological_process (GO:0008150) | 17791 | 2191 | 2050.55 | + | 1.07 | 1.63E-21 | 1.93E-19 |
| catabolic process (GO:0009056) | 1944 | 172 | 224.06 | - | 0.77 | 7.44E-05 | 1.52E-03 |
| organonitrogen compound catabolic process (GO:1901565) | 1116 | 98 | 128.63 | - | 0.76 | 2.77E-03 | 3.23E-02 |
| small molecule metabolic process (GO:0044281) | 1629 | 133 | 187.75 | - | 0.71 | 4.70E-06 | 1.28E-04 |
| defense response to other organism (GO:0098542) | 1015 | 82 | 116.99 | - | 0.7 | 2.77E-04 | 4.65E-03 |

| | | | | | | | |
|---|---|---|---|---|---|---|---|
| response to bacterium (GO:0009617) | 682 | 55 | 78.61 | - | 0.7 | 3.35E-03 | 3.74E-02 |
| sensory perception (GO:0007600) | 976 | 77 | 112.49 | - | 0.68 | 1.71E-04 | 3.08E-03 |
| immune response (GO:0006955) | 1662 | 127 | 191.56 | - | 0.66 | 7.26E-08 | 2.76E-06 |
| immune effector process (GO:0002252) | 479 | 36 | 55.21 | - | 0.65 | 4.62E-03 | 4.89E-02 |
| mitochondrion organization (GO:0007005) | 442 | 32 | 50.94 | - | 0.63 | 3.24E-03 | 3.63E-02 |
| G protein-coupled receptor signaling pathway (GO:0007186) | 1255 | 90 | 144.65 | - | 0.62 | 1.79E-07 | 6.30E-06 |
| purine nucleotide metabolic process (GO:0006163) | 468 | 33 | 53.94 | - | 0.61 | 1.57E-03 | 2.03E-02 |
| lipid localization (GO:0010876) | 384 | 27 | 44.26 | - | 0.61 | 4.58E-03 | 4.92E-02 |
| oxoacid metabolic process (GO:0043436) | 813 | 57 | 93.7 | - | 0.61 | 1.50E-05 | 3.68E-04 |
| nucleotide metabolic process (GO:0009117) | 528 | 37 | 60.86 | - | 0.61 | 6.81E-04 | 1.01E-02 |
| organic acid metabolic process (GO:0006082) | 820 | 57 | 94.51 | - | 0.6 | 1.23E-05 | 3.07E-04 |
| nucleoside phosphate metabolic process (GO:0006753) | 536 | 37 | 61.78 | - | 0.6 | 4.47E-04 | 7.02E-03 |
| nucleobase-containing small molecule metabolic process (GO:0055086) | 605 | 41 | 69.73 | - | 0.59 | 9.91E-05 | 1.97E-03 |
| carboxylic acid metabolic process (GO:0019752) | 791 | 53 | 91.17 | - | 0.58 | 5.04E-06 | 1.37E-04 |
| purine-containing compound metabolic process (GO:0072521) | 498 | 33 | 57.4 | - | 0.57 | 2.76E-04 | 4.66E-03 |
| Unclassified (UNCLASSIFIED) | 2789 | 181 | 321.45 | - | 0.56 | 1.63E-21 | 1.94E-19 |

| | | | | | | | |
|---|---|---|---|---|---|---|---|
| autophagy (GO:0006914) | 327 | 21 | 37.69 | - | 0.56 | 2.84E-03 | 3.32E-02 |
| process utilizing autophagic mechanism (GO:0061919) | 327 | 21 | 37.69 | - | 0.56 | 2.84E-03 | 3.31E-02 |
| adaptive immune response based on somatic recombination of immune receptors built from immunoglobulin superfamily domains (GO:0002460) | 266 | 16 | 30.66 | - | 0.52 | 3.53E-03 | 3.91E-02 |
| establishment of protein localization to membrane (GO:0090150) | 226 | 13 | 26.05 | - | 0.5 | 4.46E-03 | 4.80E-02 |
| cellular catabolic process (GO:0044248) | 788 | 45 | 90.82 | - | 0.5 | 2.00E-08 | 8.24E-07 |
| small molecule catabolic process (GO:0044282) | 357 | 20 | 41.15 | - | 0.49 | 1.56E-04 | 2.86E-03 |
| protein targeting (GO:0006605) | 247 | 13 | 28.47 | - | 0.46 | 1.21E-03 | 1.64E-02 |
| carbohydrate derivative catabolic process (GO:1901136) | 222 | 11 | 25.59 | - | 0.43 | 9.80E-04 | 1.38E-02 |
| amino acid metabolic process (GO:0006520) | 288 | 14 | 33.19 | - | 0.42 | 1.22E-04 | 2.32E-03 |
| leukocyte mediated immunity (GO:0002443) | 313 | 15 | 36.08 | - | 0.42 | 5.10E-05 | 1.11E-03 |
| detection of stimulus (GO:0051606) | 675 | 32 | 77.8 | - | 0.41 | 6.25E-10 | 3.21E-08 |
| L-amino acid metabolic process (GO:0170033) | 175 | 8 | 20.17 | - | 0.4 | 1.89E-03 | 2.37E-02 |
| vacuole organization (GO:0007033) | 197 | 9 | 22.71 | - | 0.4 | 1.04E-03 | 1.44E-02 |
| proteinogenic amino acid metabolic process (GO:0170039) | 160 | 7 | 18.44 | - | 0.38 | 2.60E-03 | 3.06E-02 |
| defense response to bacterium (GO:0042742) | 302 | 13 | 34.81 | - | 0.37 | 1.60E-05 | 3.90E-04 |
| lymphocyte mediated immunity (GO:0002449) | 258 | 11 | 29.74 | - | 0.37 | 7.17E-05 | 1.48E-03 |

| | | | | | | | |
|---|---|---|---|---|---|---|---|
| pyridine-containing compound metabolic process (GO:0072524) | 125 | 5 | 14.41 | - | 0.35 | 4.70E-03 | 4.94E-02 |
| adaptive immune response (GO:0002250) | 677 | 27 | 78.03 | - | 0.35 | 2.55E-12 | 1.71E-10 |
| xenobiotic metabolic process (GO:0006805) | 126 | 5 | 14.52 | - | 0.34 | 4.72E-03 | 4.95E-02 |
| alpha-amino acid metabolic process (GO:1901605) | 209 | 8 | 24.09 | - | 0.33 | 1.18E-04 | 2.26E-03 |
| generation of precursor metabolites and energy (GO:0006091) | 373 | 13 | 42.99 | - | 0.3 | 2.86E-08 | 1.16E-06 |
| organic acid catabolic process (GO:0016054) | 231 | 8 | 26.62 | - | 0.3 | 1.70E-05 | 4.12E-04 |
| carboxylic acid catabolic process (GO:0046395) | 231 | 8 | 26.62 | - | 0.3 | 1.70E-05 | 4.11E-04 |
| monocarboxylic acid catabolic process (GO:0072329) | 116 | 4 | 13.37 | - | 0.3 | 3.23E-03 | 3.63E-02 |
| proton transmembrane transport (GO:1902600) | 146 | 5 | 16.83 | - | 0.3 | 9.64E-04 | 1.35E-02 |
| energy derivation by oxidation of organic compounds (GO:0015980) | 273 | 9 | 31.47 | - | 0.29 | 1.34E-06 | 4.00E-05 |
| lipoprotein metabolic process (GO:0042157) | 124 | 4 | 14.29 | - | 0.28 | 1.66E-03 | 2.15E-02 |
| mitochondrial respiratory chain complex assembly (GO:0033108) | 103 | 3 | 11.87 | - | 0.25 | 2.92E-03 | 3.40E-02 |
| regulation of T cell mediated immunity (GO:0002709) | 103 | 3 | 11.87 | - | 0.25 | 2.92E-03 | 3.39E-02 |
| detection of stimulus involved in sensory perception (GO:0050906) | 558 | 16 | 64.31 | - | 0.25 | 5.68E-14 | 4.45E-12 |
| lysosome organization (GO:0007040) | 105 | 3 | 12.1 | - | 0.25 | 3.04E-03 | 3.53E-02 |
| lytic vacuole organization (GO:0080171) | 105 | 3 | 12.1 | - | 0.25 | 3.04E-03 | 3.52E-02 |

| | | | | | | | |
|---|---|---|---|---|---|---|---|
| regulation of cell killing (GO:0031341) | 110 | 3 | 12.68 | - | 0.24 | 1.45E-03 | 1.91E-02 |
| immunoglobulin mediated immune response (GO:0016064) | 186 | 5 | 21.44 | - | 0.23 | 2.32E-05 | 5.49E-04 |
| B cell mediated immunity (GO:0019724) | 191 | 5 | 22.01 | - | 0.23 | 1.09E-05 | 2.76E-04 |
| cellular respiration (GO:0045333) | 194 | 5 | 22.36 | - | 0.22 | 7.65E-06 | 1.98E-04 |
| detection of chemical stimulus (GO:0009593) | 520 | 12 | 59.93 | - | 0.2 | 5.38E-15 | 4.42E-13 |
| regulation of intracellular pH (GO:0051453) | 90 | 2 | 10.37 | - | 0.19 | 2.45E-03 | 2.91E-02 |
| immunoglobulin production (GO:0002377) | 141 | 3 | 16.25 | - | 0.18 | 8.71E-05 | 1.75E-03 |
| regulation of cellular pH (GO:0030641) | 95 | 2 | 10.95 | - | 0.18 | 1.77E-03 | 2.24E-02 |
| production of molecular mediator of immune response (GO:0002440) | 147 | 3 | 16.94 | - | 0.18 | 4.02E-05 | 8.94E-04 |
| sensory perception of chemical stimulus (GO:0007606) | 542 | 11 | 62.47 | - | 0.18 | 9.83E-17 | 9.65E-15 |
| sensory perception of smell (GO:0007608) | 466 | 8 | 53.71 | - | 0.15 | 8.36E-16 | 7.35E-14 |
| respiratory electron transport chain (GO:0022904) | 117 | 2 | 13.49 | - | 0.15 | 1.23E-04 | 2.34E-03 |
| oxidative phosphorylation (GO:0006119) | 119 | 2 | 13.72 | - | 0.15 | 1.27E-04 | 2.38E-03 |
| detection of chemical stimulus involved in sensory perception (GO:0050907) | 488 | 8 | 56.25 | - | 0.14 | 8.58E-17 | 8.48E-15 |
| electron transport chain (GO:0022900) | 127 | 2 | 14.64 | - | 0.14 | 6.19E-05 | 1.30E-03 |
| positive regulation of leukocyte mediated cytotoxicity (GO:0001912) | 69 | 1 | 7.95 | - | 0.13 | 3.87E-03 | 4.26E-02 |

| | | | | | | | |
|---|---|---|---|---|---|---|---|
| positive regulation of cell killing (GO:0031343) | 76 | 1 | 8.76 | - | 0.11 | 1.78E-03 | 2.25E-02 |
| aerobic respiration (GO:0009060) | 163 | 2 | 18.79 | - | 0.11 | 8.40E-07 | 2.65E-05 |
| aerobic electron transport chain (GO:0019646) | 87 | 1 | 10.03 | - | 0.1 | 5.57E-04 | 8.55E-03 |
| detection of chemical stimulus involved in sensory perception of smell (GO:0050911) | 440 | 5 | 50.71 | - | 0.1 | 2.27E-17 | 2.31E-15 |
| mitochondrial ATP synthesis coupled electron transport (GO:0042775) | 92 | 1 | 10.6 | - | 0.09 | 2.49E-04 | 4.28E-03 |
| ATP synthesis coupled electron transport (GO:0042773) | 92 | 1 | 10.6 | - | 0.09 | 2.49E-04 | 4.28E-03 |
| autophagosome maturation (GO:0097352) | 51 | 0 | 5.88 | - | < 0.01 | 3.37E-03 | 3.76E-02 |

**Supplementary table 5: Ciliary genes differentially expressed in ciliary mutants**

**NPHP1_Diiferentially_expressed:**

| | logFC | logCPM | F | PValue | FDR | NPHP1 | gene |
|---|---|---|---|---|---|---|---|
| NOTCH1 | 7.002083 | 3.27269 | 27.66089 | 1.03E-05 | 0.00014 | Up | NOTCH1 |
| RFX4 | 6.316626 | 1.597428 | 21.62063 | 7.83E-05 | 0.000794 | Up | RFX4 |
| FGFR3 | 5.124479 | 1.800335 | 14.08994 | 0.000722 | 0.00512 | Up | FGFR3 |
| ZIC2 | 4.870054 | 5.205141 | 431.3302 | 9.87E-20 | 6.4E-17 | Up | ZIC2 |
| YAP1 | 4.645513 | 2.420976 | 27.17636 | 1.17E-05 | 0.000159 | Up | YAP1 |
| KCNJ10 | 4.508516 | -0.64113 | 10.54321 | 0.003316 | 0.017614 | Up | KCNJ10 |
| NEK2 | 4.273711 | -0.00681 | 8.450045 | 0.007217 | 0.032775 | Up | NEK2 |
| ALPK1 | 3.659693 | 0.515827 | 16.69543 | 0.000288 | 0.002411 | Up | ALPK1 |
| NEDD9 | 3.611086 | 2.33371 | 61.72024 | 7.31E-09 | 2.58E-07 | Up | NEDD9 |
| CENPF | 3.450419 | 3.607574 | 21.12136 | 6.84E-05 | 0.000707 | Up | CENPF |
| GLI3 | 2.834795 | 2.08058 | 20.98505 | 7.13E-05 | 0.000733 | Up | GLI3 |
| ADAMTS9 | 2.811958 | 2.482344 | 39.33623 | 5.71E-07 | 1.18E-05 | Up | ADAMTS9 |
| RAB34 | 2.710198 | 0.762692 | 8.87326 | 0.005803 | 0.027516 | Up | RAB34 |
| SMO | 2.099735 | 2.172323 | 11.61144 | 0.001836 | 0.010977 | Up | SMO |
| SMAD3 | 2.090896 | 3.537693 | 60.68618 | 8.72E-09 | 3.03E-07 | Up | SMAD3 |
| EZH2 | 1.883204 | 3.352984 | 32.21534 | 3.11E-06 | 5.01E-05 | Up | EZH2 |
| FSTL1 | 1.586042 | 3.624483 | 15.82113 | 0.00039 | 0.00308 | Up | FSTL1 |
| CRHR2 | 1.579355 | 1.207559 | 7.932398 | 0.008378 | 0.036783 | Up | CRHR2 |
| PLK1 | 1.543881 | 1.873905 | 9.865941 | 0.003691 | 0.019228 | Up | PLK1 |
| FLNA | 1.495666 | 6.858224 | 68.74036 | 2.33E-09 | 9.48E-08 | Up | FLNA |
| KCNF1 | 1.486561 | 2.180407 | 19.4833 | 0.000115 | 0.001098 | Up | KCNF1 |
| LIMA1 | 1.44052 | 3.151745 | 20.40574 | 8.55E-05 | 0.000853 | Up | LIMA1 |
| DNAH2 | 1.42007 | 1.266991 | 9.483394 | 0.004323 | 0.021758 | Up | DNAH2 |
| DNAH6 | 1.395123 | 2.679993 | 21.32389 | 6.42E-05 | 0.000671 | Up | DNAH6 |
| VANGL2 | 1.351146 | 7.137197 | 177.2657 | 2.37E-14 | 3.41E-12 | Up | VANGL2 |
| PRICKLE1 | 1.278427 | 5.500342 | 97.37683 | 4.47E-11 | 2.85E-09 | Up | PRICKLE1 |

| | | | | | | | |
|---|---|---|---|---|---|---|---|
| SYNE2 | 1.239287 | 4.698629 | 26.2864 | 1.5E-05 | 0.000195 | Up | SYNE2 |
| SALL1 | 1.226472 | 5.918038 | 96.24538 | 5.13E-11 | 3.22E-09 | Up | SALL1 |
| WDR90 | 1.19414 | 4.124898 | 25.52757 | 1.85E-05 | 0.000233 | Up | WDR90 |
| GPR161 | 1.153142 | 5.750879 | 59.38844 | 1.09E-08 | 3.67E-07 | Up | GPR161 |
| PTCH1 | 1.040107 | 4.532843 | 30.77157 | 4.49E-06 | 6.84E-05 | Up | PTCH1 |
| STK36 | 1.013957 | 4.445574 | 30.43281 | 4.91E-06 | 7.35E-05 | Up | STK36 |
| PKD2 | 0.930572 | 4.488882 | 32.26414 | 3.07E-06 | 4.96E-05 | Up | PKD2 |
| ZNF423 | 0.924123 | 6.285238 | 34.88397 | 1.61E-06 | 2.86E-05 | Up | ZNF423 |
| KIZ | 0.917979 | 4.211933 | 28.87643 | 7.39E-06 | 0.000105 | Up | KIZ |
| TTLL3 | 0.840619 | 3.289584 | 13.20456 | 0.001001 | 0.006686 | Up | TTLL3 |
| LIMK2 | 0.775557 | 6.059562 | 54.46959 | 2.63E-08 | 7.96E-07 | Up | LIMK2 |
| SUFU | 0.733639 | 4.431881 | 19.87403 | 0.000101 | 0.000987 | Up | SUFU |
| TUBB3 | 0.727889 | 3.215996 | 9.727329 | 0.003908 | 0.020099 | Up | TUBB3 |
| CFAP70 | 0.726257 | 2.938161 | 8.599702 | 0.006275 | 0.029255 | Up | CFAP70 |
| IGF1R | 0.719801 | 7.016766 | 45.10096 | 1.64E-07 | 4E-06 | Up | IGF1R |
| ANKS6 | 0.715986 | 3.888533 | 11.32383 | 0.002055 | 0.011999 | Up | ANKS6 |
| PKD1 | 0.709126 | 7.866929 | 14.5752 | 0.000606 | 0.004426 | Up | PKD1 |
| CROCC | 0.701792 | 4.632598 | 11.36635 | 0.002021 | 0.011846 | Up | CROCC |
| CELSR3 | 0.684897 | 7.022431 | 21.1047 | 6.87E-05 | 0.000709 | Up | CELSR3 |
| NUP93 | 0.68292 | 6.844128 | 49.57069 | 6.67E-08 | 1.82E-06 | Up | NUP93 |
| TUBE1 | 0.676805 | 4.928846 | 14.29285 | 0.000671 | 0.004823 | Up | TUBE1 |
| CCDC40 | 0.662619 | 4.560758 | 12.65996 | 0.001228 | 0.007911 | Up | CCDC40 |
| FAM161A | 0.652386 | 4.130534 | 13.5155 | 0.000891 | 0.006106 | Up | FAM161A |
| CENPJ | 0.643064 | 5.32017 | 23.53332 | 3.3E-05 | 0.000381 | Up | CENPJ |
| ODF2 | 0.636869 | 5.539953 | 27.78331 | 9.92E-06 | 0.000136 | Up | ODF2 |
| ONECUT1 | 0.62356 | 6.056979 | 16.7122 | 0.000286 | 0.0024 | Up | ONECUT1 |
| FNBP1L | 0.616222 | 8.411331 | 42.87747 | 2.62E-07 | 6E-06 | Up | FNBP1L |
| CEP350 | 0.600397 | 6.349488 | 26.6775 | 1.34E-05 | 0.000178 | Up | CEP350 |
| TUBGCP6 | 0.572928 | 6.34091 | 17.24895 | 0.000239 | 0.002052 | Up | TUBGCP6 |
| CEP78 | 0.555618 | 5.225073 | 18.501 | 0.000157 | 0.001446 | Up | CEP78 |

| | | | | | | | |
|---|---|---|---|---|---|---|---|
| PCNT | 0.55085 | 6.217907 | 19.84407 | 0.000102 | 0.000995 | Up | PCNT |
| CEP164 | 0.547415 | 5.519364 | 17.21735 | 0.000241 | 0.00207 | Up | CEP164 |
| KIAA1549 | 0.532494 | 7.124807 | 25.3316 | 1.96E-05 | 0.000245 | Up | KIAA1549 |
| RFX7 | 0.530164 | 6.26241 | 25.05264 | 2.12E-05 | 0.000261 | Up | RFX7 |
| IFT122 | 0.513554 | 4.987063 | 14.25725 | 0.000679 | 0.004876 | Up | IFT122 |
| KCTD10 | 0.512396 | 6.361699 | 22.15141 | 4.99E-05 | 0.000541 | Up | KCTD10 |
| TUBGCP2 | 0.495974 | 6.113132 | 12.34811 | 0.001382 | 0.008682 | Up | TUBGCP2 |
| IRS1 | 0.495908 | 6.550505 | 16.91984 | 0.000267 | 0.002262 | Up | IRS1 |
| ALMS1 | 0.49476 | 5.395662 | 13.24508 | 0.000986 | 0.006611 | Up | ALMS1 |
| VHL | 0.487658 | 6.749435 | 24.22767 | 2.69E-05 | 0.000321 | Up | VHL |
| FLCN | 0.481101 | 5.007791 | 9.68833 | 0.003971 | 0.02036 | Up | FLCN |
| IFT140 | 0.47387 | 5.831827 | 12.89828 | 0.001122 | 0.007326 | Up | IFT140 |
| CEP170 | 0.469001 | 8.908474 | 25.34023 | 1.96E-05 | 0.000244 | Up | CEP170 |
| SDCCAG8 | 0.45617 | 4.743467 | 7.612175 | 0.009647 | 0.041266 | Up | SDCCAG8 |
| ATAT1 | 0.440196 | 6.850168 | 17.70354 | 0.000205 | 0.001807 | Up | ATAT1 |
| TSC1 | 0.438478 | 6.811839 | 17.82042 | 0.000197 | 0.001749 | Up | TSC1 |
| IQCE | 0.43011 | 6.091825 | 8.651243 | 0.006138 | 0.028759 | Up | IQCE |
| POMK | 0.424555 | 6.522318 | 11.77056 | 0.001726 | 0.010451 | Up | POMK |
| NINL | 0.419115 | 5.314653 | 9.555627 | 0.004195 | 0.021283 | Up | NINL |
| KIAA0753 | 0.413466 | 4.982409 | 9.381019 | 0.004511 | 0.022496 | Up | KIAA0753 |
| EFHC1 | 0.399108 | 5.284514 | 7.242023 | 0.01138 | 0.046947 | Up | EFHC1 |
| CNGB1 | 0.395681 | 5.223128 | 7.237905 | 0.011401 | 0.047015 | Up | CNGB1 |
| GPR173 | 0.395083 | 6.305477 | 11.46407 | 0.001945 | 0.011466 | Up | GPR173 |
| TSC2 | 0.392077 | 7.183904 | 9.162401 | 0.004943 | 0.024164 | Up | TSC2 |
| TUBA1A | 0.389807 | 11.97996 | 15.95235 | 0.000372 | 0.002966 | Up | TUBA1A |
| CEP68 | 0.382805 | 5.684769 | 7.578137 | 0.009794 | 0.041748 | Up | CEP68 |
| CTNNB1 | 0.381439 | 8.582741 | 18.31921 | 0.000167 | 0.001522 | Up | CTNNB1 |
| CSPP1 | 0.362637 | 4.975161 | 7.746915 | 0.009089 | 0.0393 | Up | CSPP1 |
| CEP290 | 0.355078 | 6.089405 | 9.65454 | 0.004027 | 0.02057 | Up | CEP290 |
| IFT172 | 0.343706 | 5.783554 | 8.225485 | 0.007373 | 0.033332 | Up | IFT172 |

| | | | | | | | |
|---|---|---|---|---|---|---|---|
| TRRAP | 0.340669 | 6.93256 | 9.735719 | 0.003894 | 0.020055 | Up | TRRAP |
| VPS45 | 0.33692 | 5.993975 | 9.304197 | 0.004658 | 0.023074 | Up | VPS45 |
| TUBB2B | 0.312879 | 9.413041 | 8.708537 | 0.00599 | 0.028211 | Up | TUBB2B |
| MACF1 | 0.312154 | 9.124954 | 8.743981 | 0.0059 | 0.02787 | Up | MACF1 |
| SRGAP3 | 0.295781 | 7.69778 | 9.666493 | 0.004007 | 0.020504 | Up | SRGAP3 |
| VDAC3 | -0.26947 | 7.627275 | 8.17151 | 0.007548 | 0.033988 | Down | VDAC3 |
| TXNDC15 | -0.28495 | 6.437572 | 7.774738 | 0.008978 | 0.038944 | Down | TXNDC15 |
| HIF1A | -0.30004 | 6.985919 | 8.608316 | 0.006252 | 0.02918 | Down | HIF1A |
| ABLIM3 | -0.3181 | 6.911876 | 9.875736 | 0.003676 | 0.01917 | Down | ABLIM3 |
| GALNT11 | -0.34399 | 6.647011 | 11.053 | 0.002287 | 0.013119 | Down | GALNT11 |
| SNX17 | -0.35367 | 6.155697 | 10.97186 | 0.002362 | 0.013451 | Down | SNX17 |
| PRKACA | -0.35443 | 7.959171 | 13.50501 | 0.000895 | 0.006118 | Down | PRKACA |
| MAGI2 | -0.35782 | 6.614887 | 11.82612 | 0.001689 | 0.010261 | Down | MAGI2 |
| HTT | -0.35813 | 7.13253 | 11.18277 | 0.002172 | 0.012553 | Down | HTT |
| CUL7 | -0.3739 | 5.518359 | 10.56428 | 0.002779 | 0.015272 | Down | CUL7 |
| NUDC | -0.37652 | 6.374815 | 12.44542 | 0.001332 | 0.008433 | Down | NUDC |
| USP8 | -0.38689 | 6.023915 | 9.343251 | 0.004583 | 0.022775 | Down | USP8 |
| RAB11A | -0.38802 | 7.92137 | 14.73455 | 0.000572 | 0.004228 | Down | RAB11A |
| BBS4 | -0.39133 | 5.528563 | 12.40886 | 0.001351 | 0.008525 | Down | BBS4 |
| TTBK2 | -0.39619 | 7.517112 | 16.26985 | 0.000333 | 0.00272 | Down | TTBK2 |
| CDK20 | -0.41278 | 5.030824 | 10.25792 | 0.003145 | 0.016876 | Down | CDK20 |
| TUBB4B | -0.41817 | 7.400419 | 9.63756 | 0.004055 | 0.020679 | Down | TUBB4B |
| GPR83 | -0.42357 | 5.083282 | 9.086987 | 0.005103 | 0.02475 | Down | GPR83 |
| RAB23 | -0.42562 | 4.947558 | 8.178371 | 0.007525 | 0.033902 | Down | RAB23 |
| RCAN2 | -0.43621 | 8.338393 | 22.77091 | 4.14E-05 | 0.000465 | Down | RCAN2 |
| SNX10 | -0.43712 | 7.185969 | 12.53429 | 0.001288 | 0.008215 | Down | SNX10 |
| PRICKLE2 | -0.43878 | 7.015075 | 16.55633 | 0.000302 | 0.002512 | Down | PRICKLE2 |
| TMEM237 | -0.45433 | 5.429492 | 12.96251 | 0.001095 | 0.007174 | Down | TMEM237 |
| ATG3 | -0.46355 | 5.192869 | 11.85769 | 0.001669 | 0.010161 | Down | ATG3 |
| PIK3R4 | -0.47153 | 4.900067 | 9.825564 | 0.003753 | 0.019466 | Down | PIK3R4 |

| | | | | | | | |
|---|---|---|---|---|---|---|---|
| MLF1 | -0.47732 | 4.476462 | 7.286359 | 0.011156 | 0.04626 | Down | MLF1 |
| MAPK3 | -0.47947 | 5.787759 | 13.32868 | 0.000955 | 0.006453 | Down | MAPK3 |
| IFT57 | -0.48023 | 5.52716 | 15.6112 | 0.000419 | 0.003278 | Down | IFT57 |
| TUBGCP5 | -0.48319 | 5.252056 | 14.92407 | 0.000535 | 0.004006 | Down | TUBGCP5 |
| ATG5 | -0.50757 | 4.735654 | 11.09932 | 0.002245 | 0.012912 | Down | ATG5 |
| MAP1LC3A | -0.5506 | 6.641583 | 13.40119 | 0.00093 | 0.006318 | Down | MAP1LC3A |
| TBC1D7 | -0.64083 | 4.47013 | 18.14237 | 0.000177 | 0.001598 | Down | TBC1D7 |
| AHI1 | -0.64863 | 7.511954 | 34.09439 | 1.95E-06 | 3.35E-05 | Down | AHI1 |
| NEK10 | -0.72009 | 3.286683 | 9.113202 | 0.005047 | 0.024547 | Down | NEK10 |
| DISC1 | -0.73442 | 3.736934 | 10.74949 | 0.00258 | 0.014428 | Down | DISC1 |
| GPR22 | -0.73615 | 4.796821 | 19.91987 | 9.97E-05 | 0.000975 | Down | GPR22 |
| CEP19 | -0.77281 | 4.954546 | 32.27542 | 3.06E-06 | 4.95E-05 | Down | CEP19 |
| LZTFL1 | -0.80487 | 4.591056 | 26.38722 | 1.46E-05 | 0.000191 | Down | LZTFL1 |
| PARD6A | -0.80667 | 3.564395 | 10.68239 | 0.002651 | 0.014717 | Down | PARD6A |
| SNAP25 | -0.8304 | 8.555284 | 79.88094 | 4.42E-10 | 2.18E-08 | Down | SNAP25 |
| PDE6D | -0.84685 | 5.676413 | 47.5003 | 1.01E-07 | 2.61E-06 | Down | PDE6D |
| CEP83 | -0.84878 | 2.962219 | 10.49693 | 0.002856 | 0.015592 | Down | CEP83 |
| CC2D2A | -0.86509 | 4.357617 | 22.49167 | 4.5E-05 | 0.000498 | Down | CC2D2A |
| PACRG | -0.90009 | 3.255101 | 13.00109 | 0.00108 | 0.007089 | Down | PACRG |
| RAB28 | -0.90786 | 3.569399 | 14.43074 | 0.000638 | 0.004629 | Down | RAB28 |
| PRPH2 | -1.02247 | 3.179464 | 22.17538 | 4.95E-05 | 0.000538 | Down | PRPH2 |
| DRD2 | -1.0439 | 5.196893 | 56.41303 | 1.85E-08 | 5.81E-07 | Down | DRD2 |
| TMEM107 | -1.04705 | 3.36659 | 22.28899 | 4.78E-05 | 0.000523 | Down | TMEM107 |
| ARL6 | -1.0839 | 4.885161 | 47.68629 | 9.69E-08 | 2.54E-06 | Down | ARL6 |
| JHY | -1.09603 | 3.641016 | 27.54048 | 1.06E-05 | 0.000145 | Down | JHY |
| MCHR1 | -1.12035 | 4.767294 | 51.61273 | 4.5E-08 | 1.28E-06 | Down | MCHR1 |
| USH2A | -1.19056 | 1.995709 | 11.68593 | 0.001784 | 0.010725 | Down | USH2A |
| RSPH1 | -1.19576 | 1.65299 | 7.808091 | 0.008847 | 0.038456 | Down | RSPH1 |
| HAP1 | -1.32459 | 5.879163 | 95.13612 | 5.88E-11 | 3.66E-09 | Down | HAP1 |
| KIF17 | -1.34699 | 3.188803 | 32.21067 | 3.11E-06 | 5.01E-05 | Down | KIF17 |

| | | | | | | | |
|---|---|---|---|---|---|---|---|
| TGFB1 | -1.36246 | 3.195538 | 24.74343 | 2.32E-05 | 0.000282 | Down | TGFB1 |
| PTGER4 | -1.51636 | 1.535657 | 11.54914 | 0.001881 | 0.011199 | Down | PTGER4 |
| EVC | -1.52296 | 3.224598 | 36.26101 | 1.16E-06 | 2.16E-05 | Down | EVC |
| AK7 | -1.6255 | 1.297064 | 12.20173 | 0.001462 | 0.009105 | Down | AK7 |
| CFAP47 | -1.74349 | 0.845595 | 8.865438 | 0.005603 | 0.026742 | Down | CFAP47 |
| HTR6 | -2.07359 | 3.251247 | 76.67105 | 7.01E-10 | 3.3E-08 | Down | HTR6 |
| LRRC34 | -2.15291 | 0.168123 | 7.841511 | 0.008718 | 0.038045 | Down | LRRC34 |
| TUBA4A | -2.43287 | 4.956097 | 247.0995 | 2.64E-16 | 6.52E-14 | Down | TUBA4A |
| ORC1 | -2.49397 | 1.249769 | 31.05872 | 4.17E-06 | 6.42E-05 | Down | ORC1 |
| NGFR | -3.12922 | 7.260694 | 458.3551 | 4.09E-20 | 3.26E-17 | Down | NGFR |
| DRD5 | -3.39882 | -0.50525 | 10.27575 | 0.003275 | 0.017444 | Down | DRD5 |
| DAW1 | -3.51128 | -0.13025 | 11.19757 | 0.002423 | 0.013756 | Down | DAW1 |
| DRD1 | -3.67407 | 0.462026 | 30.46806 | 4.86E-06 | 7.29E-05 | Down | DRD1 |
| ARHGAP36 | -4.50359 | 3.19445 | 131.693 | 1.11E-12 | 1.05E-10 | Down | ARHGAP36 |
| GMNC | -4.69015 | 0.227193 | 26.24466 | 1.81E-05 | 0.000229 | Down | GMNC |
| CNGA1 | -4.79372 | -0.45775 | 11.08947 | 0.002701 | 0.014943 | Down | CNGA1 |
| PCARE | -4.87494 | -0.33968 | 16.48018 | 0.000487 | 0.003714 | Down | PCARE |

**CEP290_Differentially_expressed:**

| | logFC | logCPM | F | PValue | FDR | CEP290 | gene |
|---|---|---|---|---|---|---|---|
| RFX4 | 7.341121 | 1.597428 | 30.30036 | 7.92E-06 | 0.000218 | Up | RFX4 |
| NOTCH1 | 6.182864 | 3.27269 | 13.69031 | 0.000836 | 0.009884 | Up | NOTCH1 |
| FGFR3 | 5.609851 | 1.800335 | 14.78831 | 0.000561 | 0.007274 | Up | FGFR3 |
| YAP1 | 4.845513 | 2.420976 | 24.04506 | 2.84E-05 | 0.000635 | Up | YAP1 |
| ZIC2 | 4.628374 | 5.205141 | 334.2901 | 3.82E-18 | 3.96E-15 | Up | ZIC2 |
| CAV1 | 4.086288 | 0.188693 | 14.51255 | 0.000619 | 0.00783 | Up | CAV1 |
| CENPF | 4.032999 | 3.607574 | 25.16834 | 2.05E-05 | 0.000483 | Up | CENPF |
| NEDD9 | 4.008665 | 2.33371 | 65.20234 | 4.1E-09 | 3.06E-07 | Up | NEDD9 |
| FSTL1 | 2.638385 | 3.624483 | 40.48838 | 4.42E-07 | 1.84E-05 | Up | FSTL1 |
| SMO | 2.450345 | 2.172323 | 12.48467 | 0.001312 | 0.014074 | Up | SMO |
| ADAMTS9 | 2.399768 | 2.482344 | 19.98867 | 9.75E-05 | 0.001764 | Up | ADAMTS9 |
| CRHR2 | 1.881846 | 1.207559 | 8.83207 | 0.005683 | 0.043354 | Up | CRHR2 |
| SMAD3 | 1.726289 | 3.537693 | 30.38239 | 4.97E-06 | 0.000147 | Up | SMAD3 |
| EZH2 | 1.70095 | 3.352984 | 19.27808 | 0.000122 | 0.002121 | Up | EZH2 |
| DNAH6 | 1.550468 | 2.679993 | 20.57395 | 8.11E-05 | 0.001526 | Up | DNAH6 |
| KCNF1 | 1.506853 | 2.180407 | 14.92772 | 0.000534 | 0.006981 | Up | KCNF1 |
| LIMA1 | 1.385894 | 3.151745 | 14.04817 | 0.000733 | 0.008897 | Up | LIMA1 |
| TUBA1C | 1.111038 | 4.097115 | 18.75837 | 0.000145 | 0.002453 | Up | TUBA1C |
| FLNA | 0.954917 | 6.858224 | 24.74732 | 2.32E-05 | 0.000536 | Up | FLNA |
| GPR161 | 0.876815 | 5.750879 | 27.8384 | 9.77E-06 | 0.000261 | Up | GPR161 |
| NUP93 | 0.857404 | 6.844128 | 71.4322 | 1.53E-09 | 1.31E-07 | Up | NUP93 |
| PTCH1 | 0.847436 | 4.532843 | 15.46965 | 0.000441 | 0.005961 | Up | PTCH1 |
| ZNF423 | 0.809292 | 6.285238 | 23.12764 | 3.72E-05 | 0.000792 | Up | ZNF423 |
| PACSIN1 | 0.732295 | 4.301168 | 13.58978 | 0.000867 | 0.010146 | Up | PACSIN1 |
| VANGL2 | 0.675182 | 7.137197 | 39.42205 | 5.6E-07 | 2.28E-05 | Up | VANGL2 |
| CEP78 | 0.671588 | 5.225073 | 21.94063 | 5.32E-05 | 0.001085 | Up | CEP78 |
| STK36 | 0.635047 | 4.445574 | 8.747402 | 0.005892 | 0.044483 | Up | STK36 |

| | | | | | | | |
|---|---|---|---|---|---|---|---|
| IGF1R | 0.599866 | 7.016766 | 28.26828 | 8.7E-06 | 0.000235 | Up | IGF1R |
| CCDC66 | 0.547725 | 4.548384 | 8.678619 | 0.006067 | 0.045492 | Up | CCDC66 |
| FNBP1L | 0.546837 | 8.411331 | 32.35191 | 3.01E-06 | 9.54E-05 | Up | FNBP1L |
| IFT172 | 0.485831 | 5.783554 | 13.84478 | 0.00079 | 0.009446 | Up | IFT172 |
| CPLANE1 | 0.478229 | 6.429032 | 14.33654 | 0.00066 | 0.008204 | Up | CPLANE1 |
| LIMK2 | 0.472924 | 6.059562 | 16.34684 | 0.000325 | 0.004657 | Up | LIMK2 |
| KIAA1549 | 0.42101 | 7.124807 | 14.37645 | 0.000651 | 0.008113 | Up | KIAA1549 |
| CEP350 | 0.411739 | 6.349488 | 10.68944 | 0.002643 | 0.024349 | Up | CEP350 |
| PCNT | 0.402752 | 6.217907 | 9.026594 | 0.005234 | 0.040687 | Up | PCNT |
| POMK | 0.383849 | 6.522318 | 8.406194 | 0.006819 | 0.049555 | Up | POMK |
| MACF1 | 0.366894 | 9.124954 | 11.76889 | 0.001727 | 0.017488 | Up | MACF1 |
| VHL | 0.355889 | 6.749435 | 11.33683 | 0.002045 | 0.020083 | Up | VHL |
| TUBB2B | 0.313169 | 9.413041 | 8.547194 | 0.006418 | 0.047455 | Up | TUBB2B |
| TNPO1 | 0.298457 | 8.502184 | 10.09337 | 0.003363 | 0.029314 | Up | TNPO1 |
| SRGAP3 | 0.297944 | 7.69778 | 9.159164 | 0.00495 | 0.039079 | Up | SRGAP3 |
| ABLIM1 | 0.293312 | 8.283591 | 8.813031 | 0.00573 | 0.043625 | Up | ABLIM1 |
| TXNDC15 | -0.33844 | 6.437572 | 9.405515 | 0.004465 | 0.036258 | Down | TXNDC15 |
| MAGI2 | -0.36271 | 6.614887 | 10.55334 | 0.002791 | 0.025388 | Down | MAGI2 |
| ABLIM3 | -0.37235 | 6.911876 | 11.96665 | 0.0016 | 0.016514 | Down | ABLIM3 |
| VCP | -0.39134 | 8.302272 | 15.74821 | 0.0004 | 0.005533 | Down | VCP |
| BBS4 | -0.40171 | 5.528563 | 10.15149 | 0.003284 | 0.028895 | Down | BBS4 |
| TUBB4B | -0.41757 | 7.400419 | 8.846447 | 0.005649 | 0.043199 | Down | TUBB4B |
| SNAP25 | -0.44014 | 8.555284 | 21.92388 | 5.34E-05 | 0.001089 | Down | SNAP25 |
| TRAPPC9 | -0.4473 | 6.420271 | 14.88101 | 0.000543 | 0.007085 | Down | TRAPPC9 |
| MAPK3 | -0.44954 | 5.787759 | 9.475556 | 0.004337 | 0.035424 | Down | MAPK3 |
| CDK20 | -0.489 | 5.030824 | 10.69976 | 0.002632 | 0.024276 | Down | CDK20 |
| VPS4A | -0.49166 | 5.300549 | 10.68585 | 0.002647 | 0.024373 | Down | VPS4A |
| FSD1 | -0.50577 | 6.577939 | 12.14016 | 0.001497 | 0.015681 | Down | FSD1 |
| YIF1B | -0.51623 | 4.940148 | 8.858014 | 0.005621 | 0.04305 | Down | YIF1B |
| CEP290 | -0.54396 | 6.089405 | 16.92753 | 0.000266 | 0.003977 | Down | CEP290 |

| | | | | | | | |
|---|---|---|---|---|---|---|---|
| PRICKLE2 | -0.58914 | 7.015075 | 26.21309 | 1.53E-05 | 0.00038 | Down | PRICKLE2 |
| MAP1LC3A | -0.60965 | 6.641583 | 14.2713 | 0.000676 | 0.008336 | Down | MAP1LC3A |
| EHD3 | -0.63174 | 5.830855 | 20.29774 | 8.84E-05 | 0.001637 | Down | EHD3 |
| HAP1 | -0.63917 | 5.879163 | 19.94688 | 9.88E-05 | 0.001779 | Down | HAP1 |
| PDE6D | -0.64258 | 5.676413 | 22.47348 | 4.53E-05 | 0.000944 | Down | PDE6D |
| XPNPEP3 | -0.65371 | 4.141607 | 10.57124 | 0.002771 | 0.025262 | Down | XPNPEP3 |
| ARL6 | -0.66528 | 4.885161 | 14.34741 | 0.000658 | 0.008179 | Down | ARL6 |
| AHI1 | -0.68234 | 7.511954 | 34.7256 | 1.67E-06 | 5.83E-05 | Down | AHI1 |
| CUL7 | -0.6844 | 5.518359 | 26.43519 | 1.44E-05 | 0.000359 | Down | CUL7 |
| GPR22 | -0.69295 | 4.796821 | 13.12322 | 0.001031 | 0.011656 | Down | GPR22 |
| HYDIN | -0.7072 | 4.071532 | 8.709263 | 0.005988 | 0.045064 | Down | HYDIN |
| CEP19 | -0.72883 | 4.954546 | 21.47494 | 6.13E-05 | 0.001218 | Down | CEP19 |
| CC2D2A | -0.73438 | 4.357617 | 11.70009 | 0.001774 | 0.017857 | Down | CC2D2A |
| ARL2 | -0.74167 | 4.498717 | 9.743497 | 0.003882 | 0.032603 | Down | ARL2 |
| PRPH2 | -0.7441 | 3.179464 | 8.661185 | 0.006112 | 0.045767 | Down | PRPH2 |
| DNAH1 | -0.79538 | 3.941162 | 11.88018 | 0.001654 | 0.01694 | Down | DNAH1 |
| RCAN2 | -0.96963 | 8.338393 | 104.1032 | 2.01E-11 | 3.05E-09 | Down | RCAN2 |
| MCHR1 | -0.99715 | 4.767294 | 30.60446 | 4.69E-06 | 0.00014 | Down | MCHR1 |
| NME3 | -1.0088 | 3.678997 | 9.2163 | 0.004833 | 0.038431 | Down | NME3 |
| DRD2 | -1.33647 | 5.196893 | 63.60189 | 5.33E-09 | 3.82E-07 | Down | DRD2 |
| TGFB1 | -1.33662 | 3.195538 | 16.24944 | 0.000336 | 0.004783 | Down | TGFB1 |
| SLC9A3R1 | -1.34066 | 3.164973 | 15.40702 | 0.00045 | 0.00607 | Down | SLC9A3R1 |
| DISC1 | -1.49044 | 3.736934 | 27.29992 | 1.13E-05 | 0.000295 | Down | DISC1 |
| TTC29 | -1.66698 | 2.102947 | 10.5871 | 0.002754 | 0.025157 | Down | TTC29 |
| TUBA4A | -1.89672 | 4.956097 | 120.162 | 3.48E-12 | 6.58E-10 | Down | TUBA4A |
| RSPH9 | -1.98269 | 1.185654 | 10.76014 | 0.002569 | 0.02386 | Down | RSPH9 |
| HTR6 | -2.12378 | 3.251247 | 51.73708 | 4.39E-08 | 2.39E-06 | Down | HTR6 |
| NGFR | -2.65851 | 7.260694 | 303.9979 | 1.46E-17 | 1.32E-14 | Down | NGFR |
| DRD1 | -2.79603 | 0.462026 | 12.6311 | 0.001241 | 0.013473 | Down | DRD1 |
| MYO7A | -2.86886 | 2.182527 | 16.00336 | 0.000366 | 0.005119 | Down | MYO7A |

| | | | | | | | |
|---|---|---|---|---|---|---|---|
| ORC1 | -3.03433 | 1.249769 | 23.38661 | 3.45E-05 | 0.000742 | Down | ORC1 |
| ARHGAP36 | -3.31109 | 3.19445 | 56.52784 | 1.81E-08 | 1.12E-06 | Down | ARHGAP36 |
| GMNC | -6.13502 | 0.227193 | 15.11427 | 0.000543 | 0.007088 | Down | GMNC |

Supplementary Figures and legends:

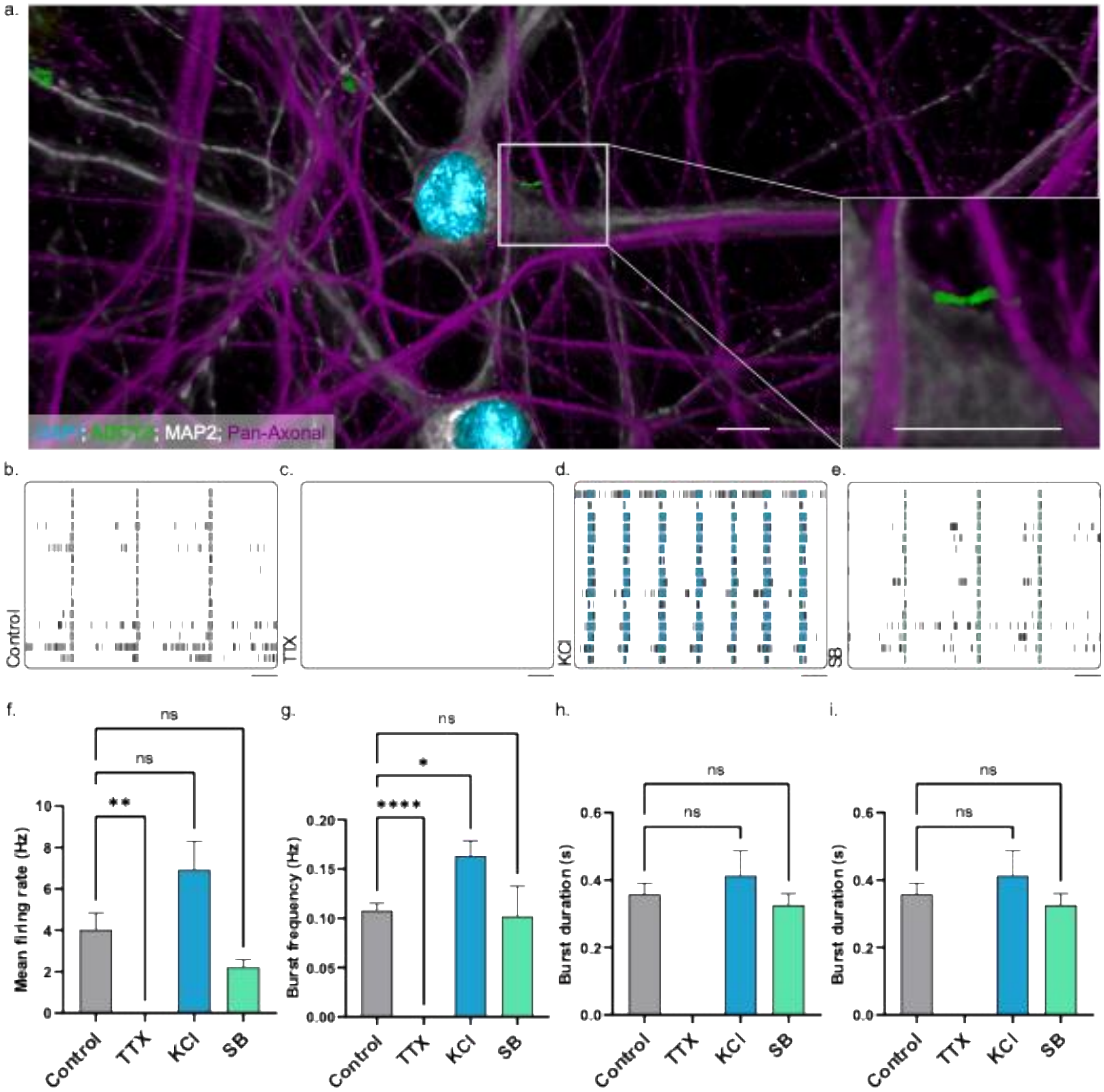


**Supplementary Figure 1: Effects of compounds on neuronal activity. (a)** 3D reconstruction of a primary cilium (ADCY3, green) making contacts with an axon (Pan-axonal, purple). Image acquired on Leica LSM900 with Airyscan 2.0, and reconstruction was made using Imaris software (Oxford Instruments, UK). Scale bar = 10 microns. **(b-e)** Representative raster plots of the activity of day 35 hiPSC-derived neurons following compound addition, 60s shown per plot and network bursts are highlighted. **(b)** shows the basal activity of neurons treated with vehicle (ddH$_2$O) for 24 hours, **(c)** shows recording following 24 hours of 1 μM TTX application **(d)** shows recording following 3 hours of 10 μM KCl **(e)** shows the recording following 24 hours of 1 μM SB-399225 application. Scale bar = 10 s. **(f-i)** show the parameters from these recordings (N=2 independent experiments and n=6 wells per condition. Data presented as mean ± SEM. Significance denoted following one-way ANOVA followed by Dunnett's multiple comparisons, ns = not significant,*p < 0.05, **p < 0.01, ****p < 0.0001).

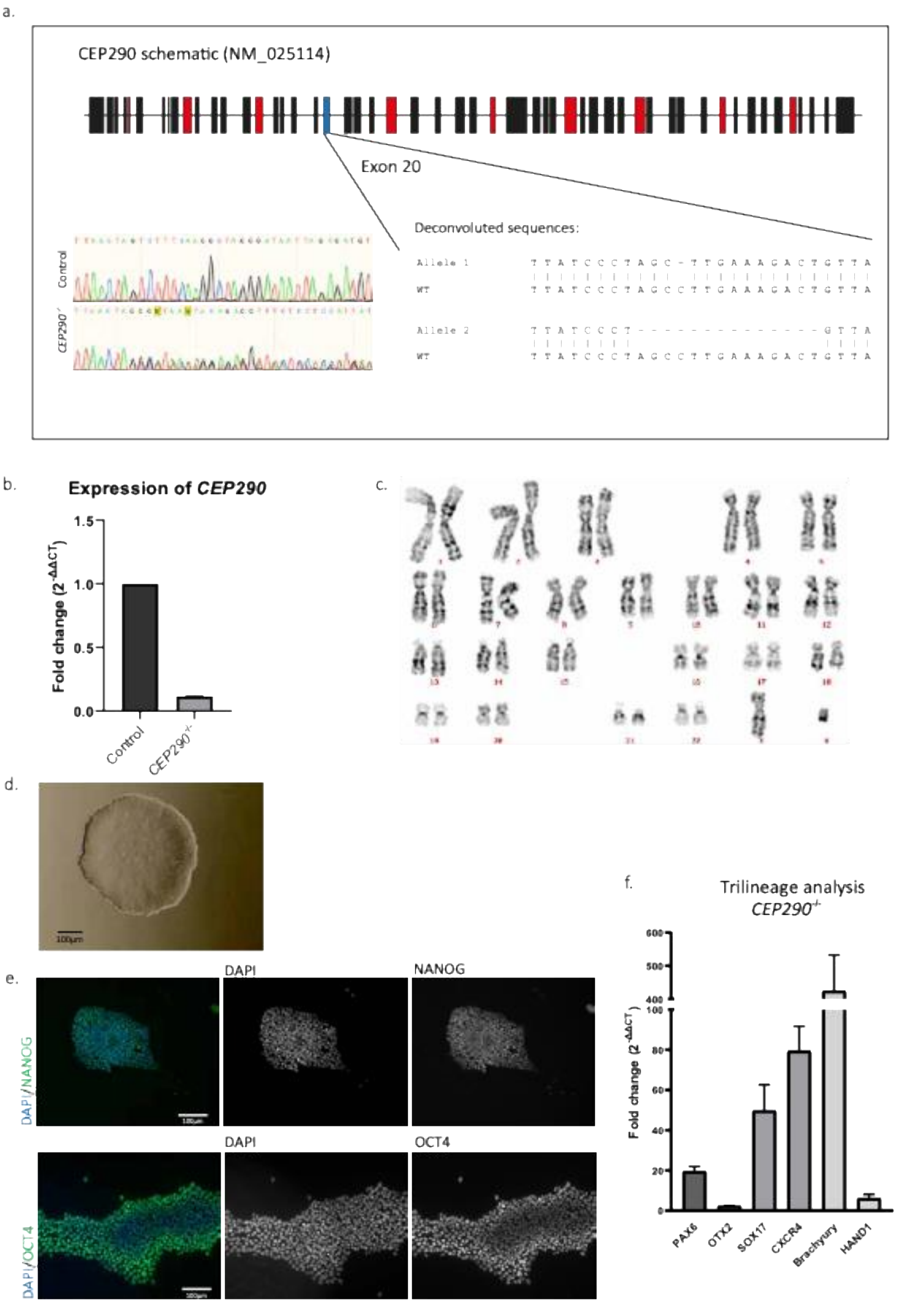


**Supplementary Figure 2: Generation of induced pluripotent stem cell line carrying frameshift variants in CEP290. (a)** Schematic of CEP290 gene showing location of mutations with chromatogram from Sanger sequencing of control and CEP290-/- lines, and sequence deconvolution showing precise nature of mutation. **(b)** Expression of CEP290 assessed by RT-qPCR (average expression data based on 3 biological replicates and 6 technical replicates using two sets of primers).

**(c)** Karyotype of resultant cell line acquired with resolution of 300-500 bhps. **(d)** Brightfield image of an hiPSC colony showing a characteristic appearance. **(e)** Immunofluorescent images of colonies to assess expression of pluripotency markers NANOG (Upper panel) and OCT4 (lower panel). Pluripotency markers shown in green and DAPI in blue. **(f)** RT-qPCR results following trilineage of cells. Results show the expression of genes for ectoderm (PAX6 and OTX2), mesoderm (BRACHYURY and HAND1), and endoderm (SOX17 and CXCR4) with respect to undifferentiated hiPSCs (n=2 biological replicates across 8x technical replicates).

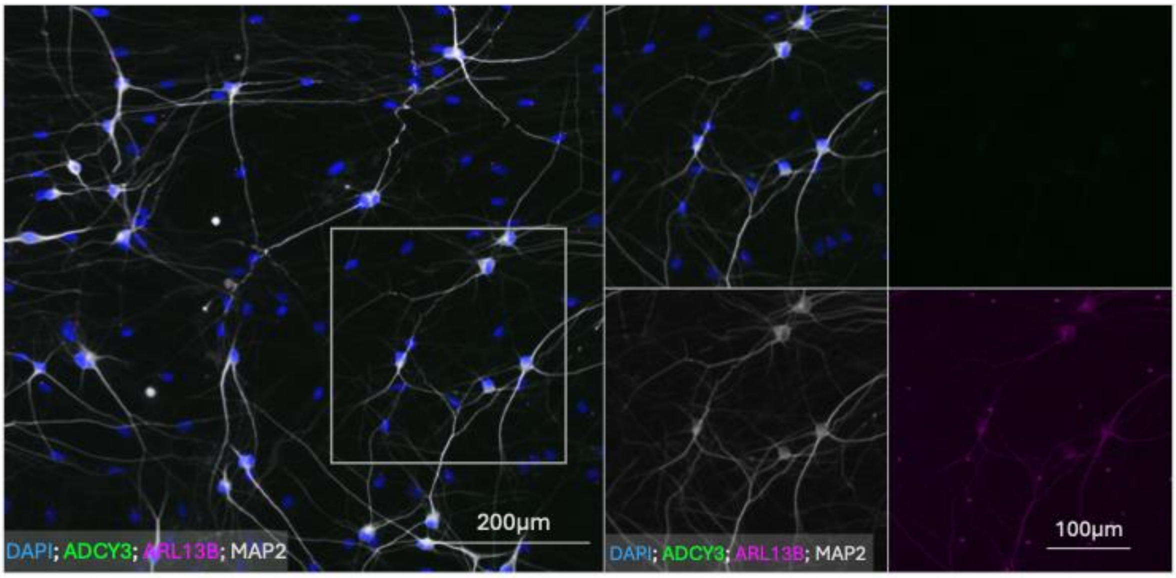


**Supplementary Figure 3: *CEP290*$^{-/-}$ neurons lack a primary cilium.** Immunofluorescent images of day 28 neurons stained against ADCY3 (green), ARL13B (magenta), and MAP2 (white) showing no cilia associated with *CEP290*$^{-/-}$ neurons, but ARL13B cilia associated with non-MAP2 positive cells (control astrocytes). Scale bar 200 µm, inset scale bar 100 µm.

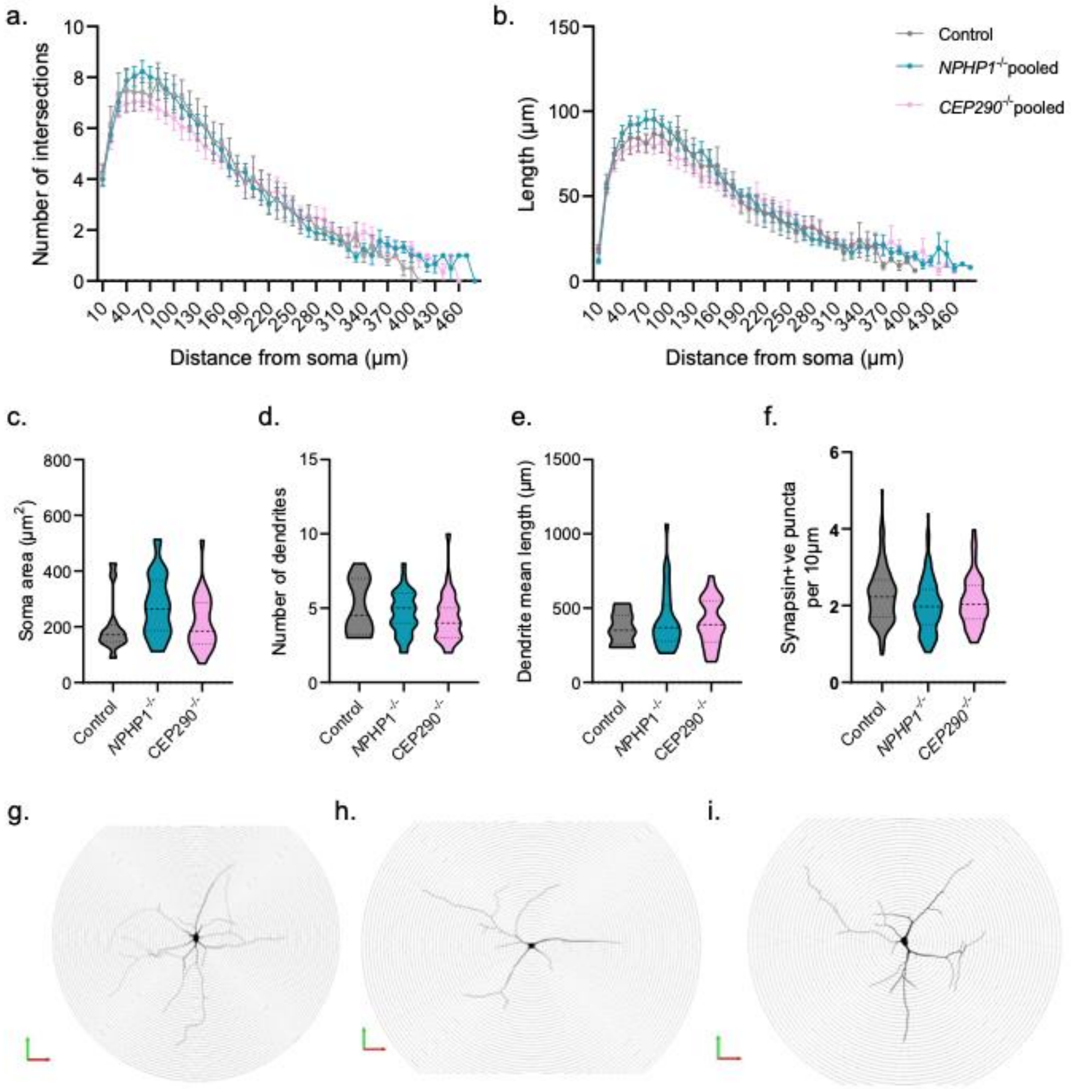


**Supplementary Figure 4: Neuronal morphology and quantitative analysis**
Quantifications of reconstructions of 28-day old neurons using MAP2 as the marker for reconstruction. **(a-b)** output of Sholl analysis of dendritic complexity showing the number of intersections or dendritic length as a function of radial distance from the soma. **(c-e)** Quantification of soma area, number of dendrites, and dendrite mean length from reconstructed neurons. **(f)** Density of synapsin-positive puncta per 100 µm dendrite length. **(g-i)** Representative neuronal reconstructions from control (g), *NPHP1*$^{-/-}$ (h), and *CEP290*$^{-/-}$ neurons at day 28. Grid spacing = 10 µm. Data are presented as mean ± SEM (a-b) or violin plots with median (dotted line) indicated (c-f); n=28-31 neurons across N=2 independent experiments. Data showed no significant differences following one-way ANOVA analysis followed by Dunnett's multiple comparisons ($p > 0.05$).

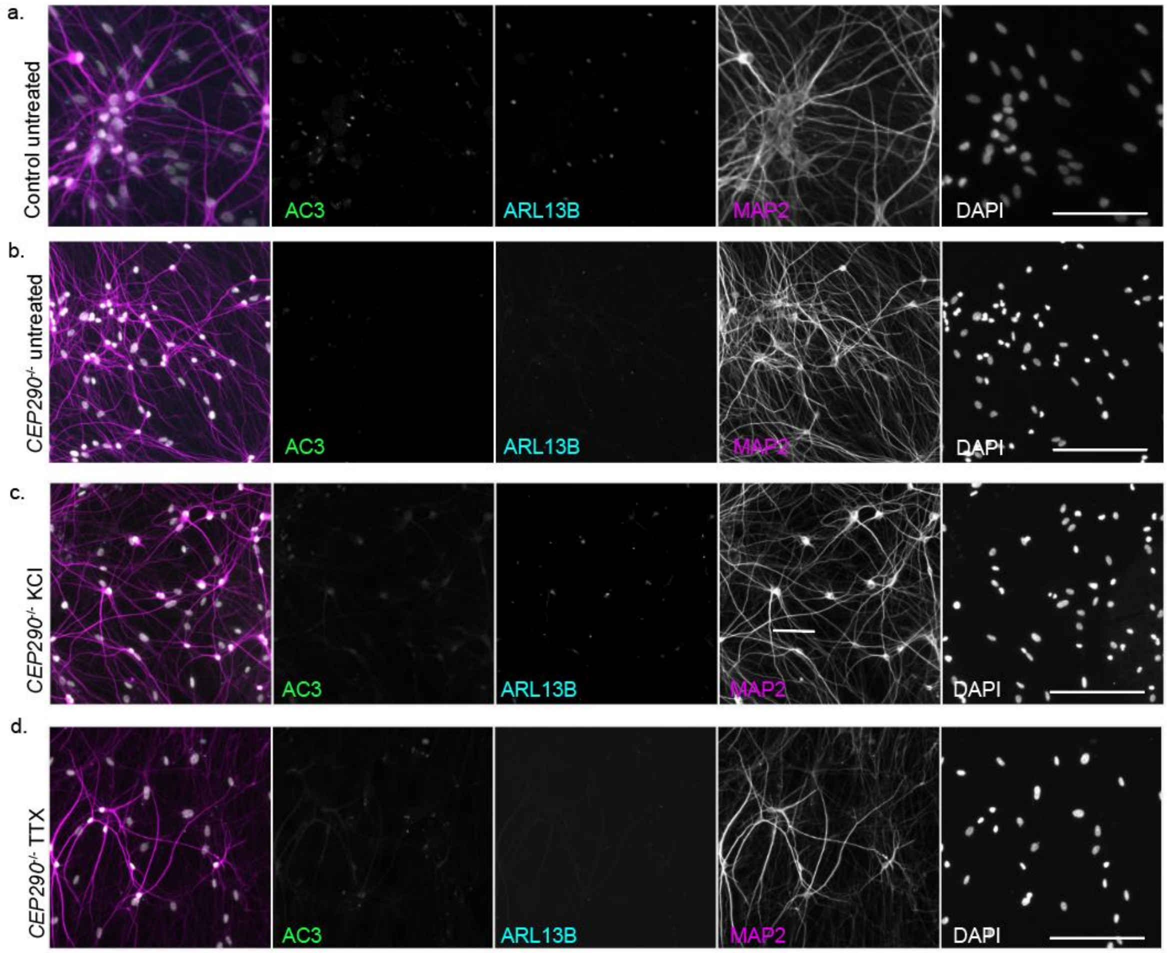


**Supplementary Figure 5: *CEP290*$^{-/-}$ neurons remain unciliated following treatment with compounds that alter activity**

Representative immunofluorescence images of day 35 neurons. Composite images depict ADCY3 in green, and MAP2 in magenta, ARL13B in cyan, and DAPI in white. **(a)** Control, untreated neurons, **(b)** *CEP290*$^{-/-}$ untreated neurons, **(c)** *CEP290*$^{-/-}$ neurons treated with 10mM KCl for 3h, and **(d)** *CEP290*$^{-/-}$ neurons treated with 1 µM TTX for 24 hours. Scale bar: 200 µm.

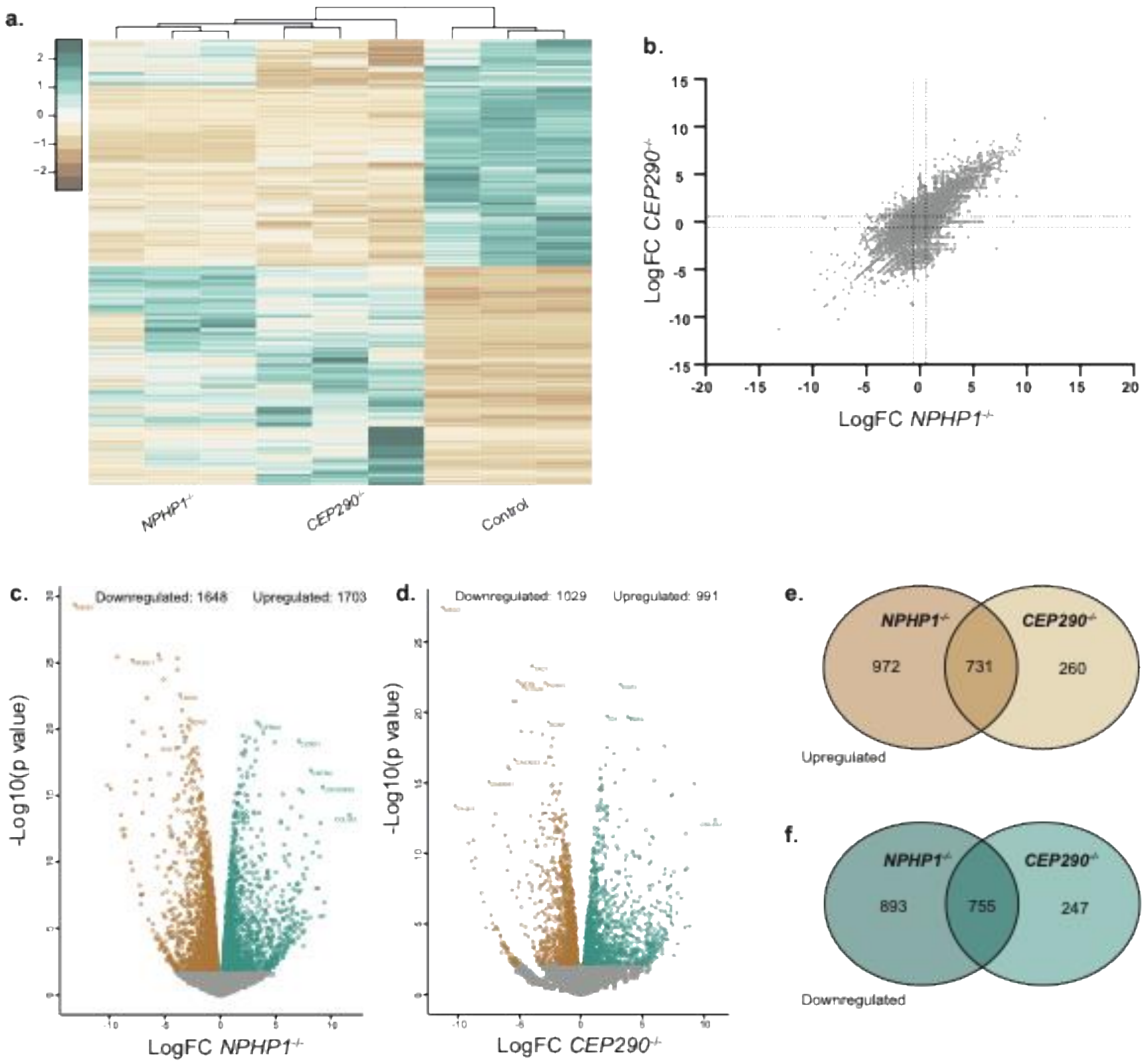


**Supplementary Figure 6: Comparison of RNAseq data between control, *NPHP1*$^{-/-}$ and *CEP290*$^{-/-}$ neurons.**

**(a)** Heatmap of differentially expressed genes (DEGs) visualised with heatmap2 tool which utilises the heatmap.2 function from the R gplots package, depicting hierarchical clustering of samples from *NPHP1*$^{-/-}$ and *CEP290*$^{-/-}$ 35-day old neurons. Gene expression values are shown as row Z-scores of normalised counts. **(b)** Scatter plot comparing $\log_2$ fold change (LogFC) values in *NPHP1*$^{-/-}$ versus *CEP290*$^{-/-}$ neurons, with dotted lines indicating ±1.5 LogFC and $p < 0.05$ thresholds. **(c, d)** Volcano plots of DEGs in *NPHP1*$^{-/-}$ **(c)** and *CEP290*$^{-/-}$ **(d)** neurons relative to control, highlighting significantly upregulated (green) and downregulated (brown) genes. Numbers of upregulated and downregulated genes are indicated above each plot. **(e, f)** Venn diagrams showing overlap of upregulated **(e)** and downregulated **(f)** DEGs between *NPHP1*$^{-/-}$ and *CEP290*$^{-/-}$ neurons.
